# Cross-link scrambling in peptide pairs

**DOI:** 10.1101/2022.12.27.522012

**Authors:** Luitzen de Jong, Winfried Roseboom, Gertjan Kramer

**Affiliations:** University of Amsterdam

## Abstract

Identification of peptides and their linked amino acids from chemically cross-linked protein complexes with bifunctional N-hydroxysuccinimidyl (NHS) esters can reveal interacting proteins and their spatial arrangements. With NHS esters both amide- and ester cross-links can be formed. Retention time prediction for strong cation exchange chromatography (SCXC) of cross-linked peptides at pH 3 can distinguish between ester and amide cross-links based on their charge differences. By this approach we show that about 98 % of cross-links are formed by two amide bonds. However MS/MS analysis revealed the presence of an ester linkage in more than 5% of peptide pairs predicted by SCXC to be linked by amide bonds. This discrepancy can be explained by intra-peptide amide-ester rearrangement in the gas phase during MS/MS analysis. So, SCXC retention time prediction can be used to distinguish amide-amide, amide-ester and ester-ester linkages actually formed in the cross-linking reaction and to detect scrambling of cross-linked sites. This information is important for studies aimed to understand the spatial arrangement of protein complexes by cross-linking at the highest possible resolution.

## Introduction

Chemical protein cross-linking (CX) combined with mass spectrometric (MS) identification of proteolytically obtained peptides can reveal the spatial arrangements of proteins in biomolecular assemblies. Since spatial distance constraints between linked residues are defined by the length of the spacer of the cross-linking agent, CXMS has been used widely to study the 3-D structure of isolated protein complexes, often in combination with molecular modeling and cryo-electron microscopy[1–3]. Interestingly, started by pioneering work several years ago [4], CX-MS can also be applied to detect binary protein-proteins interactions (PPIs) in living cells [5,6]. The attractiveness of this approach is the possibility to obtain a proteome-wide view of both stable and dynamic proteinprotein interactions in a single experiment and at an extremely low experimental false discovery rate [7]. Moreover, if the structures of the two interacting proteins are known, a 3-D model of the complex in vivo can be obtained, in particular if two or more inter-protein cross-links (inter-links) have been detected. Structural information may give insight in the functional significance of an interaction and may help in drug design.

Bifunctional NHS esters are often used for CXMS. NHS esters react predominantly with amine groups in proteins at the N-terminus and at the side chain of lysine residues under formation of amide bonds. To a lesser extent NHS esters also react with the hydroxyl groups of serine, threonine and tyrosine under formation of an ester bond [8,9]. In a large dataset it was assessed that S, T or Y are involved in about 14% of the linkages. Also protein N-termini were found to be involved in cross-linking[10]. Cross-link identification in complex samples with searching in entire species-specific sequence databases has a large search space when all possible combinations of linked amino acids are taken into account. With complex samples like in vivo cross-linked material this circumstantiality requires high stringent criteria for low false discovery rate of protein-protein interactions and leads to relatively long calculation times for data analysis. On the other hand ignoring reactivity of the cross-linker with S, T and Y may diminish the number of identified cross-links.

Both modification at all four susceptible amino acids K, S, T and Y, e.g., [11,12] and at K only, e.g., [13–16] have been allowed in data analysis of cross-linked cell extracts or in vivo cross-linked material and it has been assumed that cross-linking leads to stable products of which the linked residues can be assigned unambiguously by LC-MS/MS analysis if one or more fragment ions have been generated that enable distinction between different potential cross-link sites. However, it has never been studied to which extent cross-link sites can be rearranged at any stage during analysis, in particular in the gas phase during collision induced fragmentation aimed to identify peptide pairs and linked residues.

Rearrangement of a cross-link from K to another amino acid residue requires cleavage of the crosslink amide bond and formation of a new linkage to an attacking residue. However, cross-link amide bonds are more stable than peptide amide bonds under standard mass spectrometric conditions of electrospray and CID conditions for proteomics as used here. This can be explained by a mechanism in which protonation of a peptide amide nitrogen by a mobile proton and concurrent weakening of the peptide bond is followed by a nucleophilic attack by the carbonyl oxygen of the N-terminal neighboring amino acid residue to the carbonyl C atom of the weakened bond. This leads to formation of an unmodified C-terminal peptide fragment and an N-terminal part in the form of a peptide fragment with a C-terminal 5-membered oxazolone group[17]. If protonated these fragments are called y and b ions respectively. Also the peptide amino terminus can act as a nucleophile to cleave a protonated peptide bond in the so called diketopiperizine pathway[17]. This type of nucleophile is also used to cleave one of the two cross-link amide bonds formed with BAMG after reduction of the azido group to an amine group in the spacer of the cross-link (Fig 1B). However the other cross-link amide bond with the remaining cross-link remnant has lost the cleavability by this mechanism and also lacks the favorable configuration for cleavage of peptide bonds in the oxazolone pathway. With other cleavable NHS ester cross-linkers like BNP-NHP (PIR)[18], CBDPS[19], DSSO[20] and DSBU[21], both cross-link bonds survive the cleavage induced in the cross-link spacer. Nevertheless cross-link amide bond cleavage mechanisms are conceivable in the gas phase when the cross-link bond happens to be nearby in space to a residue of which the side chain group can become a nucleophile. This can be either hydroxyl group-, carboxyl group- or amino group-containing residues on the same peptide. Such residues could donate a proton to the amide nitrogen concomitant with nucleophilic attack of the oxyanion. carboxylate or deprotonated amino group to the positive amide carbon atom leading to cleavage of the amide bond and transfer of the cross-link remnant or cross-link with the other peptide to the attacking residue. This would lead to ester linkages with serine, threonine or tyrosine, anhydride linkages with aspartic acid or glutamic acid and to a new amide bond with lysine or amino-terminus. An example of anhydride formation in the gas phase is cleavage of a peptide bond C-terminal to aspartic acid[22]. A similar mechanism can be envisaged by rearrangement of an ester cross-link bond.

**Fig. 1.**
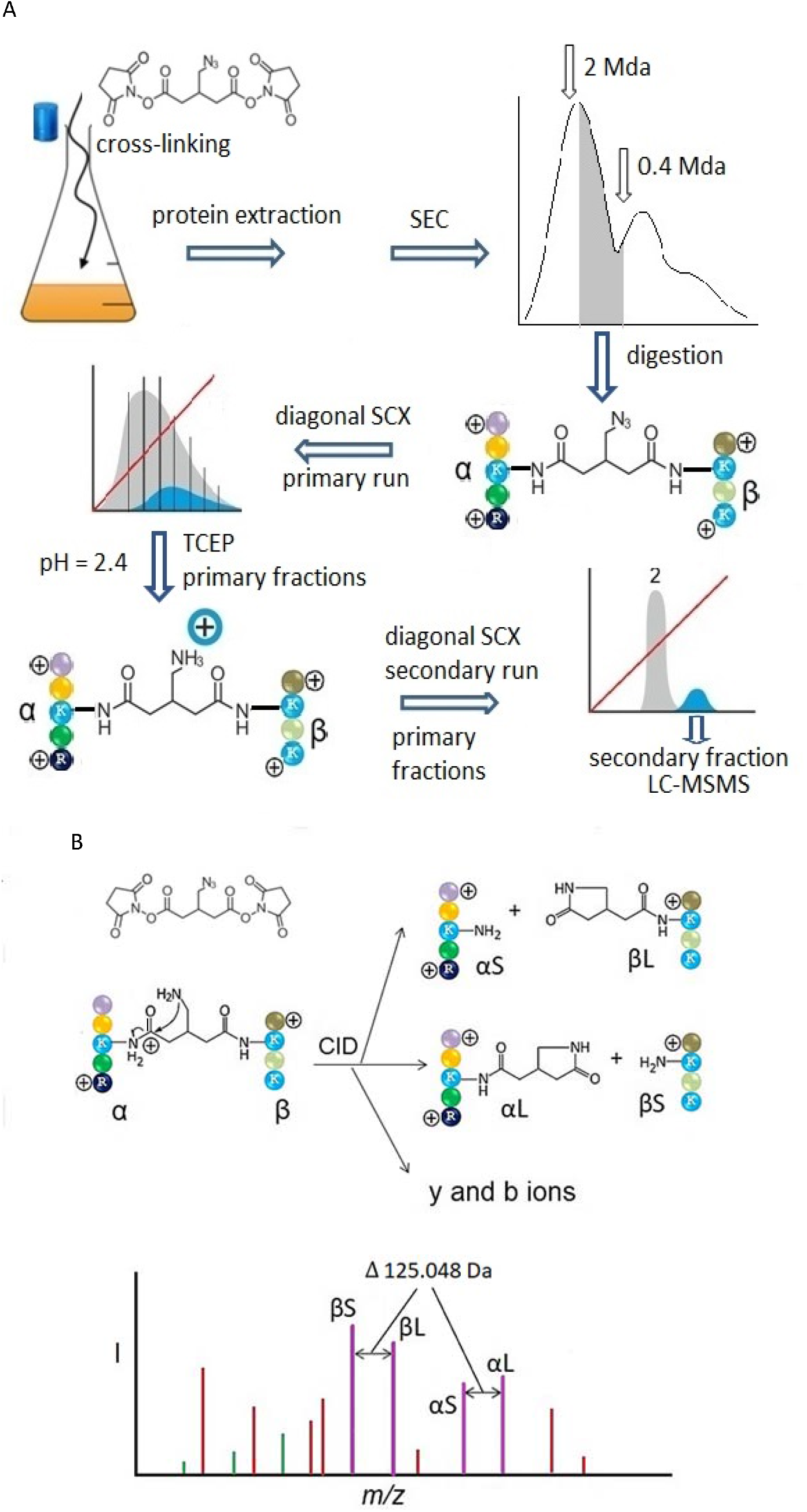
(A), Workflow from in vivo cross-linking with BAMG to LCMSMS. To a culture of exponentially growing cells BAMG (structure depicted in upper left part) is added and 5 min later cells are harvested and sonicated. The obtained soluble extract is subjected to SEC. The fraction eluting between approximately 1-2 MDa and 0.4 MDa is digested to obtain a peptide mixture with crosslinked α-β peptide pairs. An α-β peptide pair with amide linkages and amino acids represented by colored candies depicted below the carton of the SEC chromatogram. The peptide mixture is fractionated by strong cation exchange chromatography (primary run or first dimension SCX), using a mobile phase of pH 3 and a salt gradient (diagonal red line) of ammonium formate to elute bound peptides. Grey, regular peptides; cyan, cross-linked peptides. Primary fractions, indicated by vertical lines in the cartoon of the SCX chromatogram, are treated with TCEP to reduce the azido group in the spacer of the cross-linker to an amine group. At the pH of the mobile phase of SCX chromatography the amino group is protonated adding an extra positive charge to cross-linked peptides. The TCEP-treated primary fractions are then separately subjected to secondary runs of SCX. Here target peptides are sequestered from the bulk of unmodified peptides that elute at the same time as in the primary run, while the extra charge state of the cross-link peptides leads to elution at a later time (cartoon of the chromatogram of the secondary run of fraction 2 in lower right corner). Depicted peptide charge states before and after reduction of the azido group are calculated for pH 3, assuming full protonation of the two amino-termini plus 2 C-terminal basic amino-acid side chains, carboxylic acid side chains being uncharged under these conditions. (B), Gas phase cleavage reactions of BAMG-cross-linked peptides in which the azido group has been reduced to an amine group (slightly modified [31]. Collision induced dissociation (CID) of a crosslinked peptide pair leads to cleavages of the cross-link amide or ester bonds along with cleavages of peptide bonds resulting in y an b ions. Cleavage of a cross-link amide or ester bond probably occurs by nucleophilic attack of the amine in the spacer of the cross-link to a protonated carbonyl group of the amide or ester bond. This leads to formation of an unmodified peptide or short version of the cleavage product (αS or βS), the other peptide being modified by the remnant of the cross-linker in the form of a γ lactam adding 125.048 Da to the mass of the peptide. This is the longer version of the cleavage product (αL or βL). The indicated gas phase charge states of the cross-linked peptide and the cleavage products are arbitrarily. The lower part is a cartoon of a fragment mass spectrum with two pairs of cleavage products with the characteristic 125.048 Da mass difference (purple sticks) and some peaks of b (green) and y (red) ions.

To study whether such rearrangements after the inititial cross-link reaction can occur we use the NHS-ester bis(succinimidyl)-3-azidomethyl-glutarate (BAMG). Cross-linking by BAMG both enables enrichment of cross-linked peptides by diagonal strong cation exchange chromatography using reduction of the azido group to an amine group to induce a change in SCX chromatographic retention time of the target peptides (fig 1 A). Reduction of the azido group also renders the cross-link cleavable with collision induced dissociation in the sense that one of the two cross-link amide or ester bonds can be easily cleaved resulting in an unmodified peptide while the other peptide is modified with the cross-link remnant with a γ-lactam group (fig 1 B).

In this study SCXC retention time prediction of peptides is used to investigate to which extent amide-amide, amide-ester and ester-ester cross-links have been generated during treatment of exponentially growing Bacillus subtilis cells with BAMG added directly in the growth medium.

Distinction of these different cross-link types is based on a peptide charge difference at the pH of the mobile phase of SCXC (pH = 3) between peptides with an ester and an amide linkage, an amide linkage resulting in loss of a positively charged group, while an ester linkage has no effect on the charge state of the peptide. Possible amide to ester or ester to amide rearrangements after SCXC were detected by LC-MS/MS analysis of cross-linked peptides identified by database searching using pLink 2[23] as a search engine, allowing cross-linking by either amide bonds only to ε-amino groups of lysine residues or to amide or ester bonds by also including serine, threonine and tyrosine as possible cross-link sites in the search parameters. It appears that roughly 2% of a redundant dataset of about 2000 cross-linked peptide-pairs fractionated by SCXC consists of amide-ester cross-links, while 98% are linked via amide bonds only, in 3.5% of which protein N-termini are involved. Rearrangement in the gas phase, both from amide bonds at K to ester bonds at S, T and Y and to peptide N-termini were detected in about 5.5% of the cases. Ester to amide rearrangements after or during protein digestion were noticed in 4 instances.

## Materials and methods

### Preparation of cross-linked peptides after in vivo treatment of Bacillus subtilis cells with bis(succinimidyl)-3-azidomethyl-glutarate (BAMG)

We used an existing LS-MS/MS dataset for this study. The work flow for the preparation of cross-linked peptides for the LC-MS/MS dataset is schematically depicted in Fig. 1A as described in detail [24]. After cross-linking the soluble cell extract is subjected to SEC and the protein fraction eluting in the size range 0.4-2 MDa is selected for further analysis. Proteins are denatured by urea in the presence of iodoacetamide. Use of dithiothreitol to reduce possible disulfide bonds is avoided to prevent premature reduction of the azido group in the spacer of BAMG. After digested by trypsin the peptide mixture is subjected to a form of 2-D strong cation exchange chromatography (SCXC) also called diagonal chromatography[25]. Fractions containing cross-linked peptides after the primary SCX run (fraction 7-16) are treated for 2 h with tris(carboxyethyl)phosphine (TCEP) at pH 2.4 and 60 °C aimed to reduce the azido group in the spacer of the cross-link to an amine group. It should be noted that these conditions favor hydrolysis of esters[26] and also lead to partial cleavage of cross-link amide bonds[27,28]. The TCEP-treated fractions are then separately loaded on the SCX column. Since cross-linked peptides after reduction of the azido group have one positive charge more than the parent compound at the pH of the mobile phase in SCXC (pH = 3), thanks to protonation of the amine group in the spacer, their retention times increase enough for sequestering from the bulk of unmodified regular peptides in secondary runs of the particular primary fractions[29]. The shifted material containing the cross-linked peptides of each primary SCX fraction was collected in 3-5 subfractions. These were analyzed by LC-MS/MS as described, using the same CID conditions as applied for identification of regular peptides[24].

### Gas-phase cleavage reaction of BAMG-cross-linked peptides

Besides enabling efficient isolation by diagonal SCXC, reduction of the azido group also renders the cross-linked peptides cleavable by CID in the gas phase [30]. The cleavage reaction for cross-link amide bonds is depicted in Fig. 1B, but a similar mechanism can be envisaged for an ester cross-link. The largest of the two composite peptides is called the α peptide and the other one is the β peptide. Upon a cleavage event in one of the two cross-link amide or ester bonds one peptide is modified by the cross-linker remnant in the form of a stable γ-lactam moiety, resulting in a large (L) αL or βL peptide, while the other peptide is released unmodified resulting in a short (S) αS or βS peptide. Along with cleavage at a cross-link amide bond, secondary cleavages at the peptide bonds can occur enabling identification of the composite peptides and residues involved in cross-linking. Fig. S1 depicts the different types of fragment ions used for database searching that are generated by CID of BAMG-cross-linked peptides.

### LC-MS/MS dataset of cross-linked peptides

To identify cross-linked peptides and to study how cross-linked sites are distributed via linkages to the 5 different NHS-reactive groups in proteins we used the dataset of mgf files that is available via ProteomeXchange with identifier PXD006287).

### Identification of cross-linked peptides

The search engine pLink 2 [23] was used in the stepped-HCD mode to interrogate the entire *B. subtilis* sequence database for cross-linked peptides in two searches. The original 3-5 mgf files of LC-MS/MS runs of secondary SCX fractions were pooled for each primary fraction before pLink 2 analysis, besides the mgf files of the 4 secondary fractions from primary fraction 9, denoted 9.2 - 9.5. Mgf files from secondary fractions of primary fractions 14, 15 and 16 were also pooled before pLink 2 analysis. In one search with pLink 2 of the thus obtained mgf files the specificity of BAMG was set to reaction with only K, i.e. at the ε amino group in the side chains of lysine residues (K search). In the second search (KSTY search) also cross-linking via the hydroxyl groups of serine, threonine and tyrosine was taken into account. For validation purposes also results of a pLink 2 search for crosslinking at K and protein N-termini was used, denoted [K search. The other search parameters were: digestion with trypsin, a maximum of two missed cleavages per peptide, precursor and fragment ion mass error 25-75 ppm, dependent on the mgf file, a fixed carbamidomethyl modification at cysteine and oxidation of methionine as a variable modification.

### Identification of cross-linked residues

Distinction between cross-links formed by two amide bonds, two ester bonds or one amide and one ester bond is made by SCXC retention time prediction. For MS/MS assessment of the residues involved in cross-linking only fragment ions containing the intact cross-link or the γ-lactam remnant of the cross-link are taken into account (Fig. S1). Doubly charged fragment ions ≤ 700 m/z are excluded from this analysis.

### False discovery rate (FDR)

Data were analyzed at 5% FDR at cross-linked spectrum matches (CSM) with separate control of the FDR for intra-protein cross-links also called self-links[11] and inter-protein cross-links (inter-links). Since the experimentally estimated FDR for inter-links, determined by an entrapment database approach, is much higher than 5%, while no decoy sequences were found for self-links [31], the interlink data were further filtered by the following parameters: (i) unambiguous assignment of the number of y and b fragments ions, dependent on the length of α and β, double charged fragments ≤ 700 m/z not taken into account, (ii) matched intensity (MI) of fragment ions, (iii) unambiguous peptide identification and (iv) retention time in SCXC[31].

## Results

### Comparison of searches with cross-linking allowed at K only and at K, S, T and Y

The LC-MS/MS dataset of cross-linked peptides obtained from *Bacillus subtilis* treated in culture with the cleavable cross-linker BAMG was analyzed by pLink 2 in a search for cross-linking at K only and at K along with S, T and Y. The search for cross-links at K revealed 210 inter-link CSMs, 20 homodimer CSMs and 1609 self-link CSMs after about 20 min calculation by pLink 2 on a laptop with solid state drive disc and a first generation i5 processor. In the KSTY search about 10% less inter-link CSMs and about 2% more self-link CSMs were detected after about 2 h calculation (Table 1, Table S1). The increase in calculation time reflects the large search space when four possible cross-linkable residues are taken into account.

**Table 1.** CSMs in K and KSTY searches after the composite filter and assigned CSMs with amide-amide and amide-ester cross-links.

|  | *Only*<br>*in*<br>*KSTY*<br>*search* | *Only in*<br>*K*<br>*search* | *In both K*<br>*and KSTY*<br>*search* | *Total* | *Amide-*<br>*amide*<br>*cross-*<br>*links*<br>*assigned* | *Amide-*<br>*ester*<br>*cross-*<br>*links*<br>*assigned* | *% amide-*<br>*ester cross-*<br>*links* |
| --- | --- | --- | --- | --- | --- | --- | --- |
| Inter-links | 9 | 29 | 181 | 219 | 218 | 1 | 0.5 |
| Homodimers | 3 | 4 | 16 | 23 | 23 | 0 | 0 |
| Self-links | 133 | 96 | 1513 | 1742 | 1701 | 41 | 2.4 |
| Total | 145 | 129 | 1710 | 1984 | 1942 | 42 | 2.1 |
| % | 7.3 | 6.5 | 86.2 | 100 |  |  |  |

Inter-links are cross-linked peptides from two different proteins (inter-protein cross-links); Homodimers are cross-links between identical or overlapping peptides; self-links are cross-linked peptides from the same proteins (intra-protein cross-links).

### SCXC retention time prediction can be used to distinguish amide from ester cross-linking

In Fig. 2 the importance of mass and charge at pH 3 for SCXC retention time prediction in order to distinguish between ester and amide cross-links is demonstrated. In this figure the distribution of charge/mass ratios is depicted in the different SCXC fractions of the 1002 precursor ions (of which three were excluded from the analysis), that were assigned to be linked to exclusively K residues, both in the K search and in the KSTY search so that there charge at pH 3 can be calculated (Fig.S1, row 1421-3414). Importantly, it appears that for a data set with a similar mass range the charge state can probably be correctly predicted for more than 99% of the cases. Only for 2 precursors in SCX fraction 9 the charge at pH 3 could be either 3^+^-4^+^ or 4^+^-5^+^. The cross-link-peptide of charge 3.44^+^ with the lowest z/m ratio in this SCX fraction (0.8638) would have a z/m ratio of 1.1209 with one positive charge more, which is within the z/m ratio of the 4^+^-5^+^ charged peptides. With one charge less, the cross-linked peptide of charge 4.20^+^ with the highest m/z value (1.2101) would have an m/z value of 0.9164, which is within the m/z range of 3^+^-4^+^ charged peptides in this SCX fraction. In the remaining 997 precursor ions no mass overlap between the different possible charge states in each SCXC fraction occurs. This high predictability is both due to the extremely low FDR of self-links as identified by pLink 2 and inter-links after application of a composite filter[7,31] and to the large increase in retention time with an increase of one net charge. This large shift induced by an increase in net charge is illustrated by nine cross-linked peptide pairs from the used precursor collection. At pH=3 missed cleavage variants of eight peptide pairs have one protonated amino acid extra while the remaining case has four amino acids and one proton extra in the remaining case (Table S2). The species without a missed cleavage eluted in fractions 7 to 10. In all cases the missed cleavage variants eluted 4 or 5 fractions later.

**Fig 2.**
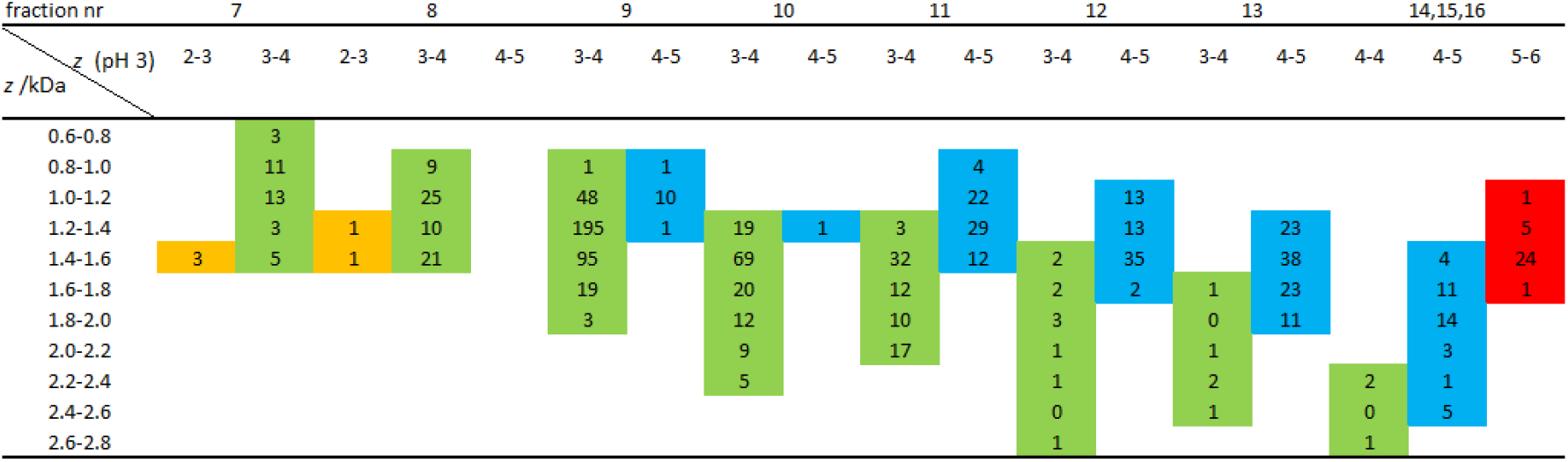
Distribution of cross-linked peptides linked via lysine residues as identified both in the K search and KSTY search from different primary fractions of SCX chromatography as a function of charge (z) at pH 3 and z/mass (kDa) ratio. For calculation of z it is assumed that N-termini, H, K and R are fully protonated, cross-linked K residues are uncharged and carboxyl groups (at D, E and peptide C-termini) are protonated for 92 % under the experimental conditions of SCX chromatography. CSMs (n) are listed in yellow(2^+^-3^+^), green (3^+^-4^+^), blue (4^+^-5^+^) and red (5^+^-6^+^) highlighted rows.

### Scrambling of cross-link sites from K to S, T and Y

Precursor ions identified in both the K and KSTY search have the same SCXC elution time and, therefore, their composite peptides must have been cross-linked via amide-amide linkages in all cases. Nevertheless more than 40% of the precursors identified in both the K search and the KSTY search were nominated by the search engine in the KSTY search to be linked by one or both ester bonds (Table S1, row 5-1420). While retention time prediction indicated amide-amide cross-links, inspection of MS/MS spectra should reveal an explanation for this apparent discrepancy. Fragment ions indicating cross-linking to a residue other than the presumed K include either Ly ions and y ions containing the intact cross-link on the C-terminal side of the concerning K residue or Lb ions and b ions containing the intact cross-link on the N-terminal side of the concerning K residues. (Materials and methods, section identification of linked residues and Fig. S1).

Scrambling became clear by MS/MS analysis following the observation that the matched intensity of 218 MS/MS spectra out of the 1710 precursors ions selected for MS/MS in the KSTY search differs from that in the K search (row 5-440 of Table S1). These mass spectra were selected for further analysis. In 104 cases in the KSTY search one or more unambiguous fragment ions were nominated that contained either the intact cross-link with the other peptide or the cross-link remnant in the form of γ-lactam group of 125.048 Da, while these fragment ions did not comprise the K residue as nominated for linkage in the K search (row 5-212 of Table S1). Discriminating fragment ions are denoted in column V of Table S1. Non-redundant MS/MS spectra (highlighted green in column C of Table S1) are depicted in supplemental material, file KTSY_Spectra. We interpret these observations to mean that an amide bond has been exchanged for an ester bond after the primary SCX run. Likewise, 5 LC-MS/MS spectra of precursors ions identified in only the KSTY search and predicted to contain amide-amide linkages based on SCXC retention time prediction contain fragment ions that point to amide-ester linkages (row 3425-3429). In cases were pLink 2 nominated the involvement of two ester linkages, we observed scrambling to a hydroxy amino acid, but in only one of the two composite peptides. These cases are highlighted blue in column J of Table S1. So, we found evidence for cross-link site rearrangements for 109 CSMs by inspection of selected MS/MS spectra generated in the KSTY search.

Cross-links mentioned in row 213-276 of Table S1 have also unequal matched intensity scores in the K and KSTY search but we could not detect discriminating fragment ions to identify the linkage site in these cases. Cross-links listed in row 277-440 of Table S1, have higher scores in the K search than in the KSTY search and in these cases cross-linking at K was unambiguously confirmed by MS/MS analysis, without evidence for cross-link scrambling.

### Unambiguous identification of the residue involved in cross-link scrambling is possible in nearly 45% of the cases

Unambiguous assignment of residues to which a cross-link has been transferred from a K residue is possible if identified fragment ions excluding K as a cross-link contain only one residue of the amino acids S, T, or Y, which are included in the KSTY search, and none of the susceptible amino acids D, E or the N-terminus, which are not included in this search. This is the case in 40 of the 109 CSMs showing evidence for scrambling, highlighted red for transfer of the cross-link to S, blue for transfer to T and brown for transfer to Y in column Q of Table S1. We also noticed in eight CSMs the presence of an Lb ion, or a b ion with the intact cross-link, while these fragments did not contain one or more of the residues K, S, T, Y, D, E or a protein amino-terminus. This indicates that the cross-link must have been transferred from K to the peptide amino terminus, highlighted orange in column Q of Table S1. In the remaining 61 cases the discriminating fragment ions contained two or more of the residues S, T, Y, D, E or a peptide amino-terminus, preventing unambiguous identification of the site of transfer.

### Cross-link rearrangement is partial and occurs predominantly in the gas phase

Cross-link scrambling is probably partial. In nearly all 109 cases the discriminating fragments not containing a K residue pointing to cross-linking at S, T, Y, D, E, or peptide amino-N-termini are relatively weak. Moreover, we found in 39 cases fragment ions that indicate both a cross-link at K and in the same peptide also at either an S or T residue (row 135-212, Table S1). Non-redundant MS/MS spectra showing this linkage at K are depicted in Supplemental material, file K_Spectra, highlighted orange in column C of Table S1. We also encountered two precursor ions with scrambling to two different residues in the same peptide (row 209-212). Probably cross-link rearrangement occurs in the gas phase. Otherwise one would expect chromatographic separation of the scrambled and unmodified molecules of cross-linked peptide pairs during LC-MS/MS.

### A spacing of three or four amino acids apart in the linear sequence may favor scrambling

The distances in the linear sequence between susceptible residues for transfer of the cross-link and the cross-linked K-residue appears to be nonrandomly distributed with a preference for two, three or four residues apart (Fig. 3), peptide amino-termini not taken into account. Amino acids spaced three or four apart are close to each other if arranged in an α-helix. This preferred spacing was observed both in the entire nonredundant set of peptide pairs by taking all possible reactive amino acids S, T, Y, D and E into account and in the non-redundant set of cross-linked K-residue pairs in which the susceptible amino acid for cross-link transfer was unambiguously identified (Fig. S2). Scrambling can also occur between amino acids relatively far apart in the sequence, about 20% of the cases concerning distances of 8 amino acids or more (Fig. 3).

**Fig. 3.**
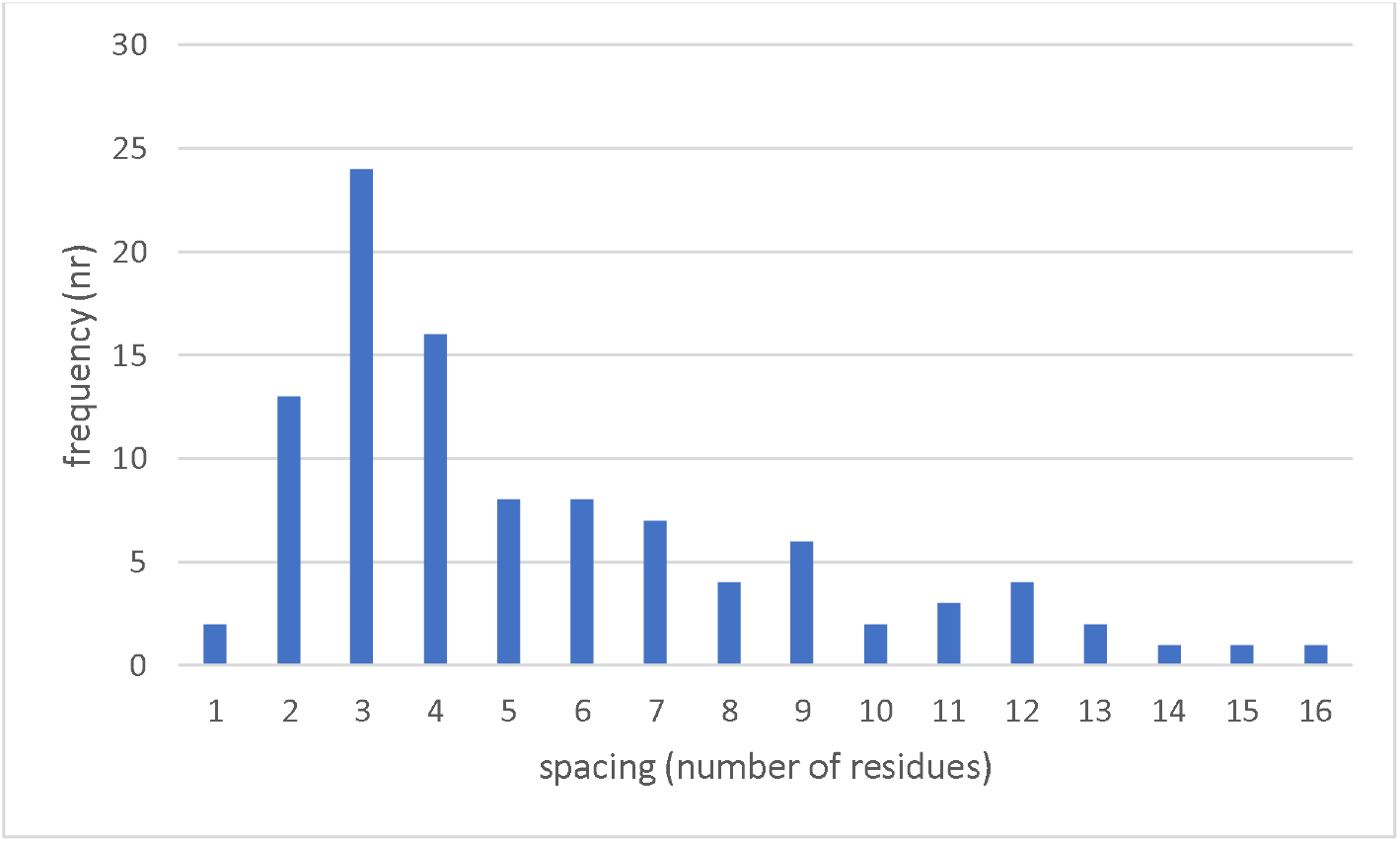
Frequency of spacing between residues susceptible for cross-link transfer and cross-linked K residues. Results are from all non-redundant peptide pairs with evidence for cross-link scrambling.

So, cross-linked peptides can undergo in the analysis phase chemical modifications that may result in a cross-link at another residue than at the amino acid that was connected during the actual cross-link reaction. This rearrangement is probably facilitated by a defined, possibly transient, spatial configuration of donor and acceptor in the gas phase.

### Cross-linking to protein N-termini

A special group of cross-linked peptide pairs is formed by species with a protein N-terminal peptide. A protein N-terminal peptide can be nominated by pLink 2 as member of a cross-linked peptide pair only if containing one or more K, S, T or Y residues, since the amino terminal amine group is not taken into account as a cross-link target in a K or KSTY search. Table S1 lists 68 peptide pairs only identified in the KSTY search of which one of the two composite peptides contains the protein N-terminus without a cross-linkable K in that peptide (row 3468-3535 of Table S1). SCXC retention times predict cross-linking at the protein N-terminus in all cases, while assumption of a free N-terminus and instead cross-linking to a S, T or Y as proposed by pLink 2 in the KSTY search would be in disagreement with SCXC retention times. A pLink 2 search allowing cross-linking at both K and the protein N-terminus (denoted [K search) revealed nearly all candidates with a cross-linkable K residue in one composite peptide and a protein N-terminus in the other one. The search including both K and protein N-termini appeared to be slightly less sensitive than a search at K only, but cross-linking at the protein N-terminus was unambiguously confirmed by MS/MS analysis in several instances (blue highlighted in column C of Table S1). The corresponding MS/MS spectra are depicted in supplemental material, file [K_Spectra. These results supports the usefulness of inclusion of a protein-N terminus as cross-link target in the search parameters and further underscores the power of SCXC retention time prediction to distinguish amide from ester cross-linking by N-hydroxy succinimidyl esters.

### Cross-linking at S,T or Y

The last group of cross-linked peptide pairs that were only nominated in the KSTY search comprised 46 CSMs of which one of the two composite peptides lacked a cross-linkable K residue or a protein amino terminus, indicating the presence of at least one ester linkage (row 3524-3569 in Table S1). Indeed, retention time prediction provided evidence for the presence of an ester-amide type crosslink in 42 cases. However in the remaining 4 cases (row 3566-3569 of Table S1) the SCX retention times correspond to 3 or more chromatographic fractions earlier than expected with the presence of one ester linkage. This indicates that these cross-links must comprise two amide linkages. We explain this observation by assuming that after digestion and before the primary SCX run an initially formed ester linkage has been attacked by the now available reactive amino terminus or a peptide C-terminal K residue resulting in an amide linkage. MS/MS analyses supported linkage to the peptidyl N-terminus (Fig. S3) or to the ε-amino group of a peptidyl C-terminal K residue (Fig. S4 and Fig S5).

### Overview of cross-linking by amide and ester linkages and the occurrence of cross-link scrambling

The results together show a limited initial cross-linking at S, T or Y, comprising 2.1 % of all precursor ions. About 3.5% of all cross-links are to protein N-termini. Scrambling of cross-link sites is more abundant with 5.5% of all CSMs. Scrambling occurs probably predominantly in the gas phase and, to a limited extent, also to peptide N-termini or to C-terminal lysine residues after digestion but before SCXC.

## Discussion

To our knowledge exchange reactions between cross-linked sites after the cross-link reaction have not been studied before, but as shown here are observed in more than 5% of the cases under our standard mass spectrometric conditions of electrospray and CID conditions for proteomics. This implies that a cross-linked residue identified by MS/MS analysis can be different from the residue actually modified in the cross-link reaction. Since only residues actually modified in the cross-link reaction are informative for the 3-D protein structure, in the sense that linked atoms cannot have been father away from each other than the length of the spacer during the cross-link reaction, control experiments may be required to validate the cross-link type in order to obtain the correct maximal possible spatial distance between linked residues. There is a certain preference for scrambling between amino acids spaced three and four apart in the linear sequence, so that they are in an α helix close in space. However, also much longer distances between donor amino acid and acceptor occur. Especially in such cases correct identification of the linked residues as formed during the cross-link reaction is important for insight at the highest possible resolution in the spatial arrangement of proteins in a complex. As shown here SCXC retention time prediction is a powerful method to distinguish amide from ester cross-linking and to detect scrambling. SCXC is already often used to diminish the complexity of digests and for enrichment of the target peptide pairs.

We have shown here that cross-linking to hydroxyl group-containing amino acids residues occurs in about 2% of the cases, the large majority of cross-link targets being the ε-amino groups of lysine residues and, to a lesser extent, protein N-termini. In another large scale investigation about the distribution of cross-links among reactive sites a frequency of about 14% was found to involve linkages at S, T or Y, with similar relative contributions as found here for scrambling[10]. The 7-fold difference in the extent of ester cross-linking between this and our study can at least partly be explained by acid hydrolysis of cross-link ester bonds under our experimental conditions. Treatment of cross-linked peptides with TCEP at low pH and high temperature is an obligatory step in sample preparation to enable isolation of BAMG-cross-linked peptides and to render these target peptides cleavable in the gas phase. It should be noted that by the TCEP treatment also partial cleavage of K-K cross-links occurs along with reduction of the azido group [29] according to the mechanisms in a report [27] on which the design of BAMG was based [32]. It is not known whether, besides low pH and elevated temperature, the reducing conditions also add to the preferential hydrolysis of ester bonds. Besides acid hydrolysis of esters under our conditions, it can probably not be excluded that gas phase scrambling may contribute to the relatively high extent of cross-links at hydroxyl group-containing amino acids as found in the work by Giese *et al*. [10].

Previously we have demonstrated the power of SCXC in the isolation of cross-linked peptides from complex mixtures [24,29] and the usefulness of SCXC retention time prediction in the lowering of the FDR of in vivo detected protein-protein interactions by CXMS [7,31]. Here we show the application of SCXC retention time prediction for determination of the relative contribution of amide and ester linkages in cross-linked peptides and for detection of cross-link scrambling in the analysis phase of peptide identification. This approach can be followed with all types of cleavable and non-cleavable NHS-esters as the cross-linking agent and may contribute to the further development of the CXMS technology for discovering in vivo protein-protein interactions and for determination of the spatial arrangement of subunits in large protein complexes. Progress in these areas will increase our understanding of the functional organization of living matter and will help in drug design and finding new drug targets.

## Supporting information

supplemental material Spectra[K_Search

supplemental material Spectra_K_Search

supplemental material Spectra_KTSY_Search

---

**Fig. S1.**
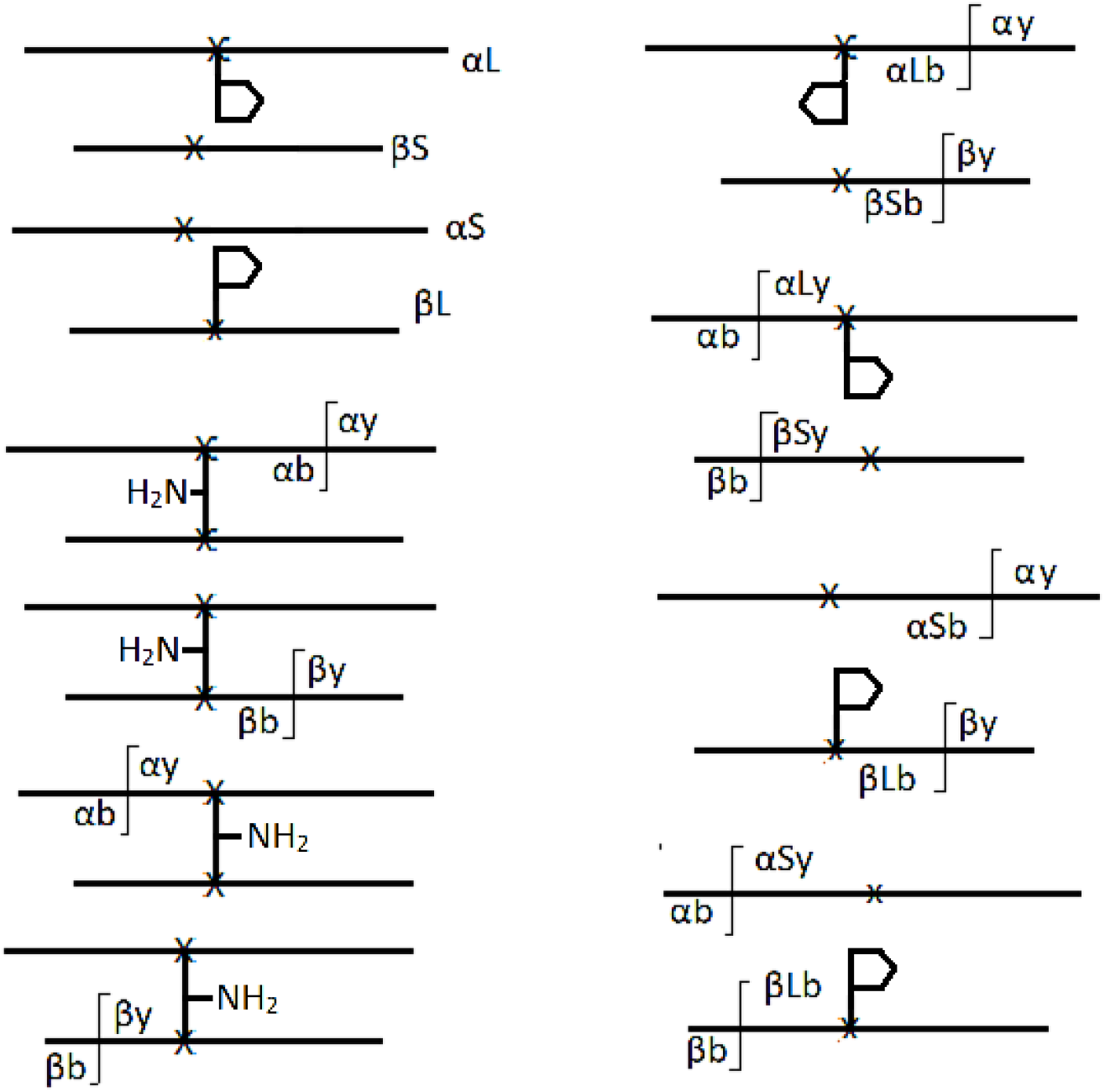
Terminology of fragment ions used for identification of BAMG-crosslink peptides with respect to amino acid sequence of composite peptides, cross-linked residues and parent proteins. X, crosslink site. Left part, possible fragment ions after one cleavage event in a cross-linked peptide pair. Upper left part, possible fragment ions after one cleavage event of a cross-link bond, resulting in αL and βs ions if the cleavage is at the β peptide, leaving the γ-lactam remnant on the α peptide, or αS and βL ions if the cleavage is at the α peptide, with the γ-lactam remnant on the β peptide. These fragment ions are depicted as pink colored sticks in mass spectra. Lower left part, possible fragment ions after one cleavage event of a peptide bond, either resulting in αy and αb or βy and βb ions with the cross-link at the b ions if the cleavage occurs on the C-terminal side of the cross-link or with the cross-link at the y ions if the cleavage event is on the N-terminal side of the cross-link. Right part, y and b fragment ion terminology after two cleavage events of a cross-linked peptides pair, one cleavage at a cross-link bond and one at a peptide bond. Upper right part, cleavage of the cross-link at the β peptide and, from the top down, at a peptide bond on the C-terminal side of the cross-linked residue of the α or β peptide and on the N-terminal side of the cross-linked residue of the α or β peptide. Lower right part, the same for cleavage of the cross-link at the α peptide. For validation of pLink 2-identifed residues involved in cross-linking only yL, bL and y and b ions with the intact crosslink to the other peptide are taken into account, since yS, bS, y and b ions after cleavage of the crosslink are unmodified peptide fragment ions, and therefore provide no evidence for the cross-link sites on the peptides. In mass spectra y ions are depicted as red sticks and b ions as green sticks.

**Fig. S2.**
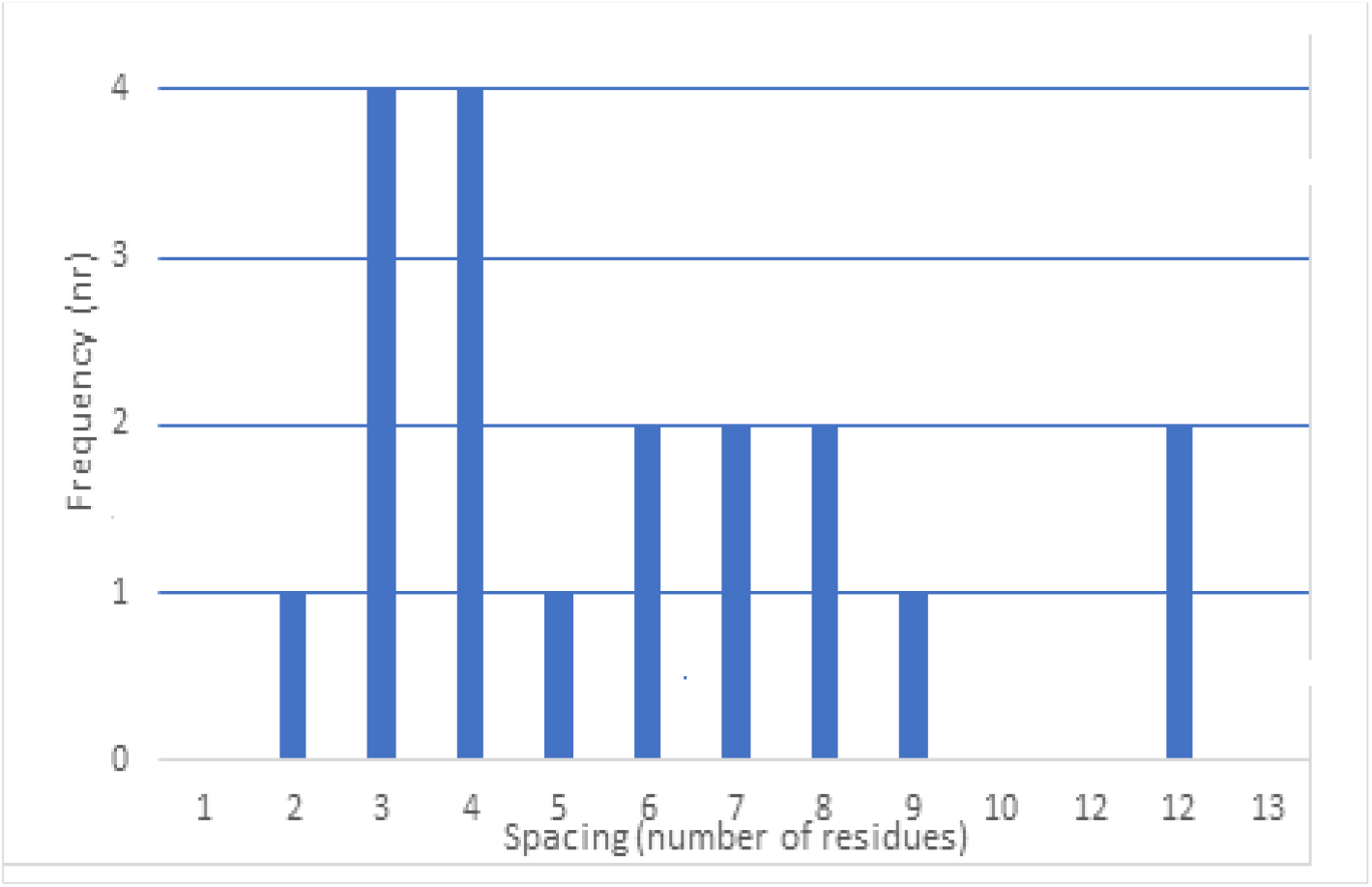
Frequency of spacing between residues susceptible for cross-link transfer and cross-linked K residues. Results are from non-redundant peptide pairs with evidence for cross-link scrambling and unambiguous identification of the residue to which the cross-link or cross-linked remnant has been transferred.

**Fig. S3.**
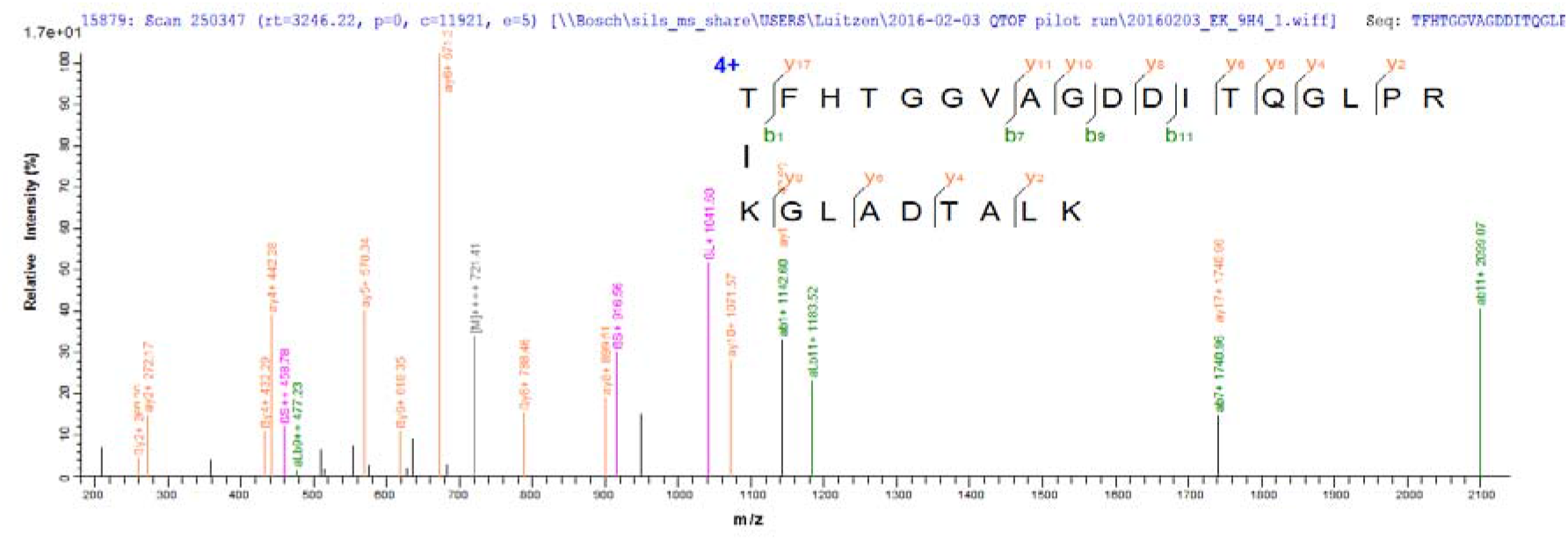
MS/MS spectrum of a cross-linked peptide from the βͰ subunit of RNA polymerase containing an amide-amide cross-link between the only cross-linkable K (K1 at the β peptide) and the α peptide N-terminus. The SCXC retention time shows that this peptide pair is linked by two amide bonds, although the α peptide lacks a lysine residue and does not contain the protein N-terminus. The αb1^+^, αLb11^+^, αb7^+^ and αb11^+^ fragment ions together with the SCXC retention time can be explained by assuming exchange between an initially formed ester cross-link to either T1 or T4 and the α peptide N-terminus after or during digestion and before SCXC.

**Fig. S4.**
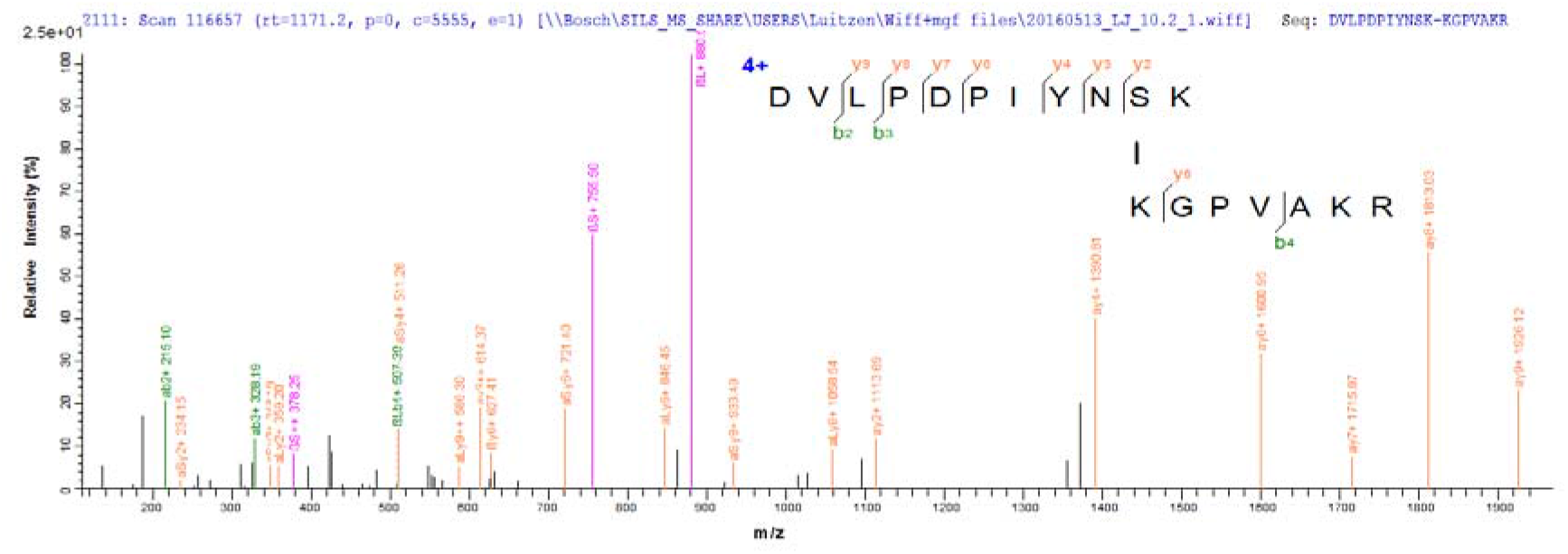
MS/MS spectrum of a self-link from the small ribosomal protein 7 showing replacement of an S10-K1 ester-amine cross-link to an amine-amine K11-K1 cross-link. In this crosslinked peptide pair the α peptide C-terminal K, not being the protein C-terminus, cannot have been cross-linked in vivo. Nevertheless according to its SCX retention time the peptide pair contains an amide-amide cross-link. Since the aLy2 fragment indicates that the cross-link must be at the C-terminus of the α peptide the S10-K1 cross-link is replaced by a K11-K1 cross-link after digestion and before SCX chromatography. Note the limited fragmentation of the β peptide in this self-link. The chance to detect by accident two peptides from the same protein in a cross-linked peptide pair by searching an entire species specific sequence database is so small that a much lower stringency for assignment based on MS/MS is required than for inter-links.

**Fig. S5.**
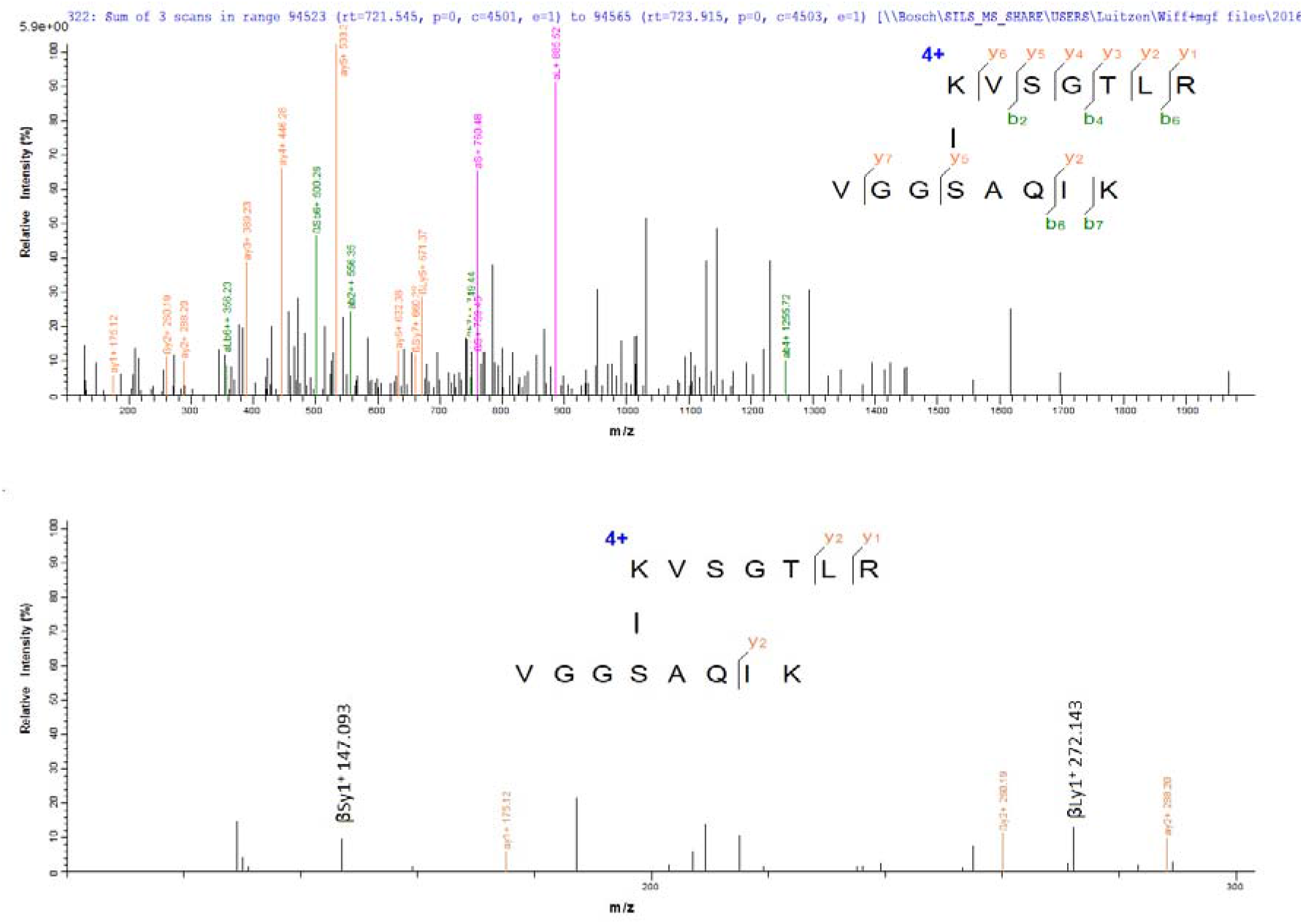
MS/MS spectrum (upper panel) of a cross-linked peptide pair from ATP synthase subunit α showing cross-linking to the C-terminal K of peptide B. This cross-linked peptide pair was identified in the KTSY search and not in the K search. The only cross-linkable residue in the β peptide is S4. However, SCXC prediction indicates that the cross-link is formed by two amide linkages. The lower panel zoomed in to the *m/z* range 100-300 shows the present of both a βSy1^+^ and a βLy1^+^ ion at *m/z* = 147.093 and 272.132, resp. An explanation for this observation is a transfer of the cross-link from S4 to the ε amino group of K8 of the β peptide. This transfer must have taken place after or during digestion and before SCXC since K8 of this peptide is not a protein C-terminus. Another possibility would be tryptic digestion at K despite the presence of a cross-link at this residue. Both possibilities are not taken into account by the search engine and therefore a βLy1^+^ ion and a βSy1^+^ ion were assigned by pLink 2 in the KTSY search.

## References

[1] E.V. Petrotchenko, C.H. Borchers, Protein Chemistry Combined with Mass Spectrometry for Protein Structure Determination, Chem Rev, 122 (2022) 7488–7499.

[2] A. Graziadei, J. Rappsilber, Leveraging crosslinking mass spectrometry in structural and cell biology, Structure, 30 (2022) 37–54.

[3] L. Piersimoni, P.L. Kastritis, C. Arlt, A. Sinz, Cross-Linking Mass Spectrometry for Investigating Protein Conformations and Protein-Protein InteractionsͰA Method for All Seasons, Chem Rev, 122 (2022) 7500–7531.

[4] H. Zhang, X. Tang, G.R. Munske, N. Tolic, G.A. Anderson, J.E. Bruce, Identification of proteinprotein interactions and topologies in living cells with chemical cross-linking and mass spectrometry, Mol Cell Proteomics, 8 (2009) 409–420.

[5] M. Matzinger, K. Mechtler, Cleavable Cross-Linkers and Mass Spectrometry for the Ultimate Task of Profiling Protein–Protein Interaction Networks in Vivo, J. Proteome Res., 20 (2021) 78–93.

[6] X. Tang, H.H. Wippel, J.D. Chavez, J.E. Bruce, Crosslinking mass spectrometry: A link between structural biology and systems biology, Protein Sci, 30 (2021) 773–784.

[7] L. de Jong, W. Roseboom, G. Kramer, Towards low false discovery rate estimation for proteinprotein interactions detected by chemical cross-linking, Biochim Biophys Acta Proteins Proteom, 1869 (2021) 140655.

[8] S. Kalkhof, A. Sinz, Chances and pitfalls of chemical cross-linking with amine-reactive N-hydroxysuccinimide esters, Anal Bioanal Chem, 392 (2008) 305–312.

[9] S. Mädler, C. Bich, D. Touboul, R. Zenobi, Chemical cross-linking with NHS esters: a systematic study on amino acid reactivities, Journal of Mass Spectrometry, 44 (2009) 694–706.

[10] S.H. Giese, L. Fischer, J. Rappsilber, A Study into the Collision-induced Dissociation (CID) Behavior of Cross-Linked Peptides, Mol Cell Proteomics, 15 (2016) 1094–1104.

[11] S. Lenz, L.R. Sinn, F.J. O’Reilly, L. Fischer, F. Wegner, J. Rappsilber, Reliable identification of protein-protein interactions by crosslinking mass spectrometry, Nat Commun, 12 (2021) 3564.

[12] I. Parfentev, S. Schilbach, P. Cramer, H. Urlaub, An experimentally generated peptide database increases the sensitivity of XL-MS with complex samples, Journal of Proteomics, 220 (2020) 103754.

[13] M. Götze, C. Iacobucci, C.H. Ihling, A. Sinz, A Simple Cross-Linking/Mass Spectrometry Workflow for Studying System-wide Protein Interactions, Anal Chem, 91 (2019) 10236–10244.

[14] P.-L. Jiang, C. Wang, A. Diehl, R. Viner, C. Etienne, P. Nandhikonda, L. Foster, R.D. Bomgarden, F. Liu, A Membrane-Permeable and Immobilized Metal Affinity Chromatography (IMAC)-Enrichable Cross-Linking Reagent to Advance In Vivo Cross-Linking Mass Spectrometry, Angew Chem Int Ed Engl, (2021).

[15] J.P. Mohr, P. Perumalla, J.D. Chavez, J.K. Eng, J.E. Bruce, Mango: A General Tool for Collision Induced Dissociation-Cleavable Cross-Linked Peptide Identification, Anal Chem, 90 (2018) 6028–6034.

[16] K. Yugandhar, T.-Y. Wang, A.K.-Y. Leung, M.C. Lanz, I. Motorykin, J. Liang, E.E. Shayhidin, M.B. Smolka, S. Zhang, H. Yu, MaXLinker: Proteome-wide Cross-link Identifications with High Specificity and Sensitivity, Mol Cell Proteomics, 19 (2020) 554–568.

[17] Béla. Paizs, S. Suhai, Towards understanding the tandem mass spectra of protonated oligopeptides 1: Mechanism of amide bond cleavage, J. Am. Soc. Mass Spectrom., 15 (2004) 103–113.

[18] D. Tan, Q. Li, M.-J. Zhang, C. Liu, C. Ma, P. Zhang, Y.-H. Ding, S.-B. Fan, L. Tao, B. Yang, X. Li, S. Ma, J. Liu, B. Feng, X. Liu, H.-W. Wang, S.-M. He, N. Gao, K. Ye, M.-Q. Dong, et al., Trifunctional cross-linker for mapping protein-protein interaction networks and comparing protein conformational states, Elife, 5 (2016).

[19] E.V. Petrotchenko, J.J. Serpa, C.H. Borchers, An isotopically coded CID-cleavable biotinylated cross-linker for structural proteomics, Mol Cell Proteomics, 10 (2011) M110.001420.

[20] A. Kao, C. Chiu, D. Vellucci, Y. Yang, V.R. Patel, S. Guan, A. Randall, P. Baldi, S.D. Rychnovsky, L. Huang, Development of a novel cross-linking strategy for fast and accurate identification of cross-linked peptides of protein complexes, Mol Cell Proteomics, 10 (2011) M110.002212.

[21] M.Q. Müller, F. Dreiocker, C.H. Ihling, M. Schäfer, A. Sinz, Cleavable cross-linker for protein structure analysis: reliable identification of cross-linking products by tandem MS, Anal Chem, 82 (2010)6958–6968.

[22] C. Gu, G. Tsaprailis, L. Breci, V.H. Wysocki, Selective gas-phase cleavage at the peptide bond C-terminal to aspartic acid in fixed-charge derivatives of Asp-containing peptides, Anal Chem, 72 (2000)5804–5813.

[23] Z.-L. Chen, J.-M. Meng, Y. Cao, J.-L. Yin, R.-Q. Fang, S.-B. Fan, C. Liu, W.-F. Zeng, Y.-H. Ding, D. Tan, L. Wu, W.-J. Zhou, H. Chi, R.-X. Sun, M.-Q. Dong, S.-M. He, A high-speed search engine pLink 2 with systematic evaluation for proteome-scale identification of cross-linked peptides, Nat Commun, 10 (2019) 3404.

[24] L. de Jong, E.A. de Koning, W. Roseboom, H. Buncherd, M.J. Wanner, I. Dapic, P.J. Jansen, J.H. van Maarseveen, G.L. Corthals, P.J. Lewis, L.W. Hamoen, C.G. de Koster, In-Culture Cross-Linking of Bacterial Cells Reveals Large-Scale Dynamic Protein-Protein Interactions at the Peptide Level, J Proteome Res, 16 (2017) 2457–2471.

[25] K. Gevaert, J. Van Damme, M. Goethals, G.R. Thomas, B. Hoorelbeke, H. Demol, L. Martens, M. Puype, A. Staes, J. Vandekerckhove, Chromatographic isolation of methionine-containing peptides for gel-free proteome analysis: identification of more than 800 Escherichia coli proteins, Mol Cell Proteomics, 1 (2002) 896–903.

[26] J.H. Richards, D.J. Cram, G.S. Hammond, Elements of Organic Chemistry, in: first, McGraw-Hill, New York, 1967: p. 224.

[27] J.W. Back, O. David, G. Kramer, G. Masson, P.T. Kasper, L.J. de Koning, L. de Jong, J.H. van Maarseveen, C.G. de Koster, Mild and chemoselective peptide-bond cleavage of peptides and proteins at azido homoalanine, Angew Chem Int Ed Engl, 44 (2005) 7946–7950.

[28] H. Buncherd, M.A. Nessen, N. Nouse, S.K. Stelder, W. Roseboom, H.L. Dekker, J.C. Arents, L.E. Smeenk, M.J. Wanner, J.H. van Maarseveen, X. Yang, P.J. Lewis, L.J. de Koning, C.G. de Koster, L. de Jong, Selective enrichment and identification of cross-linked peptides to study 3-D structures of protein complexes by mass spectrometry, J Proteomics, 75 (2012) 2205–2215.

[29] H. Buncherd, W. Roseboom, B. Ghavim, W. Du, L.J. de Koning, C.G. de Koster, L. de Jong, Isolation of cross-linked peptides by diagonal strong cation exchange chromatography for protein complex topology studies by peptide fragment fingerprinting from large sequence databases, J Chromatogr A, 1348 (2014) 34–46.

[30] H. Buncherd, W. Roseboom, L.J. de Koning, C.G. de Koster, L. de Jong, A gas phase cleavage reaction of cross-linked peptides for protein complex topology studies by peptide fragment fingerprinting from large sequence database, J Proteomics, 108 (2014) 65–77.

[31] L. de Jong, W. Roseboom, G. Kramer, A composite filter for low FDR of protein-protein interactions detected by in vivo cross-linking, J Proteomics, 230 (2021) 103987.

[32] P.T. Kasper, J.W. Back, M. Vitale, A.F. Hartog, W. Roseboom, L.J. de Koning, J.H. van Maarseveen, A.O. Muijsers, C.G. de Koster, L. de Jong, An aptly positioned azido group in the spacer of a protein cross-linker for facile mapping of lysines in close proximity, Chembiochem, 8 (2007)1281–1292.

