## Supplementary figures and images for "Cross-link scrambling in peptide pairs"

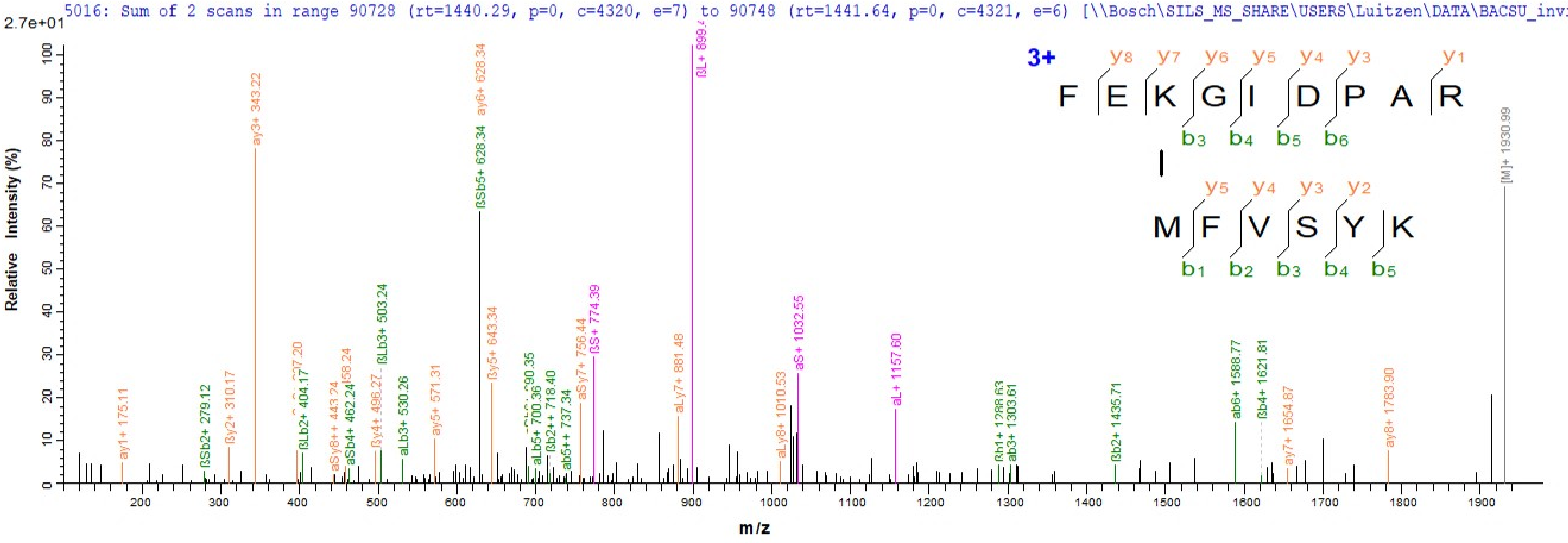

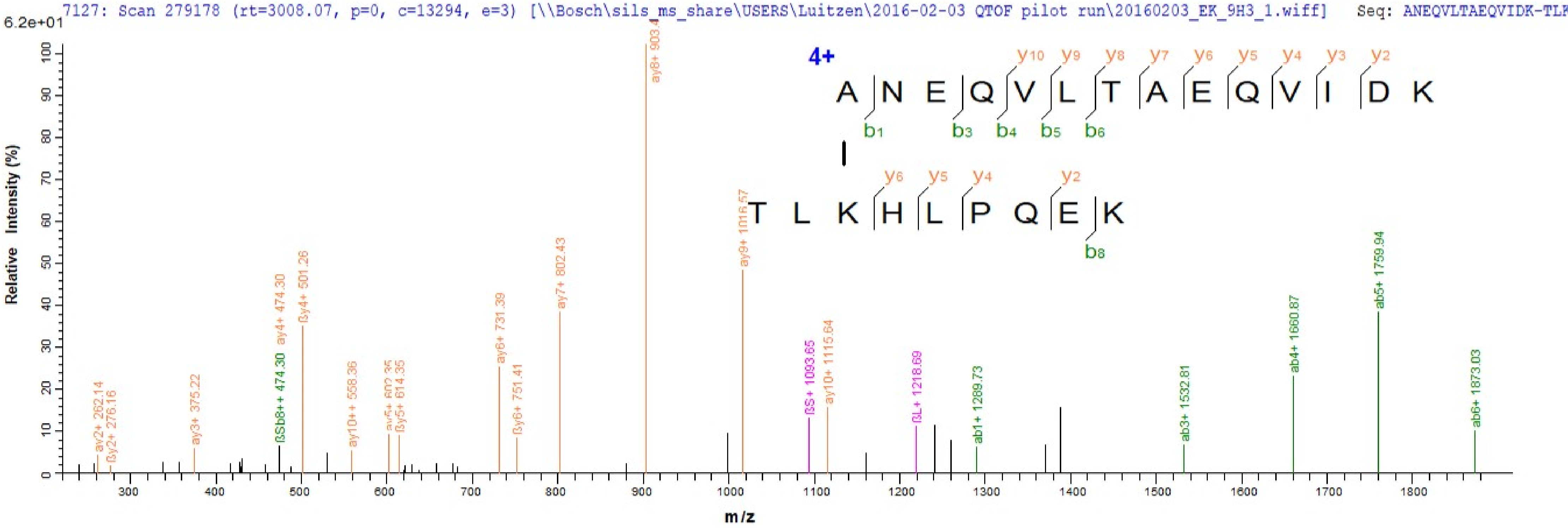

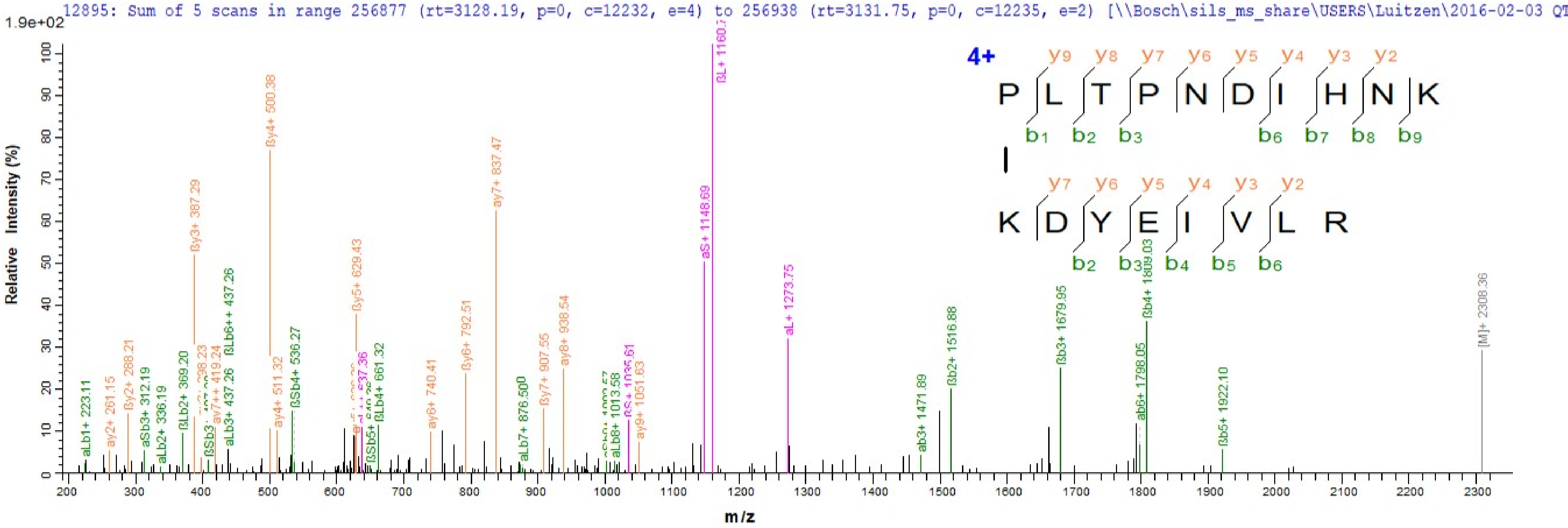

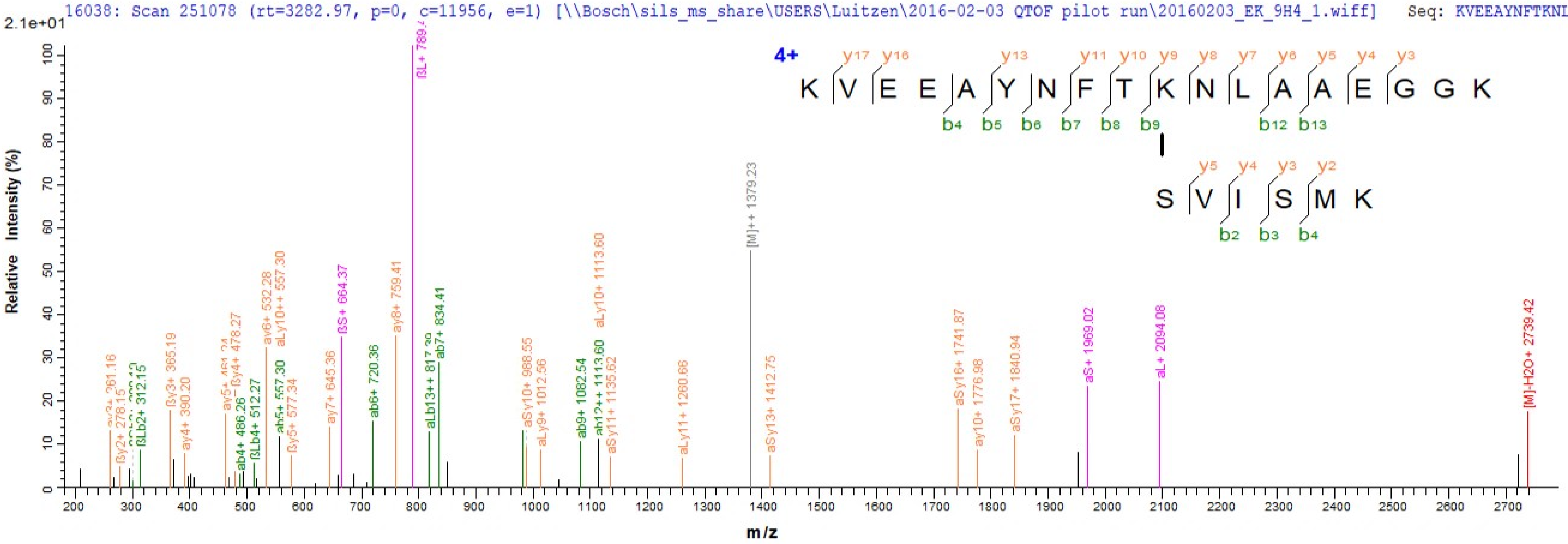

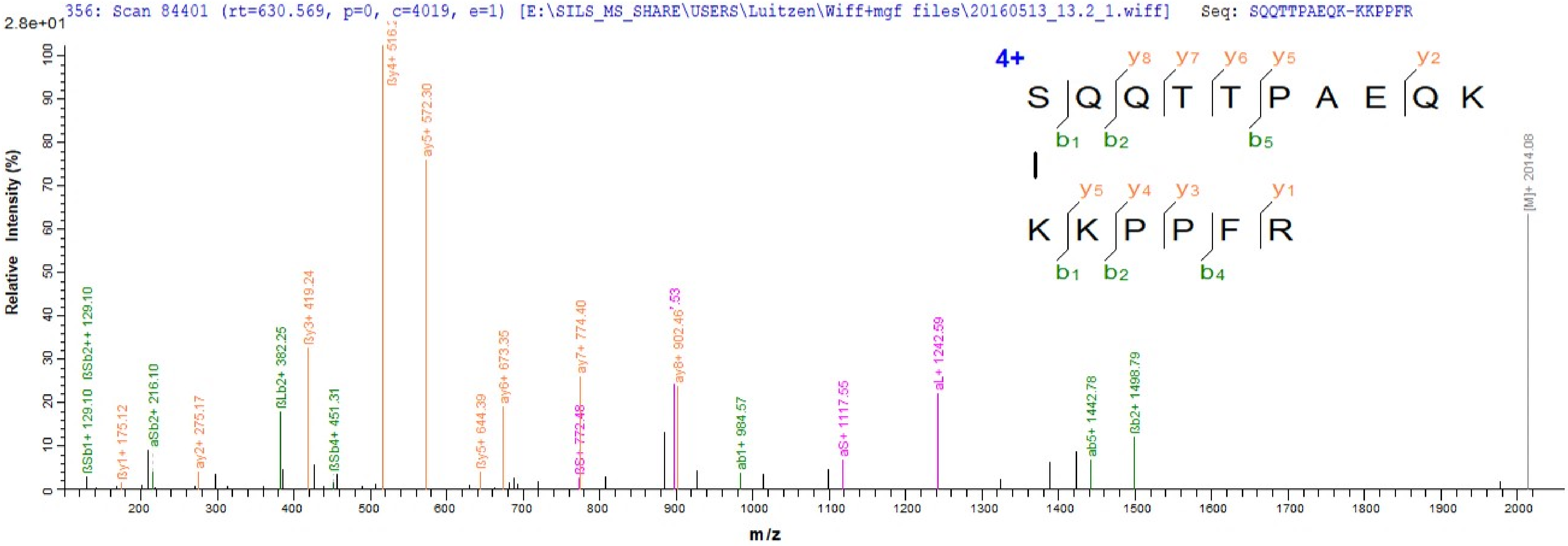

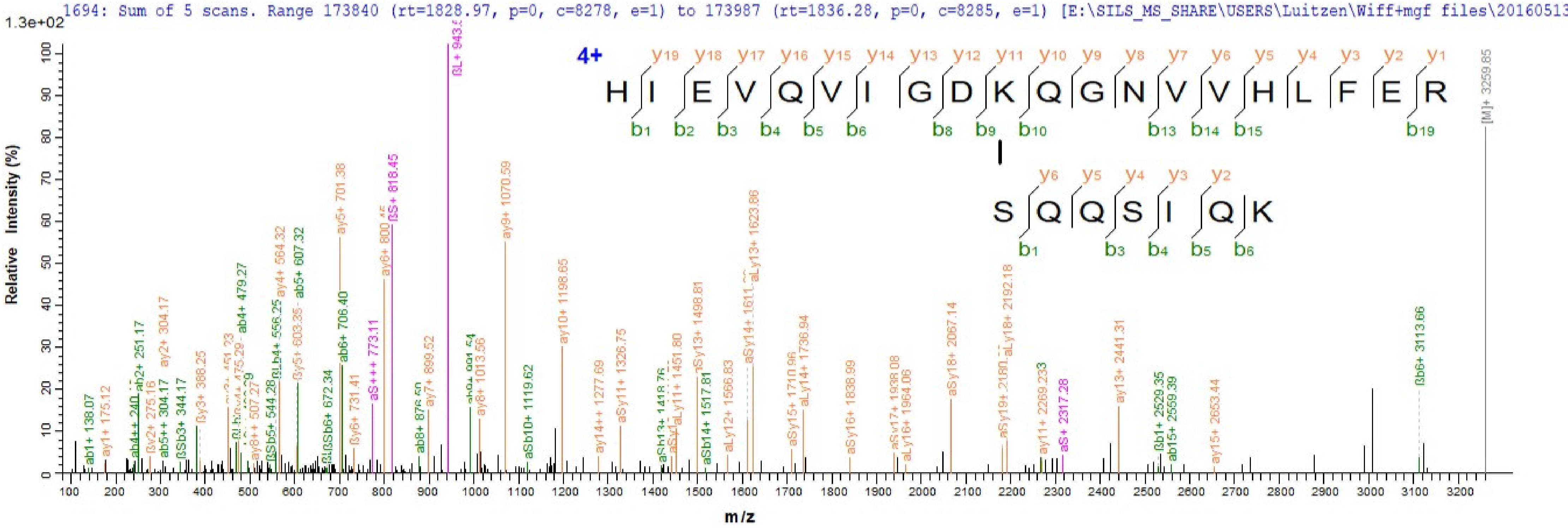

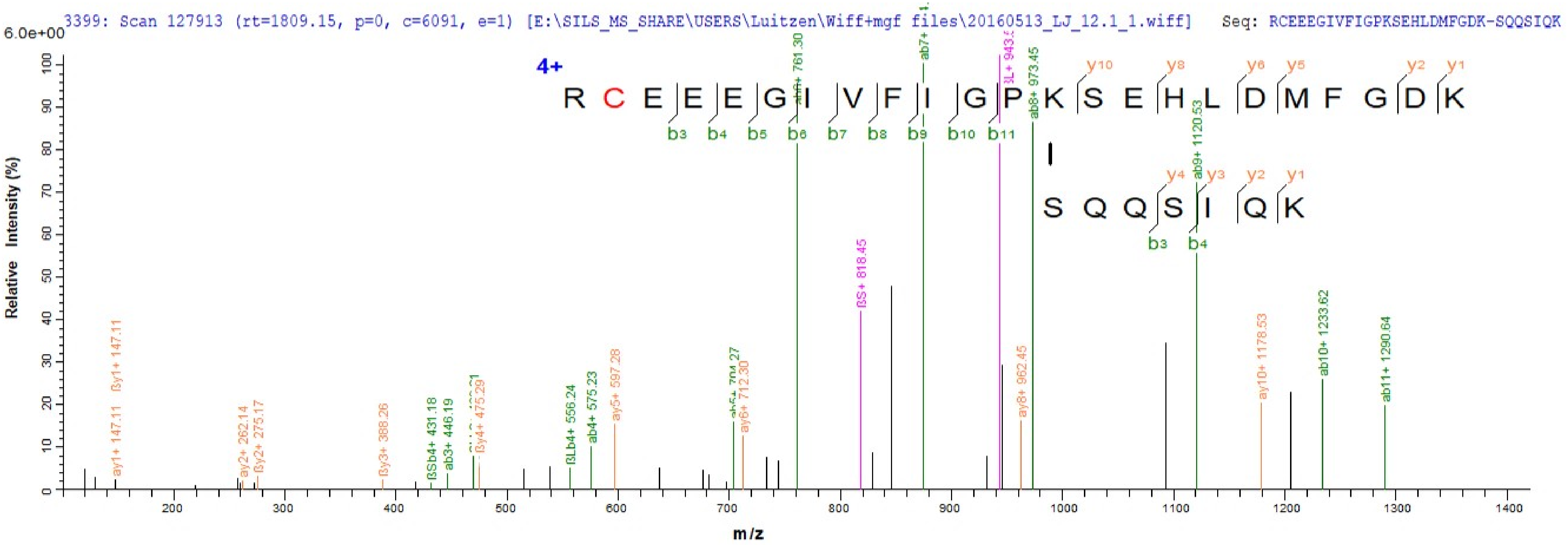

Supplement: supplemental material Spectra[K_Search [file 522012_file02.pdf]
