## supplemental material Spectra_K_Search for "Cross-link scrambling in peptide pairs"

2769: Scan 119786 (rt=1327.41, p=0, c=5704, e=1) [\\Bosch\SILS_MS_SHARE\USERS\Luitzen\Wiff+mgf files\20160513_LJ_10.2_1.wiff]   Seq: TIKNCSVTGLAR-LAIAKQLER   Mod:

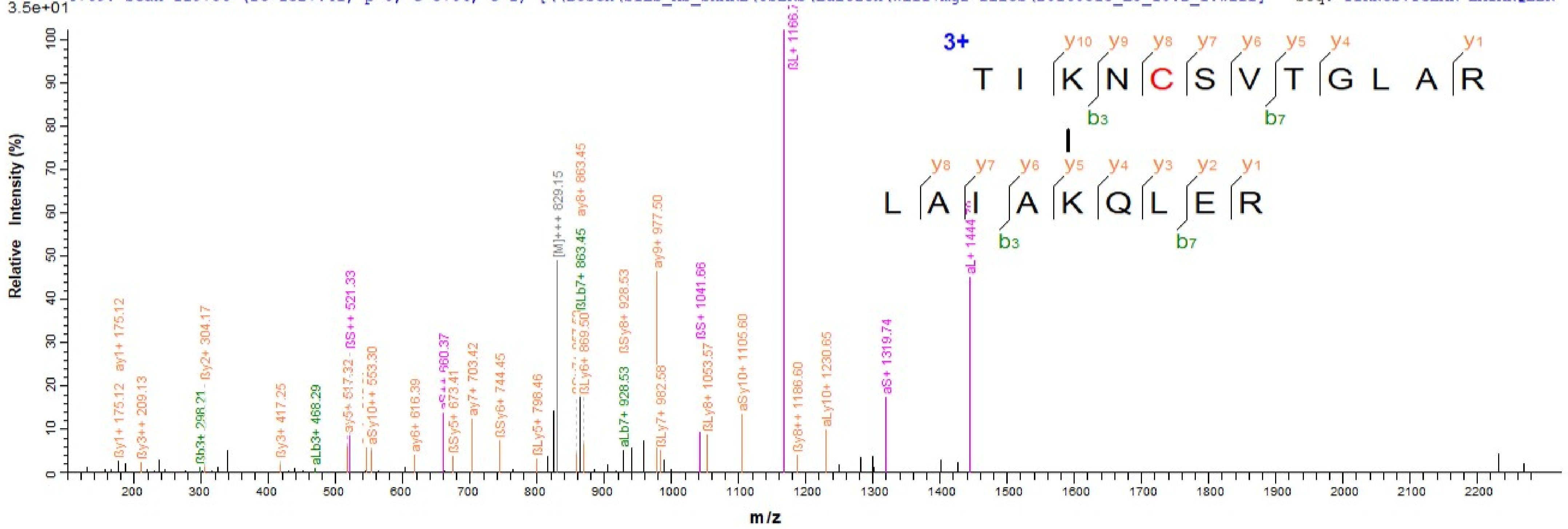

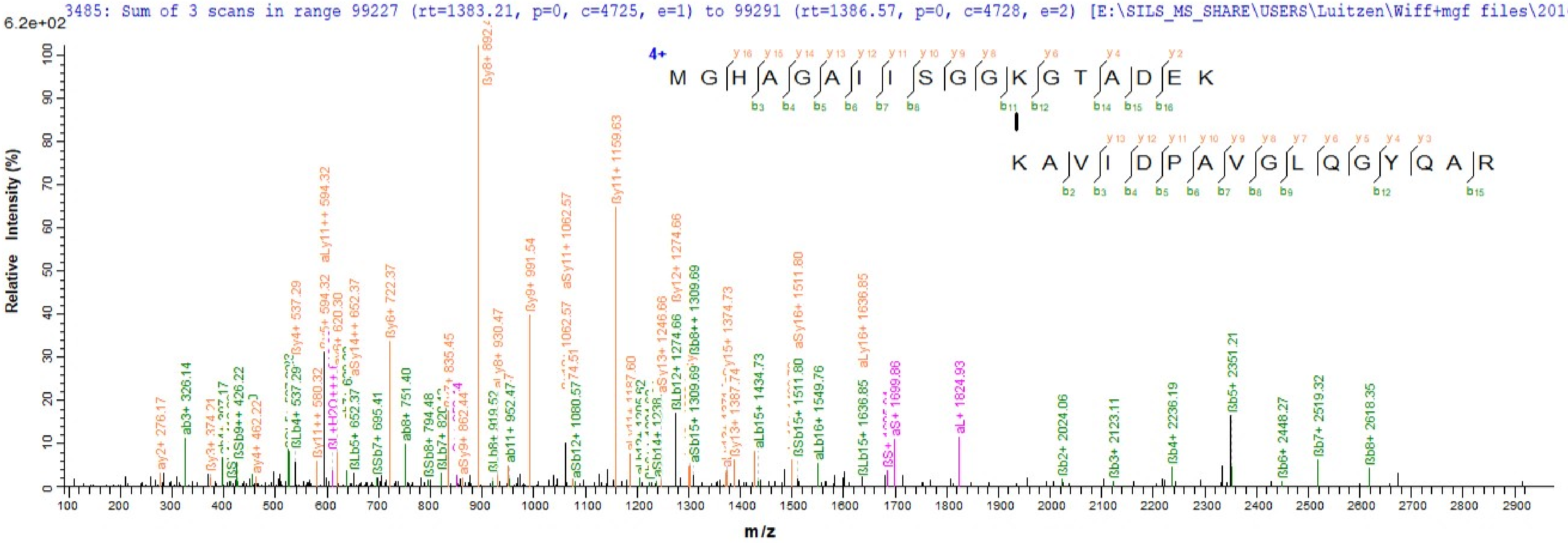
3485: Sum of 3 scans in range 99227 (rt=1383.21, p=0, c=4725, e=1) to 99291 (rt=1386.57, p=0, c=4728, e=2) [E:\SILS_MS_SHARE\USERS\Luitzen\Wiff+mgf files\20160513
6.2e+02
Relative Intensity (%)
m/z
4+
M G H A G A I I S G G K G T A D E K
K A V I D P A V G L Q G Y Q A R
ßy8+ 892.4
ßy11+ 1159.63
aSy11+ 1062.57
ßy9+ 991.54
aLy11++ 594.32
ßy6+ 722.37
ßy12+ 1274.66
aSy16+ 1511.80
aLy16+ 1636.85
ßb5+ 2351.21
aL+ 1824.93
aS+ 1699.86
ßb7+ 2519.32
ßb8+ 2618.35
ßb4+ 2236.19
ßb3+ 2123.11
ßb2+ 2024.06
ßb6+ 2448.27
ab3+ 326.14
ab8+ 751.40
ay2+ 276.17
ay4+ 462.22
aLb15+ 1434.73
aLb16+ 1549.76
ßLb15+ 1636.85
ßSb15+ 1511.80

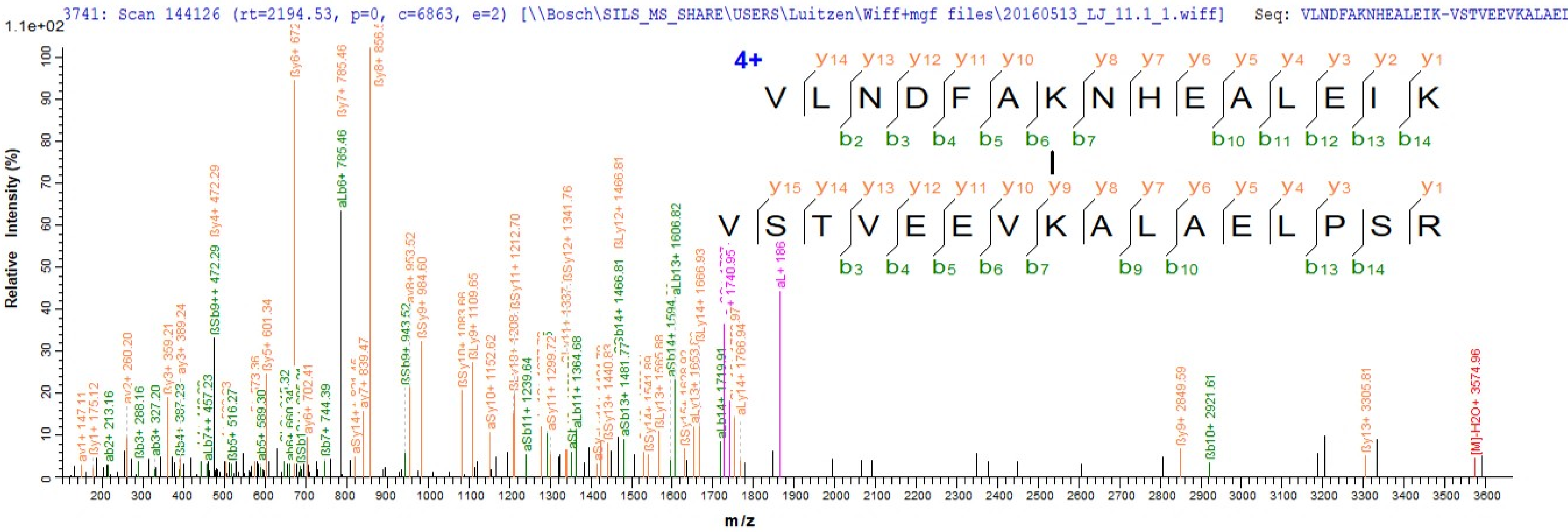

3741: Scan 144126 (rt=2194.53, p=0, c=6863, e=2) [\\Bosch\SILS_MS_SHARE\USERS\Luitzen\Wiff+mgf files\20160513_LJ_11.1_1.wiff] Seq: VLNDFAKNHEALEIK-VSTVEEVKALAEI
1.1e+02
Relative Intensity (%)
m/z
4+
V L N D F A K N H E A L E I K
V S T V E E V K A L A E L P S R
ßy6+ 672
ßy8+ 856.
ßy7+ 785.46
aLb6+ 785.46
ßy4+ 472.29
ßSb9++ 472.29
aLb13+ 1606.82
ßLy12+ 1466.81
ßSy12+ 1341.76
ßSy11+ 1212.70
aL+ 186
[M]-H2O+ 3574.96
ßy13+ 3305.81
ßy9+ 2849.59
ßb10+ 2921.61

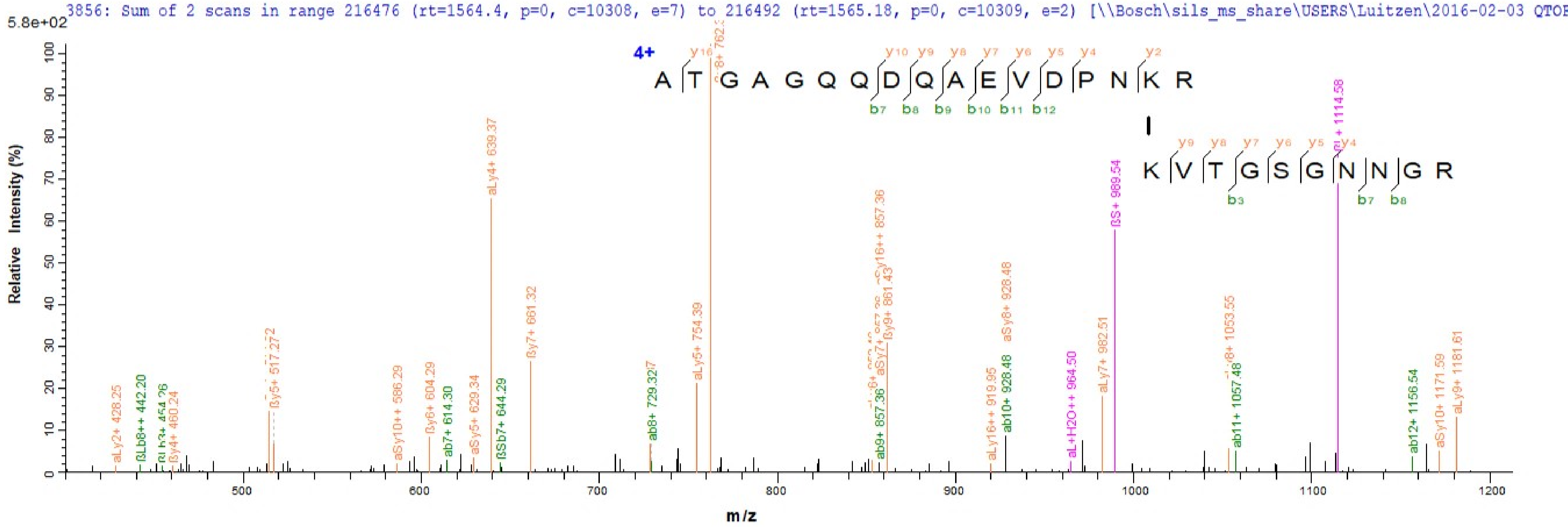
3856: Sum of 2 scans in range 216476 (rt=1564.4, p=0, c=10308, e=7) to 216492 (rt=1565.18, p=0, c=10309, e=2) [\\Bosch\sils_ms_share\USERS\Luitzen\2016-02-03 QTOF
5.8e+02
Relative Intensity (%)
m/z
4+
A T G A G Q Q D Q A E V D P N K R
K V T G S G N N G R
aLy2+ 428.25
ßLb8++ 442.20
ßy4+ 460.24
ßy5+ 517.272
aSy10++ 586.29
ßy6+ 604.29
ab7+ 614.30
aSy5+ 629.34
aLy4+ 639.37
ßSb7+ 644.29
ßy7+ 661.32
ab8+ 729.32
aLy5+ 754.39
ab9+ 857.36
ßy9+ 861.43
aLy16++ 919.95
ab10+ 928.48
aSy8+ 928.48
aL+H2O++ 964.50
aLy7+ 982.51
ßS+ 989.54
ab11+ 1057.48
ab12+ 1156.54
aSy10+ 1171.59
aLy9+ 1181.61

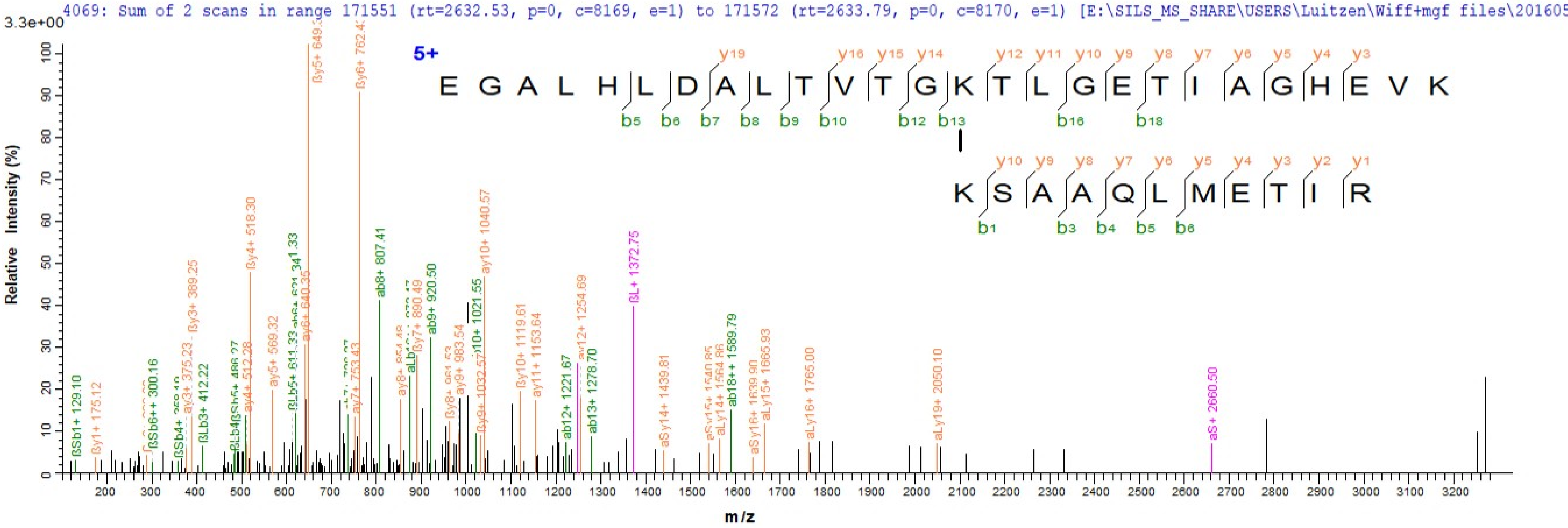
4069: Sum of 2 scans in range 171551 (rt=2632.53, p=0, c=8169, e=1) to 171572 (rt=2633.79, p=0, c=8170, e=1) [E:\SILS_MS_SHARE\USERS\Luitzen\Wiff+mgf files\201605
3.3e+00
Relative Intensity (%)
m/z
5+
E G A L H L D A L T V T G K T L G E T I A G H E V K
y19 y16 y15 y14 y12 y11 y10 y9 y8 y7 y6 y5 y4 y3
b5 b6 b7 b8 b9 b10 b12 b13 b16 b18
K S A A Q L M E T I R
y10 y9 y8 y7 y6 y5 y4 y3 y2 y1
b1 b3 b4 b5 b6
ßy5+ 649.3
ßy6+ 762.43
ßy4+ 518.30
ßy3+ 389.25
ab8+ 807.41
ay10+ 1040.57
ab9+ 920.50
ßy7+ 890.49
ßL+ 1372.75
ay12+ 1254.69
ßy10+ 1119.61
ay11+ 1153.64
ab13+ 1278.70
ab12+ 1221.67
aS+ 2660.50
ay19+ 2050.10
aLy16+ 1765.00
aLy15+ 1665.93
aSy16+ 1639.90
ab18++ 1589.79
aSy14+ 1439.81
ßSb1+ 129.10
ßy1+ 175.12
ßSb6++ 300.16
ßLb3+ 412.22
ay5+ 569.32
ay7+ 753.43
ay9+ 983.54
ay4+ 512.28
ay6+ 640.35
ay3+ 375.23

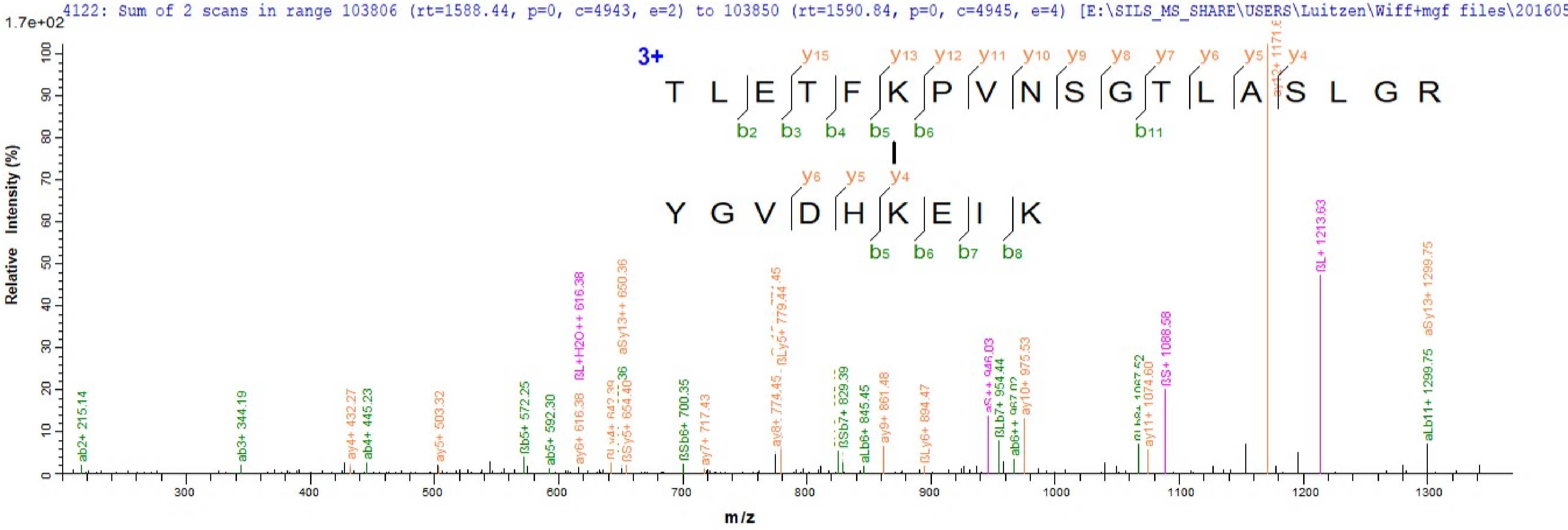

4122: Sum of 2 scans in range 103806 (rt=1588.44, p=0, c=4943, e=2) to 103850 (rt=1590.84, p=0, c=4945, e=4) [E:\SILS_MS_SHARE\USERS\Luitzen\Wiff+mgf files\201605
1.7e+02
3+
T L E T F K P V N S G T L A S L G R
Y G V D H K E I K
Relative Intensity (%)
m/z
ab2+ 215.14
ab3+ 344.19
ay4+ 432.27
ab4+ 445.23
ay5+ 503.32
ßb5+ 572.25
ab5+ 592.30
ßL+H2O++ 616.38
ay6+ 616.38
aSy13++ 650.36
ßSy5+ 654.40
ßSb6+ 700.35
ay7+ 717.43
ay8+ 774.45
ßLy5+ 779.44
ßSb7+ 829.39
aLb6+ 845.45
ay9+ 861.48
ßLy6+ 894.47
ßLb7+ 954.44
ay10+ 975.53
ay11+ 1074.60
ßS+ 1088.58
ßL+ 1213.63
aSy13+ 1299.75
aLb11+ 1299.75

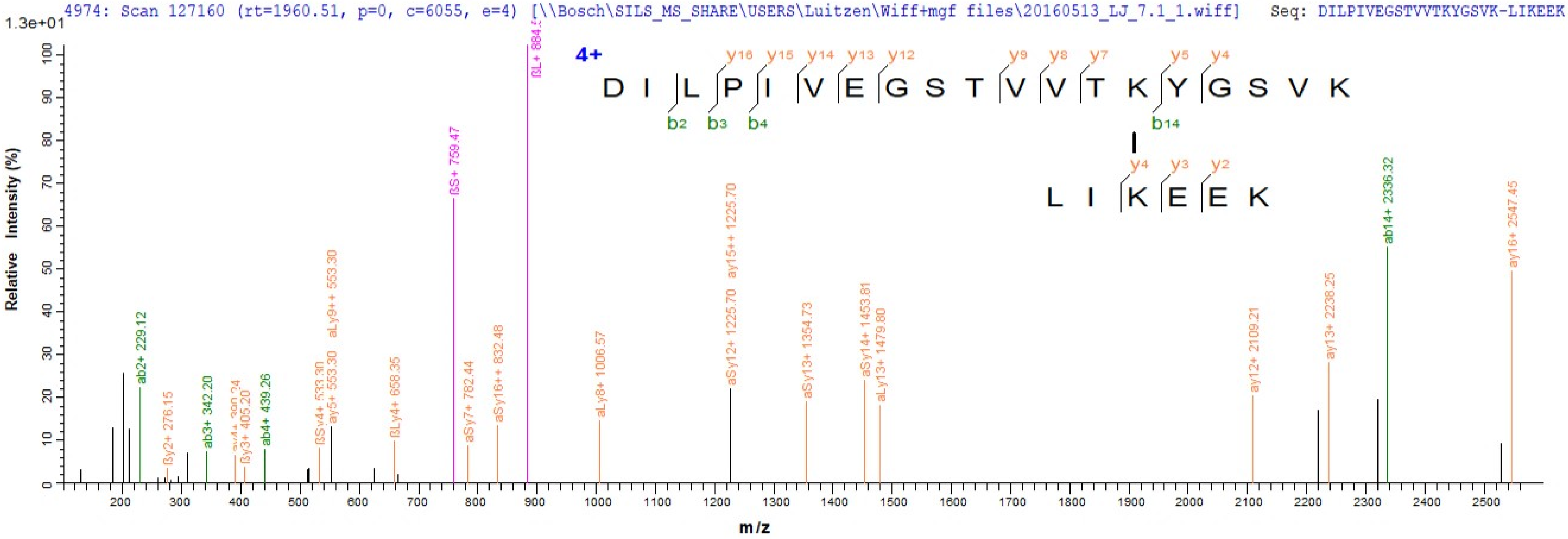

4974: Scan 127160 (rt=1960.51, p=0, c=6055, e=4) [\\Bosch\SILS_MS_SHARE\USERS\Luitzen\Wiff+mgf files\20160513_LJ_7.1_1.wiff] Seq: DILPIVEGSTVVTKYGSVK-LIKEEK
1.3e+01
4+
D I L P I V E G S T V V T K Y G S V K
L I K E E K
Relative Intensity (%)
m/z
ab2+ 229.12
ßy2+ 276.15
ab3+ 342.20
ßy3+ 405.20
ab4+ 439.26
ßSy4+ 533.30
ay5+ 553.30
aLy9++ 553.30
ßLy4+ 658.35
ßS+ 759.47
aSy7+ 782.44
aSy16++ 832.48
aLy8+ 1006.57
aSy12+ 1225.70
ay15++ 1225.70
aSy13+ 1354.73
aSy14+ 1453.81
aLy13+ 1479.80
ay12+ 2109.21
ay13+ 2238.25
ab14+ 2336.32
ay16+ 2547.45

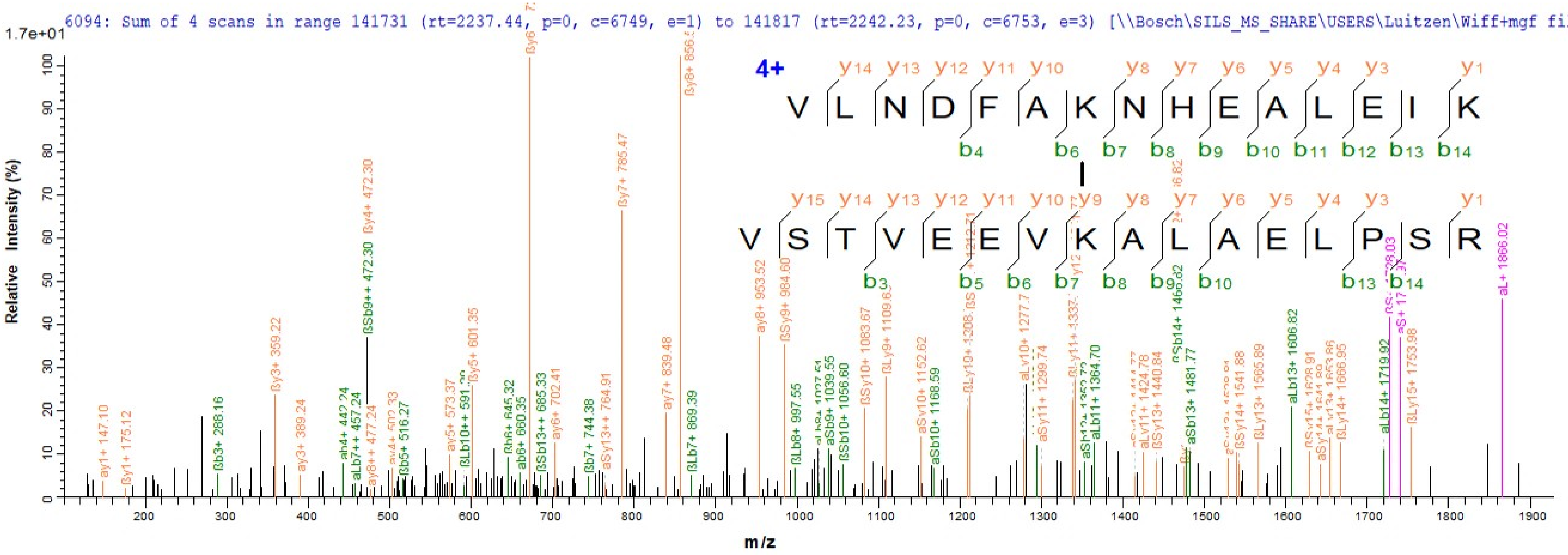
6094: Sum of 4 scans in range 141731 (rt=2237.44, p=0, c=6749, e=1) to 141817 (rt=2242.23, p=0, c=6753, e=3) [\\Bosch\SILS_MS_SHARE\USERS\Luitzen\Wiff+mgf files\2
1.7e+01
4+
V L N D F A K N H E A L E I K
V S T V E E V K A L A E L P S R
Relative Intensity (%)
m/z

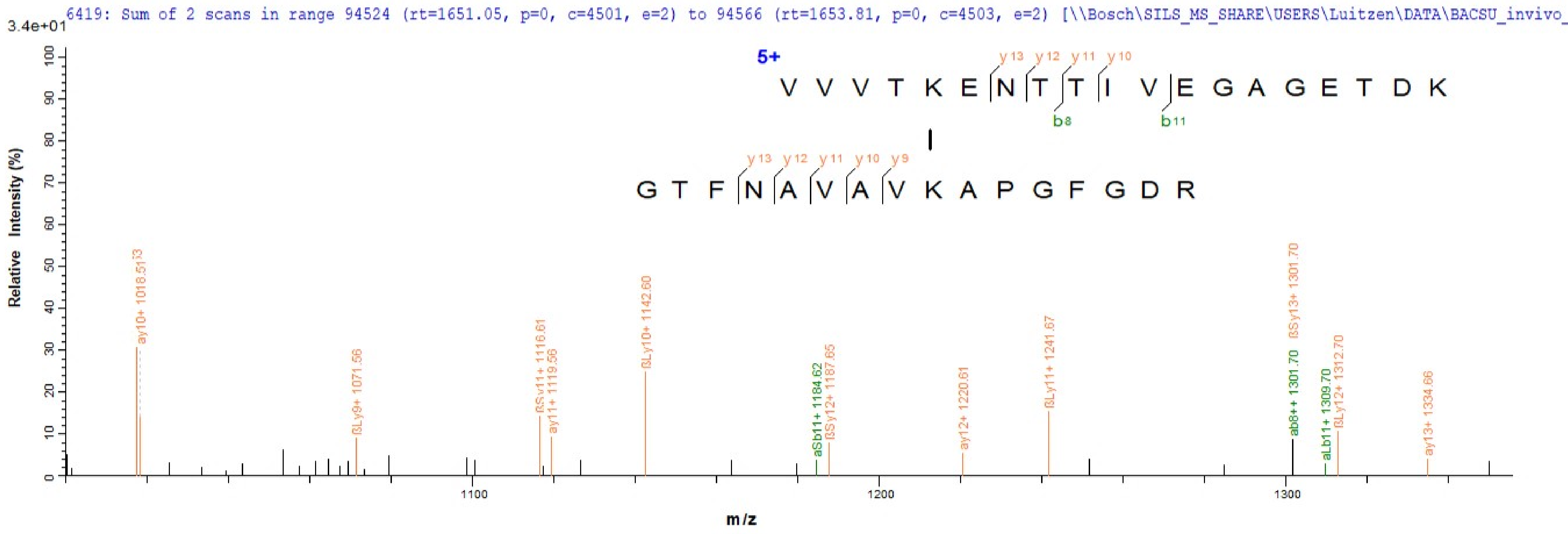
6419: Sum of 2 scans in range 94524 (rt=1651.05, p=0, c=4501, e=2) to 94566 (rt=1653.81, p=0, c=4503, e=2) [\\Bosch\SILS_MS_SHARE\USERS\Luitzen\DATA\BACSU_invivo_
3.4e+01
Relative Intensity (%)
m/z
5+
V V V T K E N T T I V E G A G E T D K
y13 y12 y11 y10
b8 b11
G T F N A V A V K A P G F G D R
y13 y12 y11 y10 y9
ay10+ 1018.513
βLy9+ 1071.56
βSy11+ 1116.61
ay11+ 1119.56
βLy10+ 1142.60
aSb11+ 1184.62
βSy12+ 1187.65
ay12+ 1220.61
βLy11+ 1241.67
ab8++ 1301.70
βSy13+ 1301.70
aLb11+ 1309.70
βLy12+ 1312.70
ay13+ 1334.66
1100
1200
1300

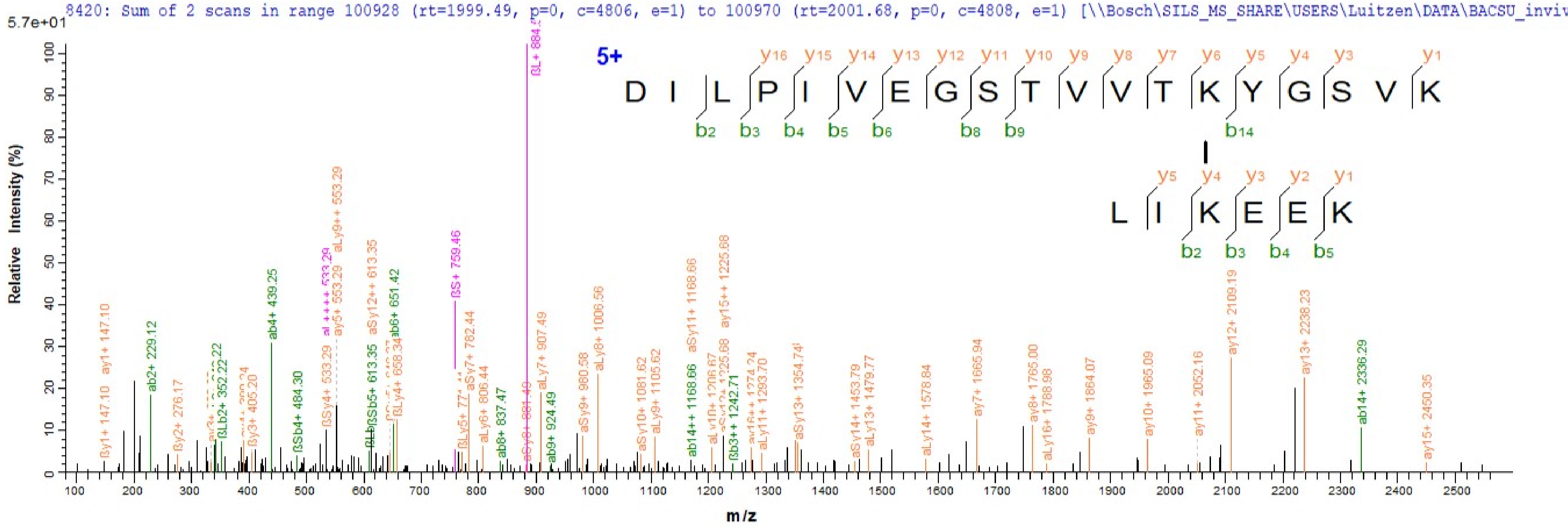
8420: Sum of 2 scans in range 100928 (rt=1999.49, p=0, c=4806, e=1) to 100970 (rt=2001.68, p=0, c=4808, e=1) [\\Bosch\SILS_MS_SHARE\USERS\Luitzen\DATA\BACSU_inviv
5.7e+01
Relative Intensity (%)
m/z
5+
D I L P I V E G S T V V T K Y G S V K
L I K E E K
ab2+ 229.12
ab4+ 439.25
ßSb4+ 484.30
ßS+ 759.46
aLy9++ 553.29
aSy12++ 613.35
ab6+ 651.42
aLy7+ 907.49
aLy8+ 1006.56
aSy11+ 1168.66
ay15++ 1225.68
ay12+ 2109.19
ay13+ 2238.23
ab14+ 2336.29
ay15+ 2450.35
ay7+ 1665.94
ay8+ 1765.00
ay9+ 1864.07
ay10+ 1965.09
ay11+ 2052.16

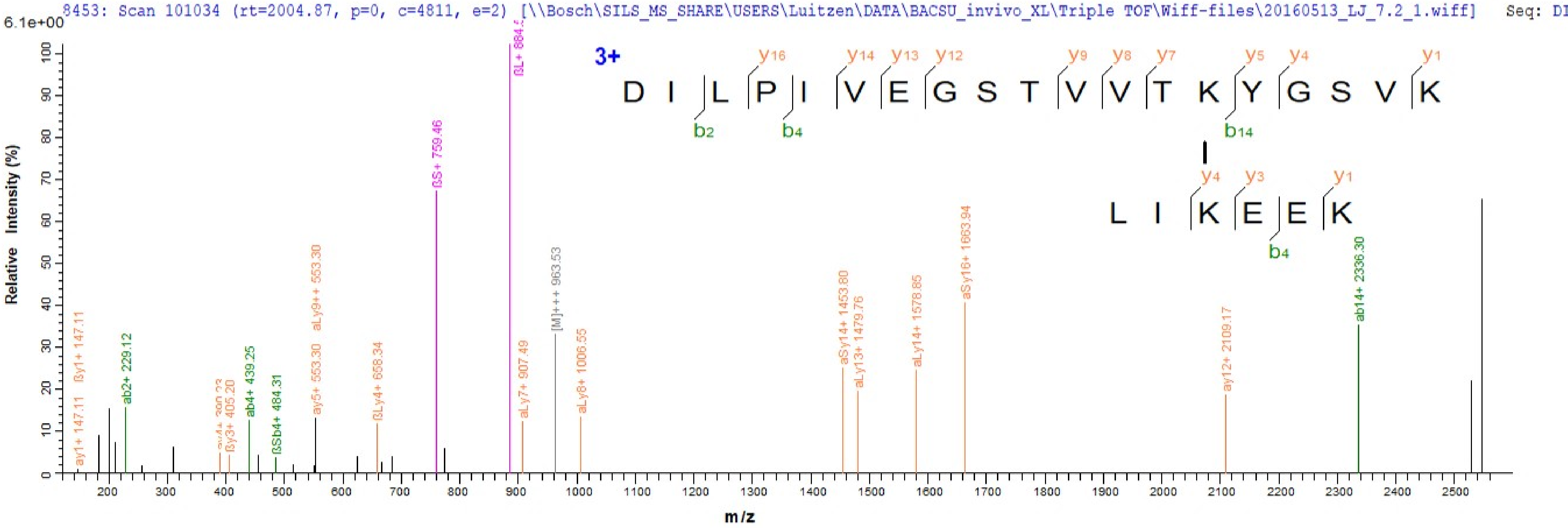
8453: Scan 101034 (rt=2004.87, p=0, c=4811, e=2) [\\Bosch\SILS_MS_SHARE\USERS\Luitzen\DATA\BACSU_invivo_XL\Triple TOF\Wiff-files\20160513_LJ_7.2_1.wiff] Seq: DI
6.1e+00
Relative Intensity (%)
m/z
3+
D I L P I V E G S T V V T K Y G S V K
y16 y14 y13 y12 y9 y8 y7 y5 y4 y1
b2 b4 b14
L I K E E K
y4 y3 y1
b4
ay1+ 147.11
ßy1+ 147.11
ab2+ 229.12
ßy3+ 405.20
ab4+ 439.25
ßSb4+ 484.31
ay5+ 553.30
aLy9++ 553.30
ßLy4+ 658.34
ßS+ 759.46
aLy7+ 907.49
[M]+++ 963.53
aLy8+ 1006.55
aSy14+ 1453.80
aLy13+ 1479.76
aLy14+ 1578.85
aSy16+ 1663.94
ay12+ 2109.17
ab14+ 2336.30

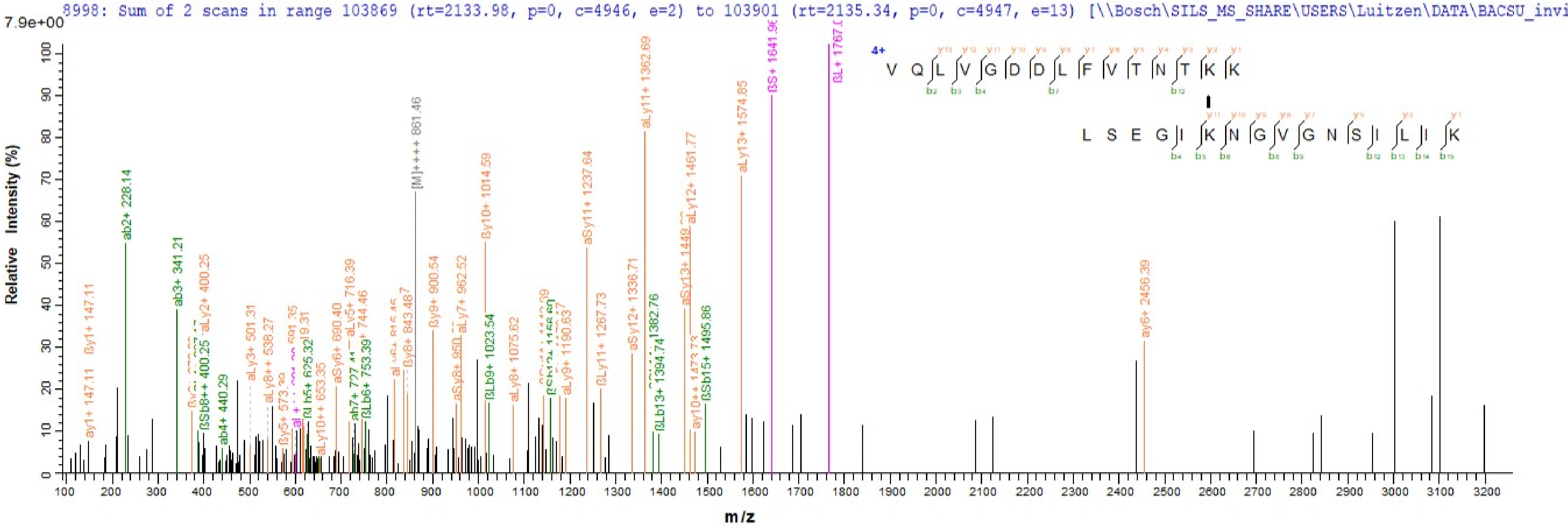
8998: Sum of 2 scans in range 103869 (rt=2133.98, p=0, c=4946, e=2) to 103901 (rt=2135.34, p=0, c=4947, e=13) [\\Bosch\SILS_MS_SHARE\USERS\Luitzen\DATA\BACSU_invi
7.9e+00
Relative Intensity (%)
m/z
4+
V Q L V G D D L F V T N T K K
L S E G I K N G V G N S I L I K
[M]++++ 861.46
aLy11+ 1362.69
aLy13+ 1574.85
ab2+ 228.14
ab3+ 341.21
ay6+ 2456.39

10788: Scan 294611 (rt=3715.93, p=0, c=14029, e=1) [\\Bosch\sils_ms_share\USERS\Luitzen\2016-02-03 QTOF pilot run\20160203_EK_9H3_1.wiff] Seq: LASEGYKEITLLGQNVN

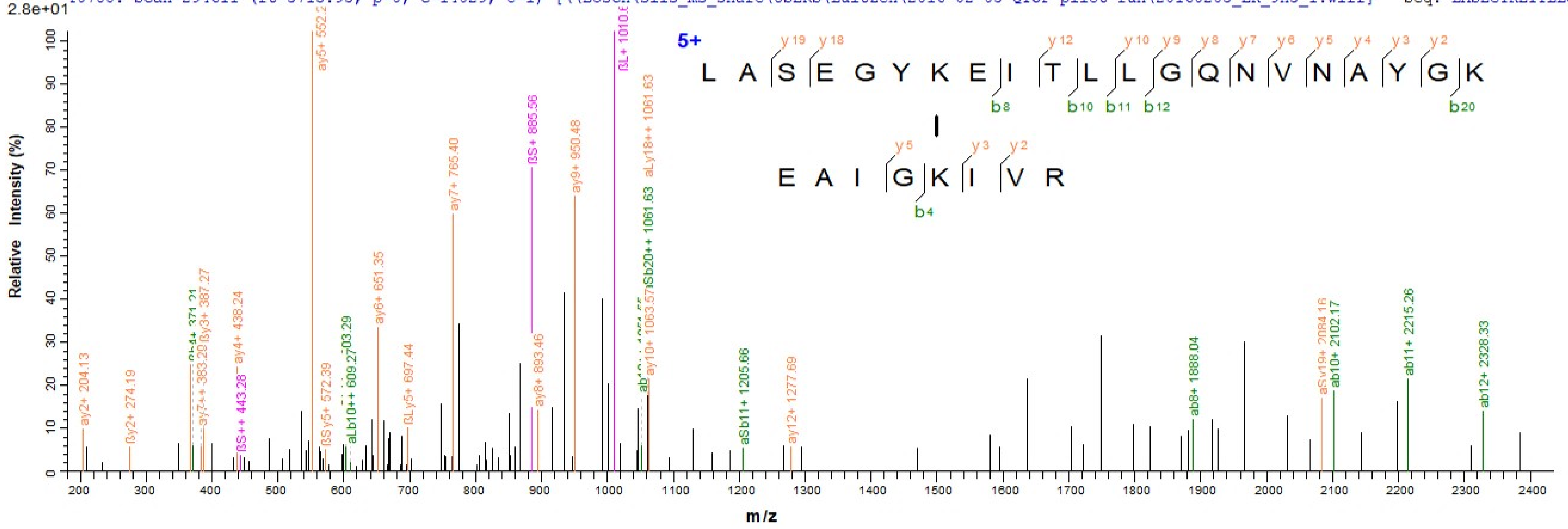

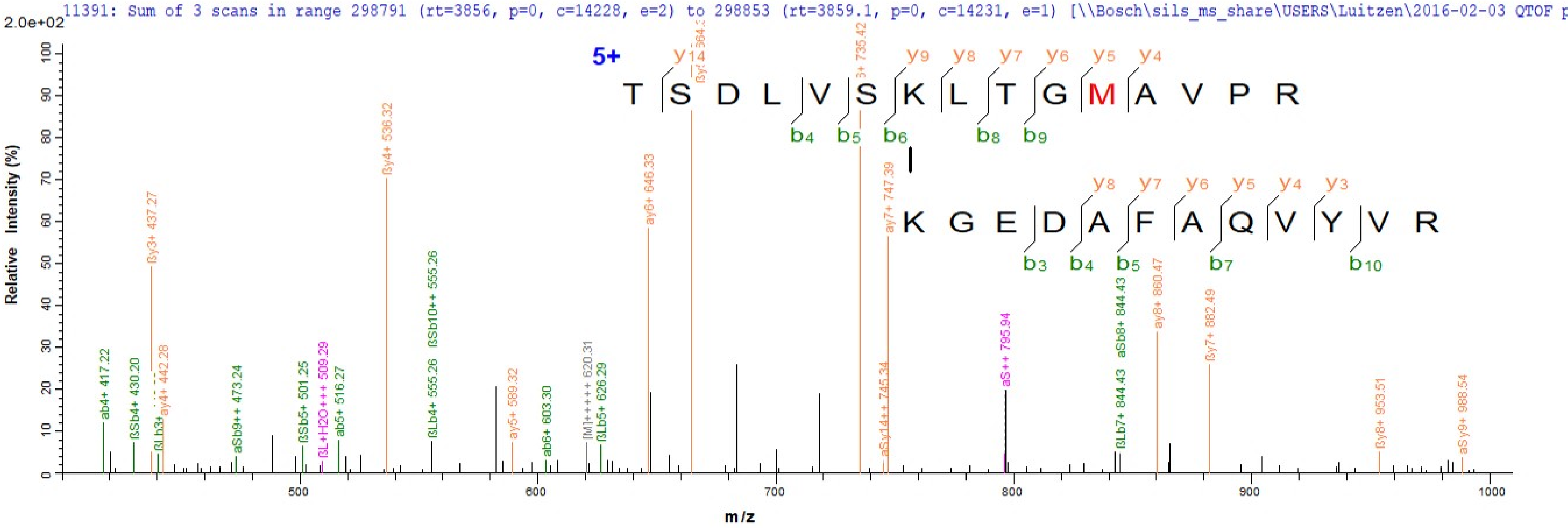
11391: Sum of 3 scans in range 298791 (rt=3856, p=0, c=14228, e=2) to 298853 (rt=3859.1, p=0, c=14231, e=1) [\\Bosch\sils_ms_share\USERS\Luitzen\2016-02-03 QTOF p
2.0e+02
5+
T S D L V S K L T G M A V P R
y14 y9 y8 y7 y6 y5 y4
b4 b5 b6 b8 b9
K G E D A F A Q V Y V R
y8 y7 y6 y5 y4 y3
b3 b4 b5 b7 b10
Relative Intensity (%)
m/z
ab4+ 417.22
ßSb4+ 430.20
ßy3+ 437.27
ay4+ 442.28
aSb9++ 473.24
ßSb5+ 501.25
ßL+H2O+++ 509.29
ab5+ 516.27
ßy4+ 536.32
ßSb10++ 555.26
ßLb4+ 555.26
ay5+ 589.32
ab6+ 603.30
[M]+++++ 620.31
ßLb5+ 626.29
ay6+ 646.33
ay7+ 747.39
aSy14++ 745.34
aS++ 795.94
aSb8+ 844.43
ßLb7+ 844.43
ay8+ 860.47
ßy7+ 882.49
ßy8+ 953.51
aSy9+ 988.54
0 10 20 30 40 50 60 70 80 90 100
500 600 700 800 900 1000

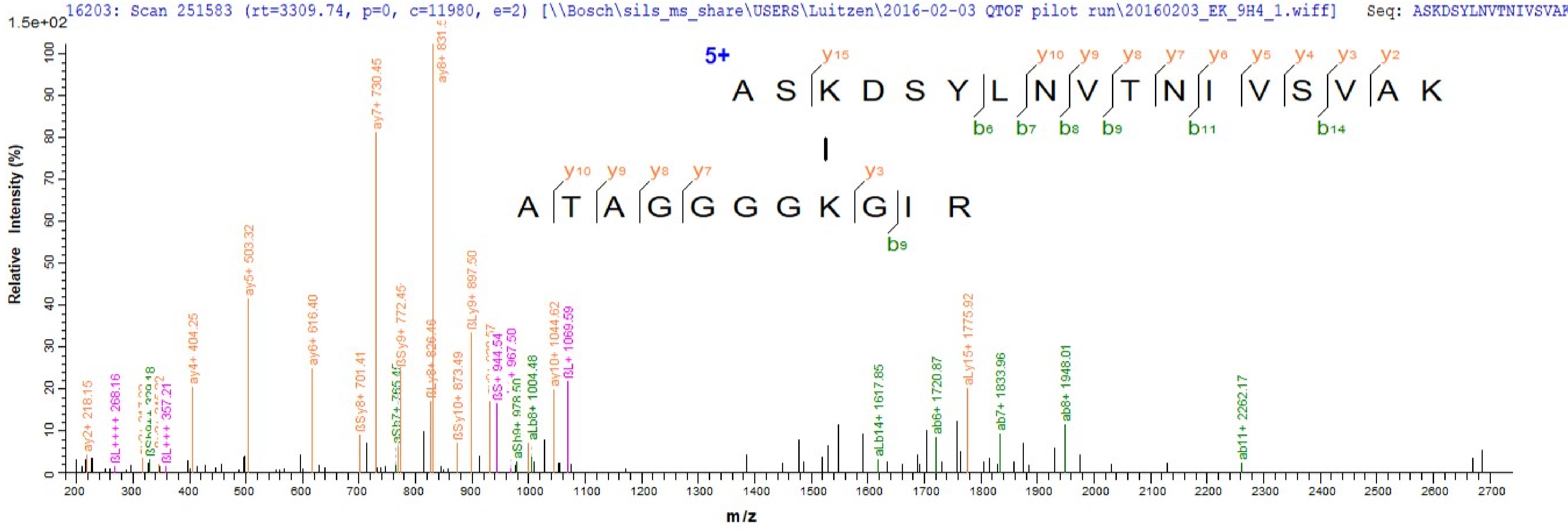
16203: Scan 251583 (rt=3309.74, p=0, c=11980, e=2) [\\Bosch\sils_ms_share\USERS\Luitzen\2016-02-03 QTOF pilot run\20160203_EK_9H4_1.wiff] Seq: ASKDSYLNVTNIVSVAK
1.5e+02
5+
A S K D S Y L N V T N I V S V A K
y15 y10 y9 y8 y7 y6 y5 y4 y3 y2
b6 b7 b8 b9 b11 b14
A T A G G G G K G I R
y10 y9 y8 y7 y3
b9
Relative Intensity (%)
m/z
ay2+ 218.15
ßL++++ 268.16
ßSh9++ 329.18
ßL+++ 357.21
ay4+ 404.25
ay5+ 503.32
ay6+ 616.40
ßSy8+ 701.41
ay7+ 730.45
ßSy9+ 772.45
ßLy8+ 826.46
ßSy10+ 873.49
ßLy9+ 897.50
ßS+ 944.54
aSh9+ 978.50
aLb8+ 1004.48
ay10+ 1044.62
ßL+ 1069.59
aLb14+ 1617.85
ab6+ 1720.87
aLy15+ 1775.92
ab7+ 1833.96
ab8+ 1948.01
ab11+ 2262.17

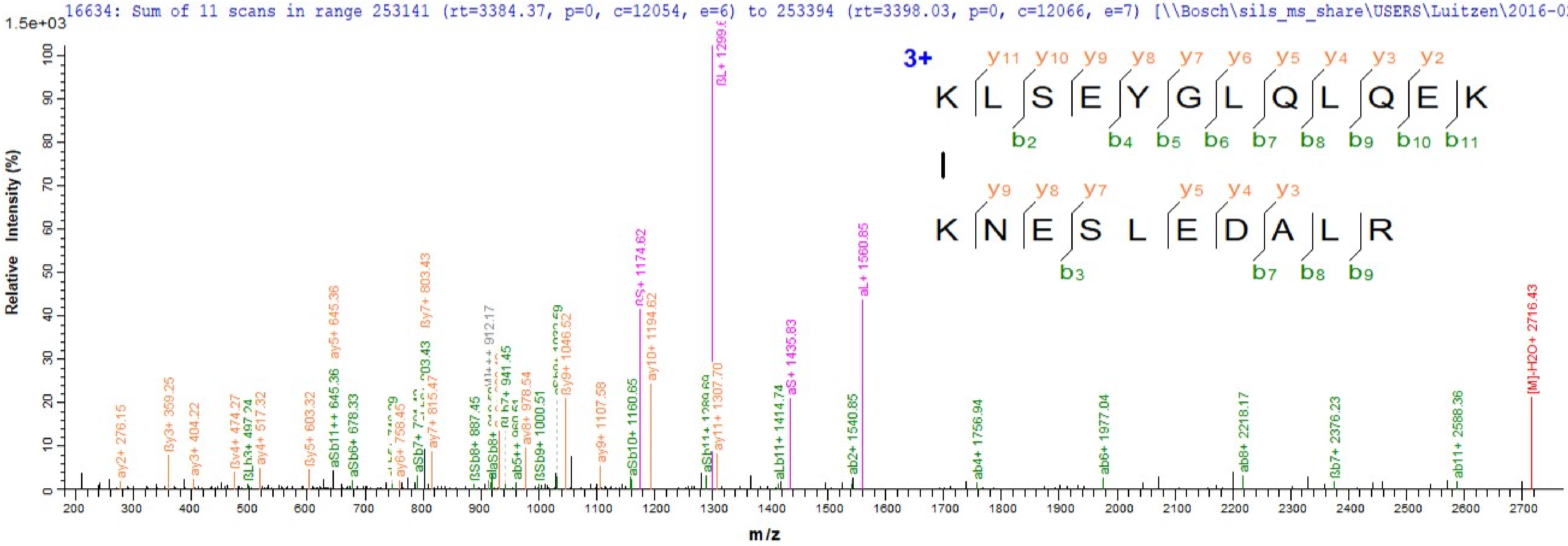
16634: Sum of 11 scans in range 253141 (rt=3384.37, p=0, c=12054, e=6) to 253394 (rt=3398.03, p=0, c=12066, e=7) [\\Bosch\sils_ms_share\USERS\Luitzen\2016-02-03
1.5e+03
Relative Intensity (%)
m/z
3+
K L S E Y G L Q L Q E K
y11 y10 y9 y8 y7 y6 y5 y4 y3 y2
b2 b4 b5 b6 b7 b8 b9 b10 b11
K N E S L E D A L R
y9 y8 y7 y5 y4 y3
b3 b7 b8 b9
ay2+ 276.15
ßy3+ 359.25
ay3+ 404.22
ßy4+ 474.27
ßLb3+ 497.24
ay4+ 517.32
ßy5+ 603.32
aSb11++ 645.36
ay5+ 645.36
aSb6+ 678.33
ay6+ 758.45
ay7+ 815.47
ßy7+ 803.43
ßSb8+ 887.45
ab5++ 960.51
ay8+ 978.54
ßSb9+ 1000.51
ßy9+ 1046.52
ay9+ 1107.58
aSb10+ 1160.65
ßS+ 1174.62
ay10+ 1194.62
ay11+ 1307.70
aLb11+ 1414.74
aS+ 1435.83
ab2+ 1540.85
aL+ 1560.85
ab4+ 1756.94
ab6+ 1977.04
ab8+ 2218.17
ßb7+ 2376.23
ab11+ 2588.36
[M]-H2O+ 2716.43

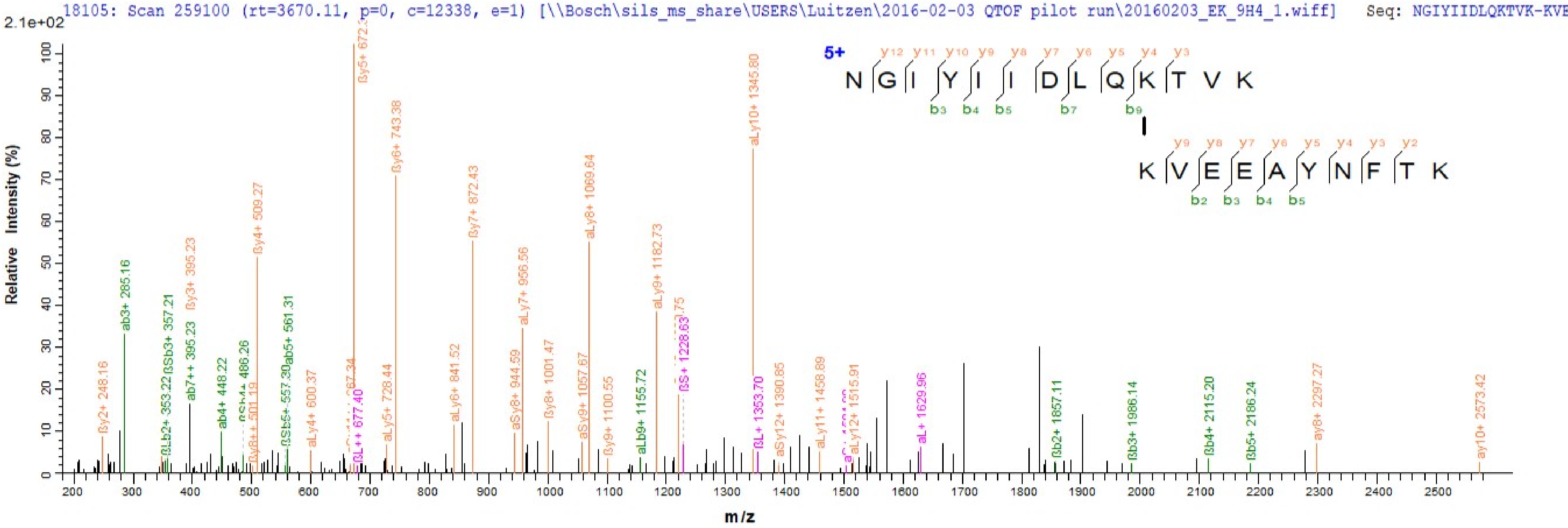
18105: Scan 259100 (rt=3670.11, p=0, c=12338, e=1) [\\Bosch\sils_ms_share\USERS\Luitzen\2016-02-03 QTOF pilot run\20160203_EK_9H4_1.wiff] Seq: NGIYIIDLQKTVK-KVE
2.1e+02
Relative Intensity (%)
m/z
5+
NGIYIIDLQKTVK
KVEEAYNFTK
ßy5+ 672
ßy6+ 743.38
ßy7+ 872.43
aLy8+ 1069.64
aLy10+ 1345.80
ßy4+ 509.27
ßy3+ 395.23
ab3+ 285.16
ßSb3+ 357.21
ab7++ 395.23
aLy9+ 1182.73
aLy7+ 956.56
ab5+ 561.31
ßy2+ 248.16
ab4+ 448.22
aLy4+ 600.37
aLy5+ 728.44
aLy6+ 841.52
aSy8+ 944.59
ßy8+ 1001.47
aSy9+ 1057.67
ßy9+ 1100.55
aLb9+ 1155.72
ßS+ 1228.63
ßL+ 1353.70
aSy12+ 1390.85
aLy11+ 1458.89
aLy12+ 1515.91
aL+ 1629.96
ßb2+ 1857.11
ßb3+ 1986.14
ßb4+ 2115.20
ßb5+ 2186.24
ay8+ 2297.27
ay10+ 2573.42
ßL++ 677.40

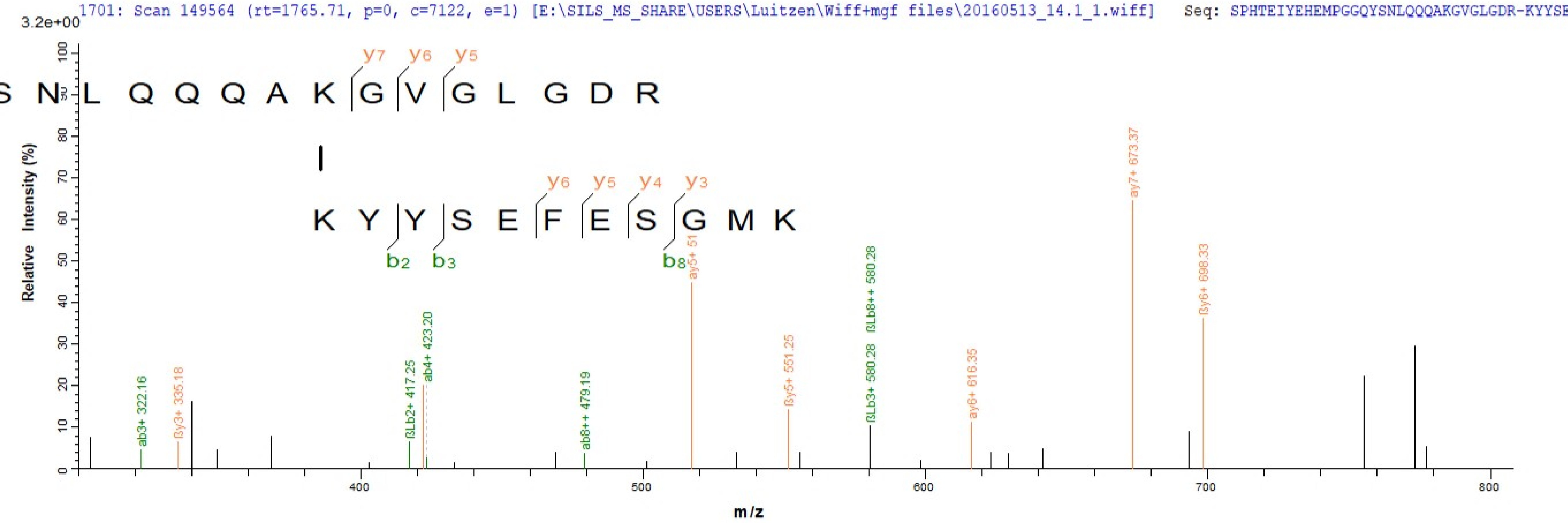
1701: Scan 149564 (rt=1765.71, p=0, c=7122, e=1) [E:\SILS_MS_SHARE\USERS\Luitzen\Wiff+mgf files\20160513_14.1_1.wiff] Seq: SPHTEIYEHEMPGGQYSNLQQQAKGVGLGDR-KYYSE
3.2e+00
Relative Intensity (%)
S N L Q Q Q A K G V G L G D R
y7 y6 y5
K Y Y S E F E S G M K
y6 y5 y4 y3
b2 b3 b8
ab3+ 322.16
ßy3+ 335.18
ßLb2+ 417.25
ab4+ 423.20
ab8++ 479.19
ay5+ 51
ßy5+ 551.25
ßLb3+ 580.28
ßLb8++ 580.28
ay6+ 616.35
ay7+ 673.37
ßy6+ 698.33
m/z
