## supplemental material Spectra_KTSY_Search for "Cross-link scrambling in peptide pairs"

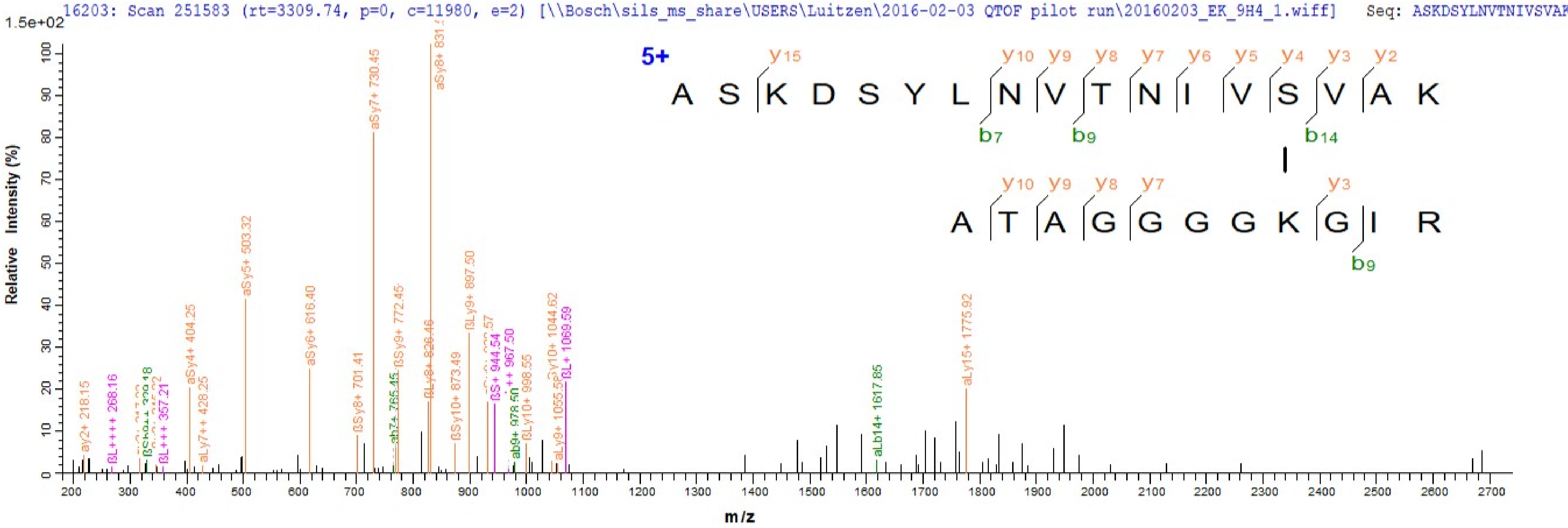
16203: Scan 251583 (rt=3309.74, p=0, c=11980, e=2) [\\Bosch\sils_ms_share\USERS\Luitzen\2016-02-03 QTOF pilot run\20160203_EK_9H4_1.wiff] Seq: ASKDSYLNVTNIVSVAK
1.5e+02
Relative Intensity (%)
m/z
5+
A S K D S Y L N V T N I V S V A K
y15 y10 y9 y8 y7 y6 y5 y4 y3 y2
b7 b9 b14
A T A G G G G K G I R
y10 y9 y8 y7 y3
b9
ay2+ 218.15
ßL++++ 268.16
ßL+++ 357.21
aSy4+ 404.25
aLy7++ 428.25
aSy5+ 503.32
aSy6+ 616.40
ßSy8+ 701.41
aSy7+ 730.45
ßSy9+ 772.45
ßLy8+ 826.46
ßSy10+ 873.49
ßLy9+ 897.50
ßS+ 944.54
ab9+ 978.50
ßLy10+ 998.55
aLy9+ 1055.56
ßL+ 1069.59
aLb14+ 1617.85
aLy15+ 1775.92

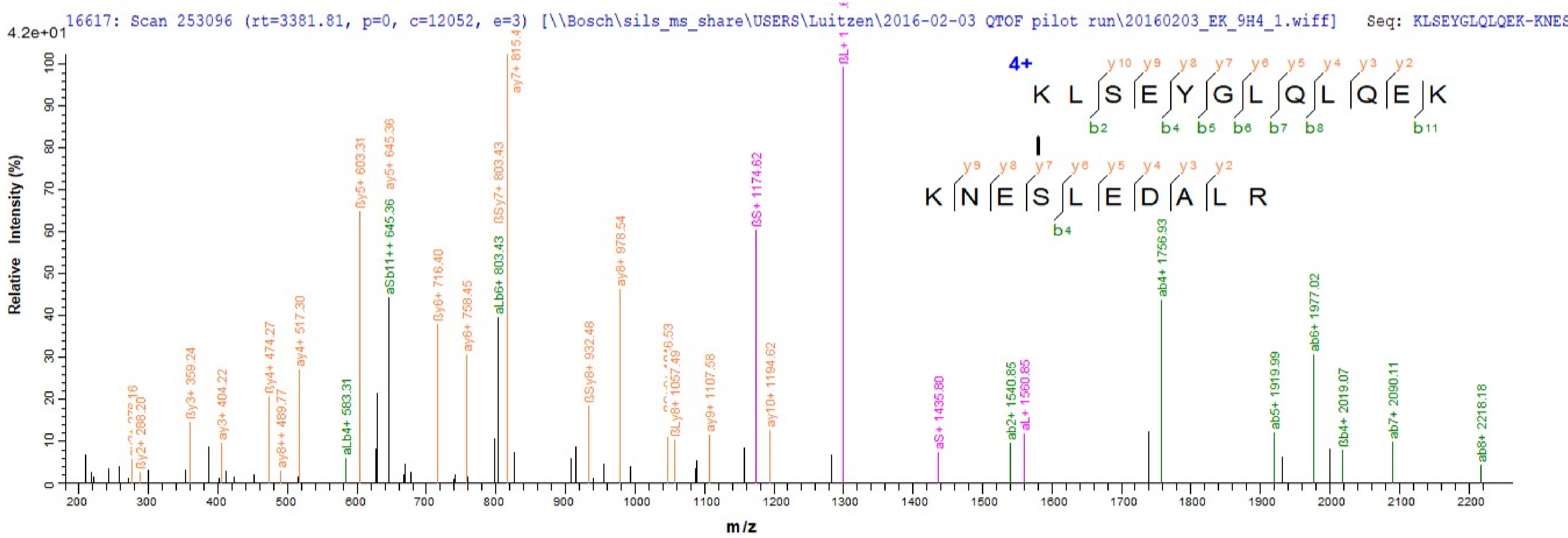
16617: Scan 253096 (rt=3381.81, p=0, c=12052, e=3) [\\Bosch\sils_ms_share\USERS\Luitzen\2016-02-03 QTOF pilot run\20160203_EK_9H4_1.wiff] Seq: KLSEYGLQLQEK-KNES
4.2e+01
Relative Intensity (%)
m/z
4+
KLSEYGLQLQEK
KNESLEDALR
y10 y9 y8 y7 y6 y5 y4 y3 y2
b2 b4 b5 b6 b7 b8 b11
y9 y8 y7 y6 y5 y4 y3 y2
b4
ßy2+ 288.20
ßy3+ 359.24
ay3+ 404.22
ßy4+ 474.27
ay8++ 489.77
ay4+ 517.30
aLb4+ 583.31
ßy5+ 603.31
ay5+ 645.36
aSb11++ 645.36
ßy6+ 716.40
ay6+ 758.45
aLb6+ 803.43
ßSy7+ 803.43
ßSy8+ 932.48
ay8+ 978.54
ßLy8+ 1057.49
ay9+ 1107.58
ßS+ 1174.62
ay10+ 1194.62
aS+ 1435.80
ab2+ 1540.85
aL+ 1560.85
ab4+ 1756.93
ab5+ 1919.99
ab6+ 1977.02
ßb4+ 2019.07
ab7+ 2090.11
ab8+ 2218.18

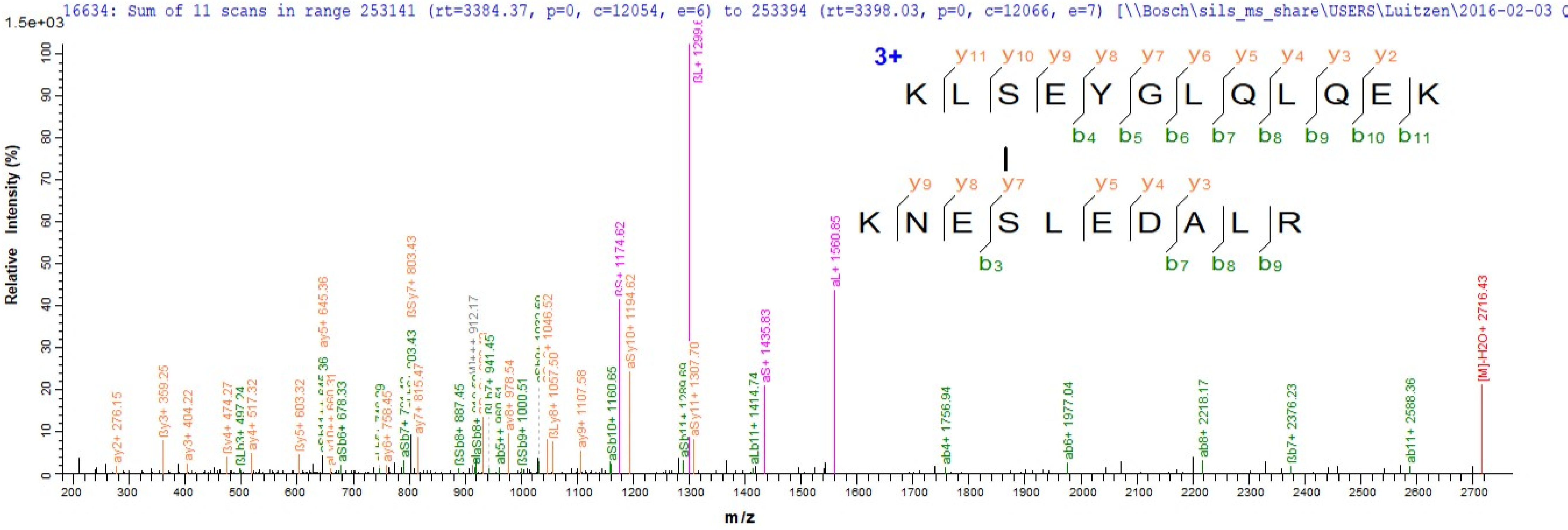
16634: Sum of 11 scans in range 253141 (rt=3384.37, p=0, c=12054, e=6) to 253394 (rt=3398.03, p=0, c=12066, e=7) [\\Bosch\sils_ms_share\USERS\Luitzen\2016-02-03
1.5e+03
Relative Intensity (%)
m/z
3+
K L S E Y G L Q L Q E K
y11 y10 y9 y8 y7 y6 y5 y4 y3 y2
b4 b5 b6 b7 b8 b9 b10 b11
K N E S L E D A L R
y9 y8 y7 y5 y4 y3
b3 b7 b8 b9
ay2+ 276.15
ßy3+ 359.25
ay3+ 404.22
ßy4+ 474.27
ßLb3+ 497.24
ay4+ 517.32
ßy5+ 603.32
ay5+ 645.36
aSb6+ 678.33
ay6+ 758.45
ßSy7+ 803.43
ay7+ 815.47
ßSb8+ 887.45
912.17
941.45
ay8+ 978.54
ßSb9+ 1000.51
1046.52
ßLy8+ 1057.50
ay9+ 1107.58
aSb10+ 1160.65
ßS+ 1174.62
aSy10+ 1194.62
ßL+ 1299.6
aSy11+ 1307.70
aLb11+ 1414.74
aS+ 1435.83
aL+ 1560.85
ab4+ 1756.94
ab6+ 1977.04
ab8+ 2218.17
ßb7+ 2376.23
ab11+ 2588.36
[M]-H2O+ 2716.43

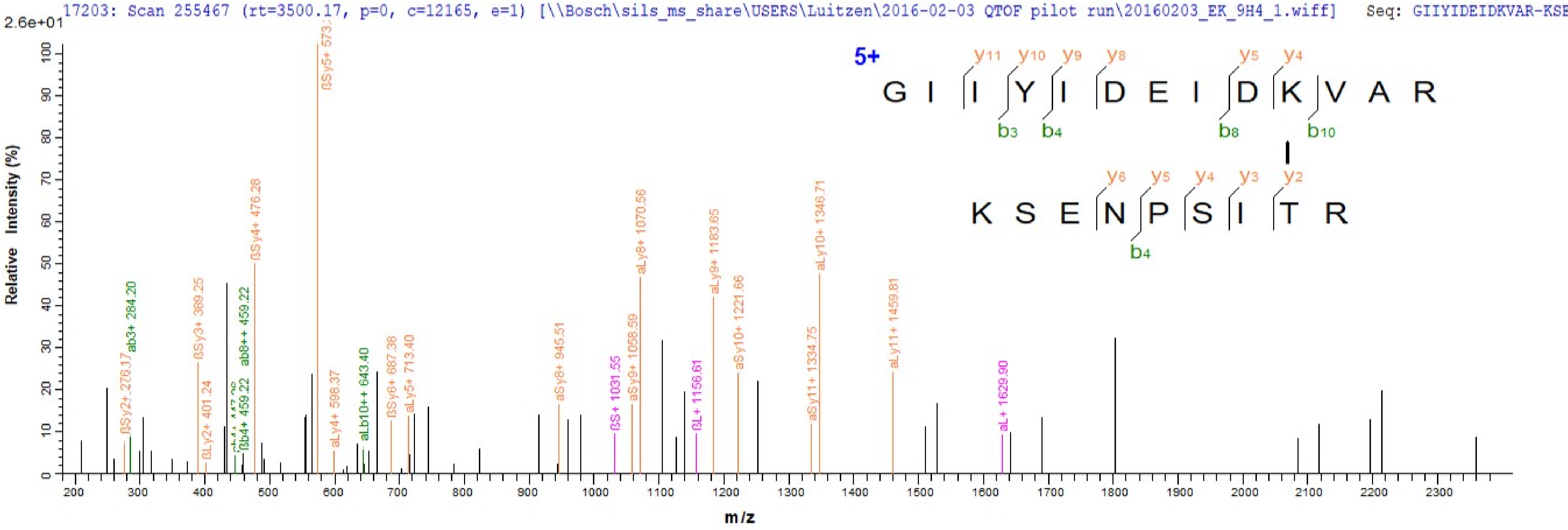
17203: Scan 255467 (rt=3500.17, p=0, c=12165, e=1) [\\Bosch\sils_ms_share\USERS\Luitzen\2016-02-03 QTOF pilot run\20160203_EK_9H4_1.wiff] Seq: GIIYIDEIDKVAR-KSE
2.6e+01
Relative Intensity (%)
m/z
5+
G I I Y I D E I D K V A R
K S E N P S I T R
ßSy5+ 573
ßSy4+ 476.28
aLy8+ 1070.56
aLy10+ 1346.71
aLy9+ 1183.65
aSy10+ 1221.66
aLy11+ 1459.81
ßSy3+ 389.25
ab3+ 284.20
ab8++ 459.22
ßSy2+ 276.17
ßLy2+ 401.24
ßb4+ 459.22
aLy4+ 598.37
aLb10++ 643.40
ßSy6+ 687.38
aLy5+ 713.40
aSy8+ 945.51
ßS+ 1031.55
aSy9+ 1058.59
ßL+ 1156.61
aSy11+ 1334.75
aL+ 1629.90

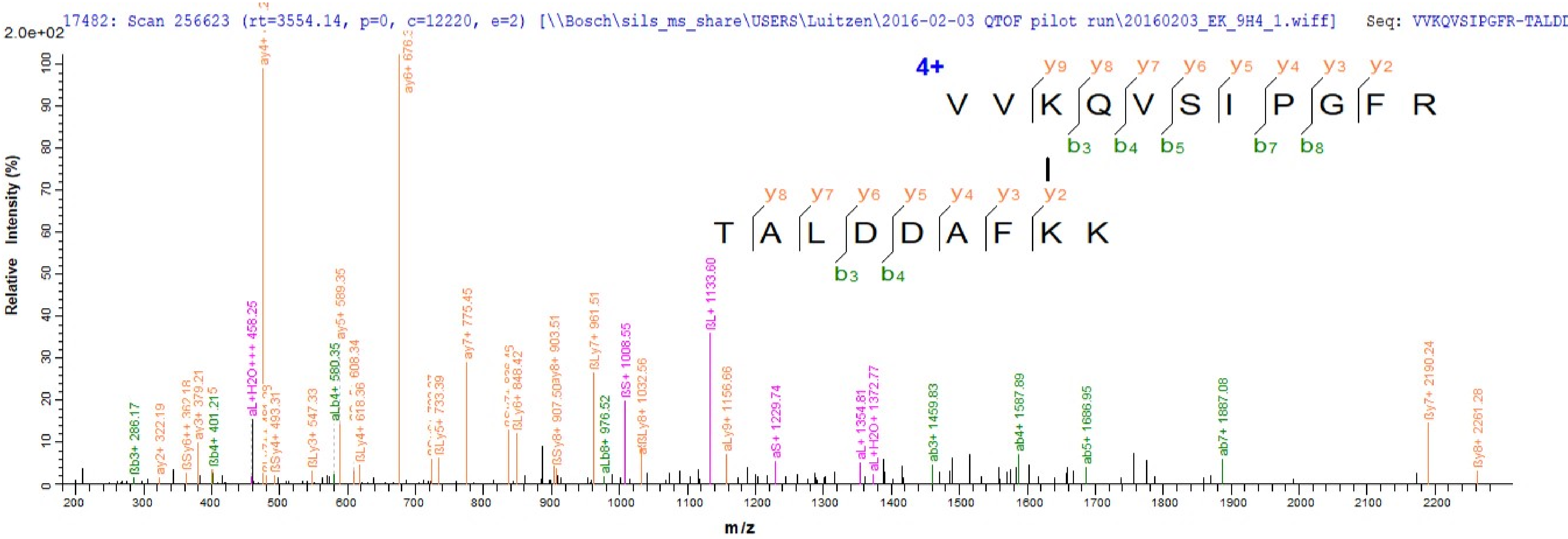
17482: Scan 256623 (rt=3554.14, p=0, c=12220, e=2) [\\Bosch\sils_ms_share\USERS\Luitzen\2016-02-03 QTOF pilot run\20160203_EK_9H4_1.wiff] Seq: VVKQVSIPGFR-TALDD
2.0e+02
Relative Intensity (%)
m/z
4+
V V K Q V S I P G F R
y9 y8 y7 y6 y5 y4 y3 y2
b3 b4 b5 b7 b8
T A L D D A F K K
y8 y7 y6 y5 y4 y3 y2
b3 b4
ßb3+ 286.17
ay2+ 322.19
ßSy6++ 362.18
ay3+ 379.21
ßb4+ 401.215
aL+H2O+++ 458.25
ßSy4+ 493.31
ßLy3+ 547.33
aLb4+ 580.35
ay5+ 589.35
608.34
ßLy4+ 618.36
ßLy5+ 733.39
ay7+ 775.45
ßLy6+ 848.42
ßSy8+ 907.50
ay8+ 903.51
ßLy7+ 961.51
aLb8+ 976.52
ßS+ 1008.55
ßLy8+ 1032.56
ßL+ 1133.60
aLy9+ 1156.66
aS+ 1229.74
aL+ 1354.81
aL+H2O+ 1372.77
ab3+ 1459.83
ab4+ 1587.89
ab5+ 1686.95
ab7+ 1887.08
ßy7+ 2190.24
ßy8+ 2261.28

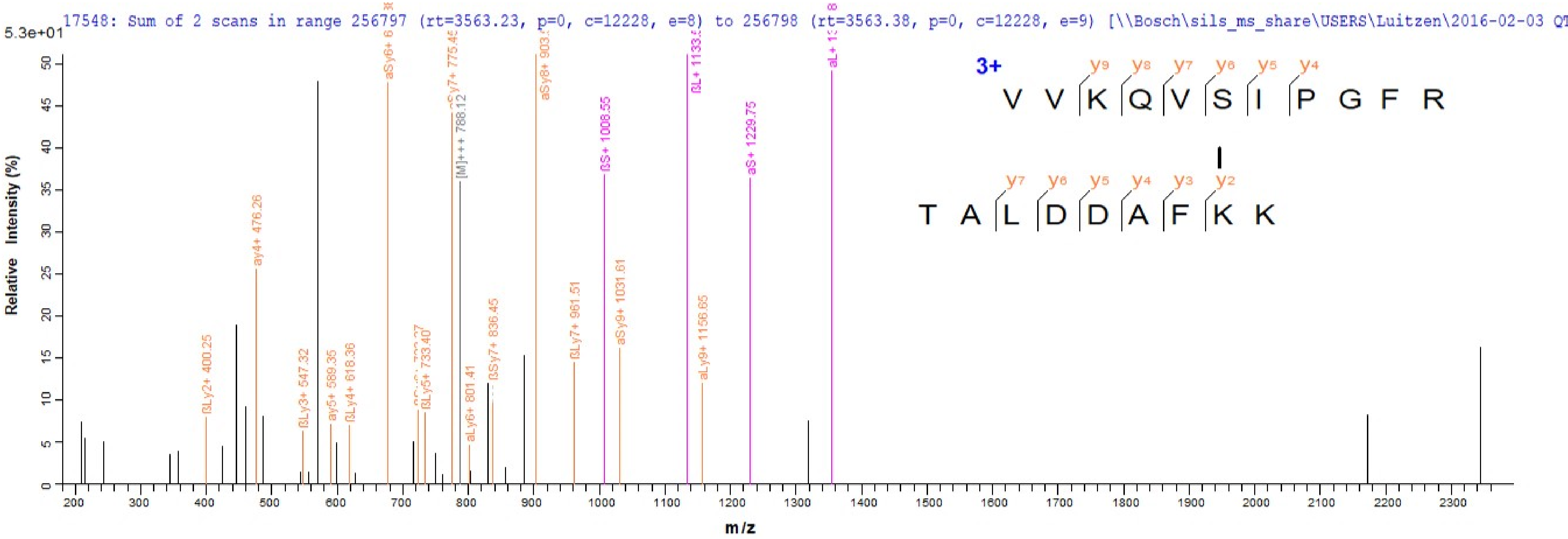
17548: Sum of 2 scans in range 256797 (rt=3563.23, p=0, c=12228, e=8) to 256798 (rt=3563.38, p=0, c=12228, e=9) [\\Bosch\sils_ms_share\USERS\Luitzen\2016-02-03 QT
5.3e+01
Relative Intensity (%)
m/z
3+
V V K Q V S I P G F R
T A L D D A F K K
y9 y8 y7 y6 y5 y4
y7 y6 y5 y4 y3 y2
ßLy2+ 400.25
ay4+ 476.26
ßLy3+ 547.32
ay5+ 589.35
ßLy4+ 618.36
ßLy5+ 733.40
[M]+++ 788.12
aLy6+ 801.41
ßSy7+ 836.45
aSy8+ 903.5
ßLy7+ 961.51
ßS+ 1008.55
aSy9+ 1031.61
aLy9+ 1156.65
aS+ 1229.75

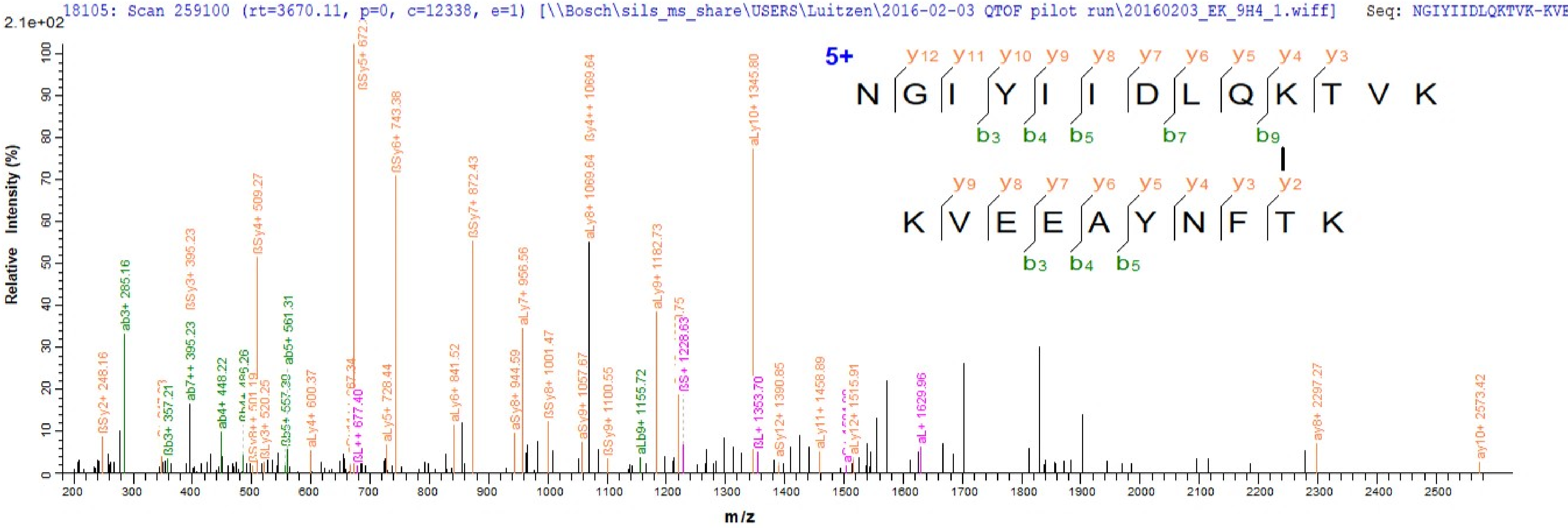
18105: Scan 259100 (rt=3670.11, p=0, c=12338, e=1) [\\Bosch\sils_ms_share\USERS\Luitzen\2016-02-03 QTOF pilot run\20160203_EK_9H4_1.wiff]
Seq: NGIYIIDLQKTVK-KVE
2.1e+02
Relative Intensity (%)
m/z
5+
N G I Y I I D L Q K T V K
K V E E A Y N F T K
ab3+ 285.16
ßSy5+ 672.
ßSy6+ 743.38
ßSy4++ 1069.64
aLy10+ 1345.80
aLy8+ 1069.64
ßSy7+ 872.43
ßSy4+ 509.27
ßSy3+ 395.23
aLy7+ 956.56
aLy9+ 1182.73
ab7++ 395.23
ab5+ 561.31
ab4+ 448.22
ßSy2+ 248.16
ay8+ 2297.27
ay10+ 2573.42
aL+ 1629.96
ßL+ 1353.70
aLy11+ 1458.89
aSy12+ 1390.85
aLy12+ 1515.91
ßS+ 1228.63
aLb9+ 1155.72
ßSy9+ 1100.55
aSy9+ 1057.67
ßSy8+ 1001.47
aSy8+ 944.59
aLy6+ 841.52
aLy5+ 728.44
aLy4+ 600.37
ßL++ 677.40
ßLy3+ 520.25
ßb3+ 357.21

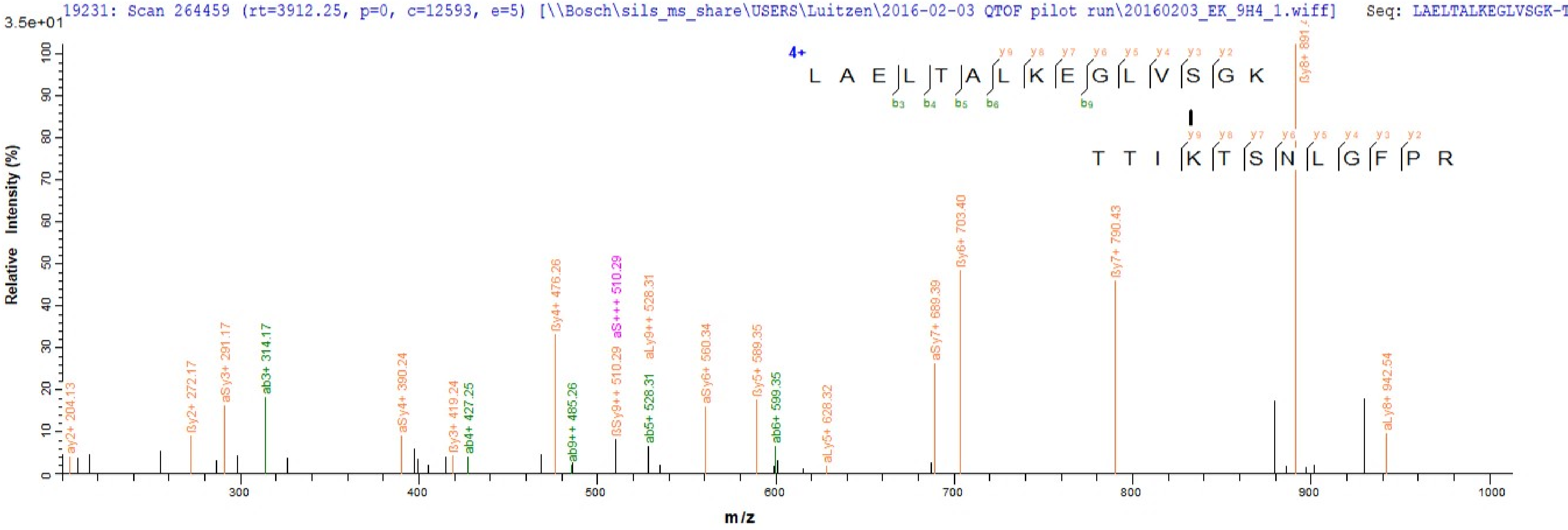

19231: Scan 264459 (rt=3912.25, p=0, c=12593, e=5) [\\Bosch\sils_ms_share\USERS\Luitzen\2016-02-03 QTOF pilot run\20160203_EK_9H4_1.wiff] Seq: LAELTALKEGLVSGK-T
3.5e+01
Relative Intensity (%)
m/z
4+
L A E L T A L K E G L V S G K
T T I K T S N L G F P R
ay2+ 204.13
ßy2+ 272.17
aSy3+ 291.17
ab3+ 314.17
aSy4+ 390.24
ßy3+ 419.24
ab4+ 427.25
ßy4+ 476.26
ab9++ 485.26
ßSy9++ 510.29
aS+++ 510.29
ab5+ 528.31
aLy9++ 528.31
aSy6+ 560.34
ßy5+ 589.35
ab6+ 599.35
aLy5+ 628.32
aSy7+ 689.39
ßy6+ 703.40
ßy7+ 790.43
ßy8+ 891.4
aLy8+ 942.54

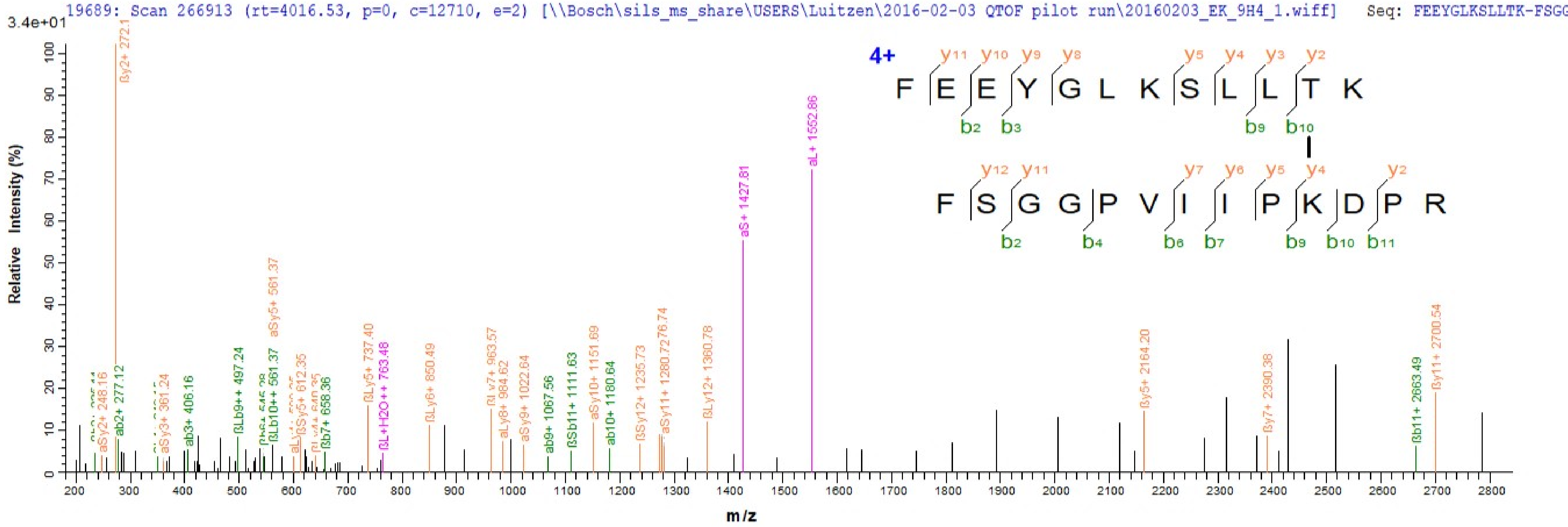
19689: Scan 266913 (rt=4016.53, p=0, c=12710, e=2) [\\Bosch\sils_ms_share\USERS\Luitzen\2016-02-03 QTOF pilot run\20160203_EK_9H4_1.wiff] Seq: FEEYGLKSLLTK-FSGG
3.4e+01
Relative Intensity (%)
m/z
4+
F E E Y G L K S L L T K
F S G G P V I I P K D P R
ßy2+ 272.1
aL+ 1552.86
aS+ 1427.81
aSy5+ 561.37
ßy11+ 2700.54
ßy5+ 2164.20
ßy7+ 2390.38
ßb11+ 2663.49
ßLy12+ 1360.78
aSy11+ 1280.72
aSy10+ 1151.69
ßSy12+ 1235.73
ab10+ 1180.64
ßSb11+ 1111.63
ab9+ 1067.56
aSy9+ 1022.64
aLy8+ 984.62
ßLy6+ 850.49
ßLy5+ 737.40
ßL+H2O++ 763.48
ßb7+ 658.36
ßSy5+ 612.35
ßLb10++ 561.37
ßLb9++ 497.24
ab3+ 406.16
aSy3+ 361.24
ab2+ 277.12
aSy2+ 248.16

20430: Scan 270188 (rt=4168.96, p=0, c=12866, e=1) [\\Bosch\sils_ms_share\USERS\Luitzen\2016-02-03 QTOF pilot run\20160203_EK_9H4_1.wiff] Seq: ISLSIKDTLPGPWNQIG

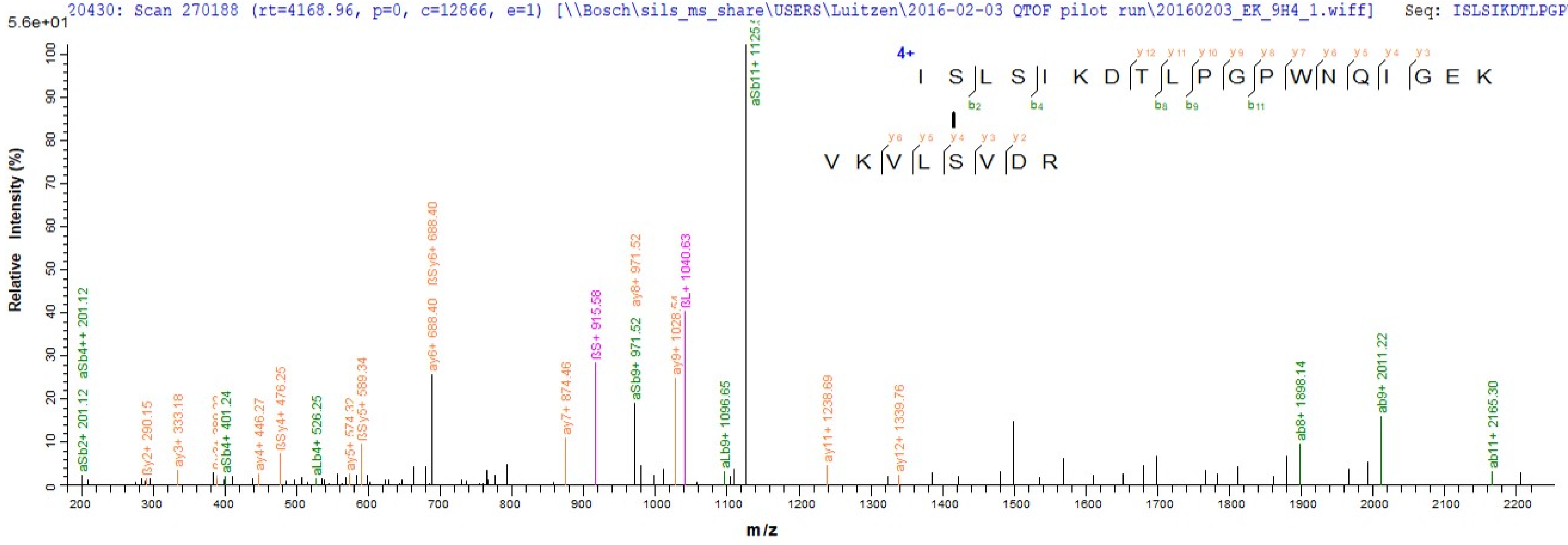

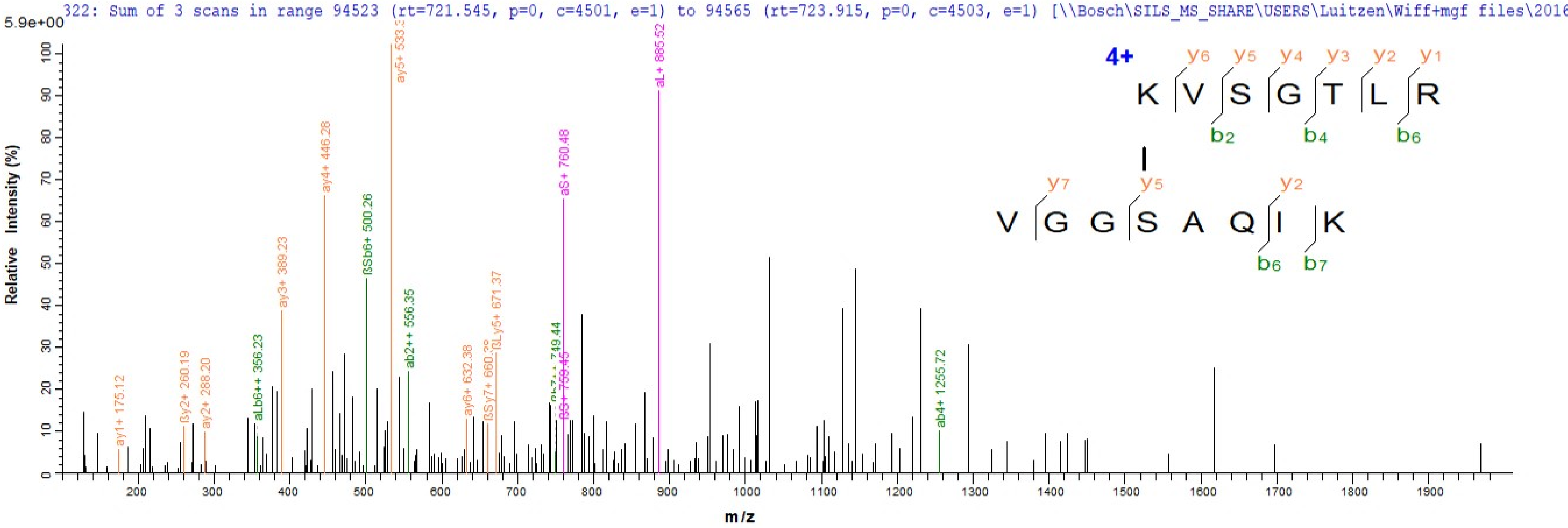
322: Sum of 3 scans in range 94523 (rt=721.545, p=0, c=4501, e=1) to 94565 (rt=723.915, p=0, c=4503, e=1) [\\Bosch\SILS_MS_SHARE\USERS\Luitzen\Wiff+mgf files\2016
5.9e+00
Relative Intensity (%)
m/z
ay1+ 175.12
βy2+ 260.19
ay2+ 288.20
aLb6++ 356.23
ay3+ 389.23
ay4+ 446.28
βSb6+ 500.26
ay5+ 533.3
ab2++ 556.35
ay6+ 632.38
βSy7+ 660.3
βLy5+ 671.37
749.44
aS+ 760.48
aL+ 885.52
ab4+ 1255.72
4+
K V S G T L R
y6 y5 y4 y3 y2 y1
b2 b4 b6
V G G S A Q I K
y7 y5 y2
b6 b7

322: Sum of 3 scans in range 94523 (rt=721.545, p=0, c=4501, e=1) to 94565 (rt=723.915, p=0, c=4503, e=1) [\\Bosch\SILS_MS_SHARE\USERS\Luitzen\Wiff+mgf files\2016

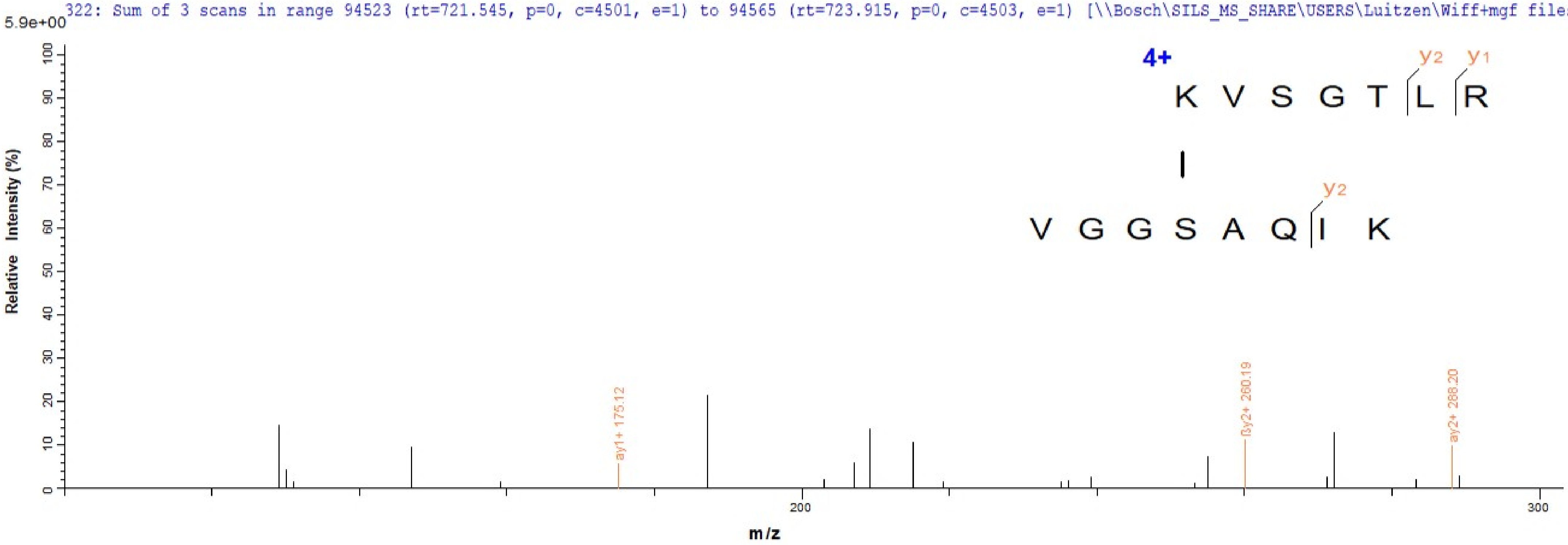

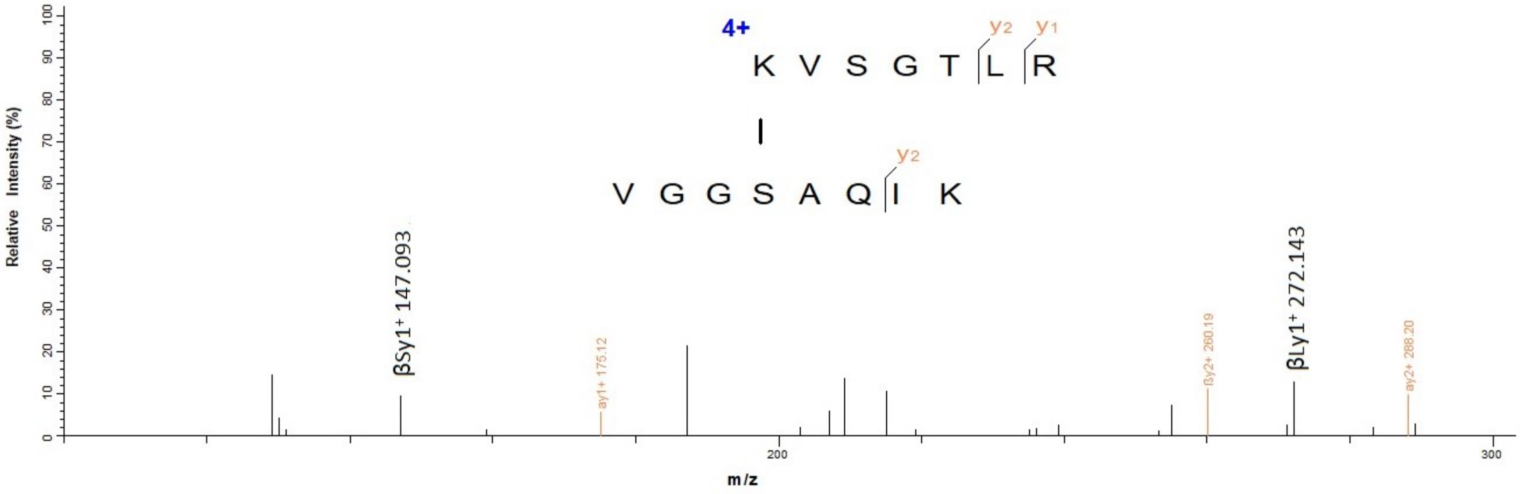
4+
KVSGTLR
VGGSAQIK
y2
y1
y2
βSy1+ 147.093
ay1+ 175.12
βy2+ 260.19
βLy1+ 272.143
ay2+ 288.20
Relative Intensity (%)
m/z
200
300

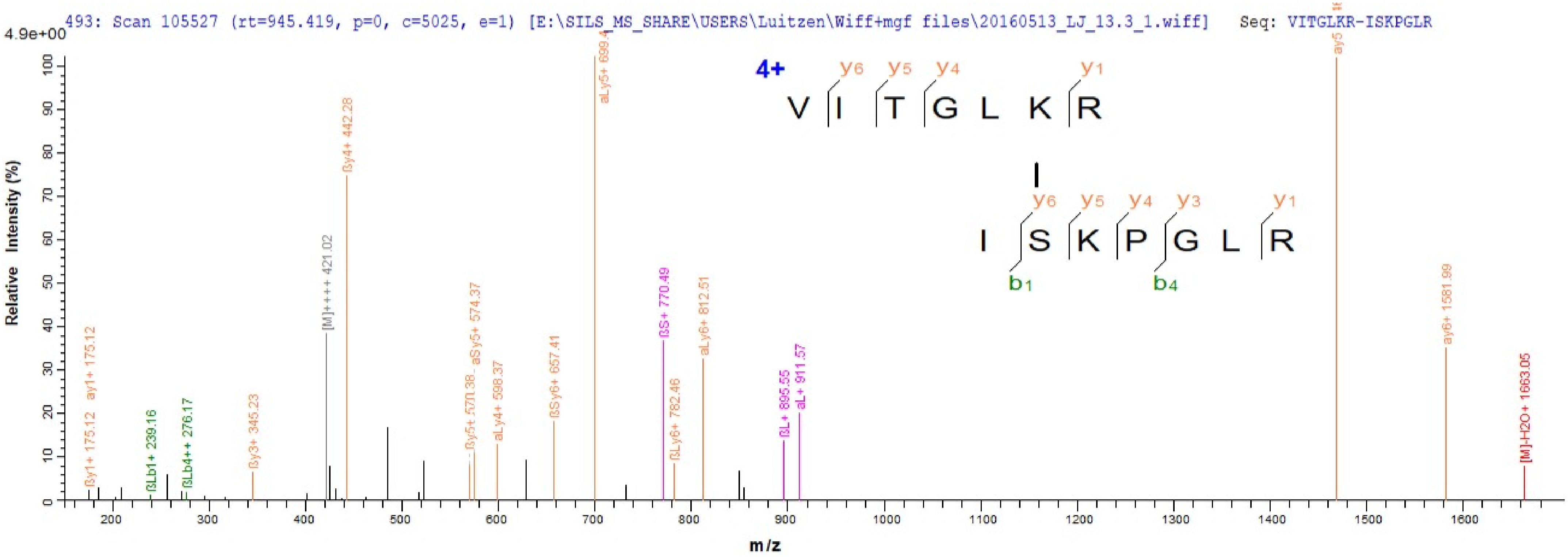

493: Scan 105527 (rt=945.419, p=0, c=5025, e=1) [E:\SILS_MS_SHARE\USERS\Luitzen\Wiff+mgf files\20160513_LJ_13.3_1.wiff] Seq: VITGLKR-ISKPGLR
4.9e+00
Relative Intensity (%)
m/z
4+
V I T G L K R
I S K P G L R
ßy1+ 175.12
ay1+ 175.12
ßLb1+ 239.16
ßLb4++ 276.17
ßy3+ 345.23
[M]++++ 421.02
ßy4+ 442.28
aSy5+ 574.37
aLy4+ 598.37
ßSy6+ 657.41
ßS+ 770.49
ßLy6+ 782.46
aLy6+ 812.51
ßL+ 895.55
aL+ 911.57
ay6+ 1581.99
[M]-H2O+ 1663.05

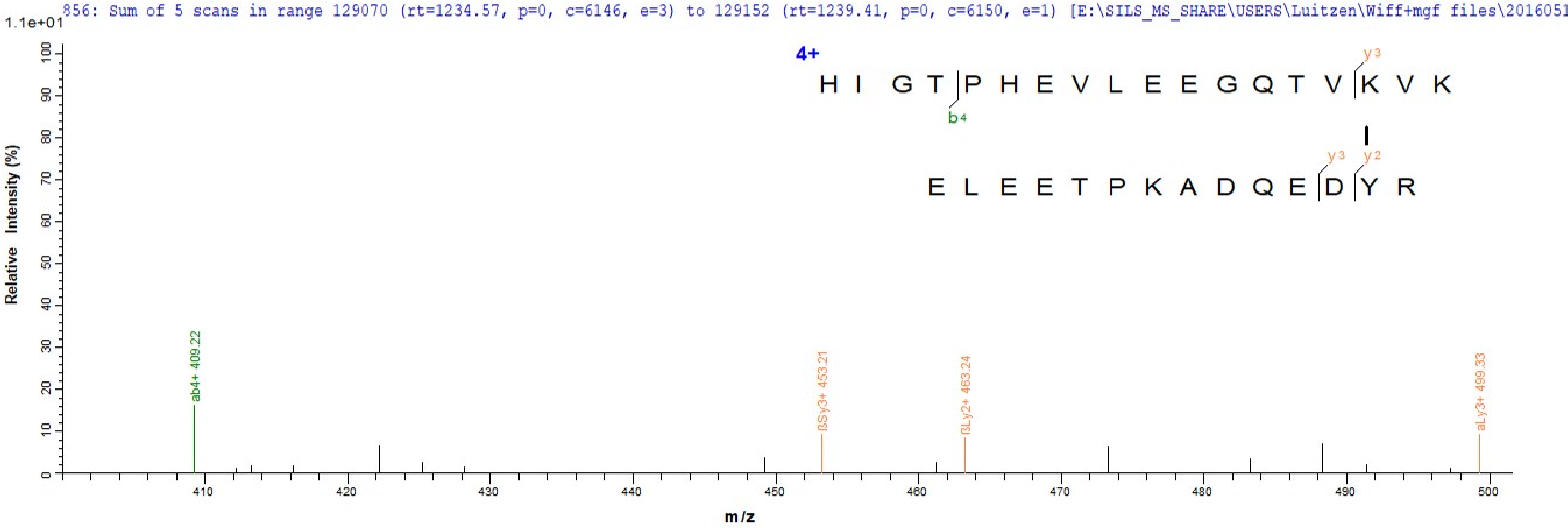
856: Sum of 5 scans in range 129070 (rt=1234.57, p=0, c=6146, e=3) to 129152 (rt=1239.41, p=0, c=6150, e=1) [E:\SILS_MS_SHARE\USERS\Luitzen\Wiff+mgf files\2016051
1.1e+01
Relative Intensity (%)
m/z
4+
H I G T P H E V L E E G Q T V K V K
b4
y3
E L E E T P K A D Q E D Y R
y3
y2
ab4+ 409.22
βSy3+ 453.21
βLy2+ 463.24
aLy3+ 499.33

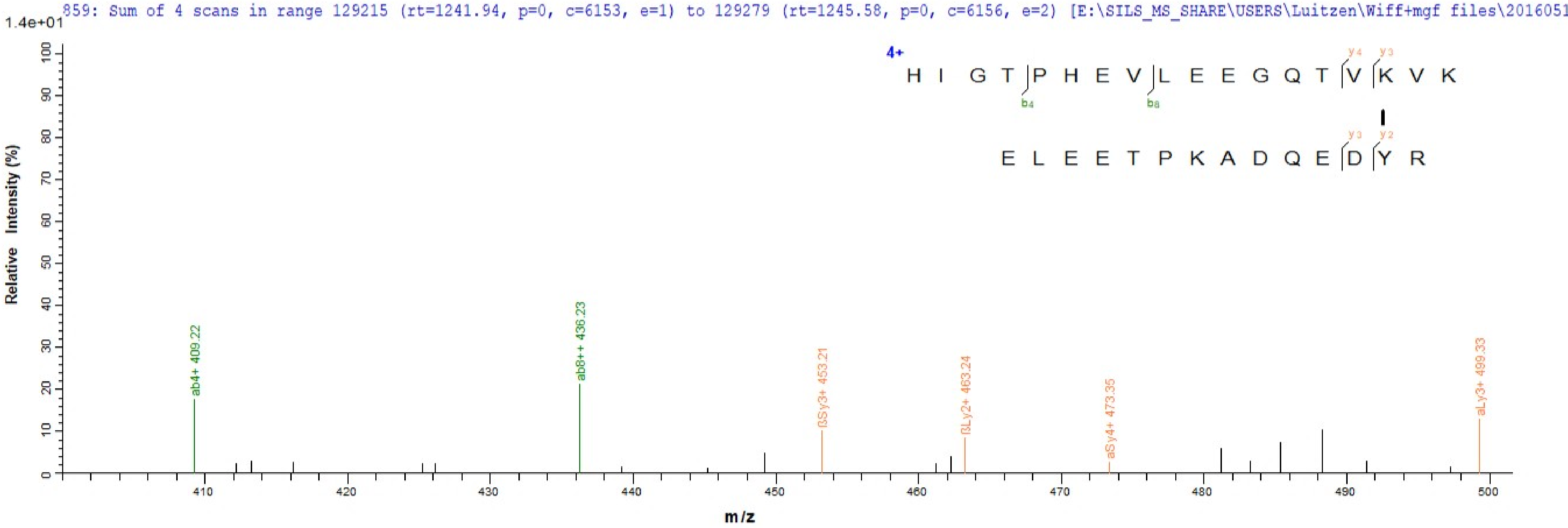

859: Sum of 4 scans in range 129215 (rt=1241.94, p=0, c=6153, e=1) to 129279 (rt=1245.58, p=0, c=6156, e=2) [E:\SILS_MS_SHARE\USERS\Luitzen\Wiff+mgf files\2016051
1.4e+01
Relative Intensity (%)
m/z
4+
H I G T P H E V L E E G Q T V K V K
E L E E T P K A D Q E D Y R
ab4+ 409.22
ab8++ 436.23
βSy3+ 453.21
βLy2+ 463.24
aSy4+ 473.35
aLy3+ 499.33

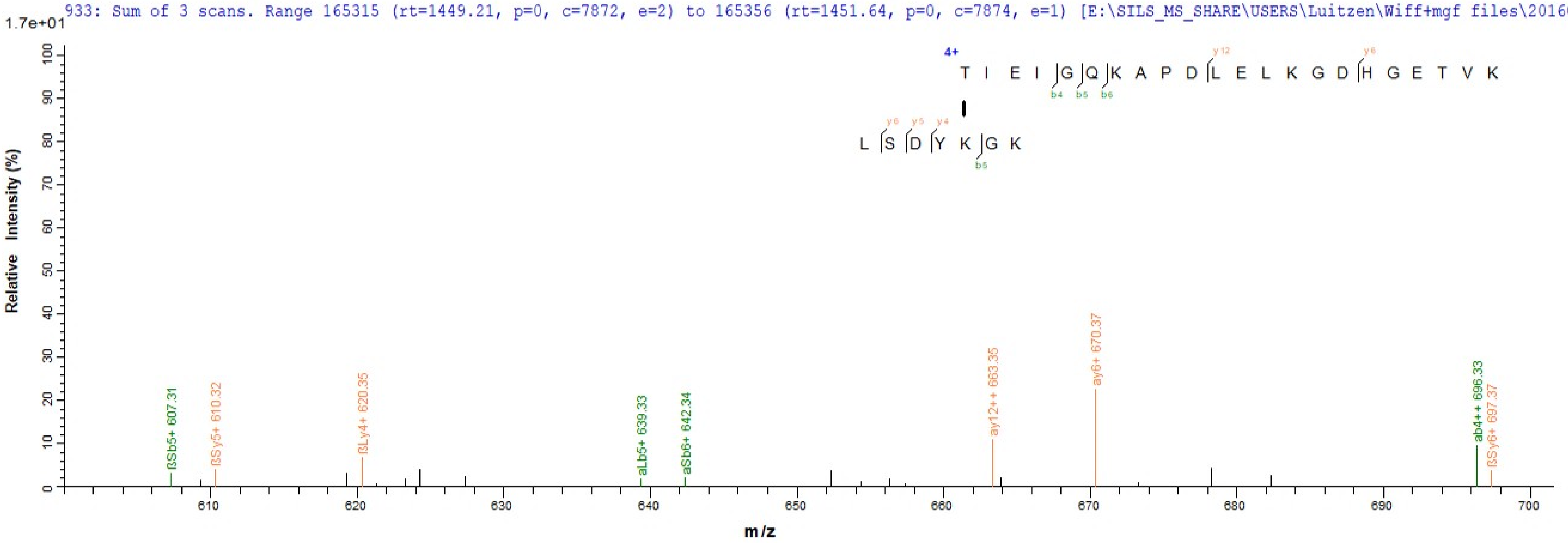

933: Sum of 3 scans. Range 165315 (rt=1449.21, p=0, c=7872, e=2) to 165356 (rt=1451.64, p=0, c=7874, e=1) [E:\SILS_MS_SHARE\USERS\Luitzen\Wiff+mgf files\20160513_
1.7e+01
4+
T I E I G Q K A P D L E L K G D H G E T V K
L S D Y K G K
ßSb5+ 607.31
ßSy5+ 610.32
ßLy4+ 620.35
aLb5+ 639.33
aSb6+ 642.34
ay12++ 663.35
ay6+ 670.37
ab4++ 696.33
ßSy6+ 697.37
Relative Intensity (%)
m/z

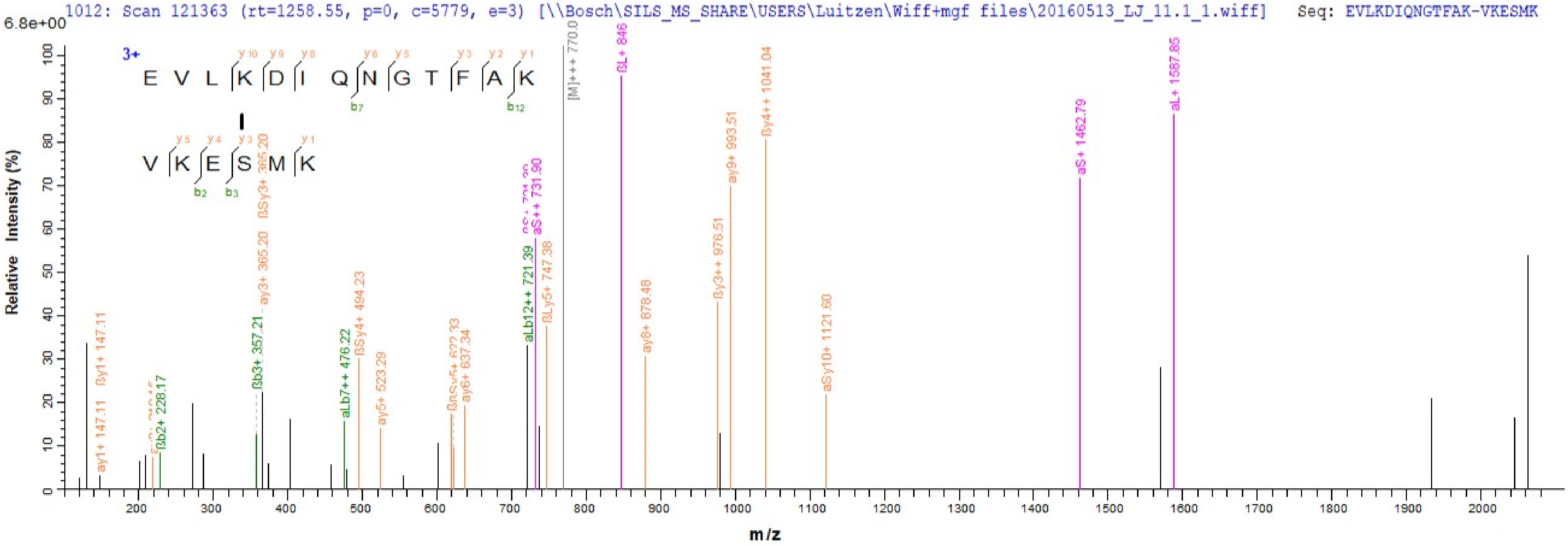
1012: Scan 121363 (rt=1258.55, p=0, c=5779, e=3) [\\Bosch\SILS_MS_SHARE\USERS\Luitzen\Wiff+mgf files\20160513_LJ_11.1_1.wiff] Seq: EVLKDIQNGTFAK-VKESMK
6.8e+00
3+
E V L K D I Q N G T F A K
V K E S M K
Relative Intensity (%)
m/z
ßy1+ 147.11
ay1+ 147.11
ßb2+ 228.17
ßb3+ 357.21
ßSy3+ 365.20
ay3+ 365.20
aLb7++ 476.22
ßSy4+ 494.23
ay5+ 523.29
ay6+ 637.34
aLb12++ 721.39
aS++ 731.90
ßLy5+ 747.38
[M]+++ 770.0
ßL+ 846
ay8+ 878.48
ßy3++ 976.51
ay9+ 993.51
ßy4++ 1041.04
aSy10+ 1121.60
aS+ 1462.79
aL+ 1587.85

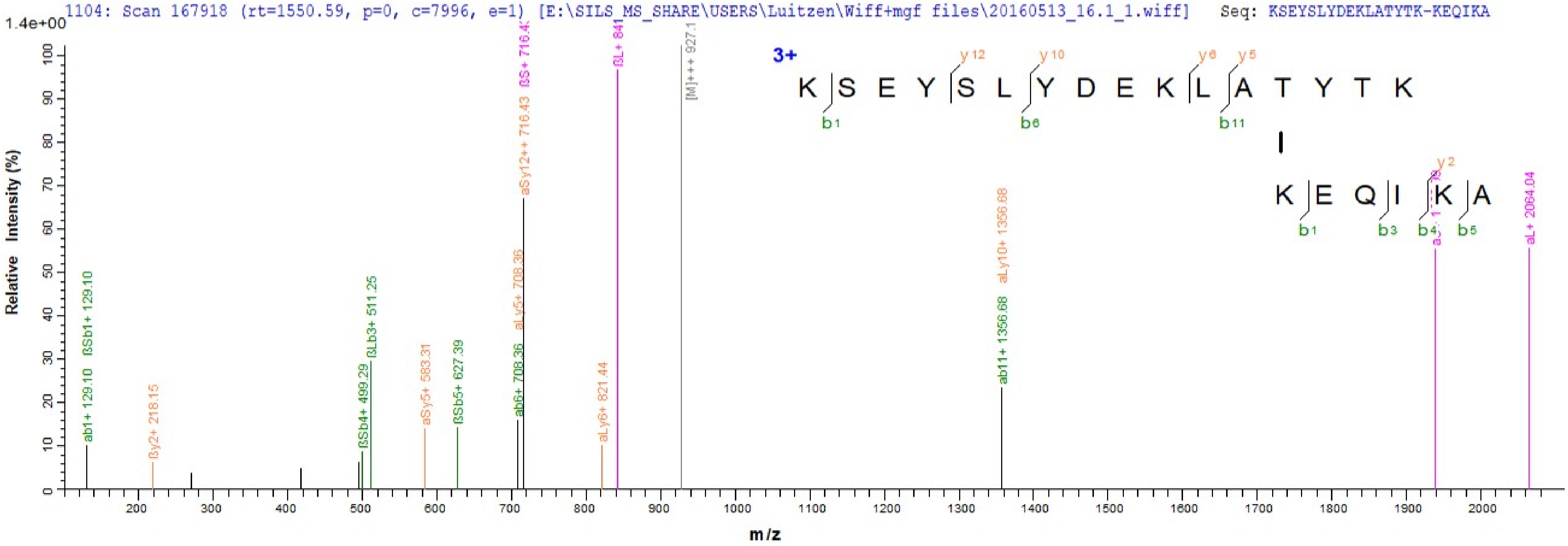
1104: Scan 167918 (rt=1550.59, p=0, c=7996, e=1) [E:\SILS_MS_SHARE\USERS\Luitzen\Wiff+mgf files\20160513_16.1_1.wiff] Seq: KSEYSLYDEKLATYTK-KEQIKA
1.4e+00
Relative Intensity (%)
m/z
3+
K S E Y S L Y D E K L A T Y T K
y12
y10
y6
y5
b1
b6
b11
K E Q I K A
y2
b1
b3
b4
b5
ab1+ 129.10
ßSb1+ 129.10
ßy2+ 218.15
ßSb4+ 499.29
ßLb3+ 511.25
aSy5+ 583.31
ßSb5+ 627.39
ab6+ 708.36
aLy5+ 708.36
aSy12++ 716.43
ßS+ 716.43
aLy6+ 821.44
ßL+ 841
[M]+++ 927.1
ab11+ 1356.68
aLy10+ 1356.68
aL+ 2064.04

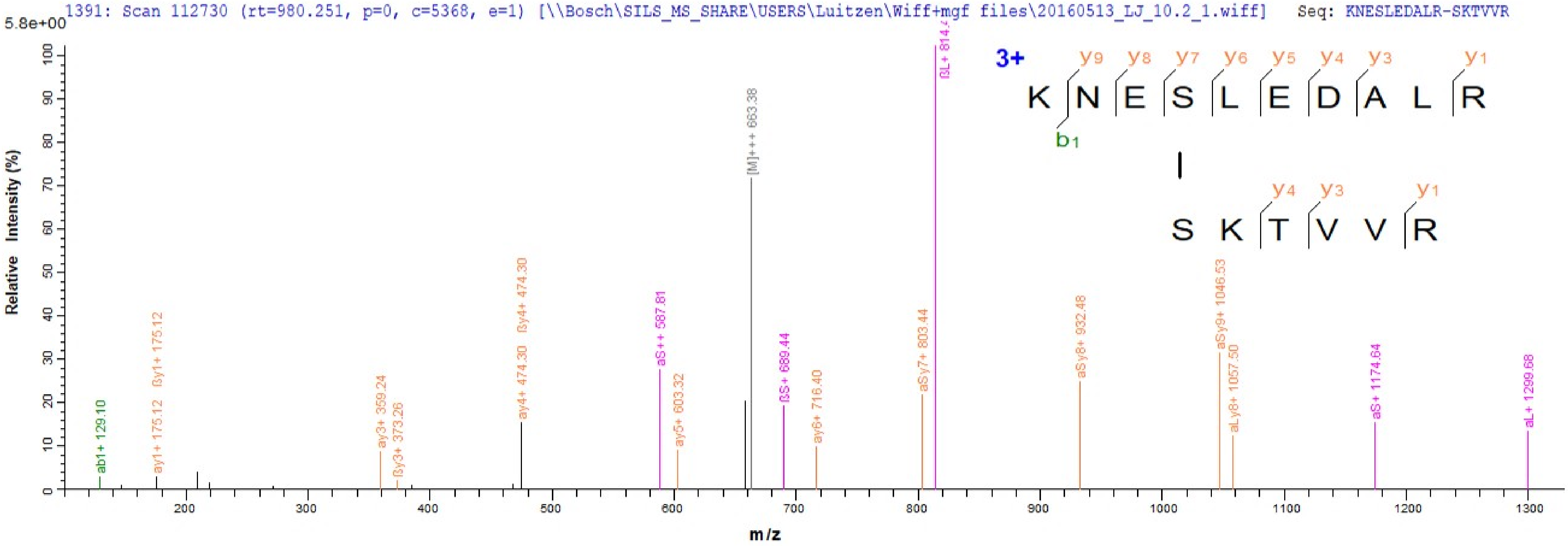
1391: Scan 112730 (rt=980.251, p=0, c=5368, e=1) [\\Bosch\SILS_MS_SHARE\USERS\Luitzen\Wiff+mgf files\20160513_LJ_10.2_1.wiff] Seq: KNESLEDALR-SKTVVR
5.8e+00
Relative Intensity (%)
m/z
3+
K N E S L E D A L R
S K T V V R
ab1+ 129.10
ay1+ 175.12
ßy1+ 175.12
ay3+ 359.24
ßy3+ 373.26
ay4+ 474.30
ßy4+ 474.30
aS++ 587.81
ay5+ 603.32
[M]+++ 663.38
ßS+ 689.44
ay6+ 716.40
aSy7+ 803.44
aSy8+ 932.48
aSy9+ 1046.53
aLy8+ 1057.50
aS+ 1174.64
aL+ 1299.68

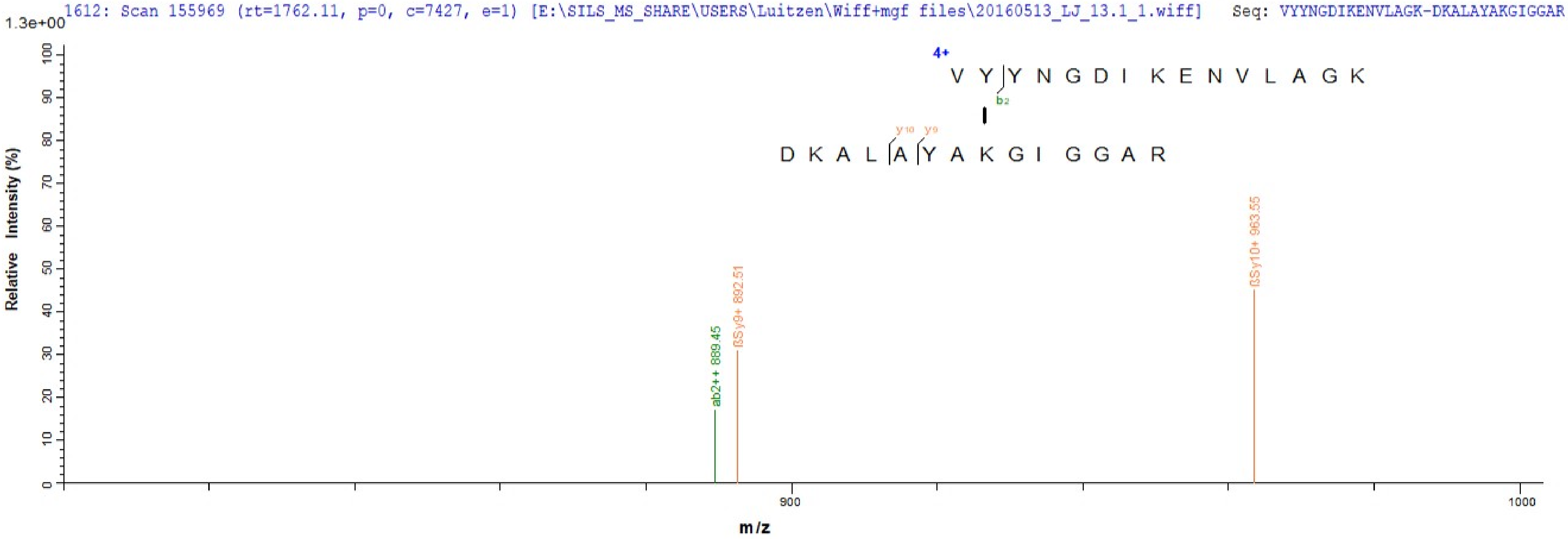
1612: Scan 155969 (rt=1762.11, p=0, c=7427, e=1) [E:\SILS_MS_SHARE\USERS\Luitzen\Wiff+mgf files\20160513_LJ_13.1_1.wiff] Seq: VYYNGDIKENVLAGK-DKALAYAKGIGGAR
1.3e+00
4+
V Y Y N G D I K E N V L A G K
b2
y10 y9
D K A L A Y A K G I G G A R
ab2++ 889.45
ßSy9+ 892.51
ßSy10+ 963.55
Relative Intensity (%)
m/z
900
1000

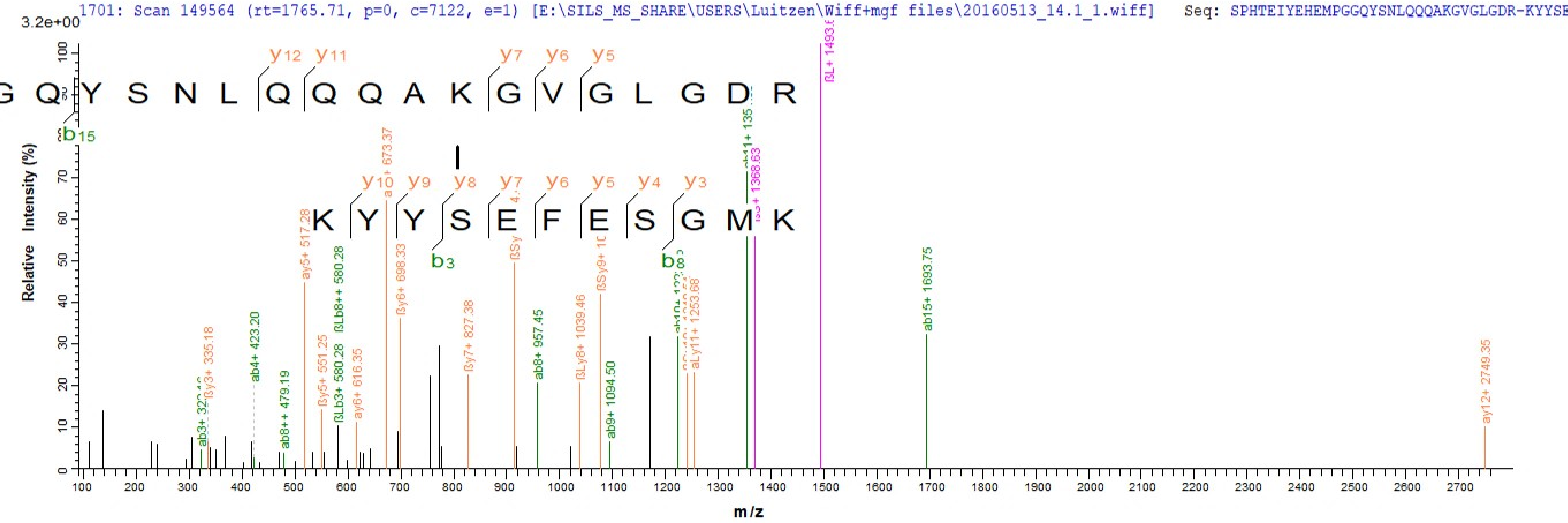

1701: Scan 149564 (rt=1765.71, p=0, c=7122, e=1) [E:\SILS_MS_SHARE\USERS\Luitzen\Wiff+mgf files\20160513_14.1_1.wiff] Seq: SPHTEIYEHEMPGGQYSNLQQQAKGVGLGDR-KYYSE
3.2e+00
G Q Y S N L Q Q Q A K G V G L G D R
K Y Y S E F E S G M K
Relative Intensity (%)
m/z
ab15+ 1693.75
ay12+ 2749.35
ay5+ 517.28
ab4+ 423.20
ab8+ 957.45
ab9+ 1094.50
ßy6+ 698.33
ßy7+ 827.38
ßLy8+ 1039.46
aLy11+ 1253.68
ab8++ 479.19
ßy3+ 335.18
ßy5+ 551.25
ay6+ 616.35
ßLb3+ 580.28
ßLb8++ 580.28

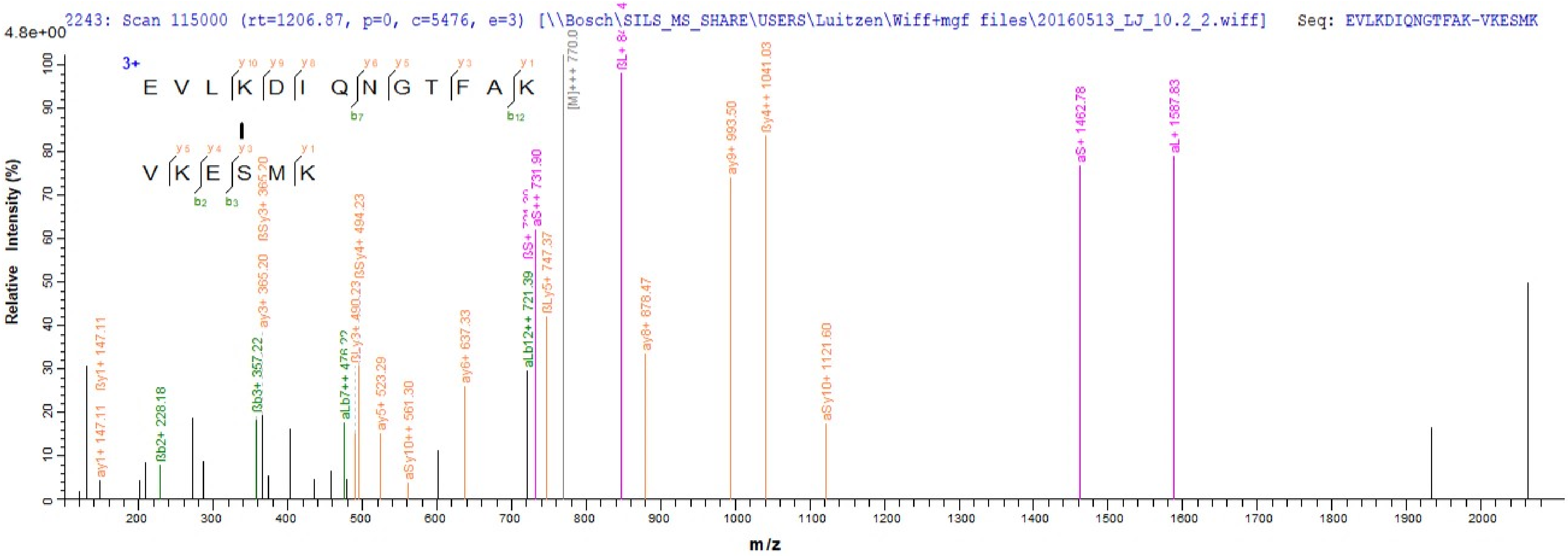
2243: Scan 115000 (rt=1206.87, p=0, c=5476, e=3) [\\Bosch\SILS_MS_SHARE\USERS\Luitzen\Wiff+mgf files\20160513_LJ_10.2_2.wiff] Seq: EVLKDIQNGTFAK-VKESMK
4.8e+00
3+
E V L K D I Q N G T F A K
V K E S M K
Relative Intensity (%)
m/z
ßy1+ 147.11
ay1+ 147.11
ßb2+ 228.18
ßb3+ 357.22
ay3+ 365.20
ßSy3+ 365.20
aLb7++ 476.22
ßLy3+ 490.23
ßSy4+ 494.23
ay5+ 523.29
aSy10++ 561.30
ay6+ 637.33
aLb12++ 721.39
aS++ 731.90
ßLy5+ 747.37
[M]+++ 770.0
ay8+ 878.47
ay9+ 993.50
ßy4++ 1041.03
aSy10+ 1121.60
aS+ 1462.78
aL+ 1587.83

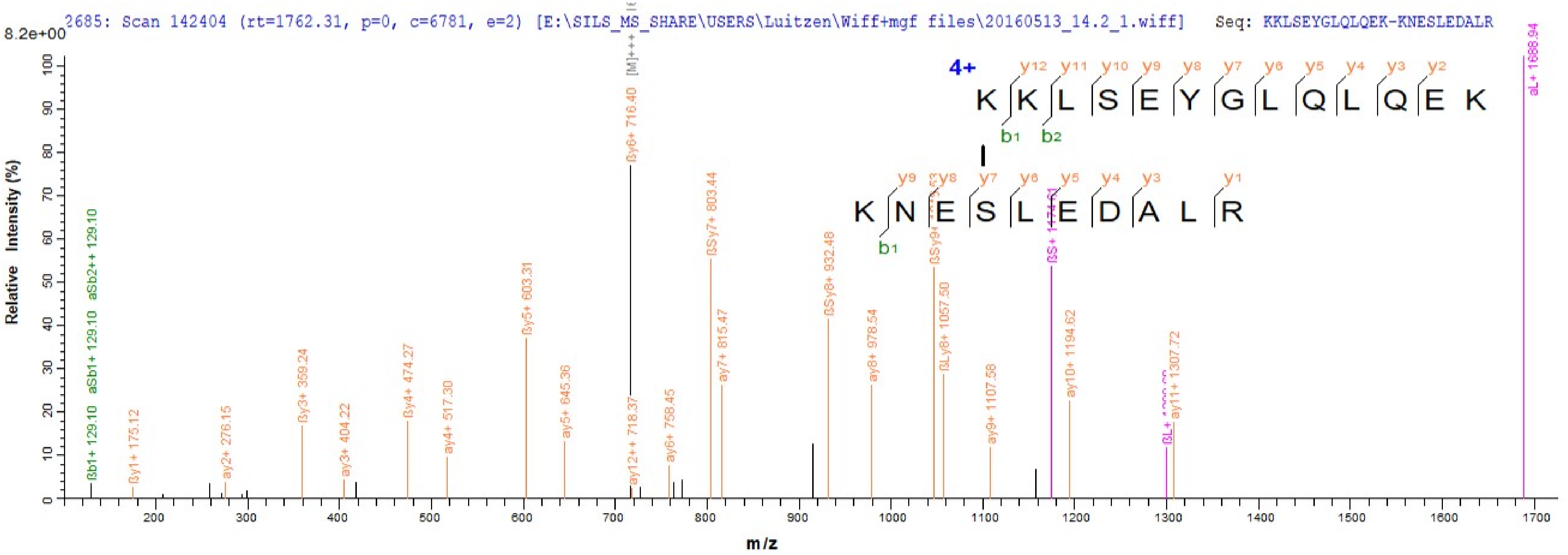
2685: Scan 142404 (rt=1762.31, p=0, c=6781, e=2) [E:\SILS_MS_SHARE\USERS\Luitzen\Wiff+mgf files\20160513_14.2_1.wiff] Seq: KKLSEYGLQLQEK-KNESLEDALR
8.2e+00
4+
KKLSEYGLQLQEK
KNESLEDALR
Relative Intensity (%)
m/z
ßb1+ 129.10
aSb1+ 129.10
aSb2++ 129.10
ßy1+ 175.12
ay2+ 276.15
ßy3+ 359.24
ay3+ 404.22
ßy4+ 474.27
ay4+ 517.30
ßy5+ 603.31
ay5+ 645.36
ßy6+ 716.40
ay12++ 718.37
ay6+ 758.45
ßSy7+ 803.44
ay7+ 815.47
ßSy8+ 932.48
ay8+ 978.54
ßLy8+ 1057.50
ay9+ 1107.58
ay10+ 1194.62
ay11+ 1307.72
aL+ 1688.94

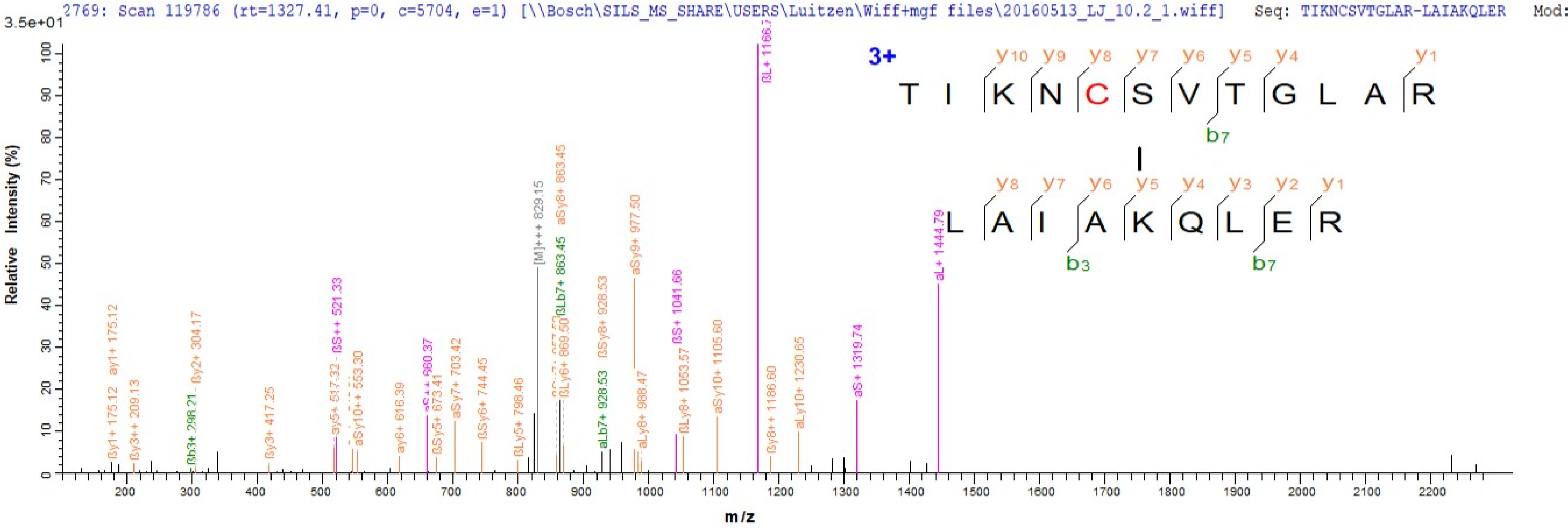
2769: Scan 119786 (rt=1327.41, p=0, c=5704, e=1) [\\Bosch\SILS_MS_SHARE\USERS\Luitzen\Wiff+mgf files\20160513_LJ_10.2_1.wiff] Seq: TIKNCSVTGLAR-LAIAKQLER Mod:
3.5e+01
Relative Intensity (%)
m/z
3+
T I K N C S V T G L A R
L A I A K Q L E R
y10 y9 y8 y7 y6 y5 y4 y1
b7
y8 y7 y6 y5 y4 y3 y2 y1
b3
b7
ßy1+ 175.12
ay1+ 175.12
ßy3++ 209.13
ßb3+ 298.21
ßy2+ 304.17
ßy3+ 417.25
ay5+ 517.32
ßS++ 521.33
aSy10++ 553.30
ay6+ 616.39
aS++ 660.37
ßSy5+ 673.41
aSy7+ 703.42
ßSy6+ 744.45
ßLy5+ 798.46
[M]+++ 829.15
aSy8+ 863.45
ßLb7+ 863.45
ßLy6+ 869.50
ßSy8+ 928.53
aLb7+ 928.53
aSy9+ 977.50
aLy8+ 988.47
ßS+ 1041.66
ßLy8+ 1053.57
aSy10+ 1105.60
ßy8++ 1186.60
aLy10+ 1230.65
aS+ 1319.74
aL+ 1444.79

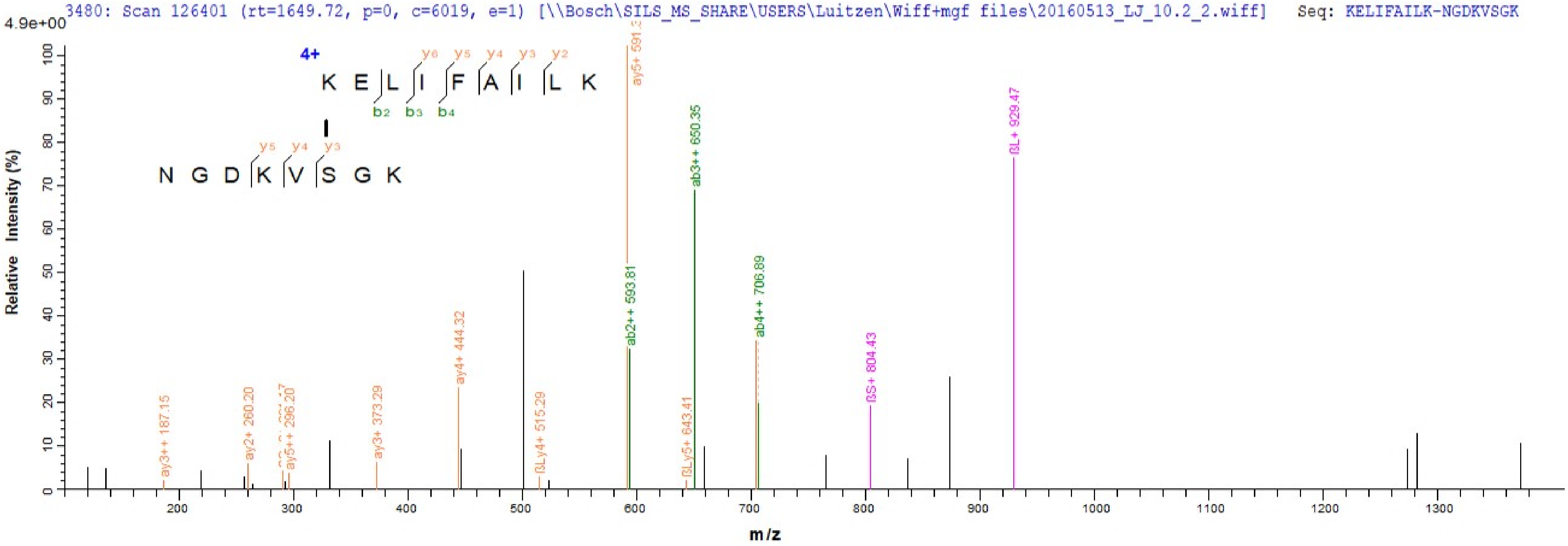
3480: Scan 126401 (rt=1649.72, p=0, c=6019, e=1) [\\Bosch\SILS_MS_SHARE\USERS\Luitzen\Wiff+mgf files\20160513_LJ_10.2_2.wiff]
Seq: KELIFAILK-NGDKVSGK
4.9e+00
Relative Intensity (%)
m/z
4+
K E L I F A I L K
NGDKVSGK
ay3++ 187.15
ay2+ 260.20
ay5++ 296.20
ay3+ 373.29
ay4+ 444.32
βLy4+ 515.29
ab2++ 593.81
βLy5+ 643.41
ab3++ 650.35
ab4++ 706.89
βS+ 804.43
βL+ 929.47

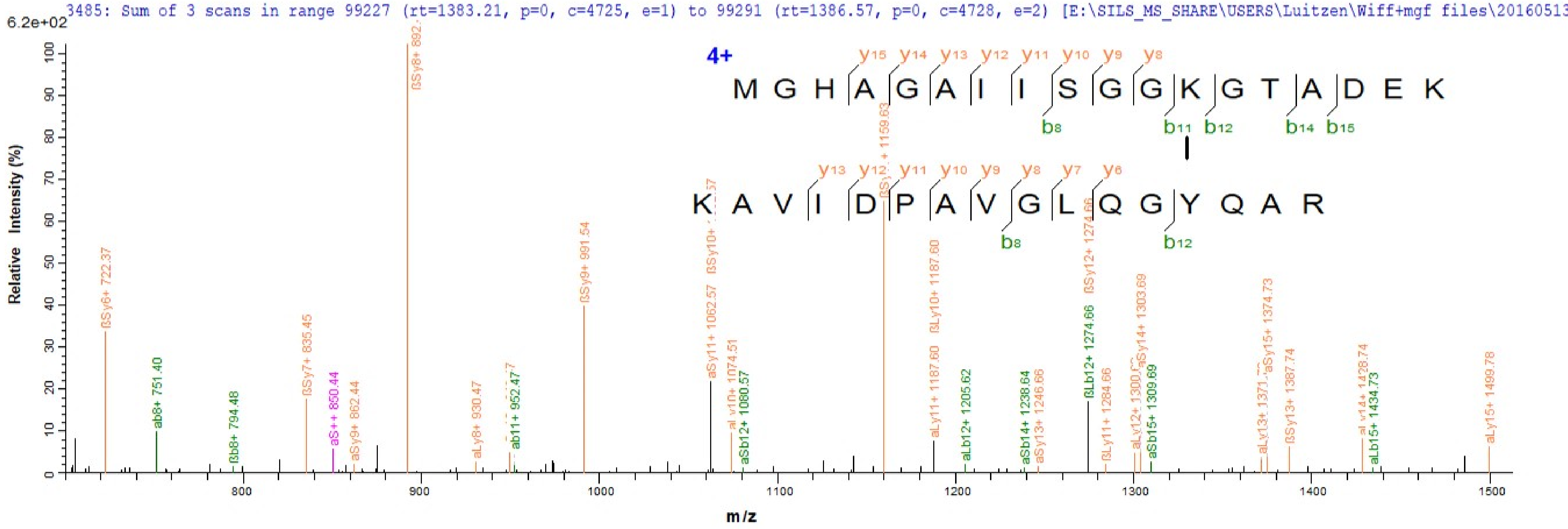
3485: Sum of 3 scans in range 99227 (rt=1383.21, p=0, c=4725, e=1) to 99291 (rt=1386.57, p=0, c=4728, e=2) [E:\SILS_MS_SHARE\USERS\Luitzen\Wiff+mgf files\20160513
6.2e+02
Relative Intensity (%)
m/z
4+
M G H A G A I I S G G K G T A D E K
y15 y14 y13 y12 y11 y10 y9 y8
b8 b11 b12 b14 b15
K A V I D P A V G L Q G Y Q A R
y13 y12 y11 y10 y9 y8 y7 y6
b8 b12
ßSy6+ 722.37
ab8+ 751.40
ßb8+ 794.48
ßSy7+ 835.45
aS++ 850.44
aSy9+ 862.44
aLy8+ 930.47
ab11+ 952.47
ßSy9+ 991.54
aSy11+ 1062.57
aSb12+ 1080.57
aLy11+ 1187.60
ßLy10+ 1187.60
aLb12+ 1205.62
aSb14+ 1238.64
aSy13+ 1246.66
ßLb12+ 1274.66
ßSy12+ 1274.66
ßLy11+ 1284.66
aSy14+ 1303.69
aSb15+ 1309.69
aSy15+ 1374.73
ßSy13+ 1387.74
aLb15+ 1434.73
aLy15+ 1499.78
800
900
1000
1100
1200
1300
1400
1500

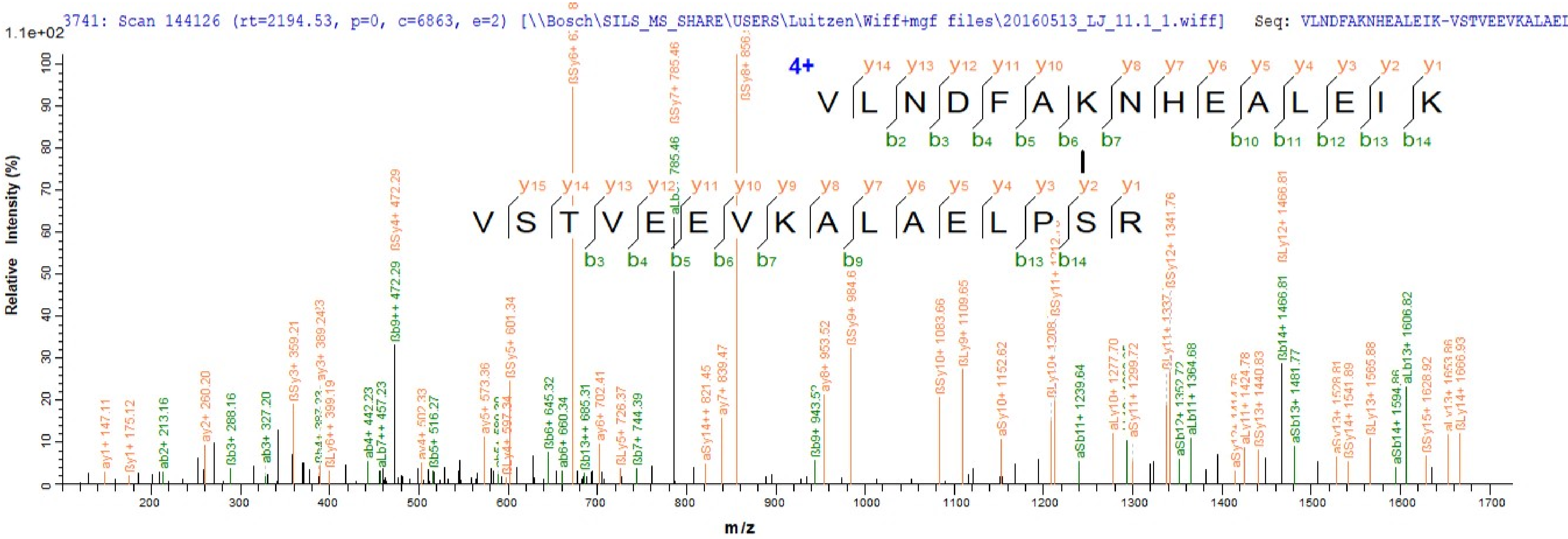
3741: Scan 144126 (rt=2194.53, p=0, c=6863, e=2) [\\Bosch\SILS_MS_SHARE\USERS\Luitzen\Wiff+mgf files\20160513_LJ_11.1_1.wiff] Seq: VLNDFAKNHEALEIK-VSTVEEVKALAEI
1.1e+02
Relative Intensity (%)
m/z
4+
VLNDFAKNHEALEIK
VSTVEEVKALAELPSR
ßSy6+
ßSy7+ 785.46
ßSy8+ 856.
aLb 785.46
ßSy4+ 472.29
ßb9++ 472.29
ßLy12+ 1466.81
ßSy12+ 1341.76
ßb14+ 1466.81
aLb13+ 1606.82
ßSy9+ 984.6
ßSy10+ 1083.66
ßLy9+ 1109.65
ßSy5+ 601.34
ßSy3+ 359.21
ay3+ 389.243
ay2+ 260.20
ay1+ 147.11
ßy1+ 175.12
ab2+ 213.16
ßb3+ 288.16
ab3+ 327.20
ßLy6++ 399.19
ab4+ 442.23
aLb7++ 457.23
av4+ 502.33
ßb5+ 516.27
ay5+ 573.36
ßLy4+ 597.34
ßb6+ 645.32
ab6+ 660.34
ßb13++ 685.31
ay6+ 702.41
ßLy5+ 726.37
ßb7+ 744.39
aSy14++ 821.45
ay7+ 839.47
ßb9+ 943.52
ay8+ 953.52
aSy10+ 1152.62
aSb11+ 1239.64
aLy10+ 1277.70
aSy11+ 1299.72
aSb12+ 1352.72
aLb11+ 1364.68
aLy11+ 1424.78
ßSy13+ 1440.83
aSb13+ 1481.77
ßSy14+ 1541.89
ßLy13+ 1565.88
aSb14+ 1594.86
ßSy15+ 1628.92
ßLy14+ 1666.93

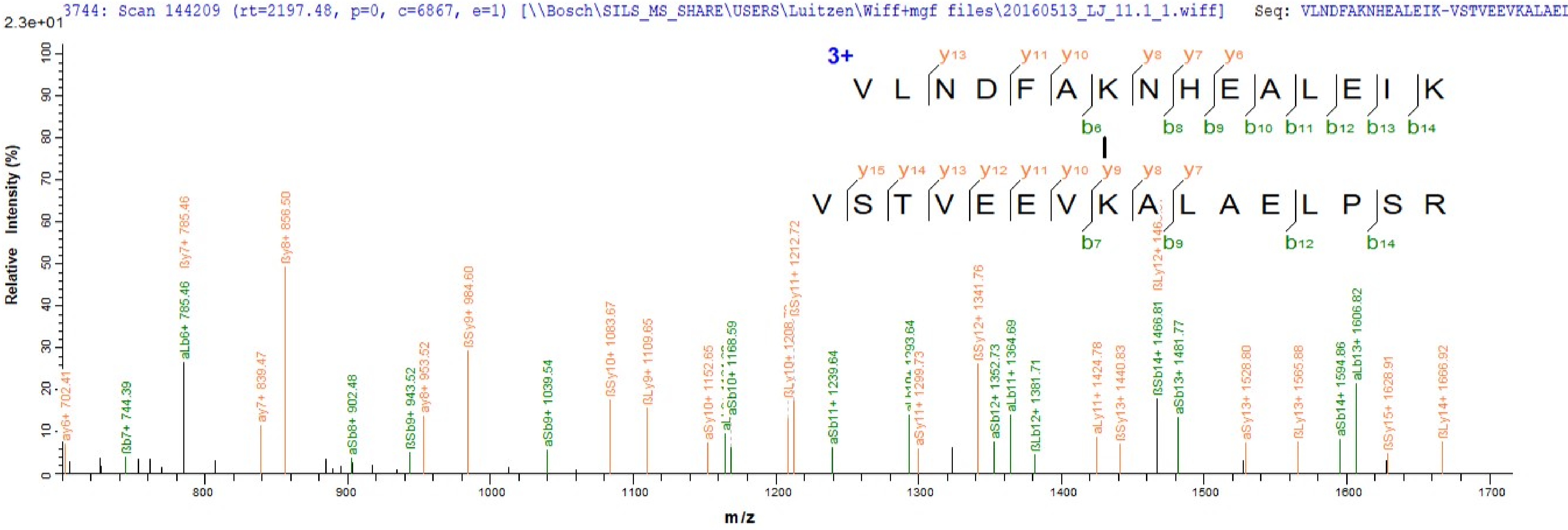
3744: Scan 144209 (rt=2197.48, p=0, c=6867, e=1) [\\Bosch\SILS_MS_SHARE\USERS\Luitzen\Wiff+mgf files\20160513_LJ_11.1_1.wiff] Seq: VLNDFAKNHEALEIK-VSTVEEVKALAEI
2.3e+01
3+
V L N D F A K N H E A L E I K
V S T V E E V K A L A E L P S R
Relative Intensity (%)
m/z
ay6+ 702.41
ßb7+ 744.39
aLb6+ 785.46
ßy7+ 785.46
ay7+ 839.47
ßy8+ 856.50
aSb8+ 902.48
ßSb9+ 943.52
ay8+ 953.52
ßSy9+ 984.60
aSb9+ 1039.54
ßSy10+ 1083.67
ßLy9+ 1109.65
aSy10+ 1152.65
aSb10+ 1168.59
ßSy11+ 1212.72
aSb11+ 1239.64
aSy11+ 1299.73
ßSy12+ 1341.76
aSb12+ 1352.73
aLb11+ 1364.69
ßLb12+ 1381.71
aLy11+ 1424.78
ßSy13+ 1440.83
ßSb14+ 1466.81
aSb13+ 1481.77
aSy13+ 1528.80
ßLy13+ 1565.88
aSb14+ 1594.86
aLb13+ 1606.82
ßSy15+ 1628.91
ßLy14+ 1666.92

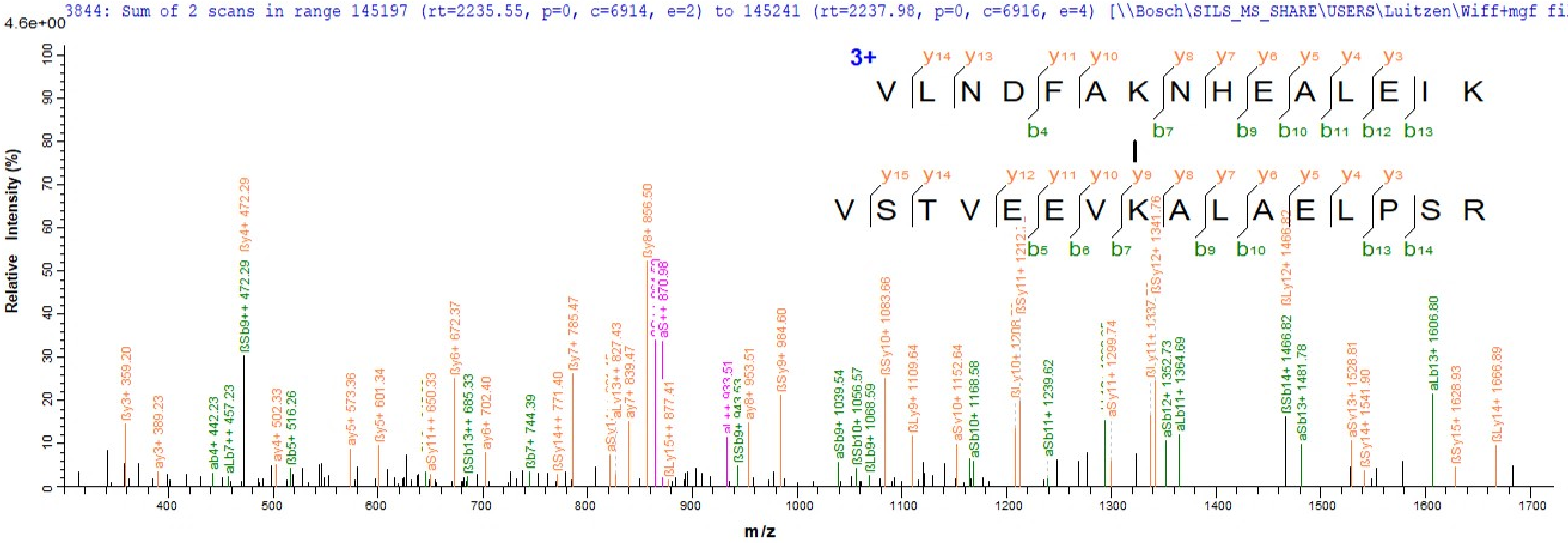
3844: Sum of 2 scans in range 145197 (rt=2235.55, p=0, c=6914, e=2) to 145241 (rt=2237.98, p=0, c=6916, e=4) [\\Bosch\SILS_MS_SHARE\USERS\Luitzen\Wiff+mgf files\2
4.6e+00
Relative Intensity (%)
m/z
3+
V L N D F A K N H E A L E I K
V S T V E E V K A L A E L P S R
ßy3+ 359.20
ay3+ 389.23
ab4+ 442.23
aLb7++ 457.23
ßSb9++ 472.29
ßy4+ 472.29
ay4+ 502.33
ßb5+ 516.26
ay5+ 573.36
ßy5+ 601.34
aSy11++ 650.33
ßy6+ 672.37
ßSb13++ 685.33
ay6+ 702.40
ßb7+ 744.39
ßSy14++ 771.40
ßy7+ 785.47
aLy13++ 827.43
ay7+ 839.47
ßy8+ 856.50
aS++ 870.98
ßLy15++ 877.41
ßSb9+ 943.53
ay8+ 953.51
ßSy9+ 984.60
aSb9+ 1039.54
ßSb10+ 1056.57
ßLb9+ 1068.59
ßSy10+ 1083.66
ßLy9+ 1109.64
aSy10+ 1152.64
aSb10+ 1168.58
aSb11+ 1239.62
aSy11+ 1299.74
ßSy12+ 1341.76
aSb12+ 1352.73
aLb11+ 1364.69
ßSb14+ 1466.82
ßLy12+ 1466.82
aSb13+ 1481.78
aSy13+ 1528.81
ßSy14+ 1541.90
aLb13+ 1606.80
ßSy15+ 1628.93
ßLy14+ 1666.89

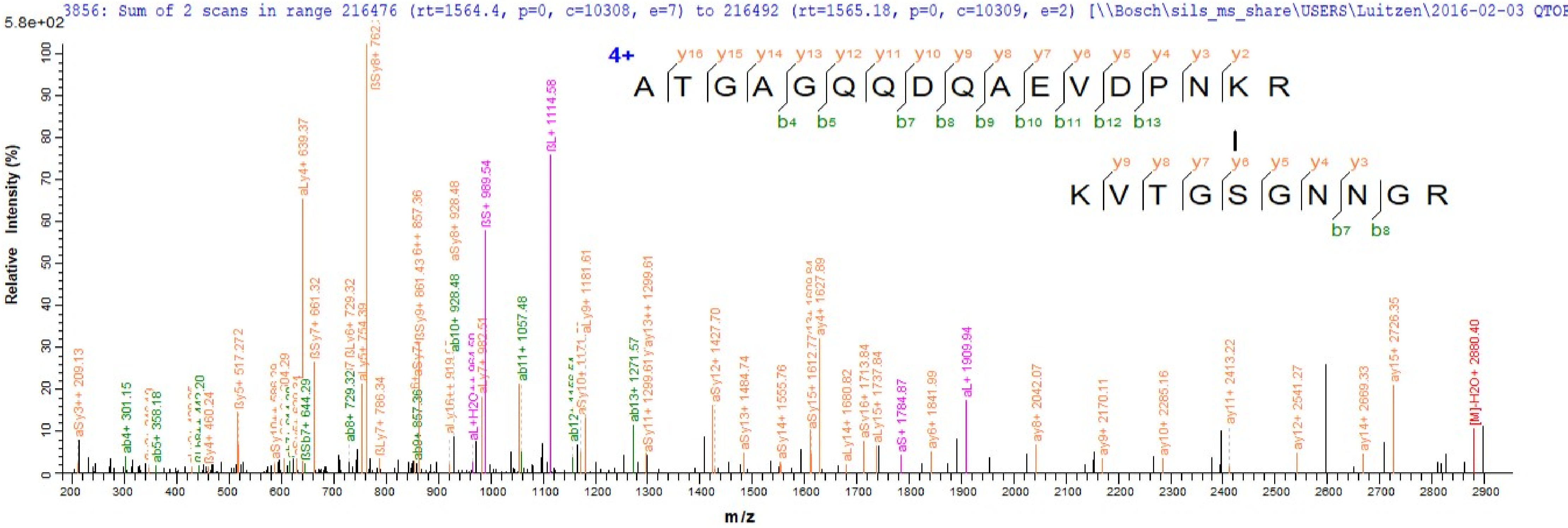
3856: Sum of 2 scans in range 216476 (rt=1564.4, p=0, c=10308, e=7) to 216492 (rt=1565.18, p=0, c=10309, e=2) [\\Bosch\sils_ms_share\USERS\Luitzen\2016-02-03 QTOF
5.8e+02
Relative Intensity (%)
m/z
4+
y16 y15 y14 y13 y12 y11 y10 y9 y8 y7 y6 y5 y4 y3 y2
A T G A G Q Q D Q A E V D P N K R
b4 b5 b7 b8 b9 b10 b11 b12 b13
y9 y8 y7 y6 y5 y4 y3
K V T G S G N N G R
b7 b8
aSy3++ 209.13
ab4+ 301.15
ab5+ 358.18
ßy4+ 460.24
ßy5+ 517.272
ßSb7+ 644.29
aLy4+ 639.37
ßSy7+ 661.32
ab8+ 729.32
aLy5+ 754.39
ßSy8+ 762.
ßLy7+ 786.34
ab9+ 857.36
aSy8+ 928.48
ab10+ 928.48
ßS+ 989.54
aLy7+ 982.51
ab11+ 1057.48
ßL+ 1114.58
aLy9+ 1181.61
ab13+ 1271.57
aSy11+ 1299.61
aSy12+ 1427.70
aSy13+ 1484.74
aSy14+ 1555.76
ay4+ 1627.89
aLy14+ 1680.82
aSy16+ 1713.84
aLy15+ 1737.84
aS+ 1784.87
ay6+ 1841.99
aL+ 1909.94
ay8+ 2042.07
ay9+ 2170.11
ay10+ 2285.16
ay11+ 2413.22
ay12+ 2541.27
ay14+ 2669.33
ay15+ 2726.35
[M]-H2O+ 2880.40

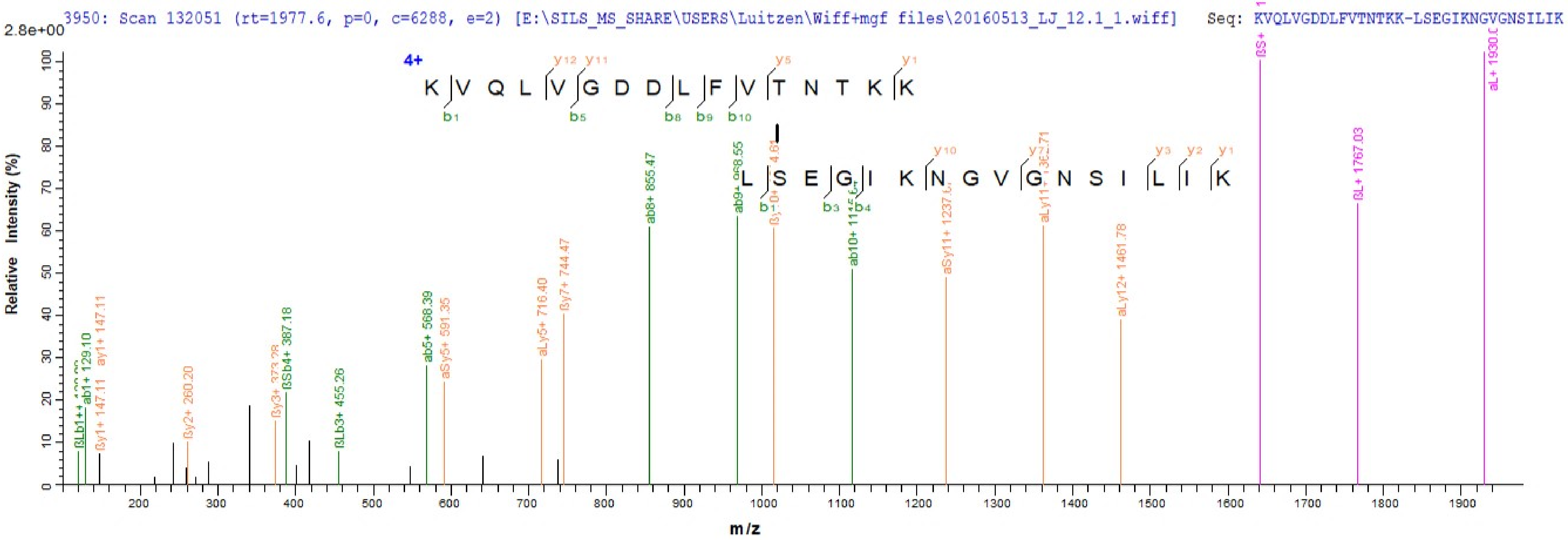

3950: Scan 132051 (rt=1977.6, p=0, c=6288, e=2) [E:\SILS_MS_SHARE\USERS\Luitzen\Wiff+mgf files\20160513_LJ_12.1_1.wiff] Seq: KVQLVGDDLFVTNTKK-LSEGIKNGVGNSILIK
2.8e+00
4+
K V Q L V G D D L F V T N T K K
L S E G I K N G V G N S I L I K
Relative Intensity (%)
m/z
ab8+ 855.47
ab5+ 568.39
aSy5+ 591.35
aLy5+ 716.40
ßy7+ 744.47
aSy11+ 1237.6
aLy12+ 1461.78
ßL+ 1767.03
ßLb3+ 455.26
ßSb4+ 387.18
ßy2+ 260.20
ab1+ 129.10
ay1+ 147.11
ßy1+ 147.11

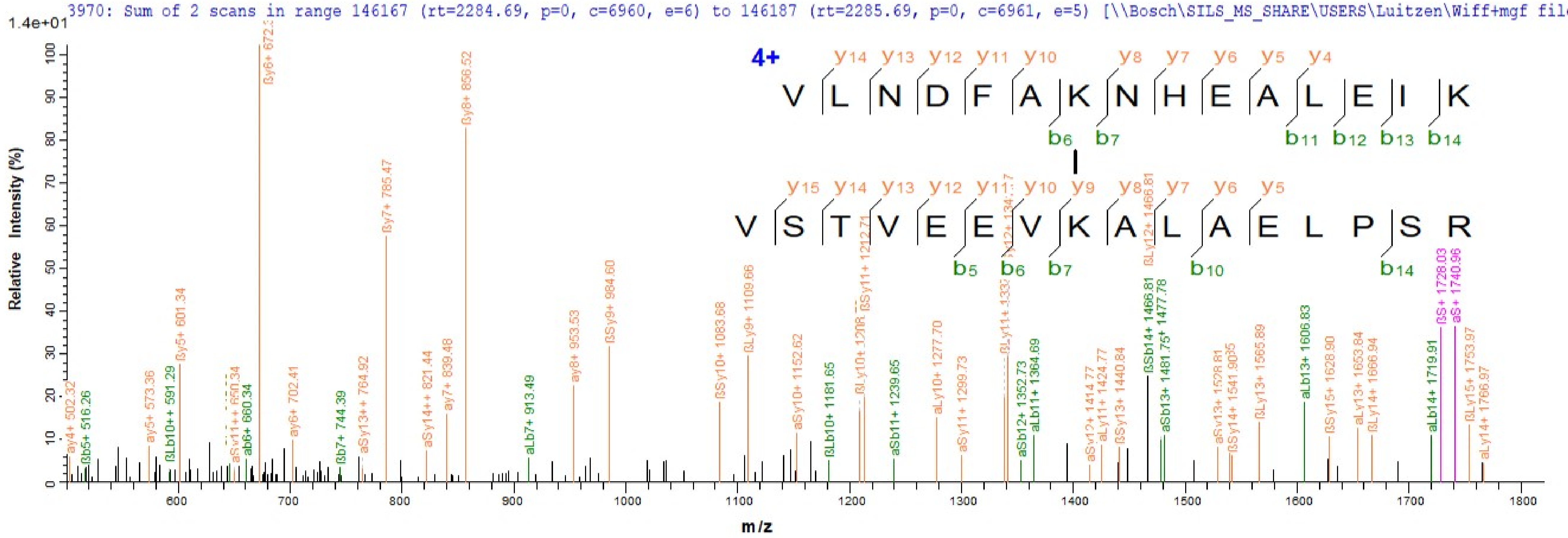
3970: Sum of 2 scans in range 146167 (rt=2284.69, p=0, c=6960, e=6) to 146187 (rt=2285.69, p=0, c=6961, e=5) [\\Bosch\SILS_MS_SHARE\USERS\Luitzen\Wiff+mgf files\2
1.4e+01
Relative Intensity (%)
m/z
4+
V L N D F A K N H E A L E I K
V S T V E E V K A L A E L P S R

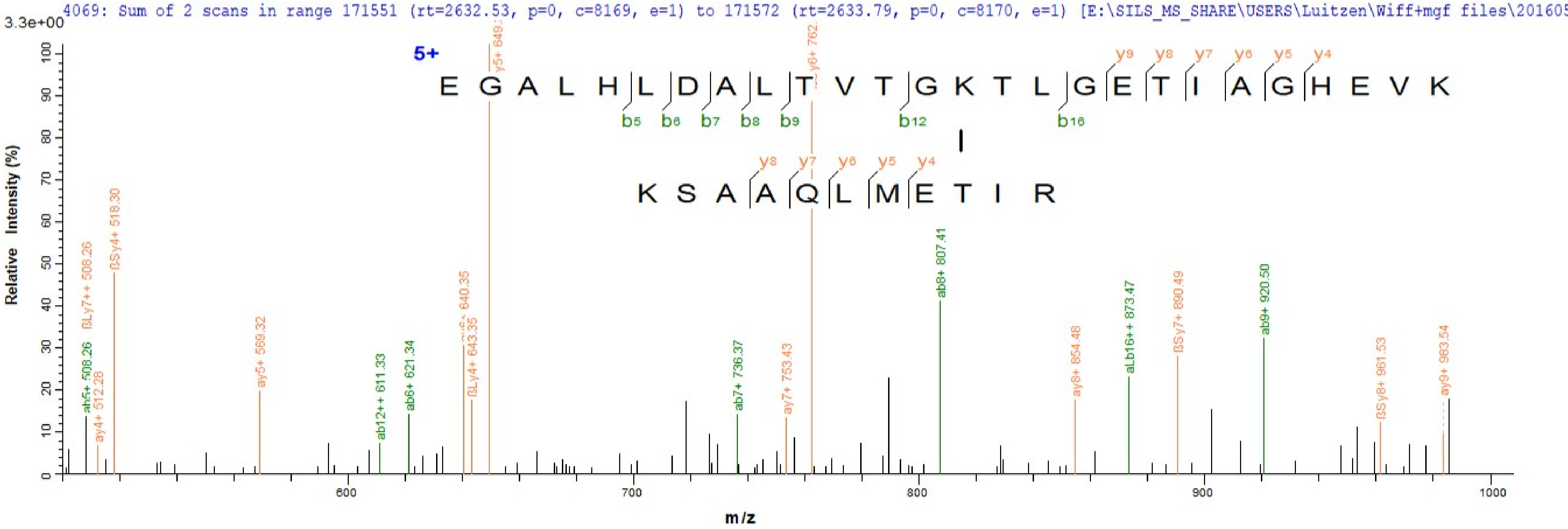

4069: Sum of 2 scans in range 171551 (rt=2632.53, p=0, c=8169, e=1) to 171572 (rt=2633.79, p=0, c=8170, e=1) [E:\SILS_MS_SHARE\USERS\Luitzen\Wiff+mgf files\201605
3.3e+00
5+
E G A L H L D A L T V T G K T L G E T I A G H E V K
K S A A Q L M E T I R
Relative Intensity (%)
m/z

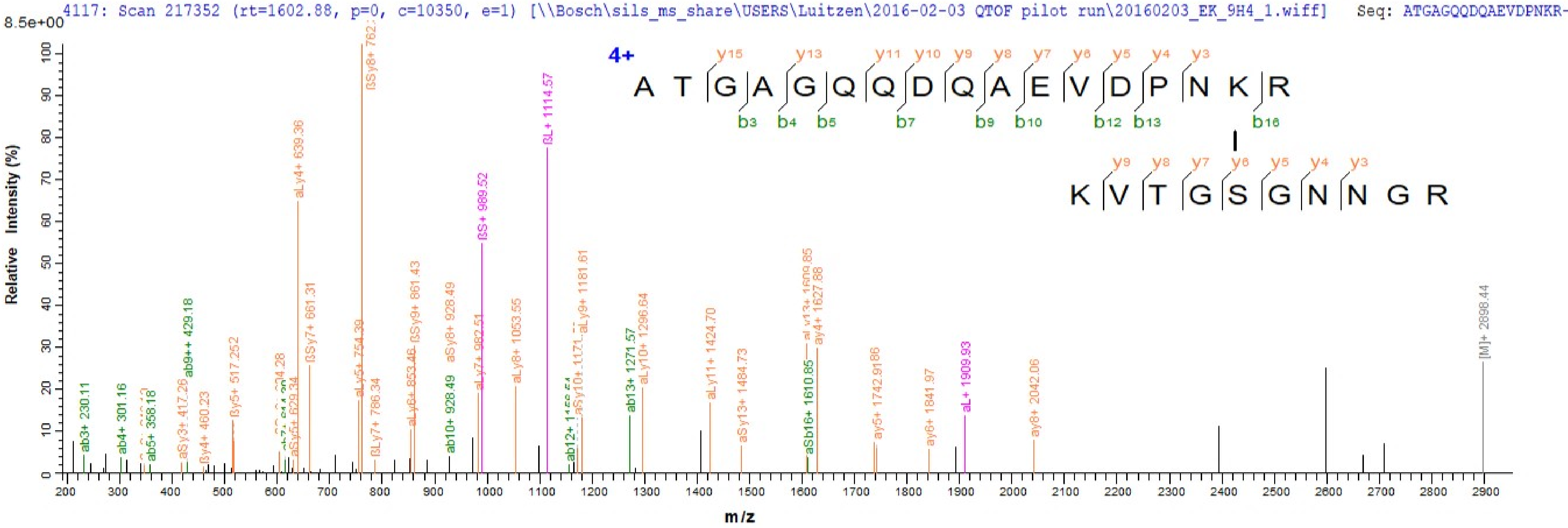
4117: Scan 217352 (rt=1602.88, p=0, c=10350, e=1) [\\Bosch\sils_ms_share\USERS\Luitzen\2016-02-03 QTOF pilot run\20160203_EK_9H4_1.wiff] Seq: ATGAGQQDQAEVDPNKR-
8.5e+00
4+
A T G A G Q Q D Q A E V D P N K R
y15 y13 y11 y10 y9 y8 y7 y6 y5 y4 y3
b3 b4 b5 b7 b9 b10 b12 b13 b16
K V T G S G N N G R
y9 y8 y7 y6 y5 y4 y3
Relative Intensity (%)
m/z
ab3+ 230.11
ab4+ 301.16
ab5+ 358.18
aSy3+ 417.26
ab9++ 429.18
ßy4+ 460.23
ßy5+ 517.252
aSy5+ 629.34
aLy4+ 639.36
ßSy7+ 661.31
aLy5+ 754.39
ßLy7+ 786.34
aLy6+ 853.46
ßSy9+ 861.43
ab10+ 928.49
aSy8+ 928.49
aLy7+ 982.51
ßS+ 989.52
aLy8+ 1053.55
ßL+ 1114.57
aLy9+ 1181.61
ab13+ 1271.57
aLy10+ 1296.64
aLy11+ 1424.70
aSy13+ 1484.73
aSb16+ 1610.85
ay4+ 1627.88
ay5+ 1742.9186
ay6+ 1841.97
aL+ 1909.93
ay8+ 2042.06
[M]+ 2898.44

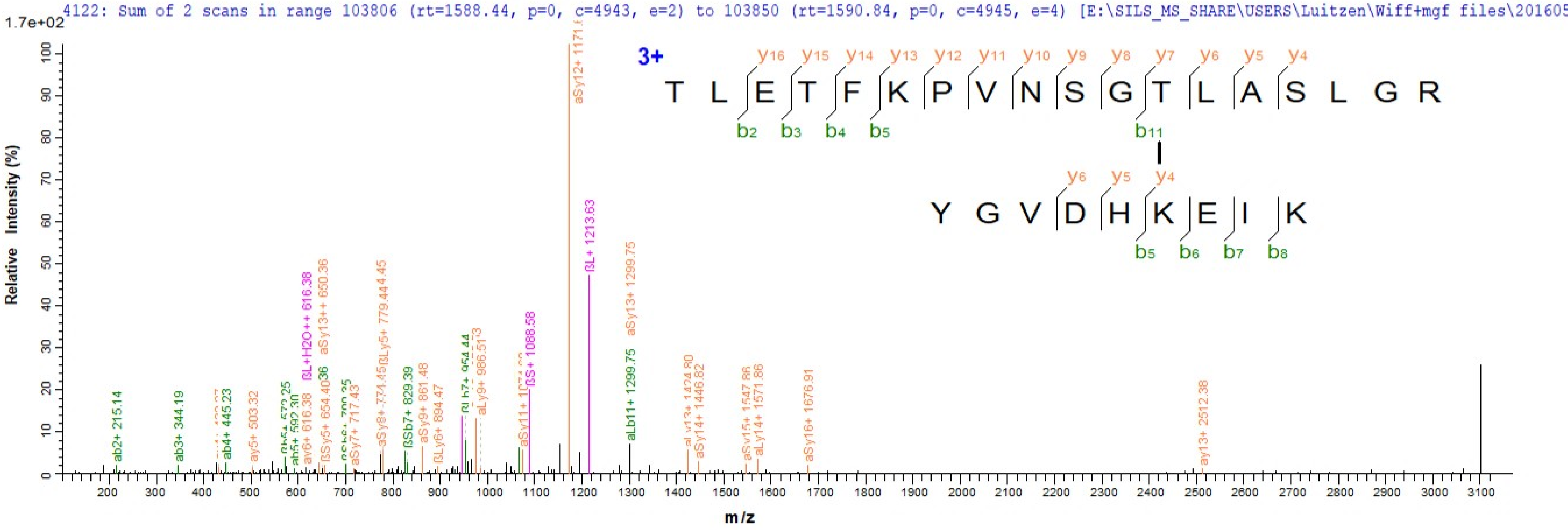
4122: Sum of 2 scans in range 103806 (rt=1588.44, p=0, c=4943, e=2) to 103850 (rt=1590.84, p=0, c=4945, e=4) [E:\SILS_MS_SHARE\USERS\Luitzen\Wiff+mgf files\201605
1.7e+02
Relative Intensity (%)
m/z
3+
y16 y15 y14 y13 y12 y11 y10 y9 y8 y7 y6 y5 y4
T L E T F K P V N S G T L A S L G R
b2 b3 b4 b5 b11
y6 y5 y4
Y G V D H K E I K
b5 b6 b7 b8
ab2+ 215.14
ab3+ 344.19
ab4+ 445.23
ay5+ 503.32
ay6+ 616.38
βL+H2O++ 616.38
aSy13++ 650.36
aSy7+ 717.43
aSy8+ 774.45
βSb7+ 829.39
aSy9+ 861.48
βLy6+ 894.47
aLy9+ 986.513
βS+ 1088.58
βL+ 1213.63
aSy13+ 1299.75
aLb11+ 1299.75
aSy14+ 1446.82
aLy14+ 1571.86
aSy16+ 1676.91
ay13+ 2512.38

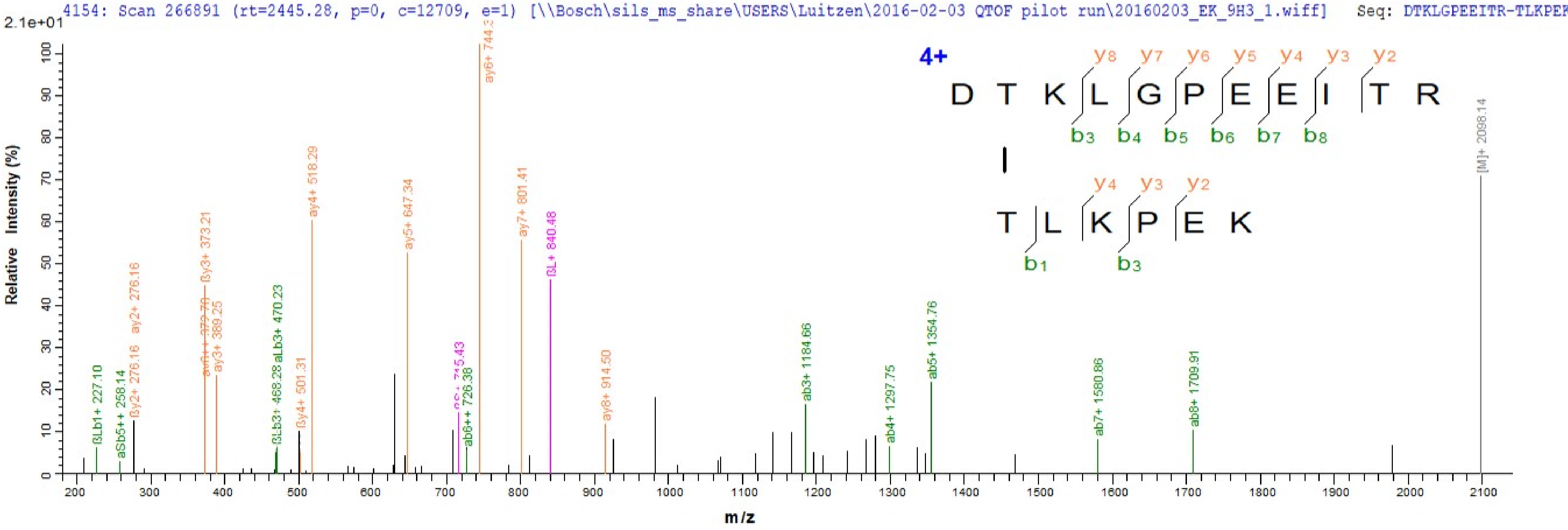

4154: Scan 266891 (rt=2445.28, p=0, c=12709, e=1) [\\Bosch\sils_ms_share\USERS\Luitzen\2016-02-03 QTOF pilot run\20160203_EK_9H3_1.wiff] Seq: DTKLGPEEITR-TLKPEK
2.1e+01
Relative Intensity (%)
m/z
4+
D T K L G P E E I T R
T L K P E K
y8 y7 y6 y5 y4 y3 y2
b3 b4 b5 b6 b7 b8
y4 y3 y2
b1 b3
ay6+ 744.3
ay4+ 518.29
ay5+ 647.34
ay7+ 801.41
ßL+ 840.48
ßy3+ 373.21
ay3+ 389.25
ay2+ 276.16
ßy2+ 276.16
ßLb1+ 227.10
aSb5++ 258.14
aLb3+ 470.23
ßLb3+ 468.28
ßy4+ 501.31
ab6++ 726.38
ay8+ 914.50
ab3+ 1184.66
ab4+ 1297.75
ab5+ 1354.76
ab7+ 1580.86
ab8+ 1709.91
[M]+ 2098.14

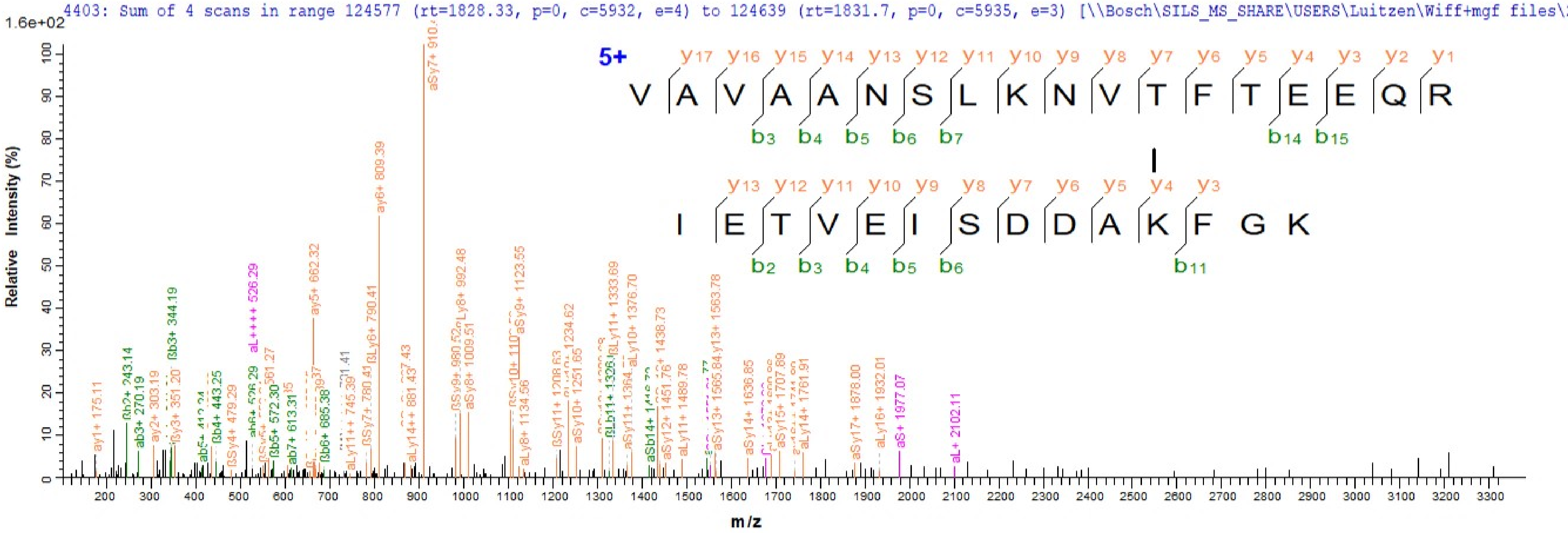
4403: Sum of 4 scans in range 124577 (rt=1828.33, p=0, c=5932, e=4) to 124639 (rt=1831.7, p=0, c=5935, e=3) [\\Bosch\SILS_MS_SHARE\USERS\Luitzen\Wiff+mgf files\20
1.6e+02
Relative Intensity (%)
m/z
5+
y17 y16 y15 y14 y13 y12 y11 y10 y9 y8 y7 y6 y5 y4 y3 y2 y1
V A V A A N S L K N V T F T E E Q R
b3 b4 b5 b6 b7 b14 b15
y13 y12 y11 y10 y9 y8 y7 y6 y5 y4 y3
I E T V E I S D D A K F G K
b2 b3 b4 b5 b6 b11
aSy7+ 910.4
ay6+ 809.39
ay5+ 662.32
aL++++ 526.29
ßb3+ 344.19
ßb2+ 243.14
ab3+ 270.19
ay2+ 303.19
ay1+ 175.11
ßb4+ 443.25
ßb5+ 572.30
ab7+ 613.31
ßb6+ 685.38
ßSy4+ 479.29
aLy11++ 745.39
ßSy7+ 780.41
ßLy6+ 790.41
aLy14++ 881.43
ßSy9+ 980.52
aLy8+ 992.48
aSy8+ 1009.51
aSy9+ 1123.55
aLy8+ 1134.56
ßSy11+ 1208.63
aSy10+ 1251.65
1234.62
ßLy11+ 1333.69
aLy10+ 1376.70
1438.73
aSy12+ 1451.76
aLy11+ 1489.78
aSy13+ 1565.84
y13+ 1563.78
aSy14+ 1636.85
aSy15+ 1707.89
aLy14+ 1761.91
aSy17+ 1878.00
aLy16+ 1932.01
aS+ 1977.07
aL+ 2102.11

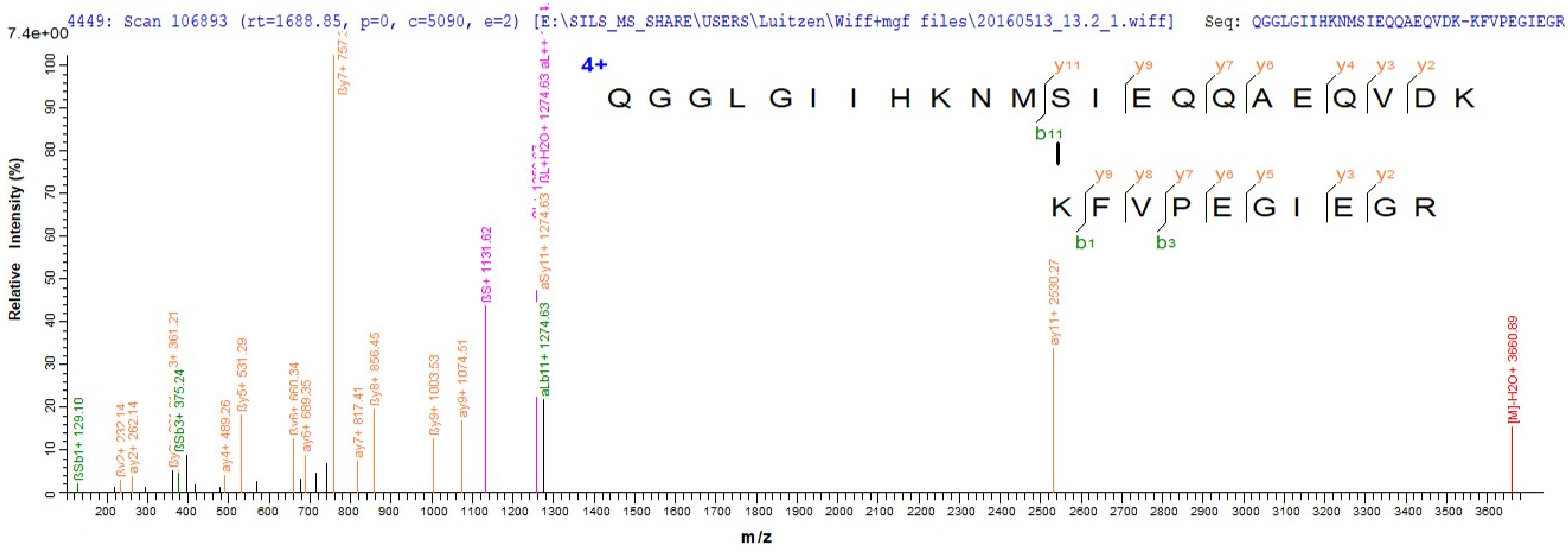
4449: Scan 106893 (rt=1688.85, p=0, c=5090, e=2) [E:\SILS_MS_SHARE\USERS\Luitzen\Wiff+mgf files\20160513_13.2_1.wiff] Seq: QGGLGIIHKNMSIEQQAEQVDK-KFVPEGIEGR
7.4e+00
4+
Q G G L G I I H K N M S I E Q Q A E Q V D K
y11 y9 y7 y6 y4 y3 y2
b11
K F V P E G I E G R
y9 y8 y7 y6 y5 y3 y2
b1 b3
Relative Intensity (%)
m/z
ßSb1+ 129.10
ay2+ 262.14
ßSb3+ 375.24
3+ 361.21
ay4+ 489.26
ßy5+ 531.29
ay6+ 689.35
ay7+ 817.41
ßy8+ 856.45
ßy9+ 1003.53
ay9+ 1074.51
ßS+ 1131.62
aLb11+ 1274.63
aSy11+ 1274.63
ßL+H2O+ 1274.63
ay11+ 2530.27
[M]-H2O+ 3660.89

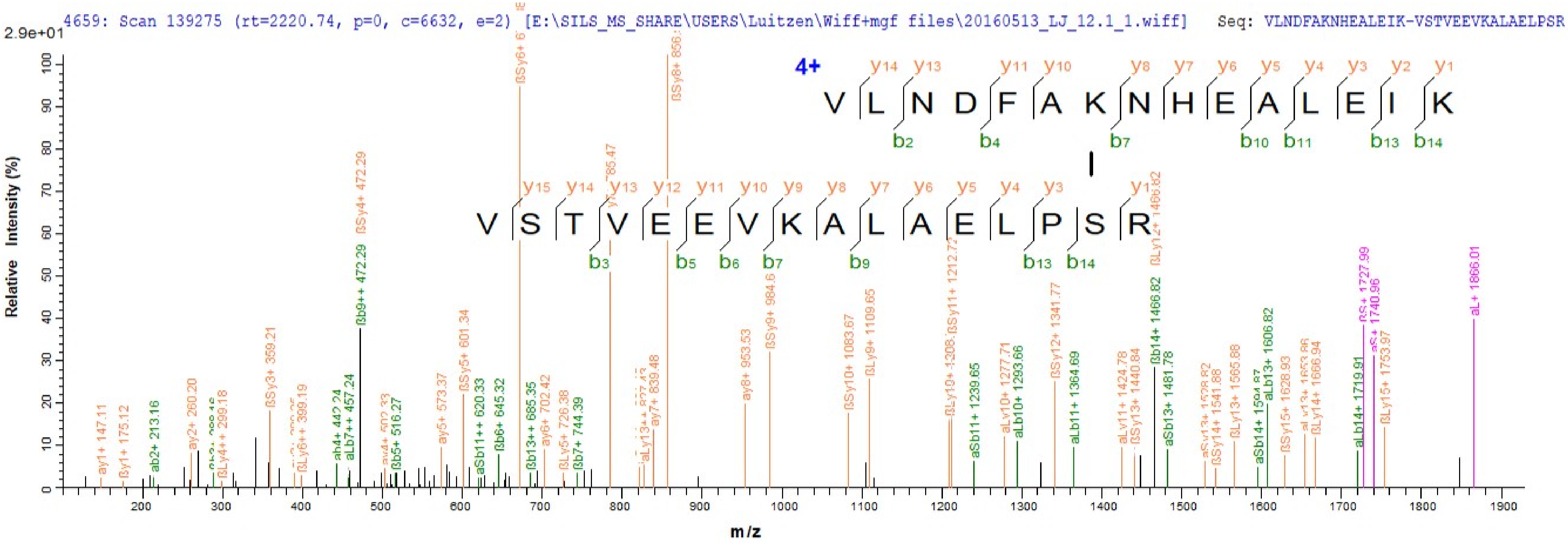
4659: Scan 139275 (rt=2220.74, p=0, c=6632, e=2) [E:\SILS_MS_SHARE\USERS\Luitzen\Wiff+mgf files\20160513_LJ_12.1_1.wiff] Seq: VLNDFAKNHEALEIK-VSTVEEVKALAELPSR
2.9e+01
Relative Intensity (%)
m/z
4+
V L N D F A K N H E A L E I K
V S T V E E V K A L A E L P S R
ay1+ 147.11
ßy1+ 175.12
ab2+ 213.16
ay2+ 260.20
ßLy4++ 299.18
ßSy3+ 359.21
ßLy6++ 399.19
aLb7++ 457.24
ßb9++ 472.29
ßSy4+ 472.29
ßb5+ 516.27
ay5+ 573.37
ßSy5+ 601.34
aSb11++ 620.33
ßb6+ 645.32
ßb13++ 685.35
ay6+ 702.42
ßLy5+ 726.38
ßb7+ 744.39
ay7+ 839.48
ay8+ 953.53
ßSy9+ 984.6
ßSy10+ 1083.67
ßLy9+ 1109.65
ßSy11+ 1212.72
aSb11+ 1239.65
aLy10+ 1277.71
aLb10+ 1293.66
ßSy12+ 1341.77
aLb11+ 1364.69
aLy11+ 1424.78
ßSy13+ 1440.84
ßb14+ 1466.82
aSb13+ 1481.78
ßSy14+ 1541.88
ßLy13+ 1565.88
aLb13+ 1606.82
ßSy15+ 1628.93
ßLy14+ 1666.94
aLb14+ 1719.91
ßS+ 1727.99
aS+ 1740.96
ßLy15+ 1753.97
aL+ 1866.01

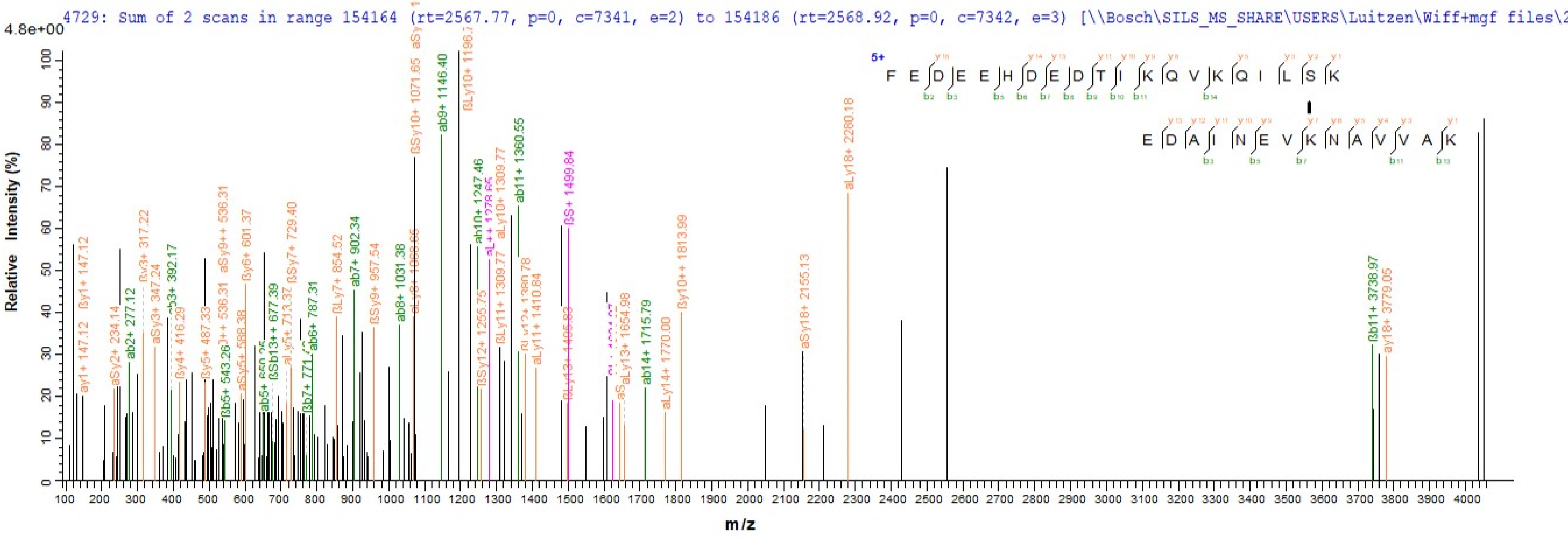
4729: Sum of 2 scans in range 154164 (rt=2567.77, p=0, c=7341, e=2) to 154186 (rt=2568.92, p=0, c=7342, e=3) [\\Bosch\SILS_MS_SHARE\USERS\Luitzen\Wiff+mgf files\2
4.8e+00
Relative Intensity (%)
m/z
0
10
20
30
40
50
60
70
80
90
100
100
200
300
400
500
600
700
800
900
1000
1100
1200
1300
1400
1500
1600
1700
1800
1900
2000
2100
2200
2300
2400
2500
2600
2700
2800
2900
3000
3100
3200
3300
3400
3500
3600
3700
3800
3900
4000
5+
F E D E E H D E D T I K Q V K Q I L S K
E D A I N E V K N A V V A K

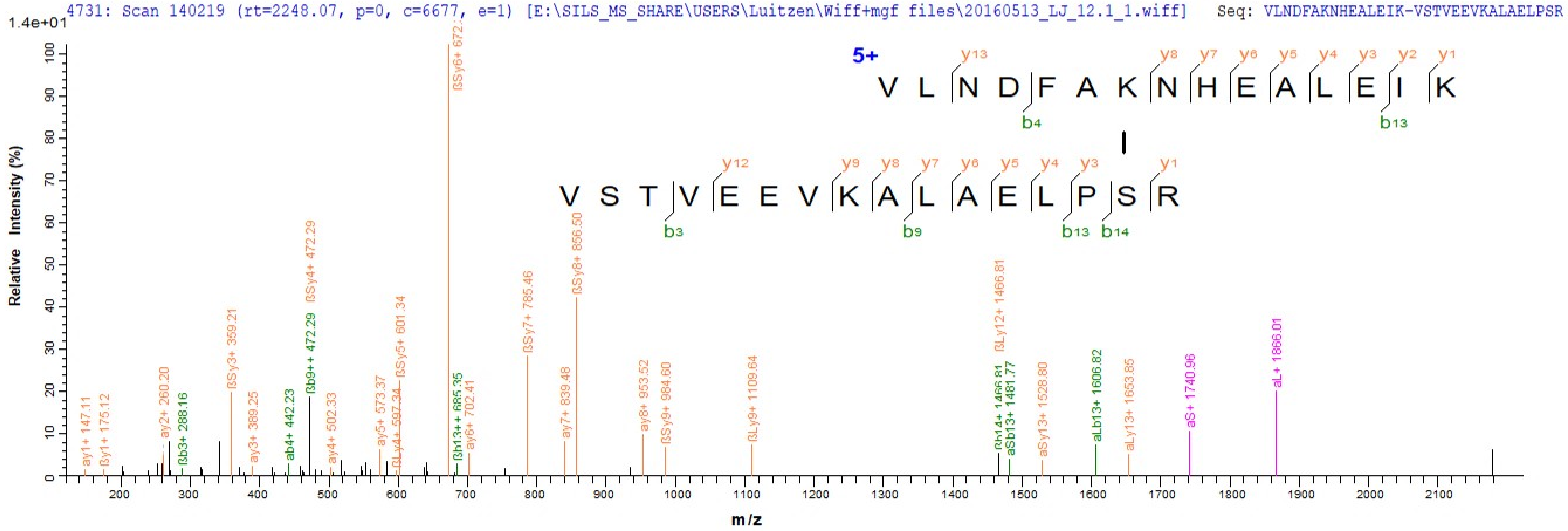
4731: Scan 140219 (rt=2248.07, p=0, c=6677, e=1) [E:\SILS_MS_SHARE\USERS\Luitzen\Wiff+mgf files\20160513_LJ_12.1_1.wiff] Seq: VLNDFAKNHEALEIK-VSTVEEVKALAELPSR
1.4e+01
5+
V L N D F A K N H E A L E I K
V S T V E E V K A L A E L P S R
Relative Intensity (%)
m/z
ay1+ 147.11
βy1+ 175.12
ay2+ 260.20
βb3+ 288.16
βSy3+ 359.21
ay3+ 389.25
ab4+ 442.23
βb9++ 472.29
βSy4+ 472.29
ay4+ 502.33
ay5+ 573.37
βLy4+ 597.34
βSy5+ 601.34
βb13++ 685.35
ay6+ 702.41
βSy7+ 785.46
ay7+ 839.48
βSy8+ 856.50
ay8+ 953.52
βSy9+ 984.60
βLy9+ 1109.64
βLy12+ 1466.81
aSb13+ 1481.77
aSy13+ 1528.80
aLb13+ 1606.82
aLy13+ 1653.85
aS+ 1740.96
aL+ 1866.01

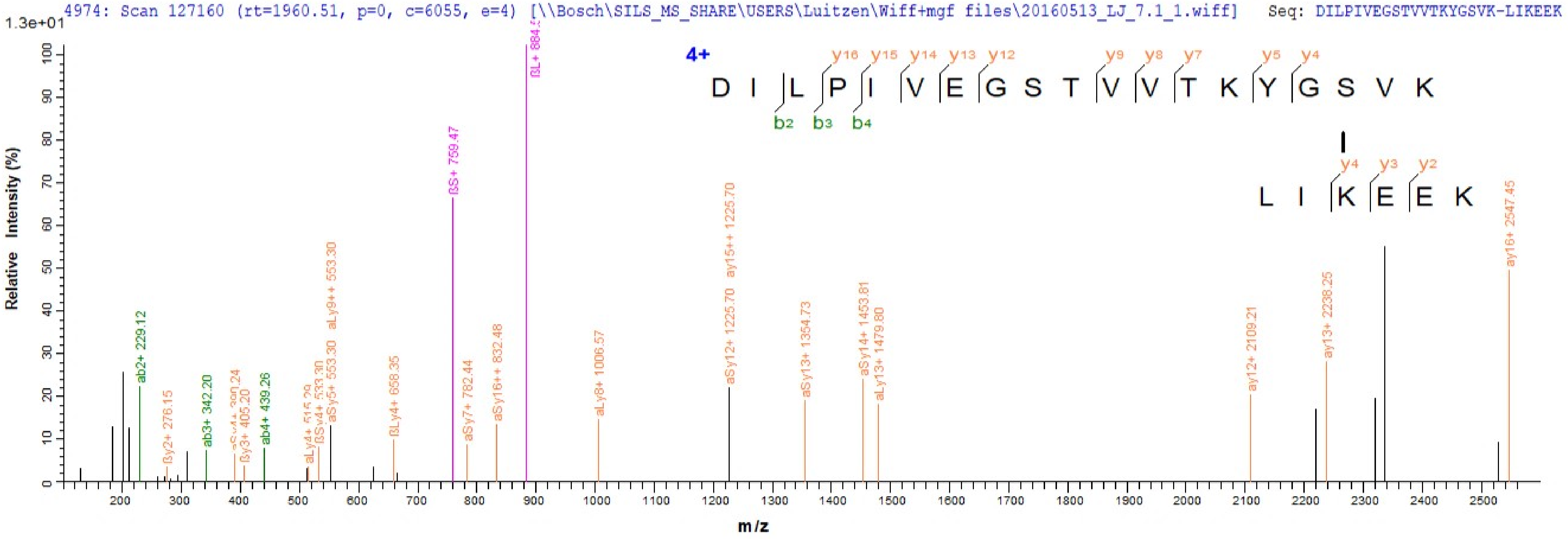
4974: Scan 127160 (rt=1960.51, p=0, c=6055, e=4) [\\Bosch\SILS_MS_SHARE\USERS\Luitzen\Wiff+mgf files\20160513_LJ_7.1_1.wiff] Seq: DILPIVEGSTVVTKYGSVK-LIKEEK
1.3e+01
Relative Intensity (%)
m/z
4+
D I L P I V E G S T V V T K Y G S V K
y16 y15 y14 y13 y12 y9 y8 y7 y5 y4
b2 b3 b4
L I K E E K
y4 y3 y2
ab2+ 229.12
βy2+ 276.15
ab3+ 342.20
βy3+ 405.20
ab4+ 439.26
βSy4+ 533.30
aSy5+ 553.30
aLy9++ 553.30
βLy4+ 658.35
βS+ 759.47
aSy7+ 782.44
aSy16++ 832.48
aLy8+ 1006.57
aSy12+ 1225.70
ay15++ 1225.70
aSy13+ 1354.73
aSy14+ 1453.81
aLy13+ 1479.80
ay12+ 2109.21
ay13+ 2238.25
ay16+ 2547.45

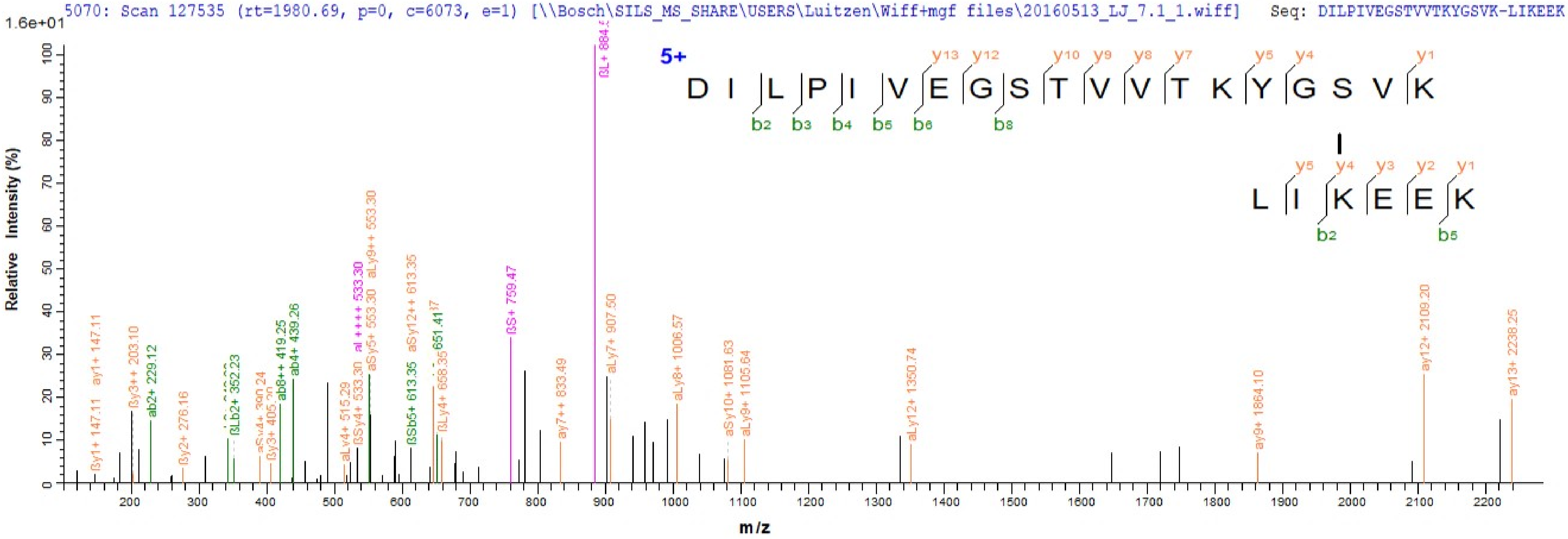
5070: Scan 127535 (rt=1980.69, p=0, c=6073, e=1) [\\Bosch\SILS_MS_SHARE\USERS\Luitzen\Wiff+mgf files\20160513_LJ_7.1_1.wiff] Seq: DILPIVEGSTVVTKYGSVK-LIKEEK
1.6e+01
5+
D I L P I V E G S T V V T K Y G S V K
y13 y12 y10 y9 y8 y7 y5 y4 y1
b2 b3 b4 b5 b6 b8
L I K E E K
y5 y4 y3 y2 y1
b2 b5
Relative Intensity (%)
m/z
ßy1+ 147.11
ay1+ 147.11
ßy3++ 203.10
ab2+ 229.12
ßy2+ 276.16
ßLb2+ 352.23
ßy3+ 405.20
ab8++ 419.25
ab4+ 439.26
aLy4+ 515.29
ßSy4+ 533.30
aL++++ 533.30
aSy5+ 553.30
aLy9++ 553.30
ßSb5+ 613.35
aSy12++ 613.35
ßLy4+ 658.35
651.41
ßS+ 759.47
ay7++ 833.49
ßL+ 884.5
aLy7+ 907.50
aLy8+ 1006.57
aSy10+ 1081.63
aLy9+ 1105.64
aLy12+ 1350.74
ay9+ 1864.10
ay12+ 2109.20
ay13+ 2238.25

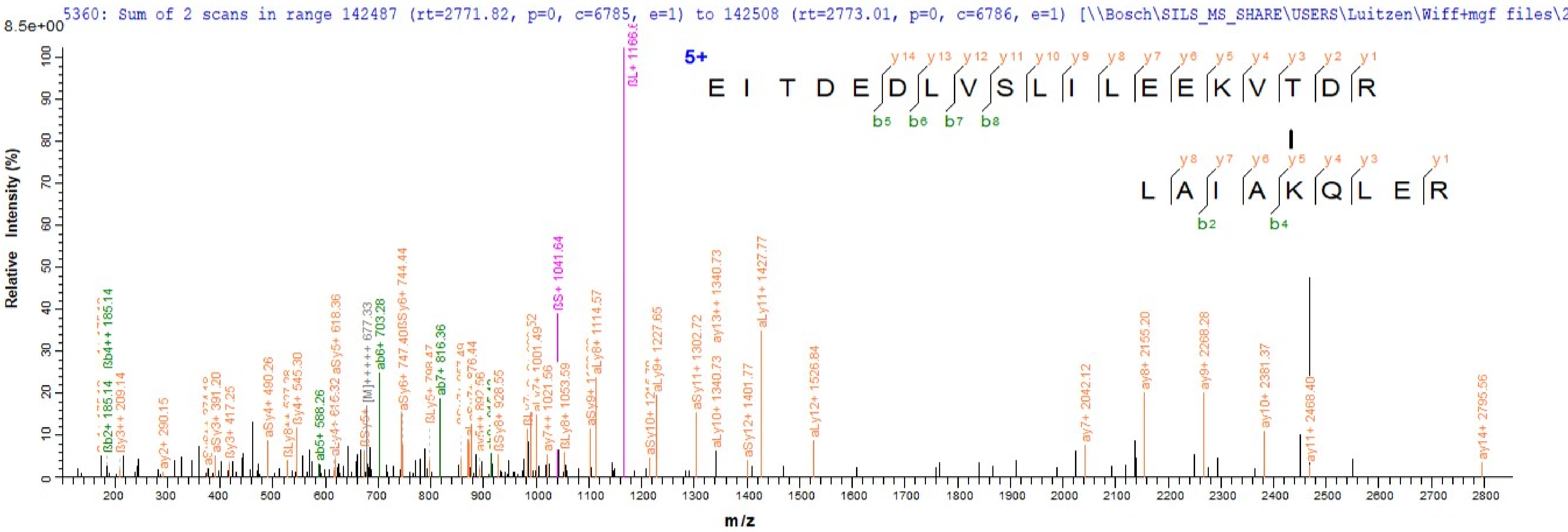

5360: Sum of 2 scans in range 142487 (rt=2771.82, p=0, c=6785, e=1) to 142508 (rt=2773.01, p=0, c=6786, e=1) [\\Bosch\SILS_MS_SHARE\USERS\Luitzen\Wiff+mgf files\2
8.5e+00
Relative Intensity (%)
m/z
5+
E I T D E D L V S L I L E E K V T D R
y14 y13 y12 y11 y10 y9 y8 y7 y6 y5 y4 y3 y2 y1
b5 b6 b7 b8
L A I A K Q L E R
y8 y7 y6 y5 y4 y3 y1
b2 b4
ßL+ 1166.6
ßS+ 1041.64
aLy11+ 1427.77
ay13++ 1340.73
aSy6+ 744.44
aSy5+ 618.36
ab6+ 703.28
ab7+ 816.36
ab5+ 588.26
aSy4+ 490.26
ßb4++ 185.14
ßb2+ 185.14
ßy3++ 209.14
ay2+ 290.15
aSy3+ 391.20
ßy3+ 417.25
ßy4+ 545.30
aLy4+ 616.32
aLy8+ 1114.57
aLy9+ 1227.65
aSy11+ 1302.72
aLy10+ 1340.73
aSy12+ 1401.77
aLy12+ 1526.84
ay7+ 2042.12
ay8+ 2155.20
ay9+ 2268.28
ay10+ 2381.37
ay11+ 2468.40
ay14+ 2795.56
ay7++ 1021.56
ßLy8+ 1053.59
ßSy8+ 928.55
ay5++ 892.56

5486: Scan 127851 (rt=2122.78, p=0, c=6088, e=2) [\\Bosch\SILS_MS_SHARE\USERS\Luitzen\Wiff+mgf files\20160513_LJ_8.1_1.wiff] Seq: VQLVGDDLFVTNTKK-LSEGIKNGVGNSII

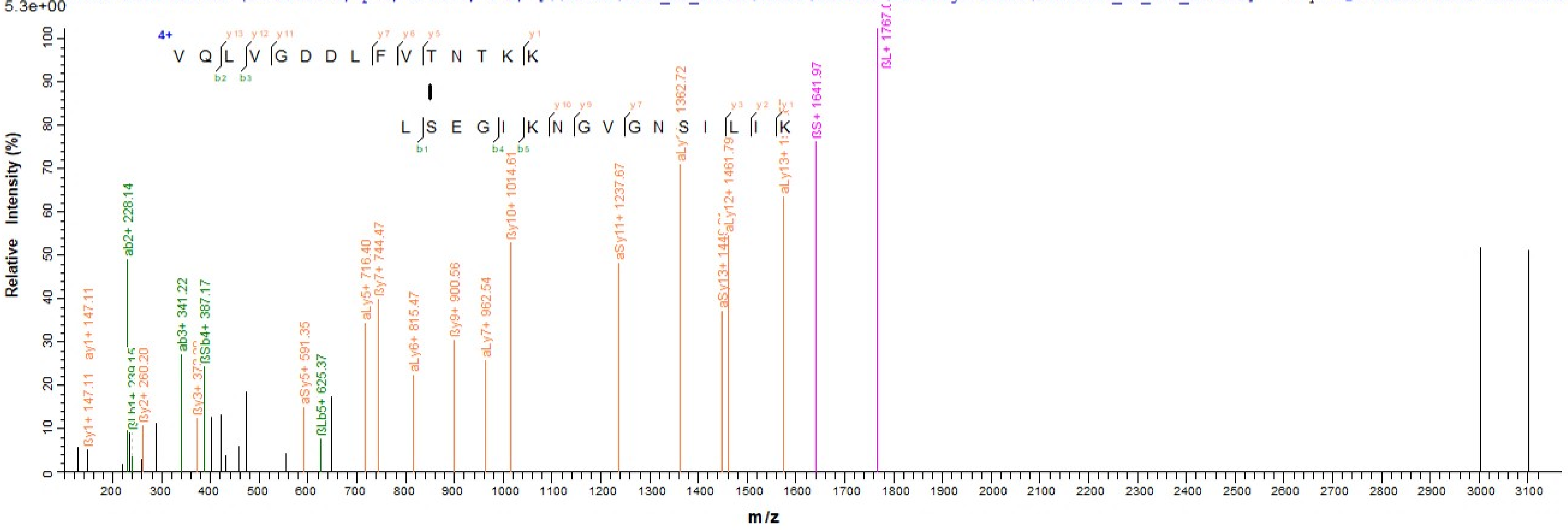

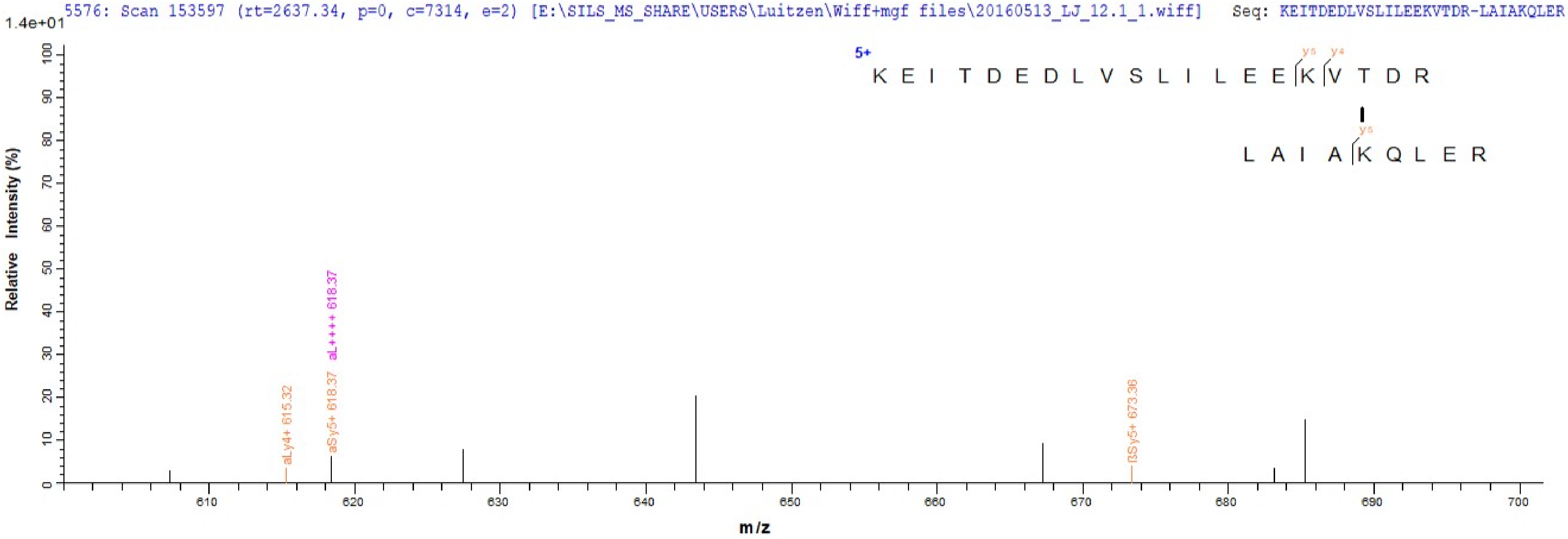
5576: Scan 153597 (rt=2637.34, p=0, c=7314, e=2) [E:\SILS_MS_SHARE\USERS\Luitzen\Wiff+mgf files\20160513_LJ_12.1_1.wiff] Seq: KEITDEDLVSLILEEKVTDR-LAIAKQLER
1.4e+01
5+
K E I T D E D L V S L I L E E K V T D R
y5 y4
y5
L A I A K Q L E R
Relative Intensity (%)
aLy4+ 615.32
aSy5+ 618.37
aL++++ 618.37
ßSy5+ 673.36
m/z

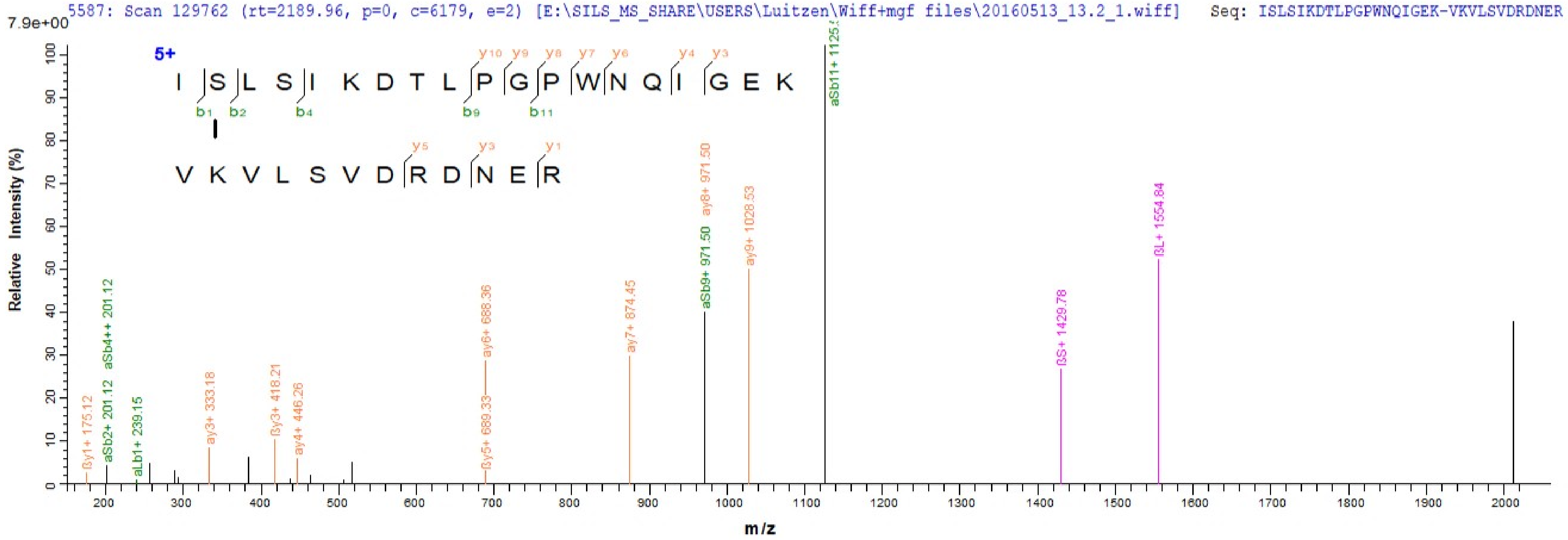
5587: Scan 129762 (rt=2189.96, p=0, c=6179, e=2) [E:\SILS_MS_SHARE\USERS\Luitzen\Wiff+mgf files\20160513_13.2_1.wiff] Seq: ISLSIKDTLPGPWNQIGEK-VKVLSVDRDNER
7.9e+00
5+
I S L S I K D T L P G P W N Q I G E K
y10 y9 y8 y7 y6 y4 y3
b1 b2 b4 b9 b11
V K V L S V D R D N E R
y5 y3 y1
Relative Intensity (%)
m/z
ßy1+ 175.12
aSb2+ 201.12
aSb4++ 201.12
aLb1+ 239.15
ay3+ 333.18
ßy3+ 418.21
ay4+ 446.26
ßy5+ 689.33
ay6+ 688.36
ay7+ 874.45
aSb9+ 971.50
ay8+ 971.50
ay9+ 1028.53
aSb11+ 1125
ßS+ 1429.78
ßL+ 1554.84

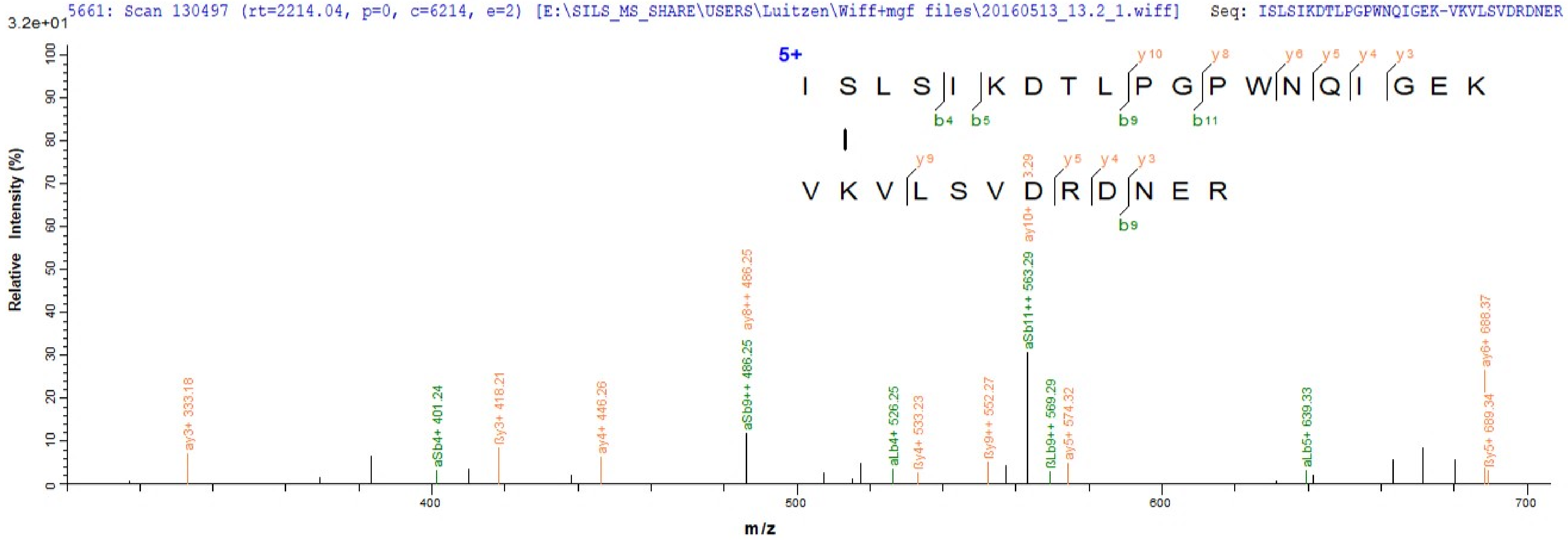
5661: Scan 130497 (rt=2214.04, p=0, c=6214, e=2) [E:\SILS_MS_SHARE\USERS\Luitzen\Wiff+mgf files\20160513_13.2_1.wiff]
Seq: ISLSIKDTLPGPWNQIGEK-VKVLSVDRDNER
3.2e+01
5+
I S L S I K D T L P G P W N Q I G E K
V K V L S V D R D N E R
y10 y8 y6 y5 y4 y3
b4 b5 b9 b11
y9 y5 y4 y3
b9
Relative Intensity (%)
m/z
ay3+ 333.18
aSb4+ 401.24
ßy3+ 418.21
ay4+ 446.26
aSb9++ 486.25
ay8++ 486.25
aLb4+ 526.25
ßy4+ 533.23
ßy9++ 552.27
aSb11++ 563.29
ay10+ 563.29
ßLb9++ 569.29
ay5+ 574.32
aLb5+ 639.33
ßy5+ 689.34
ay6+ 688.37

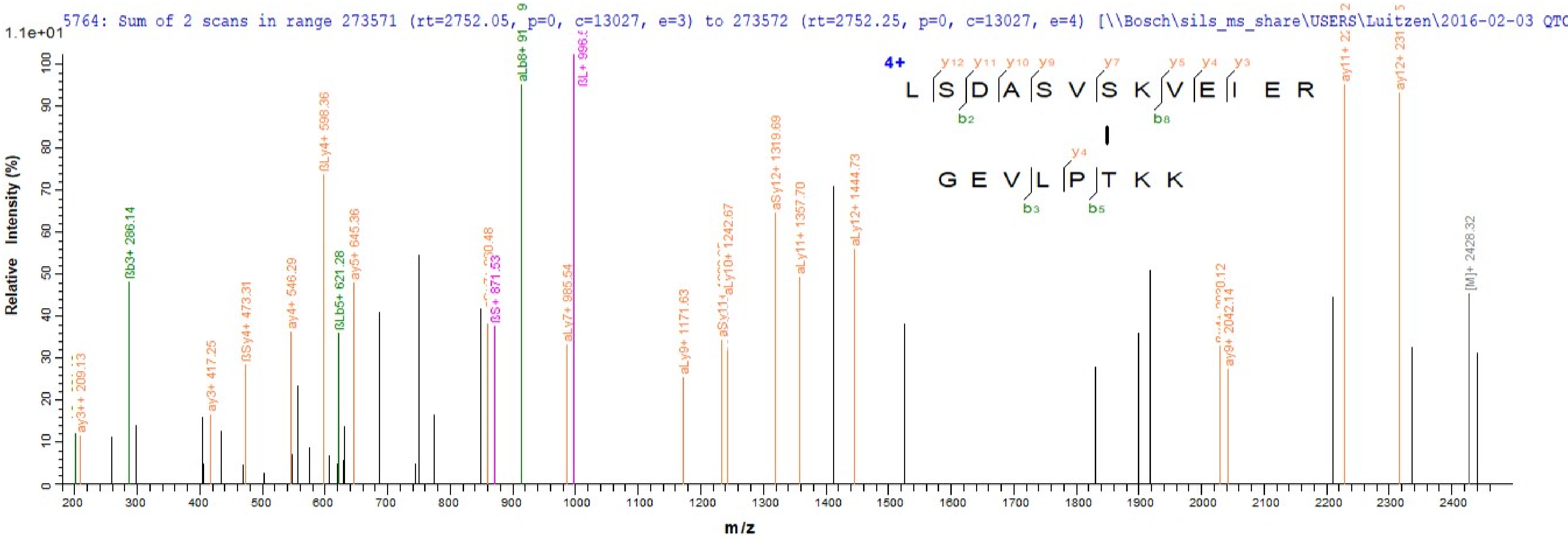

5764: Sum of 2 scans in range 273571 (rt=2752.05, p=0, c=13027, e=3) to 273572 (rt=2752.25, p=0, c=13027, e=4) [\\Bosch\sils_ms_share\USERS\Luitzen\2016-02-03 QTC
1.1e+01
Relative Intensity (%)
m/z
4+
L S D A S V S K V E I E R
G E V L P T K K
ay3++ 209.13
ßb3+ 286.14
ay3+ 417.25
ßSy4+ 473.31
ay4+ 546.29
ßLy4+ 598.36
ßLb5+ 621.28
ay5+ 645.36
ßS+ 871.53
aLy7+ 985.54
ßL+ 996.5
aLy9+ 1171.63
aLy10+ 1242.67
aSy12+ 1319.69
aLy11+ 1357.70
aLy12+ 1444.73
ay9+ 2042.14
[M]+ 2428.32

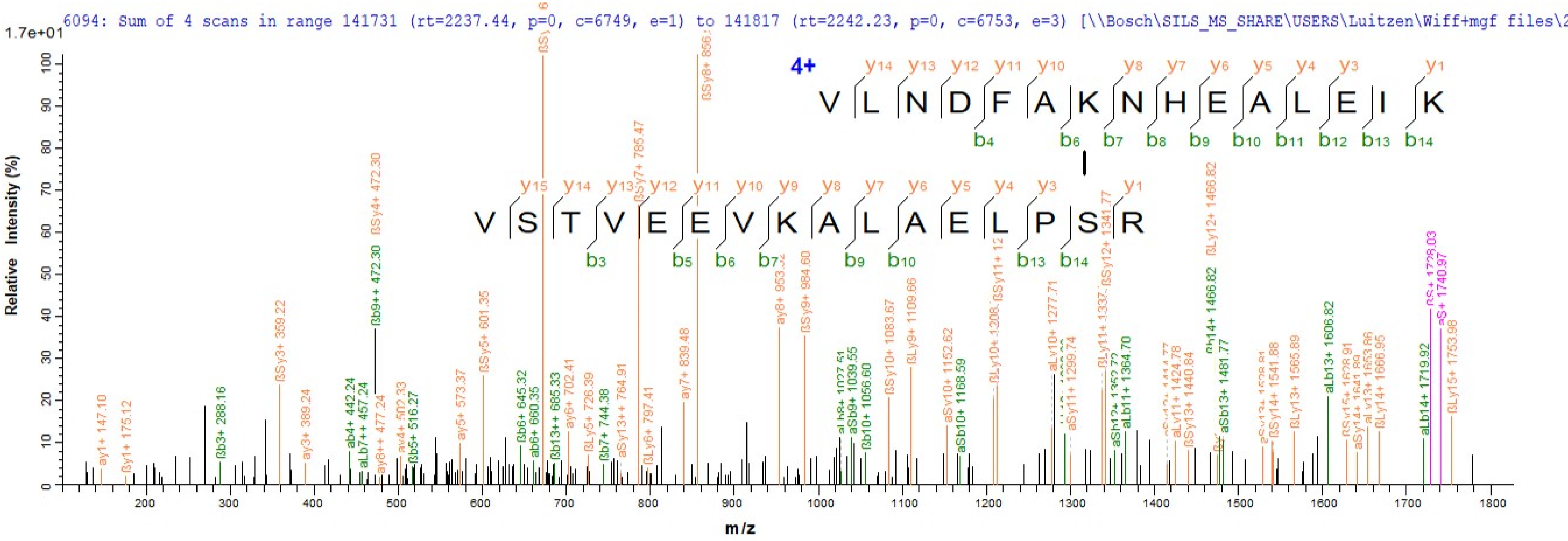
6094: Sum of 4 scans in range 141731 (rt=2237.44, p=0, c=6749, e=1) to 141817 (rt=2242.23, p=0, c=6753, e=3) [\\Bosch\SILS_MS_SHARE\USERS\Luitzen\Wiff+mgf files\2
1.7e+01
Relative Intensity (%)
m/z
4+
V L N D F A K N H E A L E I K
V S T V E E V K A L A E L P S R

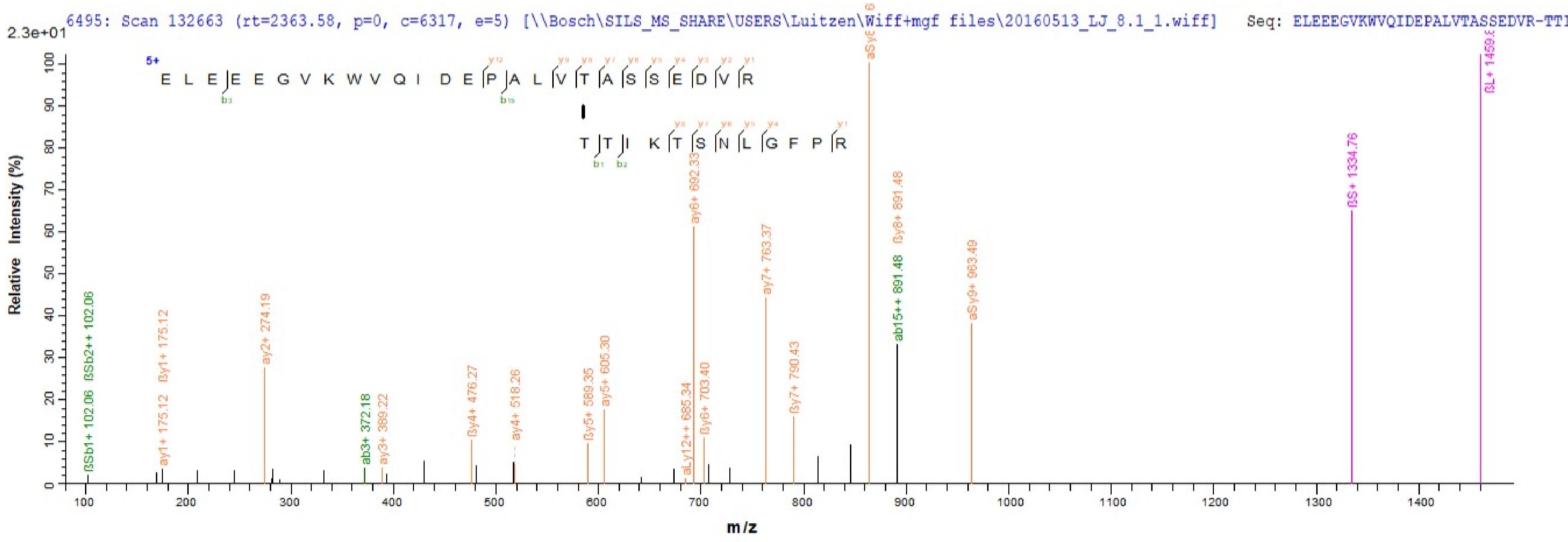
6495: Scan 132663 (rt=2363.58, p=0, c=6317, e=5) [\\Bosch\SILS_MS_SHARE\USERS\Luitzen\Wiff+mgf files\20160513_LJ_8.1_1.wiff]   Seq: ELEEEGVKWVQIDEPALVTASSEDVR-TTI
2.3e+01
5+
E L E E E G V K W V Q I D E P A L V T A S S E D V R
T T I K T S N L G F P R
Relative Intensity (%)
m/z
ßSb1+ 102.06
ßSb2++ 102.06
ay1+ 175.12
ßy1+ 175.12
ay2+ 274.19
ab3+ 372.18
ay3+ 389.22
ßy4+ 476.27
ay4+ 518.26
ßy5+ 589.35
ay5+ 605.30
aLy12++ 685.34
ay6+ 692.33
ßy6+ 703.40
ay7+ 763.37
ßy7+ 790.43
ab15++ 891.48
ßy8+ 891.48
aSy9+ 963.49
ßS+ 1334.76

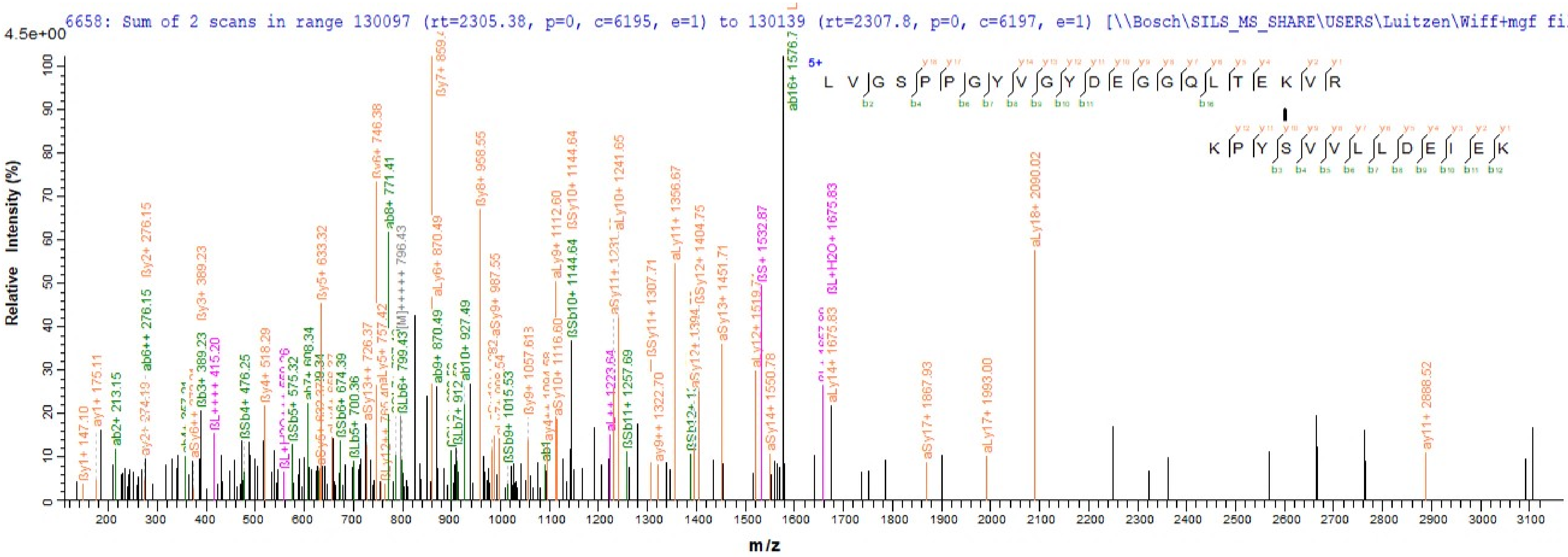

6658: Sum of 2 scans in range 130097 (rt=2305.38, p=0, c=6195, e=1) to 130139 (rt=2307.8, p=0, c=6197, e=1) [\\Bosch\SILS_MS_SHARE\USERS\Luitzen\Wiff+mgf files\20
4.5e+00
Relative Intensity (%)
m/z
5+
L V G S P P G Y V G Y D E G G Q L T E K V R
K P Y S V V L L D E I E K
ßy7+ 859.4
ab16+ 1576.7
ßy6+ 746.38
ab8+ 771.41
ßy8+ 958.55
aLy18+ 2090.02
ßSy10+ 1144.64
aLy10+ 1241.65
aLy9+ 1112.60
aLy6+ 870.49
ßL+H2O+ 1675.83
ßS+ 1532.87
aLy11+ 1356.67
ßy2+ 276.15
ßy3+ 389.23
ßy5+ 633.32
aSy13+ 1451.71
ab6++ 276.15
ab9+ 870.49
ab10+ 927.49
ay11+ 2888.52
aSy17+ 1867.93
aLy17+ 1993.00
aLy14+ 1675.83
ab2+ 213.15
ay1+ 175.11
ßy1+ 147.10

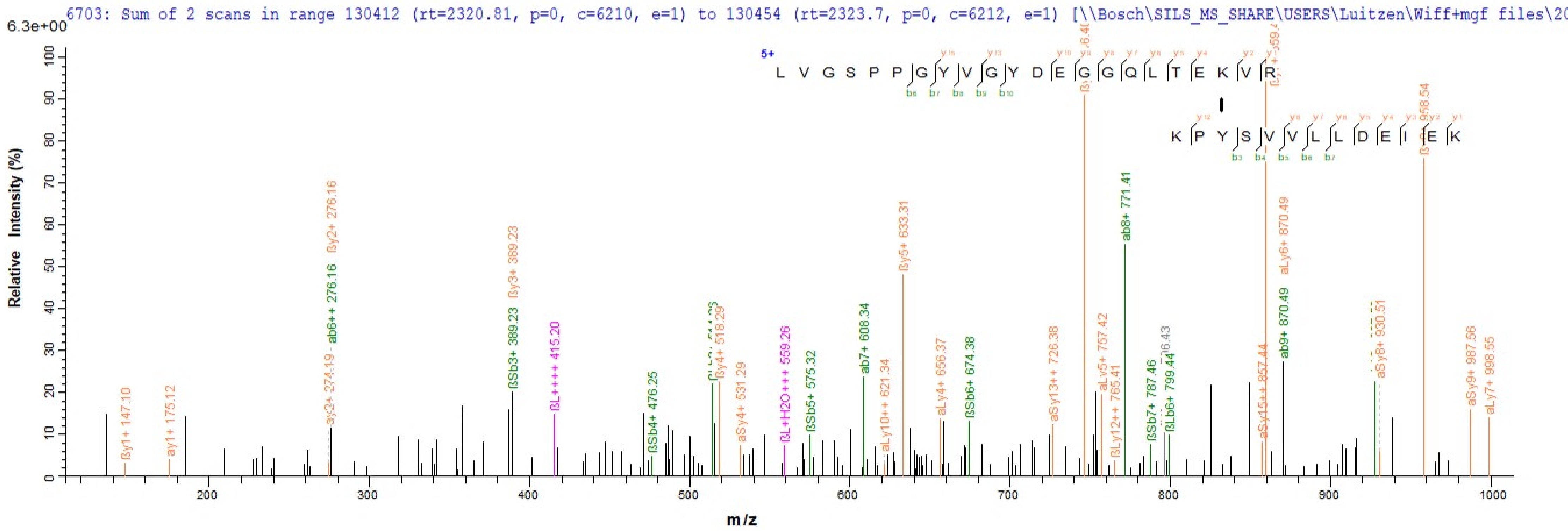
6703: Sum of 2 scans in range 130412 (rt=2320.81, p=0, c=6210, e=1) to 130454 (rt=2323.7, p=0, c=6212, e=1) [\\Bosch\SILS_MS_SHARE\USERS\Luitzen\Wiff+mgf files\20
6.3e+00
Relative Intensity (%)
m/z
5+
L V G S P P G Y V G Y D E G G Q L T E K V R
K P Y S V V L L D E I E K
ßy1+ 147.10
ay1+ 175.12
ay2+ 274.19
ßy2+ 276.16
ab6++ 276.16
ßSb3+ 389.23
ßy3+ 389.23
ßL++++ 415.20
ßSb4+ 476.25
ßy4+ 518.29
aSy4+ 531.29
ßL+H2O+++ 559.26
ßSb5+ 575.32
ab7+ 608.34
aLy10++ 621.34
ßy5+ 633.31
aLy4+ 656.37
ßSb6+ 674.38
aSy13++ 726.38
aLy5+ 757.42
ßLy12++ 765.41
ab8+ 771.41
ßSb7+ 787.46
ßLb6+ 799.44
aSy15++ 857.44
aLy6+ 870.49
ab9+ 870.49
aSy8+ 930.51
aSy9+ 987.56
aLy7+ 998.55

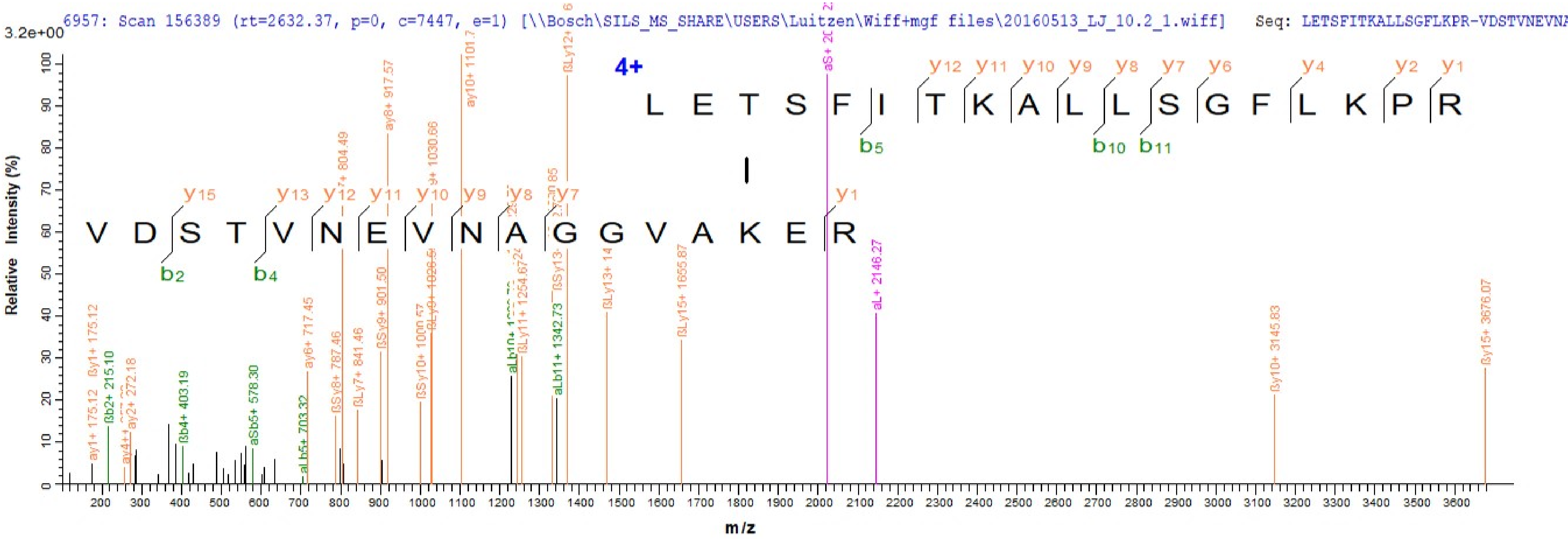
6957: Scan 156389 (rt=2632.37, p=0, c=7447, e=1) [\\Bosch\SILS_MS_SHARE\USERS\Luitzen\Wiff+mgf files\20160513_LJ_10.2_1.wiff] Seq: LETSFITKALLSGFLKPR-VDSTVNEVNA
3.2e+00
Relative Intensity (%)
m/z
4+
L E T S F I T K A L L S G F L K P R
V D S T V N E V N A G G V A K E R
y12 y11 y10 y9 y8 y7 y6 y4 y2 y1
b5 b10 b11
y15 y13 y12 y11 y10 y9 y8 y7 y1
b2 b4
ay1+ 175.12
ßy1+ 175.12
ßb2+ 215.10
ay2+ 272.18
ßb4+ 403.19
aSb5+ 578.30
aLb5+ 703.32
ay6+ 717.45
ßSy8+ 787.46
ßLy7+ 841.46
ßSy9+ 901.50
ay8+ 917.57
ßSy10+ 1000.57
y9+ 1030.66
ay10+ 1101.7
ßLy11+ 1254.67
aLb11+ 1342.73
ßLy13+ 14
ßLy15+ 1655.87
aL+ 2146.27
ßy10+ 3145.83
ßy15+ 3676.07

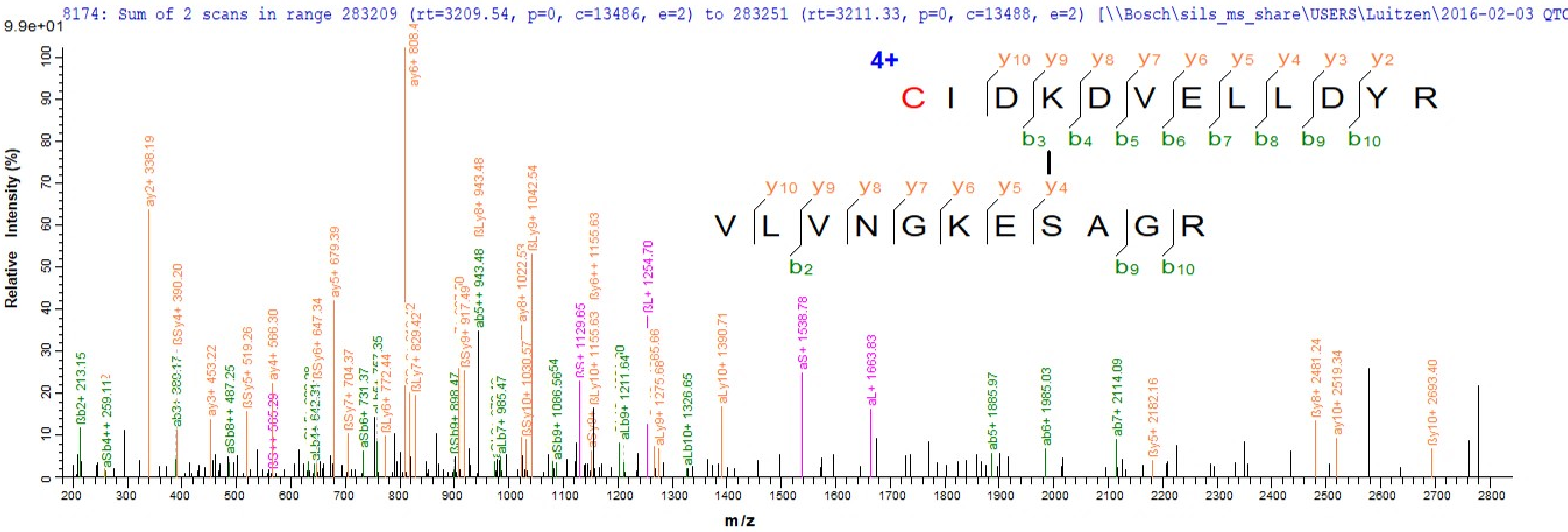
8174: Sum of 2 scans in range 283209 (rt=3209.54, p=0, c=13486, e=2) to 283251 (rt=3211.33, p=0, c=13488, e=2) [\\Bosch\sils_ms_share\USERS\Luitzen\2016-02-03 QTC
9.9e+01
Relative Intensity (%)
m/z
4+
C I D K D V E L L D Y R
y10 y9 y8 y7 y6 y5 y4 y3 y2
b3 b4 b5 b6 b7 b8 b9 b10
V L V N G K E S A G R
y10 y9 y8 y7 y6 y5 y4
b2 b9 b10
ay6+ 808.4
ay2+ 338.19
ßLy8+ 943.48
ßLy9+ 1042.54
ay5+ 679.39
ßSy4+ 390.20
ab5++ 943.48
ay8+ 1022.53
ßy6++ 1155.63
ßL+ 1254.70
aS+ 1538.78
aL+ 1663.83
aLy10+ 1390.71
ßb2+ 213.15
aSb4++ 259.11
ay3+ 453.22
aSb8++ 487.25
ßSy5+ 519.26
ay4+ 566.30
ßSy6+ 647.34
aLb4+ 642.31
ßSy7+ 704.37
aSb6+ 731.37
ßLy6+ 772.44
ßLy7+ 829.42
ßSb9+ 898.47
ßSy9+ 917.49
aLb7+ 985.47
ßSy10+ 1030.57
aSb9+ 1086.56
ßS+ 1129.65
ßLy10+ 1155.63
aLb9+ 1211.64
aLy9+ 1275.68
aLb10+ 1326.65
ab5+ 1885.97
ab6+ 1985.03
ab7+ 2114.09
ßy5+ 2182.16
ßy8+ 2481.24
ay10+ 2519.34
ßy10+ 2693.40

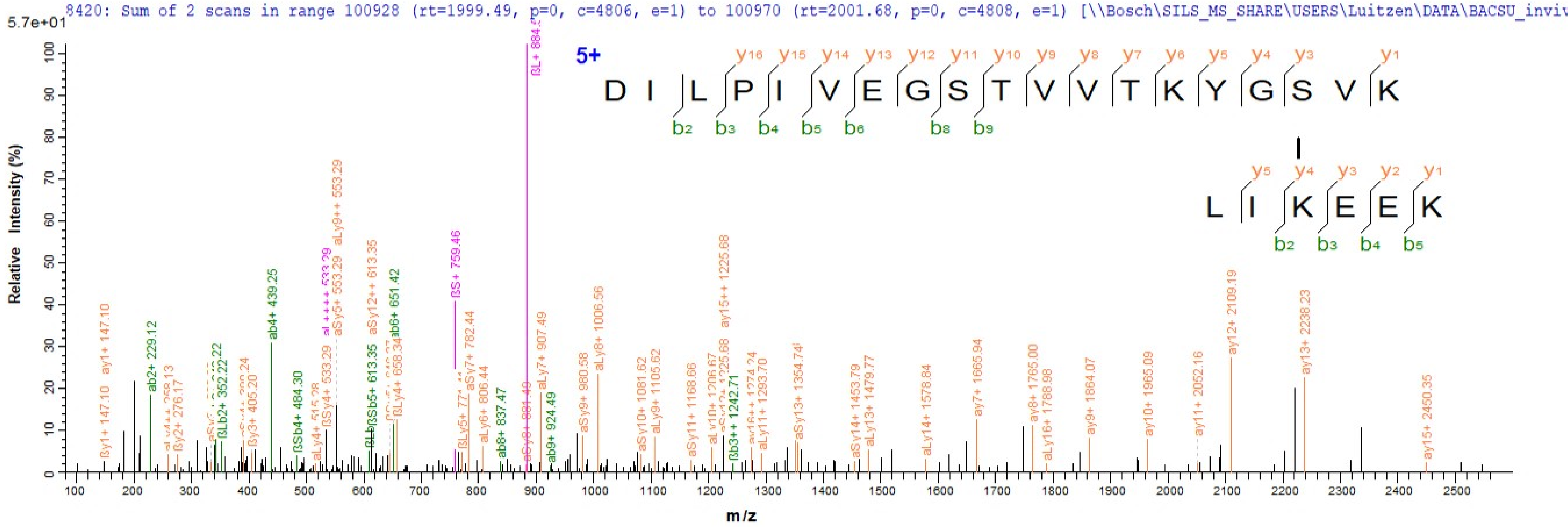
8420: Sum of 2 scans in range 100928 (rt=1999.49, p=0, c=4806, e=1) to 100970 (rt=2001.68, p=0, c=4808, e=1) [\\Bosch\SILS_MS_SHARE\USERS\Luitzen\DATA\BACSU_inviv
5.7e+01
Relative Intensity (%)
m/z
5+
D I L P I V E G S T V V T K Y G S V K
L I K E E K
y16 y15 y14 y13 y12 y11 y10 y9 y8 y7 y6 y5 y4 y3 y1
b2 b3 b4 b5 b6 b8 b9
y5 y4 y3 y2 y1
b2 b3 b4 b5
ßL+ 884.5
ay1+ 147.10
ab2+ 229.12
ab4+ 439.25
aLy9++ 553.29
aSy12++ 613.35
ab6+ 651.42
ßS+ 759.46
aSy7+ 782.44
aLy7+ 907.49
aLy8+ 1006.56
ay15++ 1225.68
ay12+ 2109.19
ay13+ 2238.23
ay15+ 2450.35

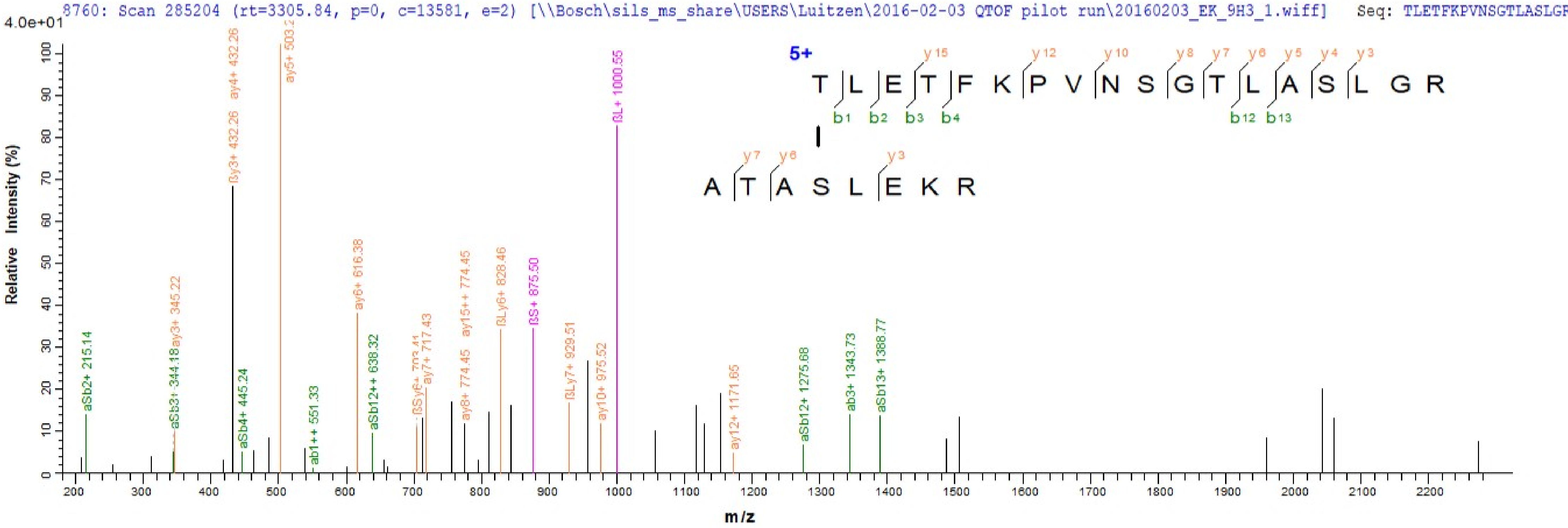
8760: Scan 285204 (rt=3305.84, p=0, c=13581, e=2) [\\Bosch\sils_ms_share\USERS\Luitzen\2016-02-03 QTOF pilot run\20160203_EK_9H3_1.wiff] Seq: TLETFKPVNSGTLASLGR
4.0e+01
Relative Intensity (%)
m/z
5+
T L E T F K P V N S G T L A S L G R
y15 y12 y10 y8 y7 y6 y5 y4 y3
b1 b2 b3 b4 b12 b13
A T A S L E K R
y7 y6 y3
aSb2+ 215.14
aSb3+ 344.18
ay3+ 345.22
βy3+ 432.26
ay4+ 432.26
aSb4+ 445.24
ab1++ 551.33
ay6+ 616.38
aSb12++ 638.32
ay7+ 717.43
ay8+ 774.45
ay15++ 774.45
βLy6+ 828.46
βS+ 875.50
βLy7+ 929.51
ay10+ 975.52
βL+ 1000.55
ay12+ 1171.65
aSb12+ 1275.68
ab3+ 1343.73
aSb13+ 1388.77

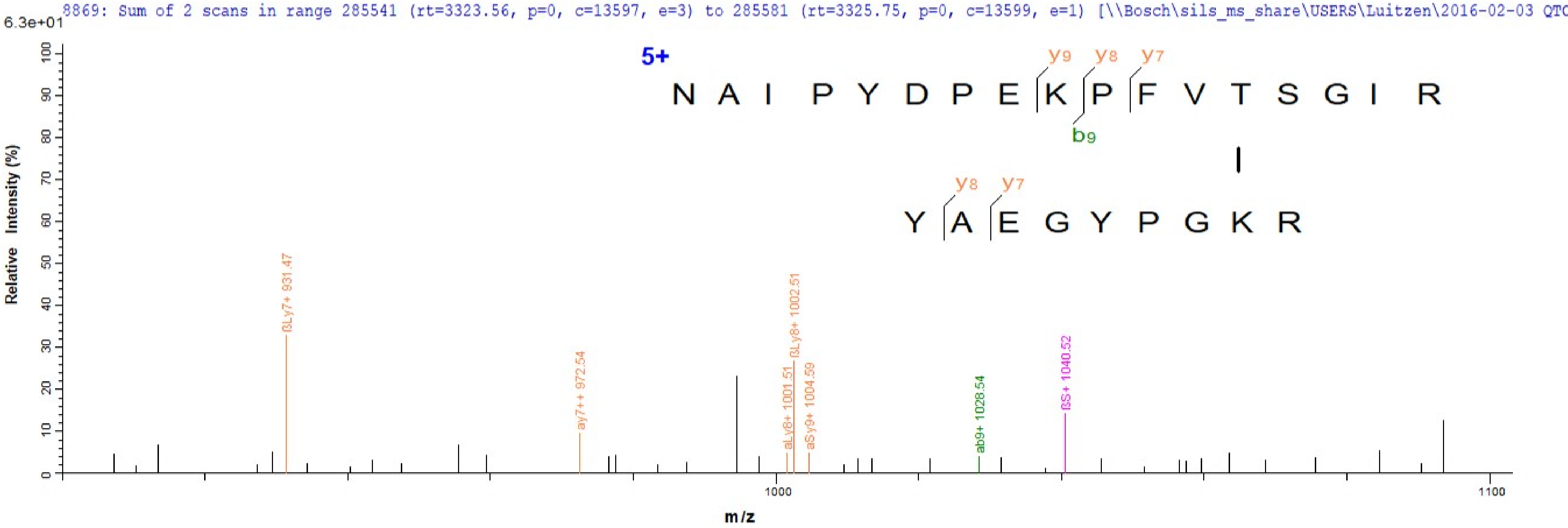
8869: Sum of 2 scans in range 285541 (rt=3323.56, p=0, c=13597, e=3) to 285581 (rt=3325.75, p=0, c=13599, e=1) [\\Bosch\sils_ms_share\USERS\Luitzen\2016-02-03 QTC
6.3e+01
5+
N A I P Y D P E K P F V T S G I R
y9 y8 y7
b9
Y A E G Y P G K R
y8 y7
ßLy7+ 931.47
ay7++ 972.54
aLy8+ 1001.51
ßLy8+ 1002.51
aSy9+ 1004.59
ab9+ 1028.54
ßS+ 1040.52
Relative Intensity (%)
m/z
1000
1100

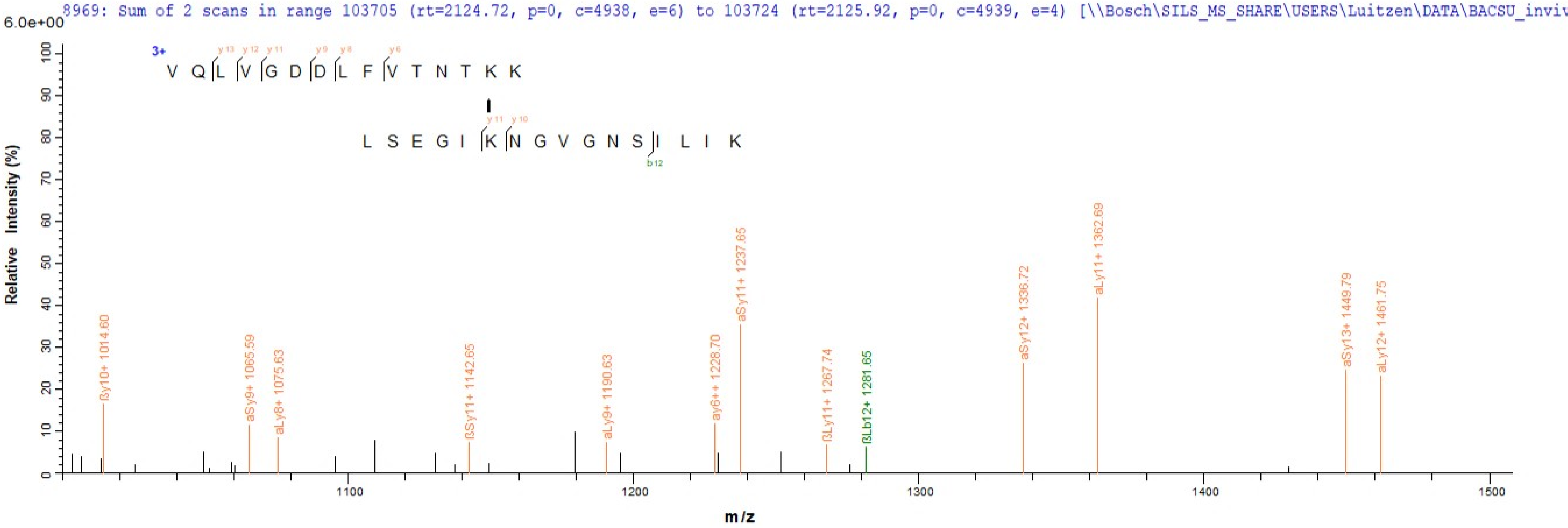
8969: Sum of 2 scans in range 103705 (rt=2124.72, p=0, c=4938, e=6) to 103724 (rt=2125.92, p=0, c=4939, e=4) [\\Bosch\SILS_MS_SHARE\USERS\Luitzen\DATA\BACSU_inviv
6.0e+00
3+
V Q L V G D D L F V T N T K K
L S E G I K N G V G N S I L I K
Relative Intensity (%)
m/z
ßy10+ 1014.60
aSy9+ 1065.59
aLy8+ 1075.63
ßSy11+ 1142.65
aLy9+ 1190.63
ay6++ 1228.70
aSy11+ 1237.65
ßLy11+ 1267.74
ßLb12+ 1281.65
aSy12+ 1336.72
aLy11+ 1362.69
aSy13+ 1449.79
aLy12+ 1461.75

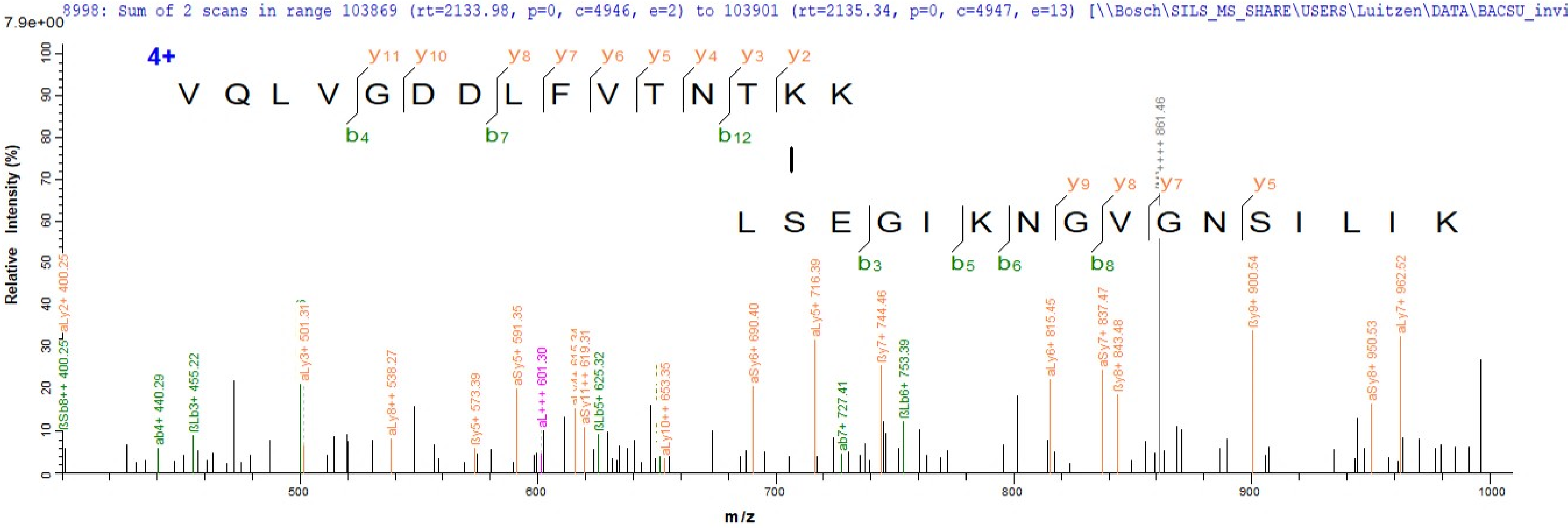
8998: Sum of 2 scans in range 103869 (rt=2133.98, p=0, c=4946, e=2) to 103901 (rt=2135.34, p=0, c=4947, e=13) [\\Bosch\SILS_MS_SHARE\USERS\Luitzen\DATA\BACSU_invi
7.9e+00
4+
V Q L V G D D L F V T N T K K
y11 y10 y8 y7 y6 y5 y4 y3 y2
b4 b7 b12
L S E G I K N G V G N S I L I K
y9 y8 y7 y5
b3 b5 b6 b8
Relative Intensity (%)
m/z
ßSb8++ 400.25
aLy2+ 400.25
ab4+ 440.29
ßLb3+ 455.22
aLy3+ 501.31
aLy8++ 538.27
ßy5+ 573.39
aSy5+ 591.35
aL+++ 601.30
aSy11++ 619.31
ßLb5+ 625.32
aLy10++ 653.35
aSy6+ 690.40
aLy5+ 716.39
ab7+ 727.41
ßy7+ 744.46
ßLb6+ 753.39
aLy6+ 815.45
aSy7+ 837.47
ßy8+ 843.48
861.46
ßy9+ 900.54
aSy8+ 950.53
aLy7+ 962.52

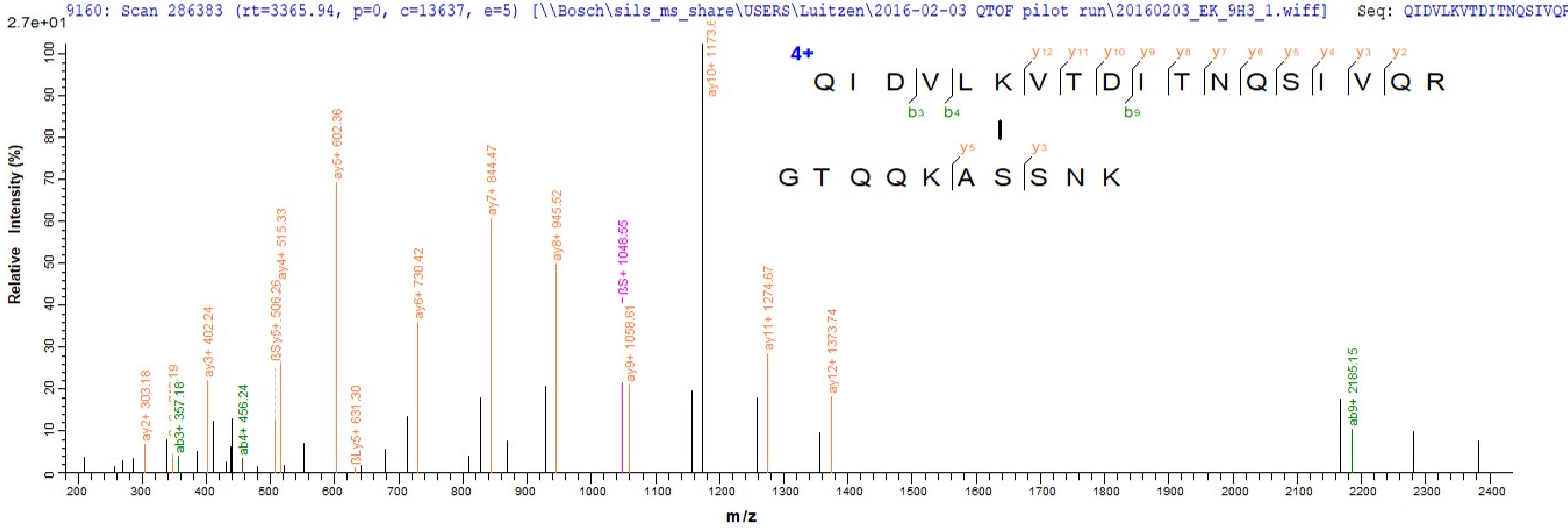
9160: Scan 286383 (rt=3365.94, p=0, c=13637, e=5) [\\Bosch\sils_ms_share\USERS\Luitzen\2016-02-03 QTOF pilot run\20160203_EK_9H3_1.wiff] Seq: QIDVLKVTDITNQSIVQR
2.7e+01
Relative Intensity (%)
m/z
4+
QIDVLKVTDITNQSIVQR
GTQQKASSNK
y12 y11 y10 y9 y8 y7 y6 y5 y4 y3 y2
b3 b4 b9
y5 y3
ay2+ 303.18
ab3+ 357.18
ay3+ 402.24
ab4+ 456.24
ßSy5+ 506.26
ay4+ 515.33
ay5+ 602.36
ßLy5+ 631.30
ay6+ 730.42
ay7+ 844.47
ay8+ 945.52
ßS+ 1048.55
ay9+ 1058.61
ay10+ 1173.6
ay11+ 1274.67
ay12+ 1373.74
ab9+ 2185.15

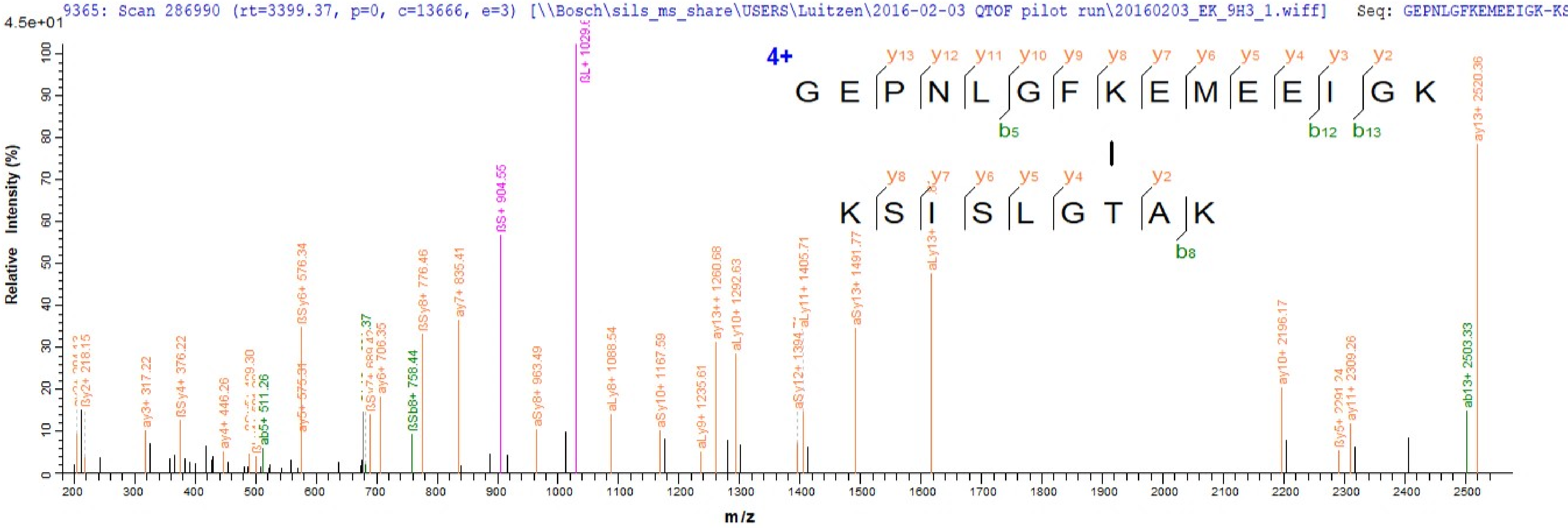

9365: Scan 286990 (rt=3399.37, p=0, c=13666, e=3) [\\Bosch\sils_ms_share\USERS\Luitzen\2016-02-03 QTOF pilot run\20160203_EK_9H3_1.wiff] Seq: GEPNLGFKEMEEIGK-KS
4.5e+01
4+
G E P N L G F K E M E E I G K
K S I S L G T A K
Relative Intensity (%)
m/z
ßL+ 1029.6
ßS+ 904.55
ay13+ 2520.36
ab13+ 2503.33
ay10+ 2196.17
ay11+ 2309.26
ßy5+ 2291.24
aLy13+
aSy13+ 1491.77
aLy11+ 1405.71
aSy12+ 1394.7
aLy10+ 1292.63
ay13++ 1260.68
aLy9+ 1235.61
aSy10+ 1167.59
aLy8+ 1088.54
aSy8+ 963.49
ay7+ 835.41
ßSy8+ 776.46
ßSb8+ 758.44
ay6+ 706.35
ßSy6+ 576.34
ay5+ 575.31
ab5+ 511.26
ay4+ 446.26
ßSy4+ 376.22
ay3+ 317.22
ßy2+ 218.15

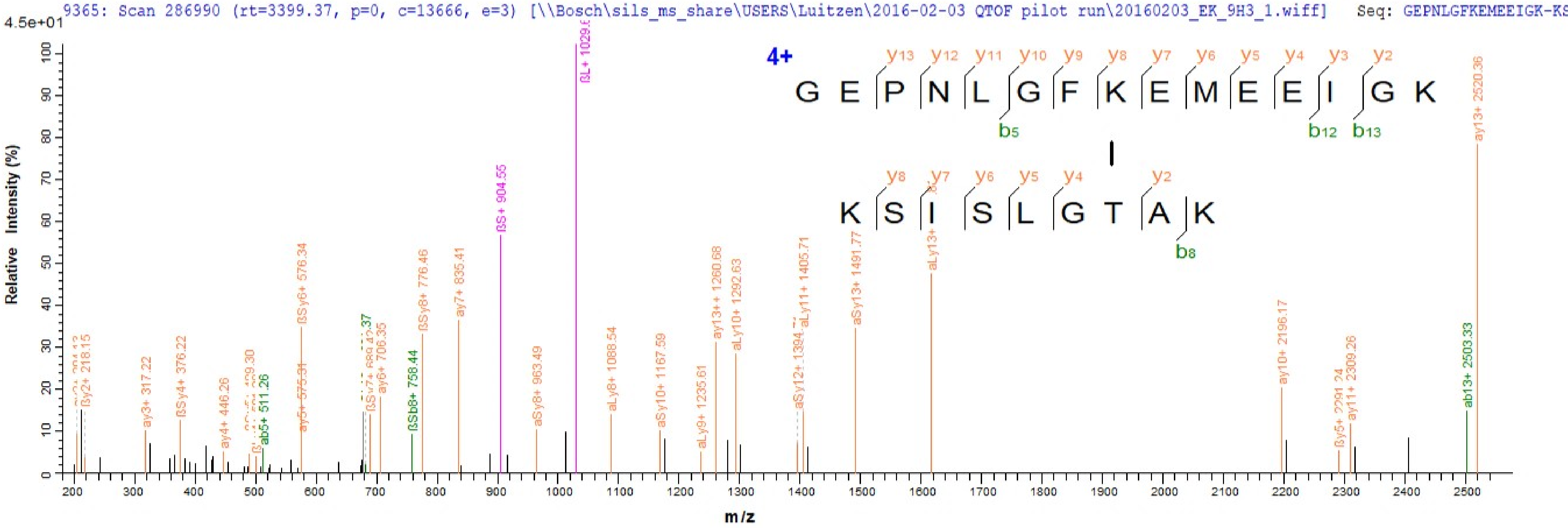
9365: Scan 286990 (rt=3399.37, p=0, c=13666, e=3) [\\Bosch\sils_ms_share\USERS\Luitzen\2016-02-03 QTOF pilot run\20160203_EK_9H3_1.wiff] Seq: GEPNLGFKEMEEIGK-KS
4.5e+01
Relative Intensity (%)
m/z
4+
G E P N L G F K E M E E I G K
y13 y12 y11 y10 y9 y8 y7 y6 y5 y4 y3 y2
b5 b12 b13
K S I S L G T A K
y8 y7 y6 y5 y4 y2
b8
ßL+ 1029.6
ßS+ 904.55
ay13+ 2520.36
ßy2+ 218.15
ay3+ 317.22
ßSy4+ 376.22
ay4+ 446.26
ab5+ 511.26
ay5+ 575.31
ßSy6+ 576.34
ay6+ 706.35
ßSb8+ 758.44
ßSy8+ 776.46
ay7+ 835.41
aSy8+ 963.49
aLy8+ 1088.54
aSy10+ 1167.59
aLy9+ 1235.61
ay13++ 1260.68
aLy10+ 1292.63
aLy11+ 1405.71
aSy13+ 1491.77
aLy13+
ay10+ 2196.17
ßy5+ 2291.24
ay11+ 2309.26
ab13+ 2503.33

10500: Scan 293312 (rt=3656.31, p=0, c=13967, e=4) [\\Bosch\sils_ms_share\USERS\Luitzen\2016-02-03 QTOF pilot run\20160203_EK_9H3_1.wiff] Seq: EAYQDDYSKTLEPGSDF

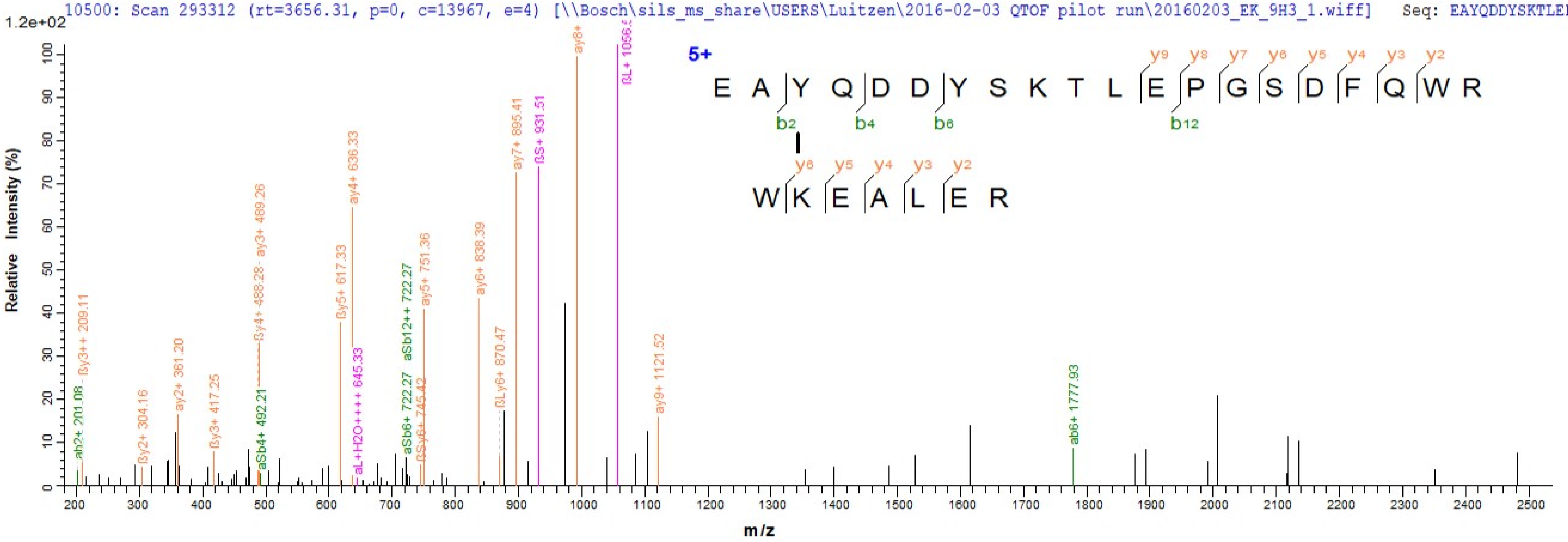

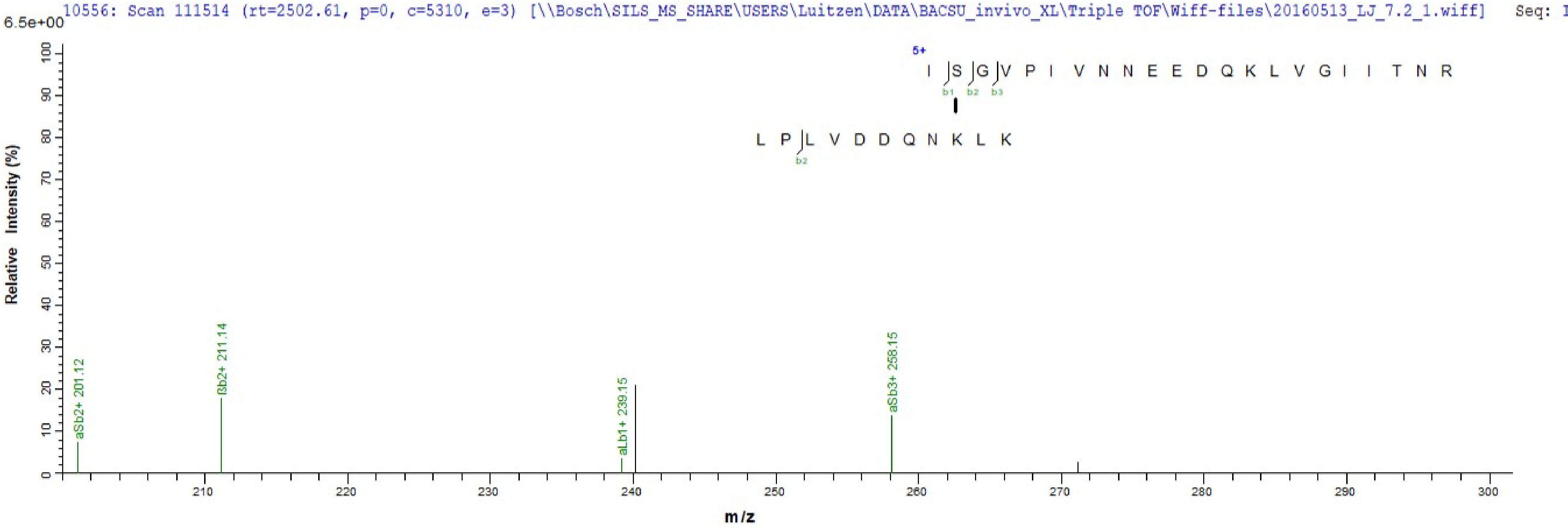
10556: Scan 111514 (rt=2502.61, p=0, c=5310, e=3) [\\Bosch\SILS_MS_SHARE\USERS\Luitzen\DATA\BACSU_invivo_XL\Triple TOF\Wiff-files\20160513_LJ_7.2_1.wiff] Seq:
6.5e+00
Relative Intensity (%)
5+
I S G V P I V N N E E D Q K L V G I I T N R
b1 b2 b3
L P L V D D Q N K L K
b2
aSb2+ 201.12
ßb2+ 211.14
aLb1+ 239.15
aSb3+ 258.15
m/z

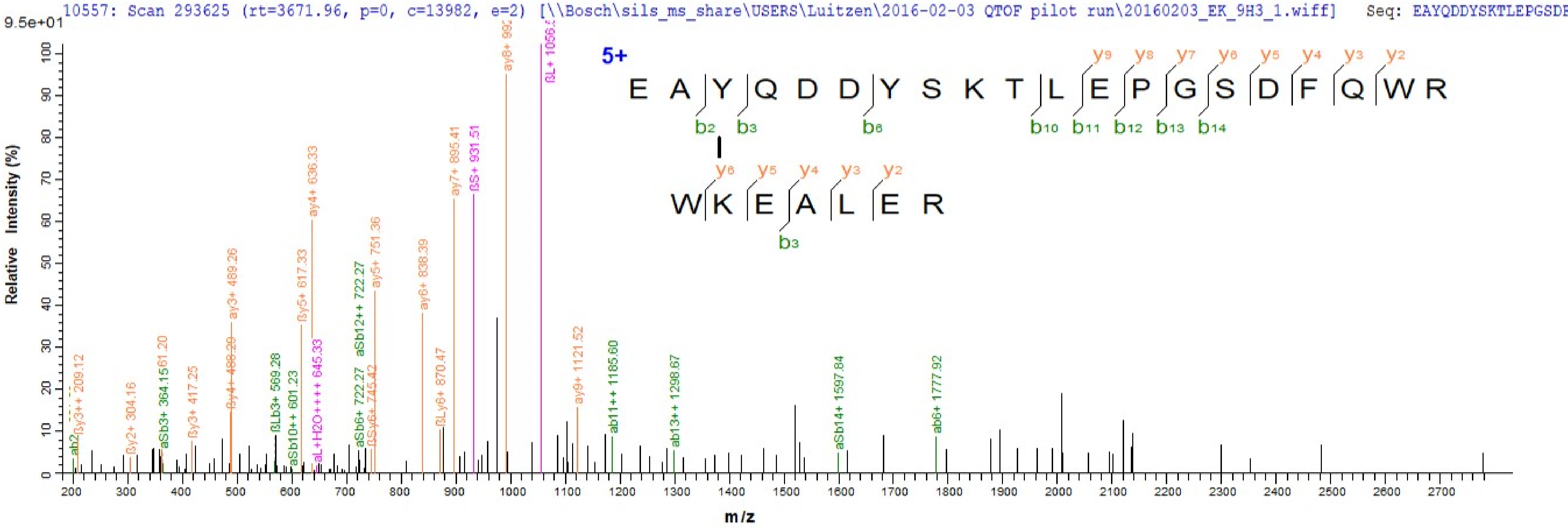
10557: Scan 293625 (rt=3671.96, p=0, c=13982, e=2) [\\Bosch\sils_ms_share\USERS\Luitzen\2016-02-03 QTOF pilot run\20160203_EK_9H3_1.wiff] Seq: EAYQDDYSKTLEPGSDF
9.5e+01
Relative Intensity (%)
m/z
5+
E A Y Q D D Y S K T L E P G S D F Q W R
y9 y8 y7 y6 y5 y4 y3 y2
b2 b3 b6 b10 b11 b12 b13 b14
W K E A L E R
y6 y5 y4 y3 y2
b3
ab2
ßy3++ 209.12
ßy2+ 304.16
aSb3+ 364.15
361.20
ßy3+ 417.25
ßy4+ 488.29
ay3+ 489.26
ßLb3+ 569.28
aSb10++ 601.23
ßy5+ 617.33
ay4+ 636.33
aL+H2O++++ 645.33
aSb6+ 722.27
aSb12++ 722.27
ßSy6+ 745.42
ay5+ 751.36
ay6+ 838.39
ßLy6+ 870.47
ay7+ 895.41
ßS+ 931.51
ay8+ 992
ßL+ 1056.
ay9+ 1121.52
ab11++ 1185.60
ab13++ 1298.67
aSb14+ 1597.84
ab6+ 1777.92

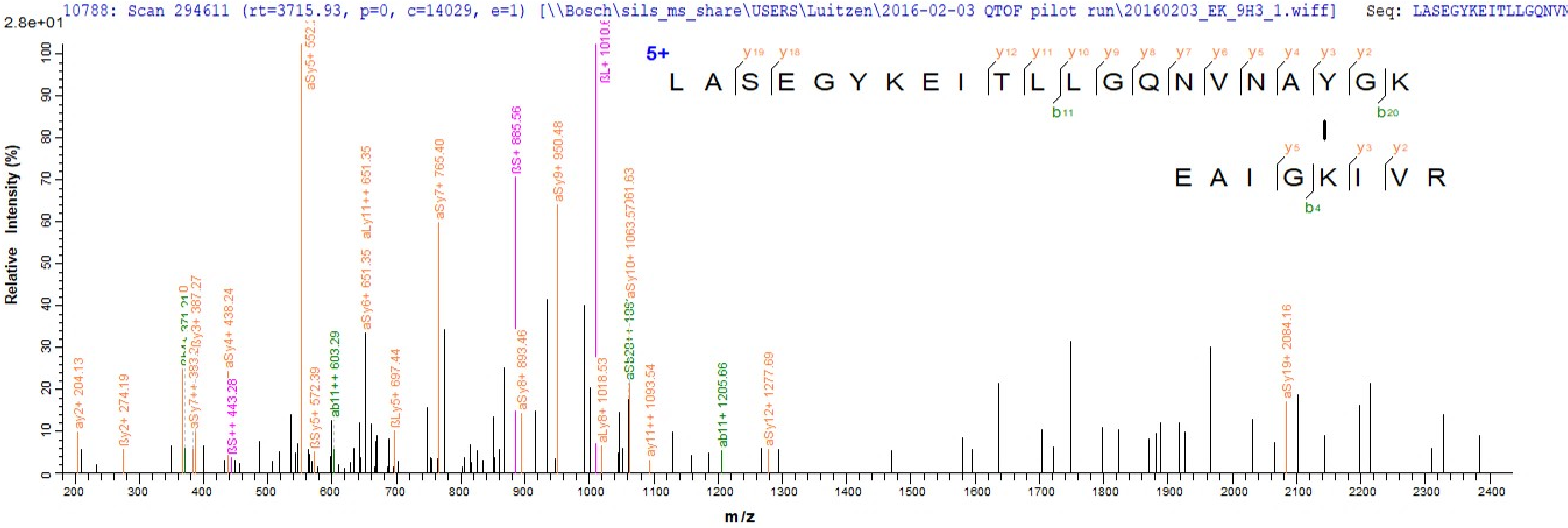
10788: Scan 294611 (rt=3715.93, p=0, c=14029, e=1) [\\Bosch\sils_ms_share\USERS\Luitzen\2016-02-03 QTOF pilot run\20160203_EK_9H3_1.wiff] Seq: LASEGYKEITLLGQNVN
2.8e+01
Relative Intensity (%)
m/z
5+
L A S E G Y K E I T L L G Q N V N A Y G K
y19 y18 y12 y11 y10 y9 y8 y7 y6 y5 y4 y3 y2
b11 b20
E A I G K I V R
y5 y3 y2
b4
ay2+ 204.13
βy2+ 274.19
aSy7++ 383.2
βy3+ 387.27
aSy4+ 438.24
βS++ 443.28
aSy5+ 552.
βSy5+ 572.39
ab11++ 603.29
aSy6+ 651.35
aLy11++ 651.35
βLy5+ 697.44
aSy7+ 765.40
βS+ 885.56
aSy8+ 893.46
aSy9+ 950.48
βL+ 1010.
aLy8+ 1018.53
aSy10+ 1063.57
ay11++ 1093.54
ab11+ 1205.66
aSy12+ 1277.69
aSy19+ 2084.16

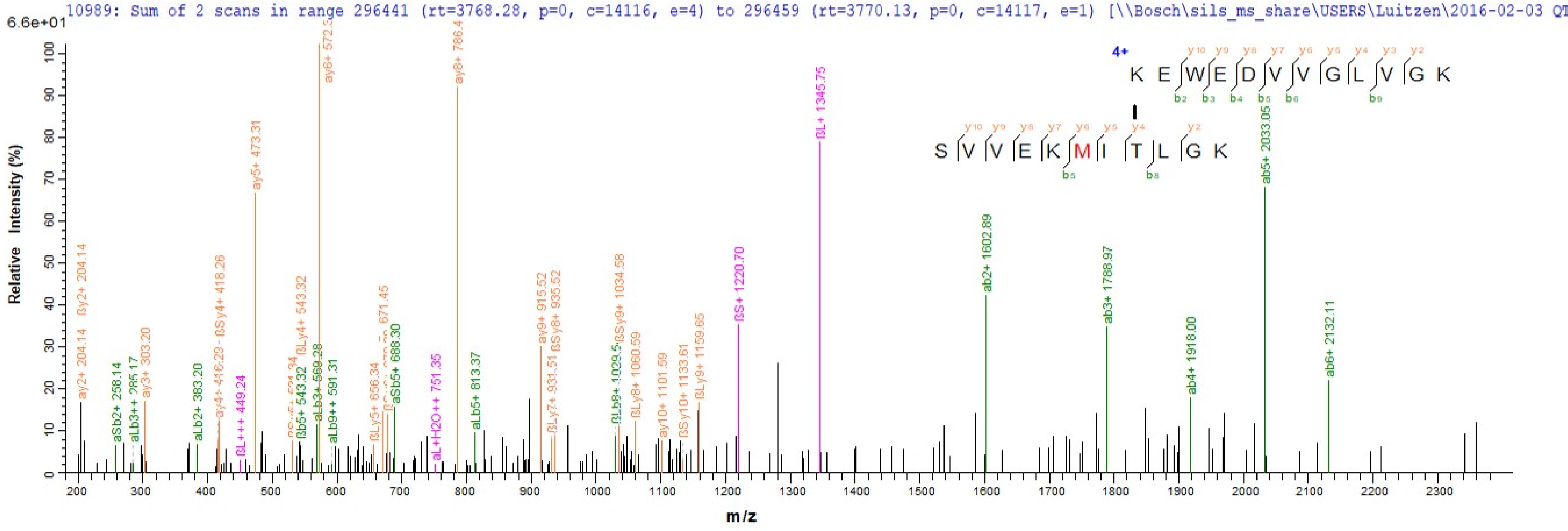
10989: Sum of 2 scans in range 296441 (rt=3768.28, p=0, c=14116, e=4) to 296459 (rt=3770.13, p=0, c=14117, e=1) [\\Bosch\sils_ms_share\USERS\Luitzen\2016-02-03 QT
6.6e+01
Relative Intensity (%)
m/z
4+
K E W E D V V G L V G K
S V V E K M I T L G K
ay6+ 572.3
ay8+ 786.4
ßL+ 1345.75
ab5+ 2033.05
ay5+ 473.31
ab2+ 1602.89
ßS+ 1220.70
ab3+ 1788.97
ab6+ 2132.11
ab4+ 1918.00
ay2+ 204.14
ßy2+ 204.14
ay3+ 303.20
aSb2+ 258.14
aLb2+ 383.20
ßL+++ 449.24
ßLy4+ 543.32
ßb5+ 543.32
aLb9++ 591.31
aSb5+ 688.30
aL+H2O++ 751.35
aLb5+ 813.37
ay9+ 915.52
ßSy8+ 935.52
ßSy9+ 1034.58
ßLy8+ 1060.59
ay10+ 1101.59
ßSy10+ 1133.61
ßLy9+ 1159.65

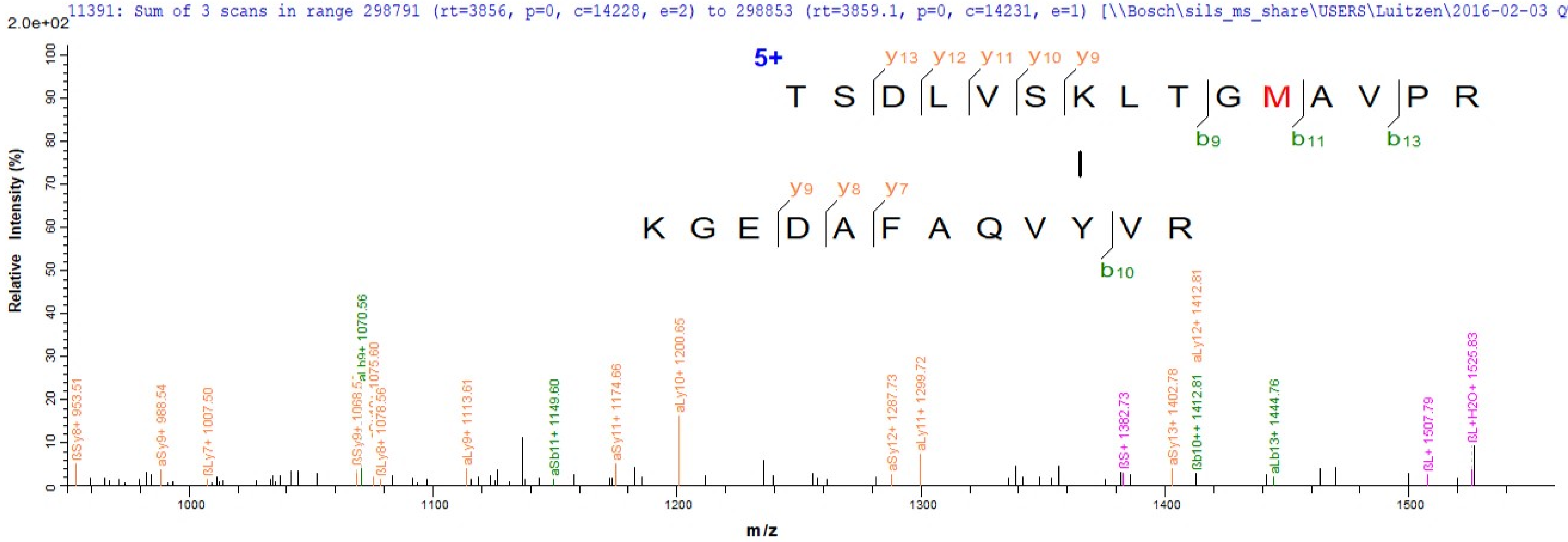

11391: Sum of 3 scans in range 298791 (rt=3856, p=0, c=14228, e=2) to 298853 (rt=3859.1, p=0, c=14231, e=1) [\\Bosch\sils_ms_share\USERS\Luitzen\2016-02-03 QTOF p
2.0e+02
5+
T S D L V S K L T G M A V P R
y13 y12 y11 y10 y9
b9 b11 b13
K G E D A F A Q V Y V R
y9 y8 y7
b10
Relative Intensity (%)
m/z
ßSy8+ 953.51
aSy9+ 988.54
ßLy7+ 1007.50
ßSy9+ 1068.5
aLb9+ 1070.56
ßLy8+ 1078.56
aLy9+ 1113.61
aSb11+ 1149.60
aSy11+ 1174.66
aLy10+ 1200.65
aSy12+ 1287.73
aLy11+ 1299.72
ßS+ 1382.73
aSy13+ 1402.78
ßb10++ 1412.81
aLy12+ 1412.81
aLb13+ 1444.76
ßL+ 1507.79
ßL+H2O+ 1525.83

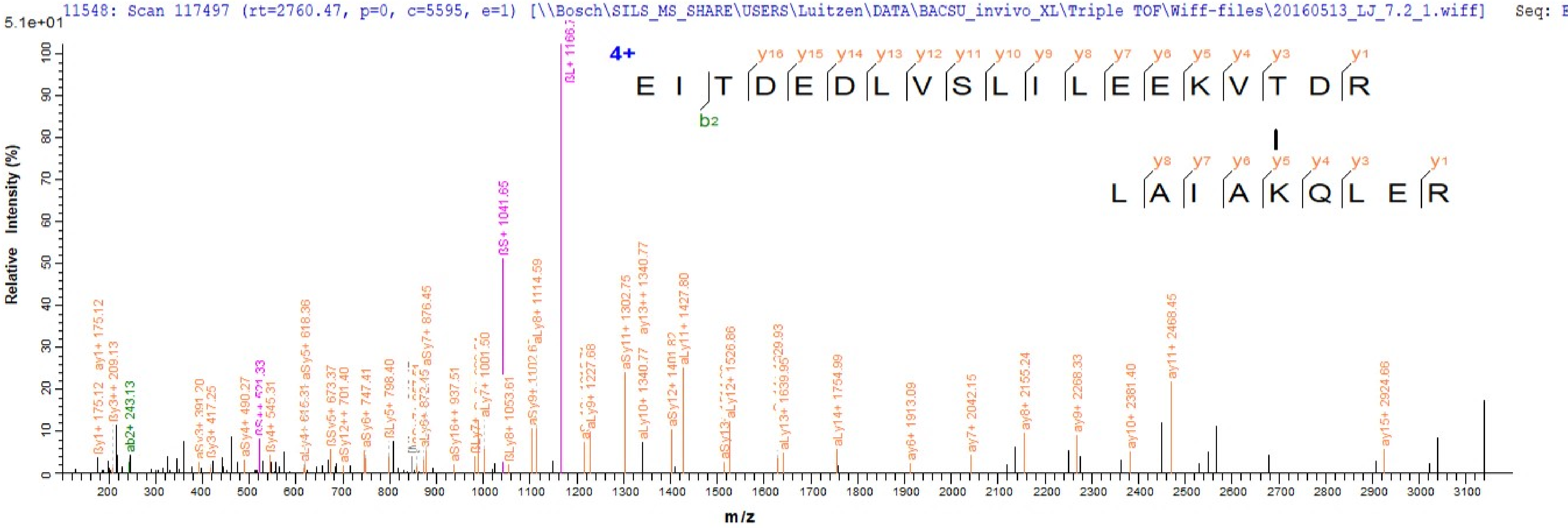
11548: Scan 117497 (rt=2760.47, p=0, c=5595, e=1) [\\Bosch\SILS_MS_SHARE\USERS\Luitzen\DATA\BACSU_invivo_XL\Triple TOF\Wiff-files\20160513_LJ_7.2_1.wiff] Seq: E
5.1e+01
4+
E I T D E D L V S L I L E E K V T D R
b2
L A I A K Q L E R
Relative Intensity (%)
m/z

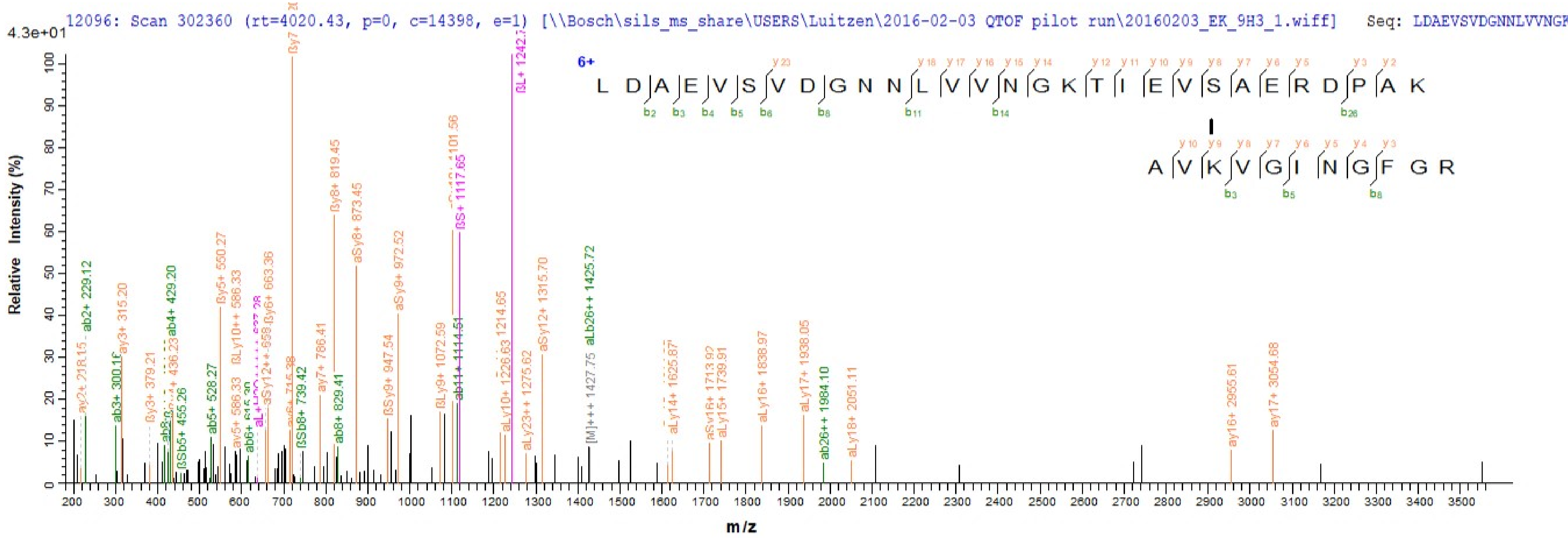

12096: Scan 302360 (rt=4020.43, p=0, c=14398, e=1) [\\Bosch\sils_ms_share\USERS\Luitzen\2016-02-03 QTOF pilot run\20160203_EK_9H3_1.wiff]
Seq: LDAEVSVDGNNLVVNGK
4.3e+01
Relative Intensity (%)
m/z
6+
L D A E V S V D G N N L V V N G K T I E V S A E R D P A K
A V K V G I N G F G R
ab2+ 229.12
ay3+ 315.20
ab4+ 429.20
ßy5+ 550.27
ßLy10++ 586.33
ßy6+ 663.36
ßy8+ 819.45
aSy8+ 873.45
aSy9+ 972.52
ßS+ 1117.65
ßL+ 1242.7
aSy12+ 1315.70
aLb26++ 1425.72
aLy16+ 1838.97
aLy17+ 1938.05
ay16+ 2955.61
ay17+ 3054.68

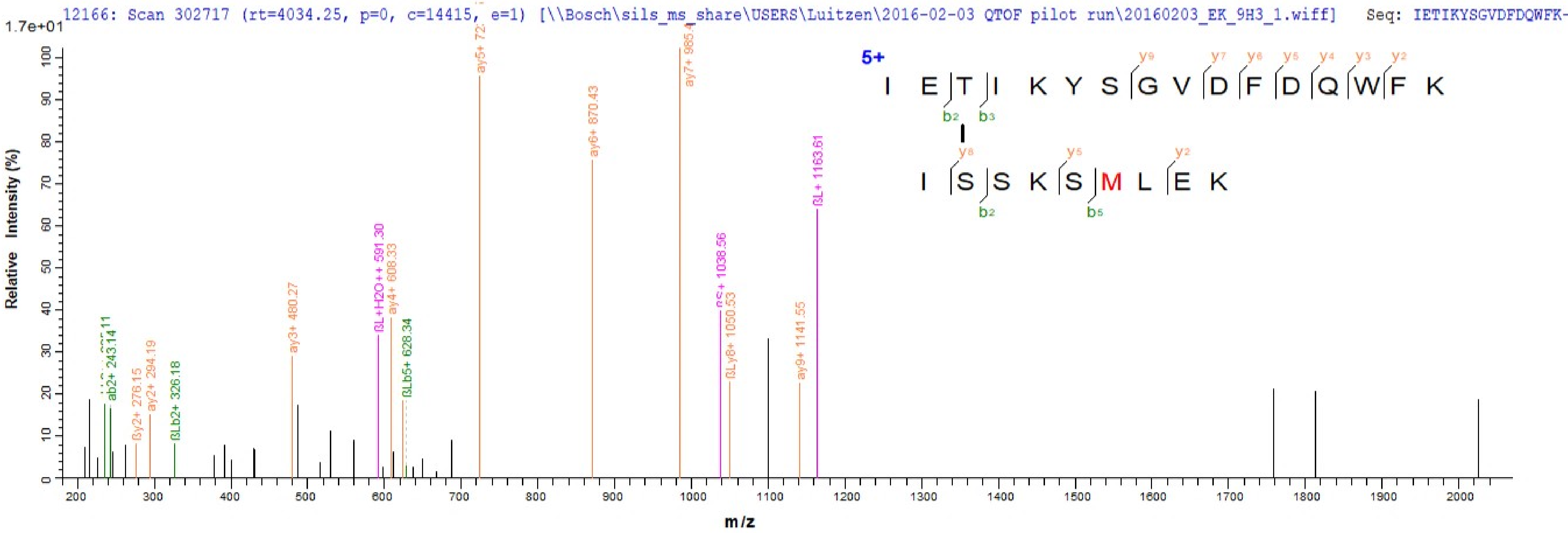

12166: Scan 302717 (rt=4034.25, p=0, c=14415, e=1) [\\Bosch\sils_ms_share\USERS\Luitzen\2016-02-03 QTOF pilot run\20160203_EK_9H3_1.wiff] Seq: IETIKYSGVDFDQWFK-
1.7e+01
Relative Intensity (%)
m/z
5+
I E T I K Y S G V D F D Q W F K
I S S K S M L E K
ab2+ 243.14
ßy2+ 276.15
ay2+ 294.19
ßLb2+ 326.18
ay3+ 480.27
ßL+H2O++ 591.30
ay4+ 608.33
ßLb5+ 628.34
ay6+ 870.43
ßLy8+ 1050.53
ßS+ 1038.56
ay9+ 1141.55
ßL+ 1163.61

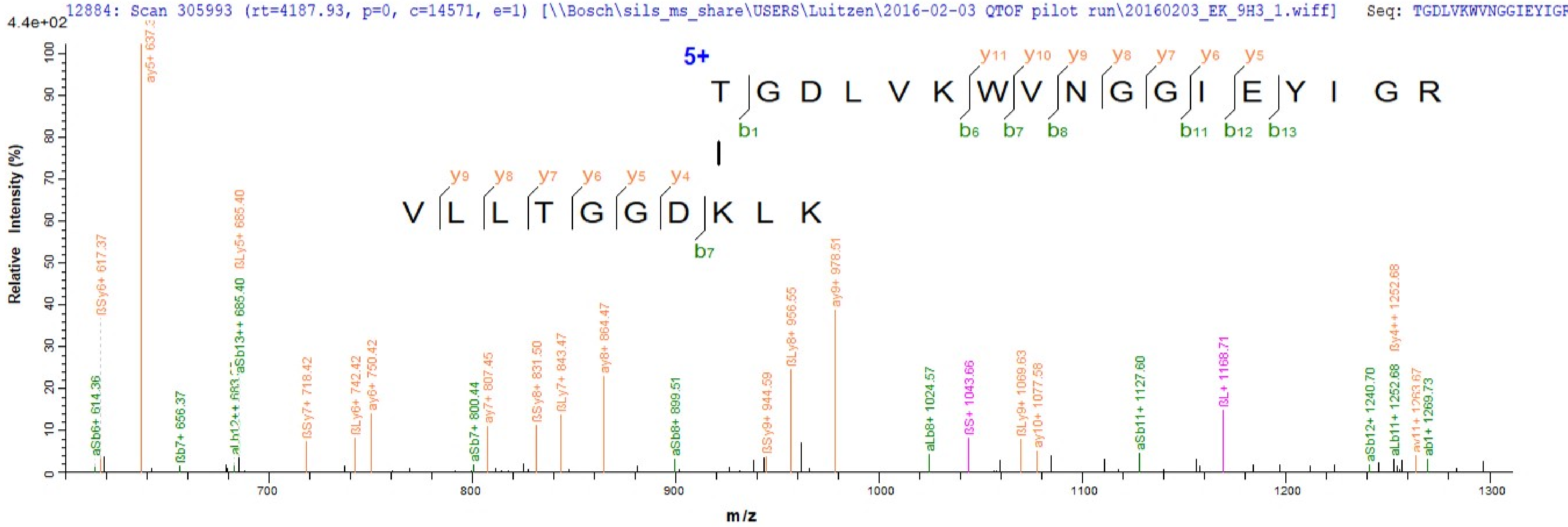
12884: Scan 305993 (rt=4187.93, p=0, c=14571, e=1) [\\Bosch\sils_ms_share\USERS\Luitzen\2016-02-03 QTOF pilot run\20160203_EK_9H3_1.wiff] Seq: TGDLVKWVNGGIEYIGR
4.4e+02
Relative Intensity (%)
m/z
5+
T G D L V K W V N G G I E Y I G R
y11 y10 y9 y8 y7 y6 y5
b1 b6 b7 b8 b11 b12 b13
V L L T G G D K L K
y9 y8 y7 y6 y5 y4
b7
aSb6+ 614.36
ßSy6+ 617.37
ßb7+ 656.37
aSb13++ 685.40
ßLy5+ 685.40
ßSy7+ 718.42
ßLy6+ 742.42
ay6+ 750.42
aSb7+ 800.44
ay7+ 807.45
ßSy8+ 831.50
ßLy7+ 843.47
ay8+ 864.47
aSb8+ 899.51
ßSy9+ 944.59
ßLy8+ 956.55
ay9+ 978.51
aLb8+ 1024.57
ßS+ 1043.66
ßLy9+ 1069.63
ay10+ 1077.58
aSb11+ 1127.60
ßL+ 1168.71
aSb12+ 1240.70
aLb11+ 1252.68
ßy4++ 1252.68
ab1+ 1269.73
700
800
900
1000
1100
1200
1300

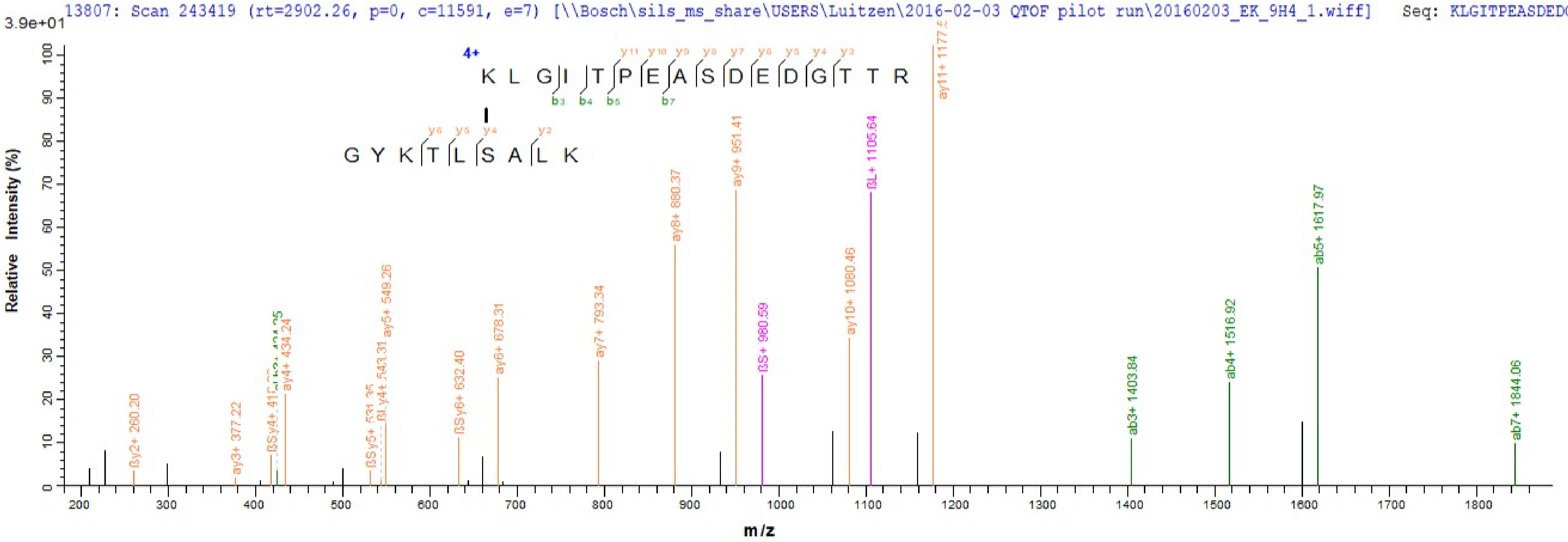

13807: Scan 243419 (rt=2902.26, p=0, c=11591, e=7) [\\Bosch\sils_ms_share\USERS\Luitzen\2016-02-03 QTOF pilot run\20160203_EK_9H4_1.wiff] Seq: KLGITPEASDEDGTTR-
3.9e+01
4+
K L G I T P E A S D E D G T T R
y11 y10 y9 y8 y7 y6 y5 y4 y3
b3 b4 b5 b7
G Y K T L S A L K
y6 y5 y4 y2
Relative Intensity (%)
m/z
ßy2+ 260.20
ay3+ 377.22
ay4+ 434.24
ßSy5+ 531.35
ßLy4+ 543.31
ay5+ 549.26
ßSy6+ 632.40
ay6+ 678.31
ay7+ 793.34
ay8+ 880.37
ay9+ 951.41
ßS+ 980.59
ay10+ 1080.46
ßL+ 1105.64
ay11+ 1177.5
ab3+ 1403.84
ab4+ 1516.92
ab5+ 1617.97
ab7+ 1844.06

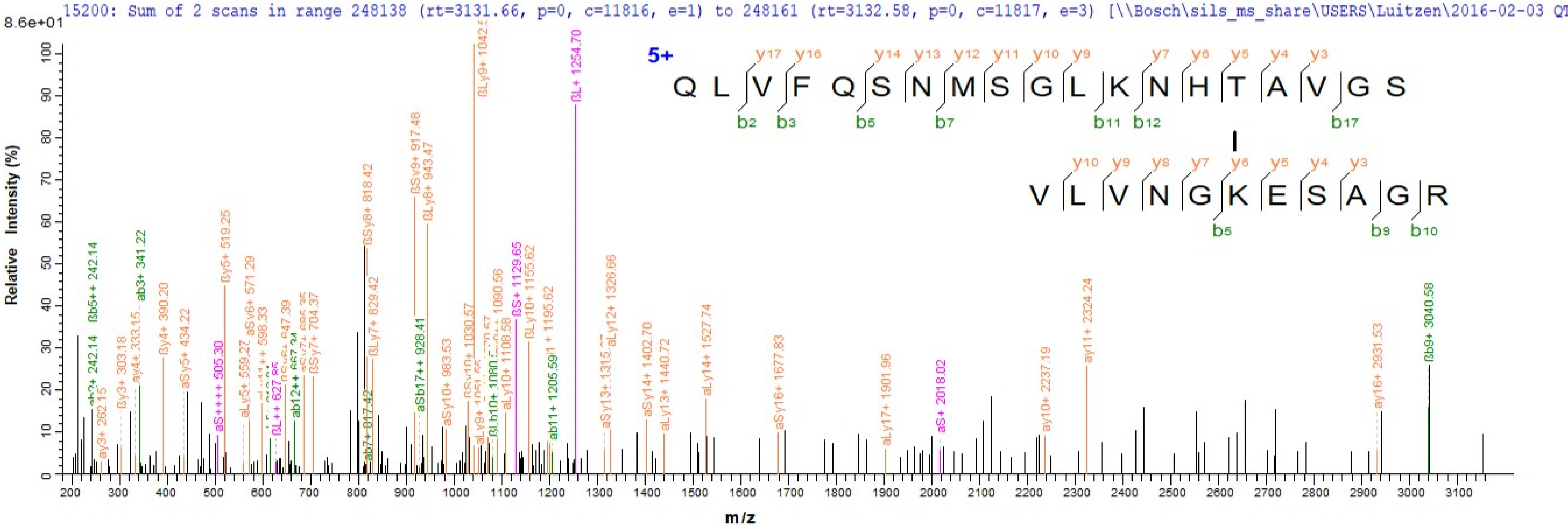
15200: Sum of 2 scans in range 248138 (rt=3131.66, p=0, c=11816, e=1) to 248161 (rt=3132.58, p=0, c=11817, e=3) [\\Bosch\sils_ms_share\USERS\Luitzen\2016-02-03 QT
8.6e+01
Relative Intensity (%)
m/z
5+
Q L V F Q S N M S G L K N H T A V G S
y17 y16 y14 y13 y12 y11 y10 y9 y7 y6 y5 y4 y3
b2 b3 b5 b7 b11 b12 b17
V L V N G K E S A G R
y10 y9 y8 y7 y6 y5 y4 y3
b5 b9 b10
ßLy9+ 1042.5
ßL+ 1254.70
ßSy8+ 818.42
ßSy9+ 917.48
ßLy8+ 943.47
ay11+ 2324.24
ßb9+ 3040.58

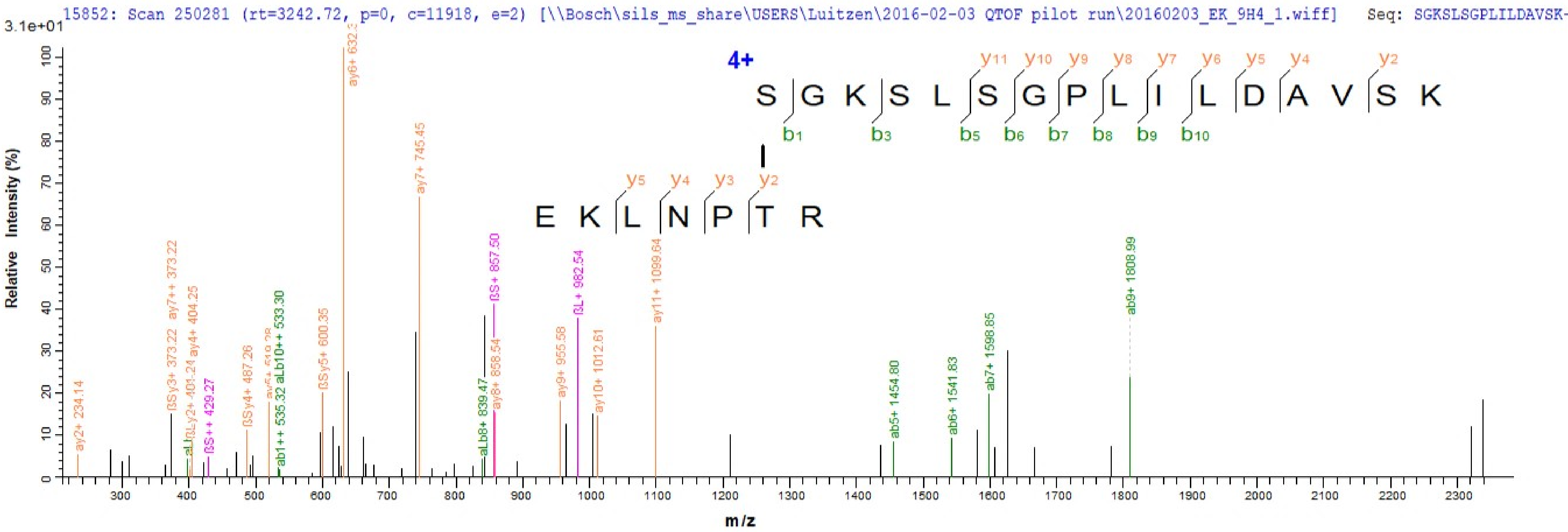
15852: Scan 250281 (rt=3242.72, p=0, c=11918, e=2) [\\Bosch\sils_ms_share\USERS\Luitzen\2016-02-03 QTOF pilot run\20160203_EK_9H4_1.wiff] Seq: SGKSLSGPLILDAVSK-
3.1e+01
Relative Intensity (%)
m/z
4+
S G K S L S G P L I L D A V S K
y11 y10 y9 y8 y7 y6 y5 y4 y2
b1 b3 b5 b6 b7 b8 b9 b10
E K L N P T R
y5 y4 y3 y2
ay2+ 234.14
ßSy3+ 373.22
ay7++ 373.22
ßLy2+ 401.24
ay4+ 404.25
ßS++ 429.27
ßSy4+ 487.26
ab1++ 535.32
aLb10++ 533.30
ßSy5+ 600.35
ay6+ 632.3
ay7+ 745.45
aLb8+ 839.47
ay8+ 858.54
ßS+ 857.50
ay9+ 955.58
ßL+ 982.54
ay10+ 1012.61
ay11+ 1099.64
ab5+ 1454.80
ab6+ 1541.83
ab7+ 1598.85
ab9+ 1808.99

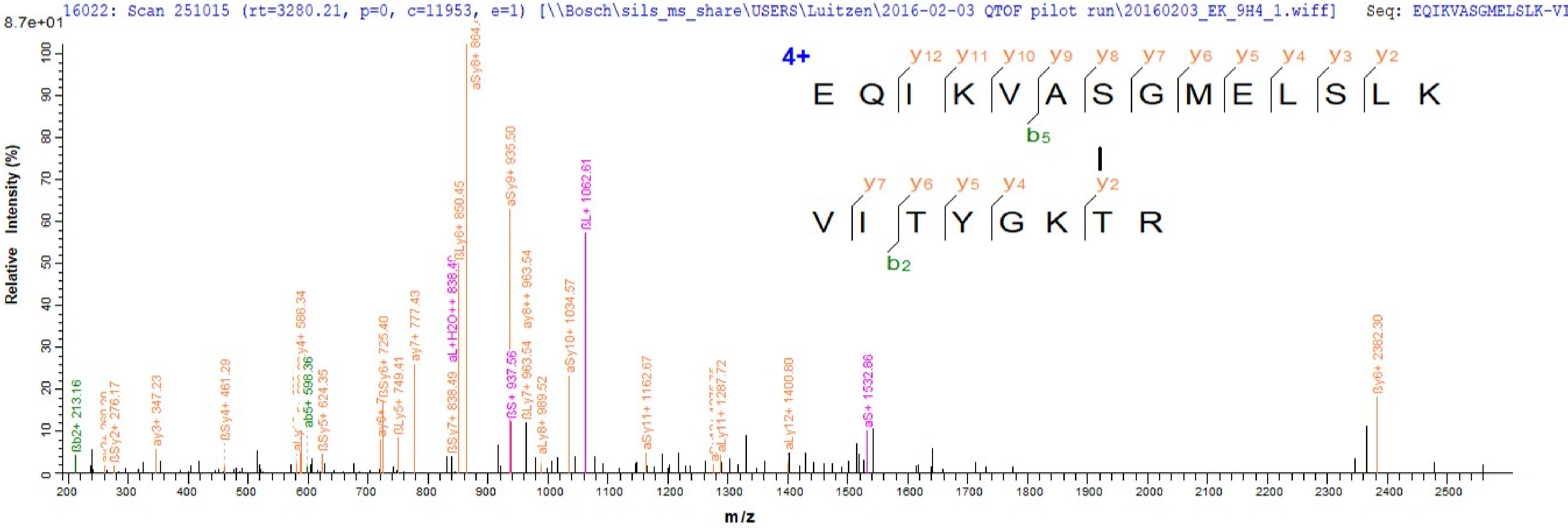
16022: Scan 251015 (rt=3280.21, p=0, c=11953, e=1) [\\Bosch\sils_ms_share\USERS\Luitzen\2016-02-03 QTOF pilot run\20160203_EK_9H4_1.wiff] Seq: EQIKVASGMELSLK-VI
8.7e+01
Relative Intensity (%)
m/z
4+
E Q I K V A S G M E L S L K
y12 y11 y10 y9 y8 y7 y6 y5 y4 y3 y2
b5
V I T Y G K T R
y7 y6 y5 y4 y2
b2
ßb2+ 213.16
ßSy2+ 276.17
ay3+ 347.23
ßSy4+ 461.29
ay4+ 586.34
ab5+ 598.36
ßSy5+ 624.35
ßSy6+ 725.40
ßLy5+ 749.41
ay7+ 777.43
ßSy7+ 838.49
aL+H2O++ 838.4
ßLy6+ 850.45
aSy8+ 864
aSy9+ 935.50
ßS+ 937.56
ßLy7+ 963.54
ay8++ 963.54
aLy8+ 989.52
aSy10+ 1034.57
ßL+ 1062.61
aSy11+ 1162.67
aLy11+ 1287.72
aLy12+ 1400.80
aS+ 1532.86
ßy6+ 2382.30

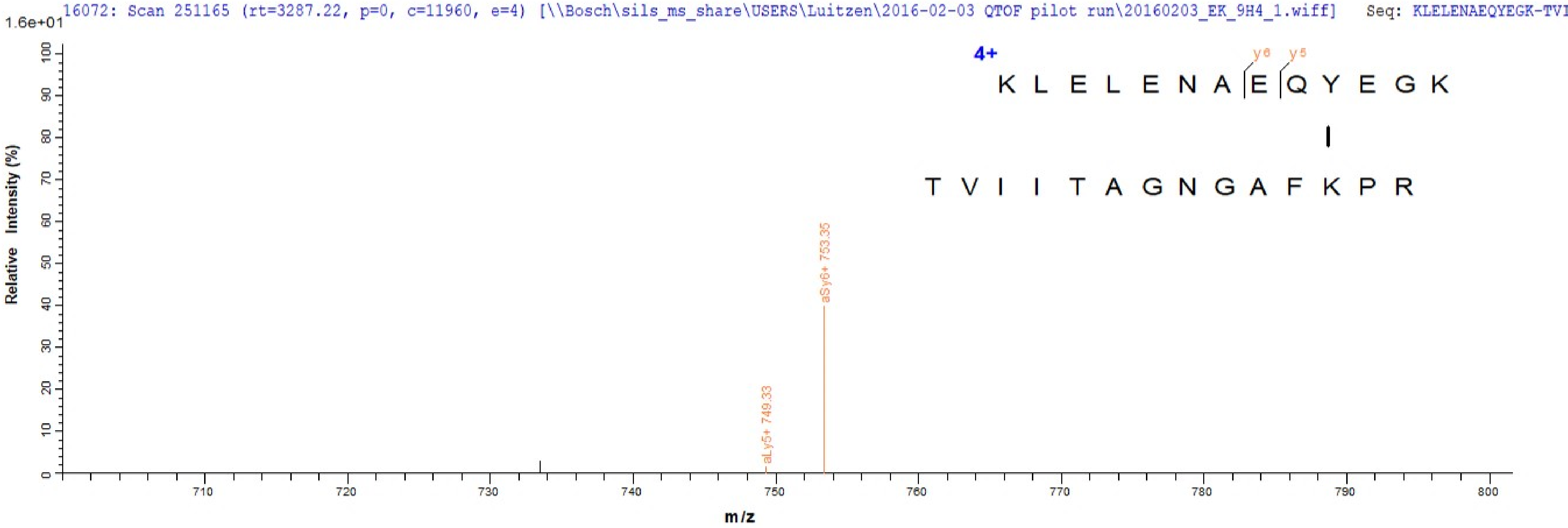
16072: Scan 251165 (rt=3287.22, p=0, c=11960, e=4) [\\Bosch\sils_ms_share\USERS\Luitzen\2016-02-03 QTOF pilot run\20160203_EK_9H4_1.wiff] Seq: KLELENAEQYEGK-TVI
1.6e+01
4+
y6 y5
K L E L E N A E Q Y E G K
T V I I T A G N G A F K P R
aLy5+ 749.33
aSy6+ 753.35
Relative Intensity (%)
m/z
0
10
20
30
40
50
60
70
80
90
100
710
720
730
740
750
760
770
780
790
800
